# Discovery, optimization, and cellular activities of 2-(aroylamino)cinnamamide derivatives against colon cancer cells

**DOI:** 10.1101/2019.12.15.876698

**Authors:** Abdelsattar M. Omar, Radwan S. Elhaggar, Martin K. Safo, Tamer M. Abdelghany, Mostafa H. Ahmed, Rio Boothello, Bhaumik B. Patel, Mohamed S. Abdel-Bakky, Moustafa E. El-Araby

## Abstract

Curcumin and *trans-*cinnamaldehyde are acrolein-based Michael acceptor compounds that are commonly found in domestic condiments, and known to cause cancer cell death *via* redox mechanisms. Based on the structural features of these compounds we designed and synthesized several 2-cinnamamido-N-substituted-cinnamamide (bis-cinnamamide) compounds. One of the derivatives, (*Z*)-2-[(*E*)-cinnamamido]-3-phenyl-N-propylacrylamide (**1512)** showed a moderate antiproliferative potency (HT116 cell line inhibition of 32.0 µM ± 2.6) with proven cellular activities leading to apoptosis. Importantly, **1512** exhibited good selectivity toxicity on cancer cells over noncancerous cells (IC_50_ of C-166 cell lines >100 µM), and low cancer cell resistance at 100 µM dose (growth rate 10.1±1.1%). We subsequently carried out structure activity relationship studies with **1512**. Derivatives with electron rich moiety at the aryl ring of the 2-aminocinnamaide moiety exhibited strong antiproliferative action while electron withdrawing groups caused loss of activity. Our most promising compound, **4112** [(*Z*)-3-(1*H*-indol-3-yl)-*N*-propyl-2-[(*E*)-3-(thien-2-yl)propenamido)propenamide] killed cancer cells at IC_50_ = 0.89 ± 0.04 µM (Caco-2), 2.85 ± 1.5 (HCT-116) and 1.65 ± 0.07 (HT-29), while exhibiting much weaker potency on C-166 and BHK normal cell lines (IC_50_ = 71 ± 5.12 and 77.6 ± 6.2 µM, respectively). Cellular studies towards identifying the compounds mechanism of cytotoxic activities revealed that apoptotic induction occurs in part due to oxidative stress. Importantly, the compounds showed inhibition of cancer stem cells that are critical for maintaining the potential for self-renewal and stemness. The results presented here show discovery of Michael addition compounds that potently kill cancer cells by a defined mechanism, with minimal effect on normal noncancerous cell.

## 1. INTRODUCTION

The Propenamide moiety is frequently recruited in designing drug-like compounds. For instance, Afatinib (**Figure 1**) a target-specific propenamide that selectively inhibits mutated HER2 kinase[1] has been recently approved by the FDA for the treatment of non-small cell lung cancer.[2] Other examples include ibrutinib (BTK inhibitor) and neratinib (HER-2 inhibitor) that have been developed for B-cell cancers and solid tumors, respectively.[3, 4] Mechanistically, the MAs cause cancer cell apoptosis by increasing the oxidative stress inside these cells. The cellular prooxidant induction by MAs is attributed to ligation of SH groups of certain targets involved in regeneration of reduced glutathione (GSH). This result in accumulation of reactive oxygen species (ROS) which cause cancer cell cycle exits and apoptosis.[5]

**Figure 1.**
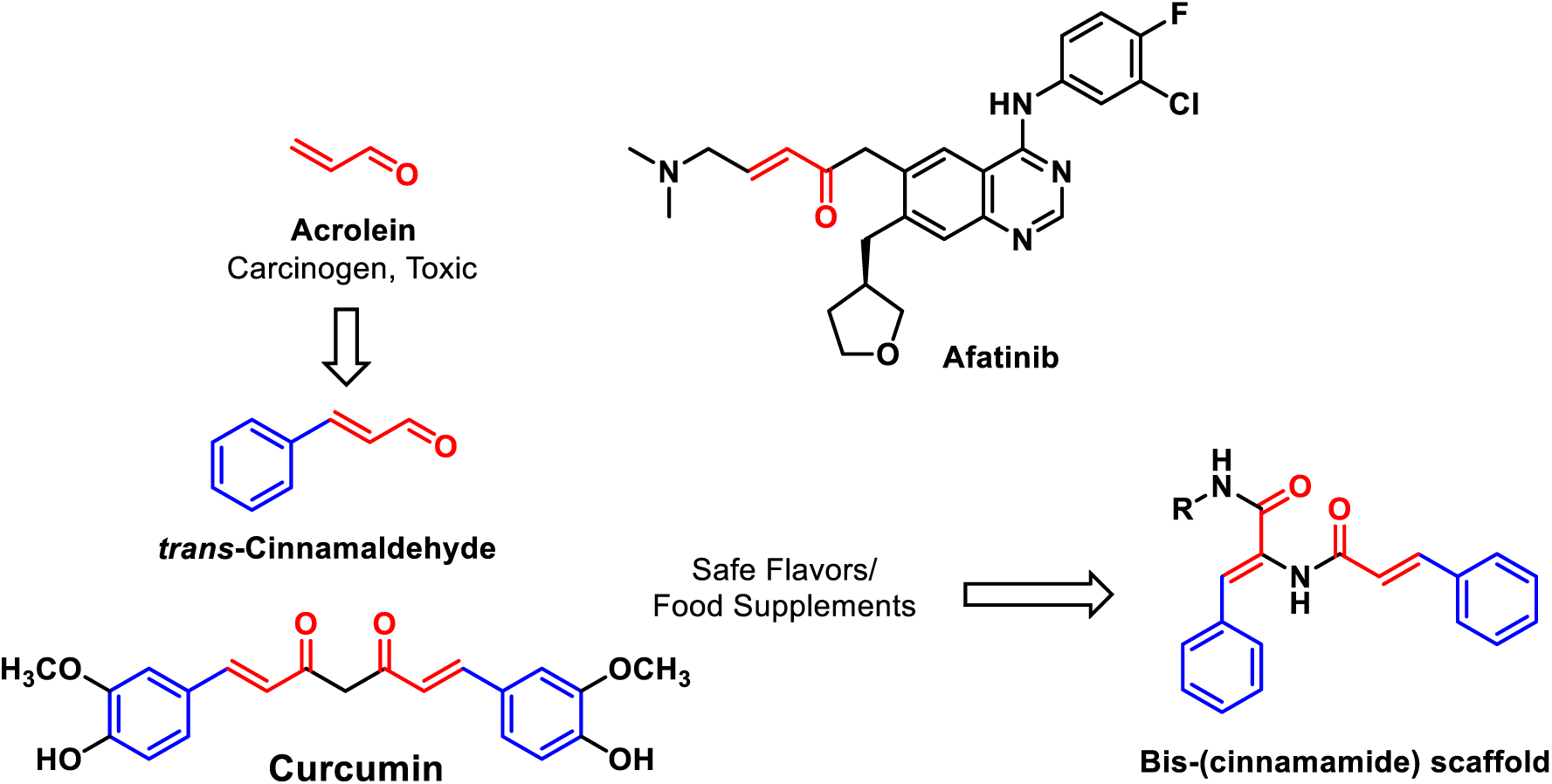
Michael Acceptors with bis-cinnamamyl moiety.

Acrolein (**Figure 1**) is a highly reactive and non-selective MA toxin and environmental pollutant with well-known carcinogenic activities. Interestingly, acrolein is also known to cause inhibition of the β-subunit of the proliferative anti-apoptotic protein NF*κ*B *via* covalent binding with the nucleophilic Cys-61 and Arg-307 leading to cytotoxicity and cell death of cancer cells.[6] Thus despite its apparent toxic effect, acrolein like other MAs has a potential as a therapeutic agent. Replacement of the aldehyde group of acrolein with amide group (acrylamide moiety) is known to attenuate the reactivity of the MA moiety leading to less toxic compounds.[7] Accordingly, safe natural MA products that are common in domestic condiments, such as *trans-*cinnamaldehyde and curcumin (**Figure 1**) are known to cause cancer cell death *via* redox mechanisms.[5] In the present study, we took advantage of *trans*-cinnamaldehyde (3-phenylpropenal) and curcumin structural features to design 2-cinnamamido-N-substituted-cinnamamide (bis-cinnamamide) compounds with a representative scaffold shown in figure 1. The choice of tCA and curcumin as building blocks to develop Michael Addition anticancer compounds with superior pharmacologic properties was clearly influenced by the lead pharmacophores presence in natural food, attractive anticancer activities, and well-known safety profiles. [5, 8–13] The ensuing derivatives as shown by our lead compound, **4112** and subsequent derivatives exhibited significant anti-proliferative effect on colorectal cancer cell lines (CaCo-2, HCT-116 and HT-29) in part via oxidative stress without significant effect on C-166 and BHK normal cell lines.

## 2. RESULTS

### 2.1. Chemical Synthesis of 2-cinnamamido-N-substituted-cinnamamide derivatives

The synthetic pathway, described in scheme 1 utilized Erlenmeyer chemistry for azlactone synthesis.[14] In this scheme, non-commercially available cinnamic acid analogues **2** were prepared by condensation of the corresponding aldehyde with malonic acid.[15] After conversion to the hippuric acid analogues (**3**), cyclocondensation was affected by their reaction with the corresponding aldehydes to afford the 2-arylvinyl-4-benzylidene-5-oxazolinone derivatives (azlactones, **4**) under Erlenmeyer conditions. The key azlactone intermediate **3** was subsequently reacted with a variety of aliphatic and aromatic amines to furnish the final compounds (Table 1). Generally, the aliphatic and benzylic amines reacted with the azlactone smoothly at room temperature in ethanol. The less reactive aromatic amines needed microwave heating in the presence of N,N-dimethylformamide (DMF) as a solvent. The ester **15EE** was prepared by solvolysis in boiling ethanol in presence of catalytic amount of 4-dimethylaminopyridine (DMAP).

**Scheme 1.**
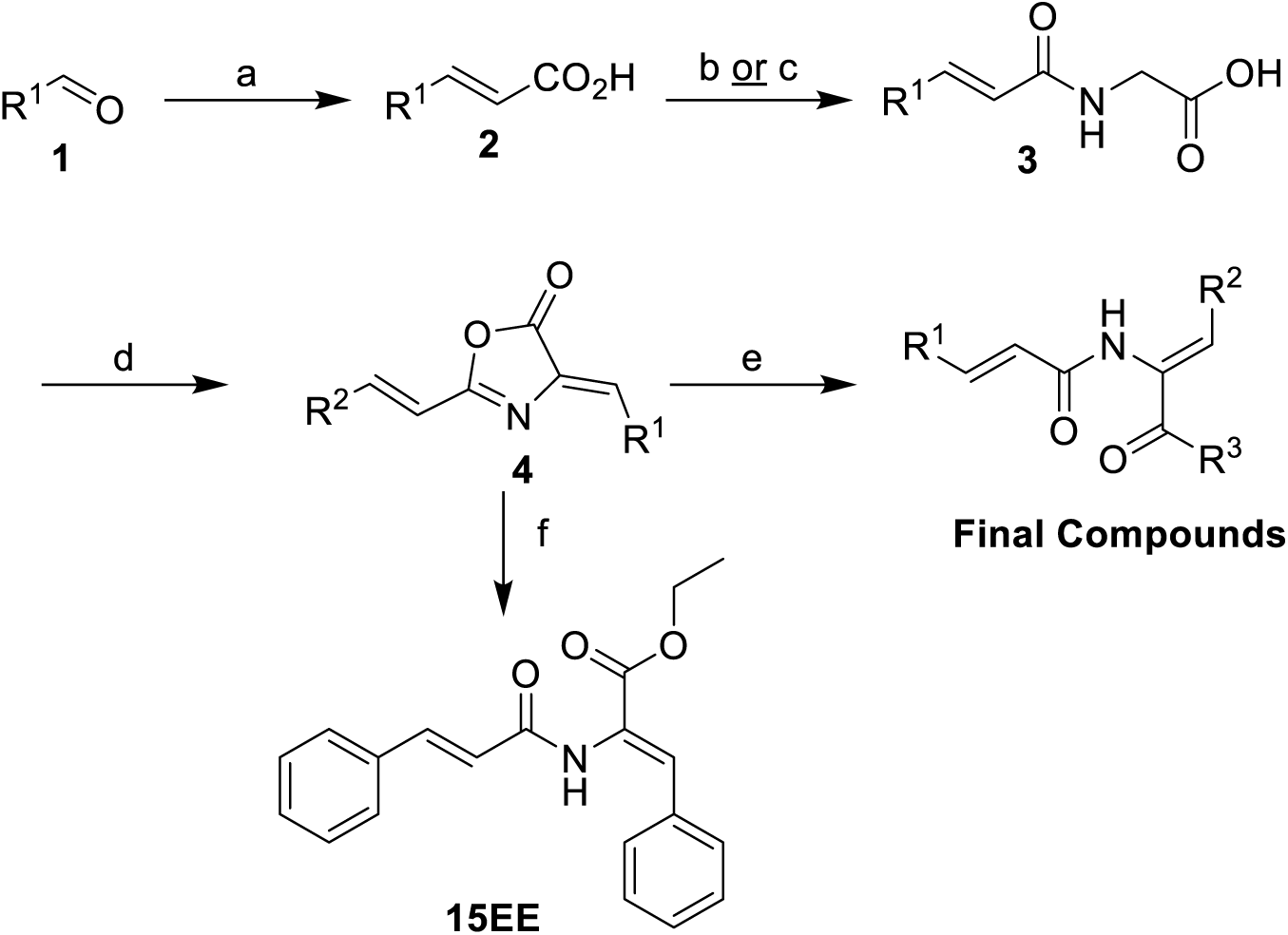
Synthesis of title compounds (a) malonic acid, pyridine, heat. (b) oxalyl chloride, DCM, DMF (cat.)then glycine Sodium carbonate, THF, Water (c) ethyl glycinate hydrochloride, Et_3_N, EDCI, DCM; OR, then NaOH, EtOH, H_2_O followed by aq. HCl (b) aryl aldehyde, acetic anhydride, heat, 15 min; (c) DMF, microwave heating; Or ethanol, room temperature; (d) EtOH, DMAP, heat, overnight.

**Table 1.** Cytotoxic activities of final compounds on colon cancer cell lines *IC50 values in µM±Standard Error of Means (SEM). **ND, Not Determined.

| Entry | Code | R<sup>1</sup> | R<sup>2</sup> | R<sup>3</sup> | HCT-116* | CACO-2* | HT-29* |
| --- | --- | --- | --- | --- | --- | --- | --- |
| 1 | 1501 | Ph | Ph | NHPh | 75.3±3.1 | ND | ND |
| 2 | 1502 | Ph | Ph | NH-CH<sub>2</sub>Ph | 53.6±0.6 | ND | ND |
| 3 | 1503 | Ph | Ph | NH-cyclopentyl | 48.5±3.1 | ND | ND |
| 4 | 1505 | Ph | Ph | NH-(1-adamantyl) | 14.7±0.3 | ND | ND |
| 5 | 1507 | Ph | Ph | NH-(3-pyridyl) | >100 | ND | ND |
| 6 | 1508 | Ph | Ph | NH<sub>2</sub> | >100 | ND** | ND |
| 7 | 1511 | Ph | Ph | NH-(4-methyl-1-piperazinyl) | >100 | ND | ND |
| 8 | 1512 | Ph | Ph | NH-(*n*-Pr) | 32.0±2.6 | 38.0±0.87 | 35.5±1.7 |
| 9 | 1513 | Ph | Ph | NH-(3-CN-Ph) | >100 | ND | ND |
| 10 | 1528 | Ph | Ph | NH-(4-F-Ph) | 91.9±4.4 | ND | ND |
| 11 | 1530 | Ph | Ph | NH-furfuryl | >100 | ND | ND |
| 12 | 1531 | Ph | Ph | NH-(2-morpholinoethyl) | >100 | ND | ND |
| 13 | 1532 | Ph | Ph | NH-(2-hydroxyethyl) | >100 | ND | ND |
| 14 | 1536 | Ph | Ph | 4-morpholinyl | >100 | ND | ND |
| 15 | 1555 | Ph | Ph | 1-pyrrolidinyl | >100 | ND | ND |
| 16 | 1556 | Ph | Ph | NH-(*n*-Bu) | 67.2±6.3 | ND | ND |
| 17 | 1557 | Ph | Ph | NH-(*sec*-Bu) | 52.4±7.1 | ND | ND |
| 18 | 1558 | Ph | Ph | NH-Et | > 100 | ND | ND |
| 19 | 1559 | Ph | Ph | N(Me)Et | 28.8±1.0 | ND | ND |
| 20 | 15EE | Ph | Ph | OEt | >100 | ND | ND |
| 21 | 1612 |  |  |  | >100 | ND | ND |
| 22 | 1712 |  |  |  | >100 | ND | ND |
| 23 | 1812 | 4-ClPh | Ph | NH-(*n*-Pr) | 10.3±0.34 | 10.7±0.5 | 7.90±0.43 |
| 24 | 1912 | 4-MeOPh | Ph | NH-(*n*-Pr) | 34.1± 1.8 | 18.5± 1.5 | 55.3± 3.2 |
| 25 | 2012 | 4-MePh | Ph | NH-(*n*-Pr) | 35.2±0.54 | 12.7±0.67 | 54.1±0.42 |
| 26 | 2112 | 2-thienyl | Ph | NH-(*n*-Pr) | 43.0±1.5 | 27.8±1.3 | 43.2±2.3 |
| 27 | 2212 | Ph | 4-FPh | NH-(*n*-Pr) | 15.2±1.2 | 13.2±1.0 | 7.20±0.31 |
| 28 | 2312 | 4-ClPh | 4-FPh | NH-(*n*-Pr) | **3.80±0.24** | **2.60±0.233** | **4.00±0.14** |
| 29 | 2412 | 4-MeOPh | 4-FPh | NH-(*n*-Pr) | 29.3±1.2 | 45.73±1.2 | 86.5±2.9 |
| 30 | 2512 | 4-MePh | 4-FPh | NH-(*n*-Pr) | 6.70±0.36 | 5.10±0.17 | 11.80±0.18 |
| 31 | 2612 | 2-thienyl | 4-FPh | NH-(*n*-Pr) | 17.3±0.13 | 10.7±0.68 | 7.68±0.05 |
| 32 | 2712 | Ph | 4-(Me<sub>2</sub>N)Ph | NH-(*n*-Pr) | 6.95±0.46 | **4.24±0.09** | 6.8±0.05 |
| 33 | 2812 | 4-ClPh | 4-(Me<sub>2</sub>N)Ph | NH-(*n*-Pr) | 20.7±0.32 | 17.0±0.8 | 9.52±0.42 |
| 34 | 2912 | 4-MeOPh | 4-(Me<sub>2</sub>N)Ph | NH-(*n*-Pr) | 12.4±0.25 | 15.3±0.05 | 36.9±0.25 |
| 35 | 3012 | 4-MePh | 4-(Me<sub>2</sub>N)Ph | NH-(*n*-Pr) | 22.1±0.10 | 29.4±0.08 | 39.6±0.22 |
| 36 | 3112 | 2-thienyl | 4-(Me<sub>2</sub>N)Ph | NH-(*n*-Pr) | 11.6±0.39 | 27.7±3.7 | 66.5±0.10 |
| 37 | 3212 | Ph | 3-MeO-4-OHPh | NH-(*n*-Pr) | >100 | 6.32±0.02 | 9.14±0.23 |
| 38 | 3312 | 4-ClPh | 3-MeO-4-OHPh | NH-(*n*-Pr) | 37.8±1.8 | 6.1±0.44 | 5.4±0.34 |
| 39 | 3412 | 4-MeOPh | 3-MeO-4-OHPh | NH-(*n*-Pr) | 38.4±2.1 | 18.0±1.2 | 48.6±0.4 |
| 40 | 3512 | 4-MePh | 3-MeO-4-OHPh | NH-(*n*-Pr) | 26.3±1.3 | 20.9±1.1 | 15.1±0.67 |
| 41 | 3612 | 2-thienyl | 3-MeO-4-OHPh | NH-(*n*-Pr) | 24.8±1.4 | 28.3±1.3 | 48.3±1.9 |
| 42 | 3712 | Ph | 3-indolyl | NH-(*n*-Pr) | **1.65±0.05** | **2.1±0.03** | **2.36±0.04** |
| 43 | 3812 | 4-ClPh | 3-indolyl | NH-(*n*-Pr) | **3.51±0.21** | **3.35±0.05** | **3.41±0.03** |
| 44 | 3912 | 4-MeOPh | 3-indolyl | NH-(*n*-Pr) | **4.35±0.29** | **2.07±0.03** | **2.49±0.04** |
| 45 | 4012 | 4-MePh | 3-indolyl | NH-(*n*-Pr) | **3.39±0.10** | **2.3±0.08** | **3.12±0.06** |
| 46 | 4112 | 2-thienyl | 3-indolyl | NH-(*n*-Pr) | **2.85±1.5** | **0.89±0.04** | **1.65±0.07** |
| 47 | 4212 | Ph | 3-pyridyl | NH-(*n*-Pr) | 73.0±2.5 | 66.0±2.2 | >100 |
| 48 | 4312 | 4-ClPh | 3-pyridyl | NH-(*n*-Pr) | 49.4±1.9 | 47.3±1.9 | 36.0±1.7 |
| 49 | 4412 | 4-MeOPh | 3-pyridyl | NH-(*n*-Pr) | 52.1±1.6 | 38.1±2.0 | 72.7±6.4 |
| 50 | 4512 | 4-MePh | 3-pyridyl | NH-(*n*-Pr) | >100 | 27.0±1.8 | >100 |
| 51 | 4612 | 2-thienyl | 3-pyridyl | NH-(*n*-Pr) | >100 | 41.8±1.4 | 68.8±6.9 |
| 52 | 4712 | Ph | 4-NO<sub>2</sub>Ph | NH-(*n*-Pr) | 17.3±1.3 | 44.2±2.7 | 23.2±1.9 |
| 53 | 4812 | 4-ClPh | 4-NO<sub>2</sub>Ph | NH-(*n*-Pr) | >100 | >100 | 36.9±1.5 |
| 54 | 4912 | 4-MeOPh | 4-NO<sub>2</sub>Ph | NH-(*n*-Pr) | >100 | >100 | 74.3±3.6 |
| 55 | 5012 | 4-MePh | 4-NO<sub>2</sub>Ph | NH-(*n*-Pr) | >100 | >100 | >100 |
| 56 | 5112 | 2-thienyl | 4-NO<sub>2</sub>Ph | NH-(*n*-Pr) | >100 | 34.2±1.1 | >100 |
| 58 | tCA |  |  |  | 12.4 ± 1.2 | ND | ND |
| 59 | Doxorubicin |  |  |  | 0.6±0.10 | 0.14±.01 | 0.3±0.01 |

The homologues **1612** and **1712** were prepared in similar ways starting with the appropriate azlactones (see experimental section).

### 2.2. Biological Screening

#### 2.2.1. Antiproliferative activities of bis-cinnamamide derivatives

Compounds in this study were tested against colorectal cancer cell line HCT-116, using sulforhodamine-B (SRB) assay.[16] The first twenty compounds (Codes **1501** to **1559**), having fixed phenyl groups at R^1^ and R^2^ positions (**Figure 2**), showed moderate to weak antiproliferative activities (**Table 1**). The compound with the highest activity in this subset was the *N*-(1-adamantyl) analogue (**1505**, Entry 4) with IC_50_ = 14.7 ± 0.3 µM, but with poor solubility (ClogP 6.08), while compound **1512** (Entry 2) with moderate activities (IC_50_ = 32.0 ± 2.6 µM) showed better solubility profile (CLogP 4.48).

**Figure 2.**
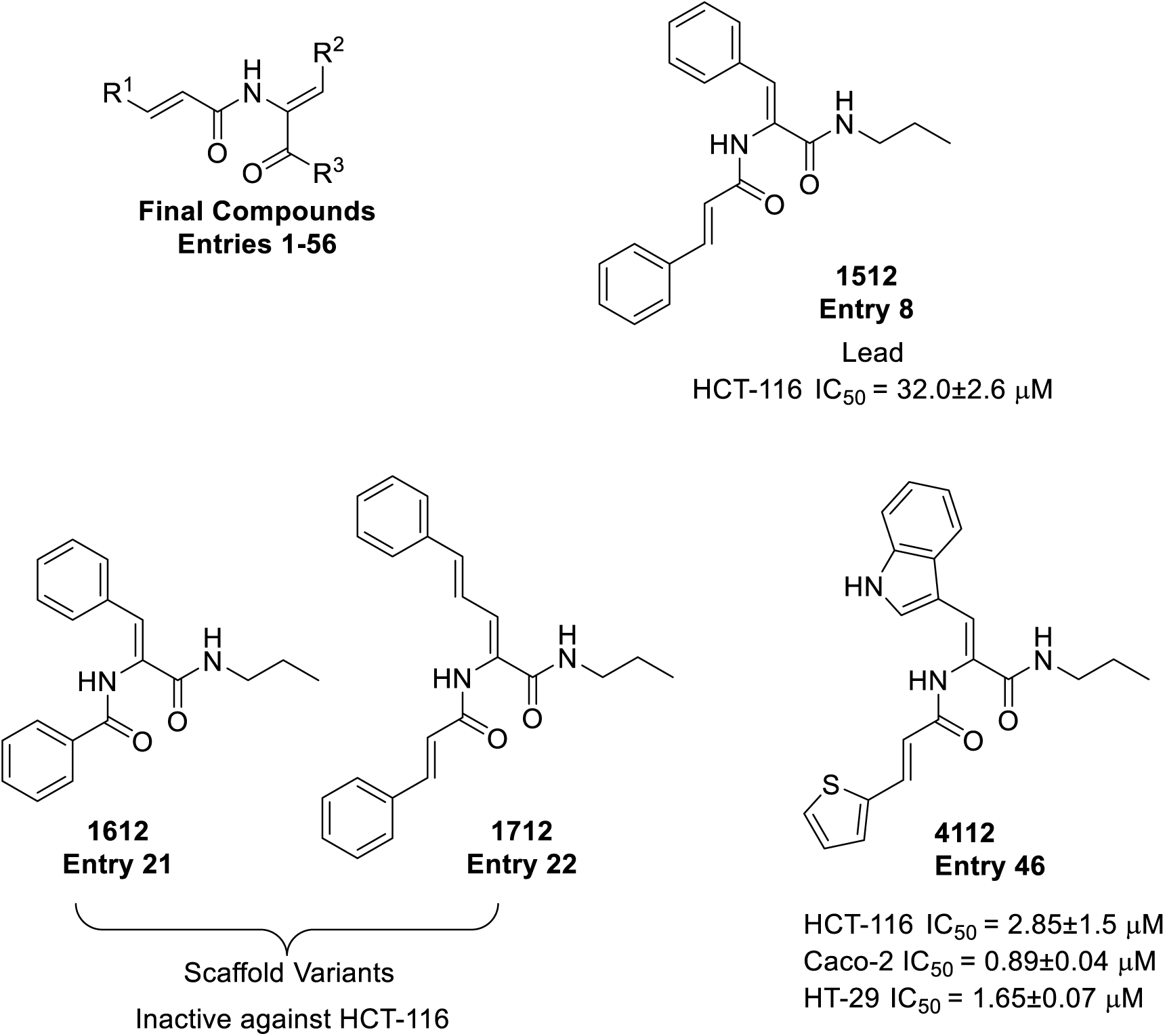
Scaffold for compounds included in table 1 and prominent entries.

Importantly, **1512** showed significantly low cancer cell resistance (R-value = 8.2%),[14] and high selectivity in killing cancer cells (HCT-116, Caco-2 and HT-29) versus highly proliferative normal cells (C-166 mouse skin fibroblasts) (**Figure 3**).

**Figure 3.**
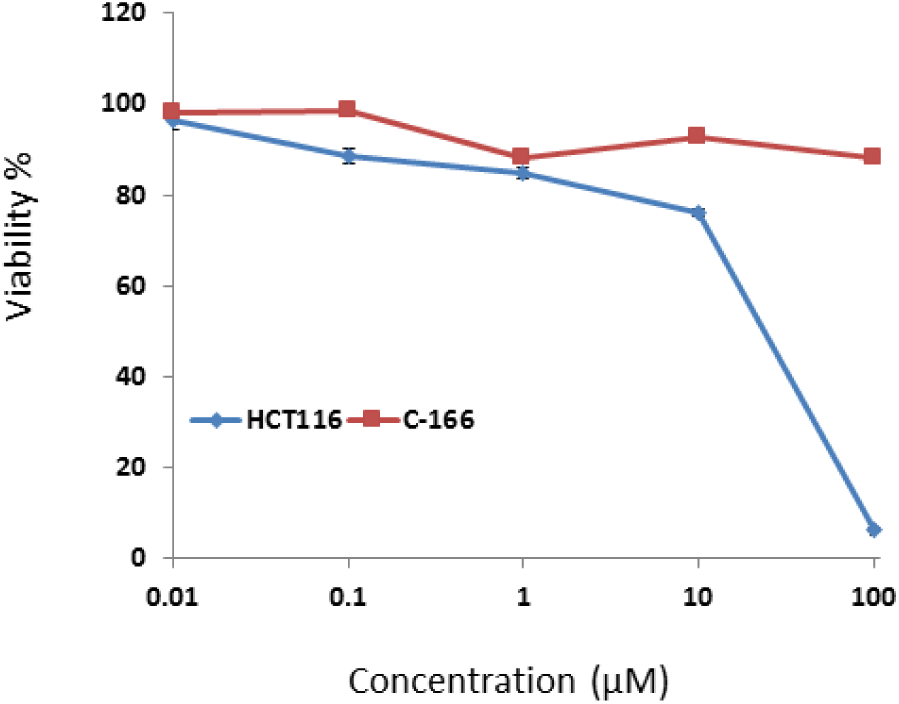
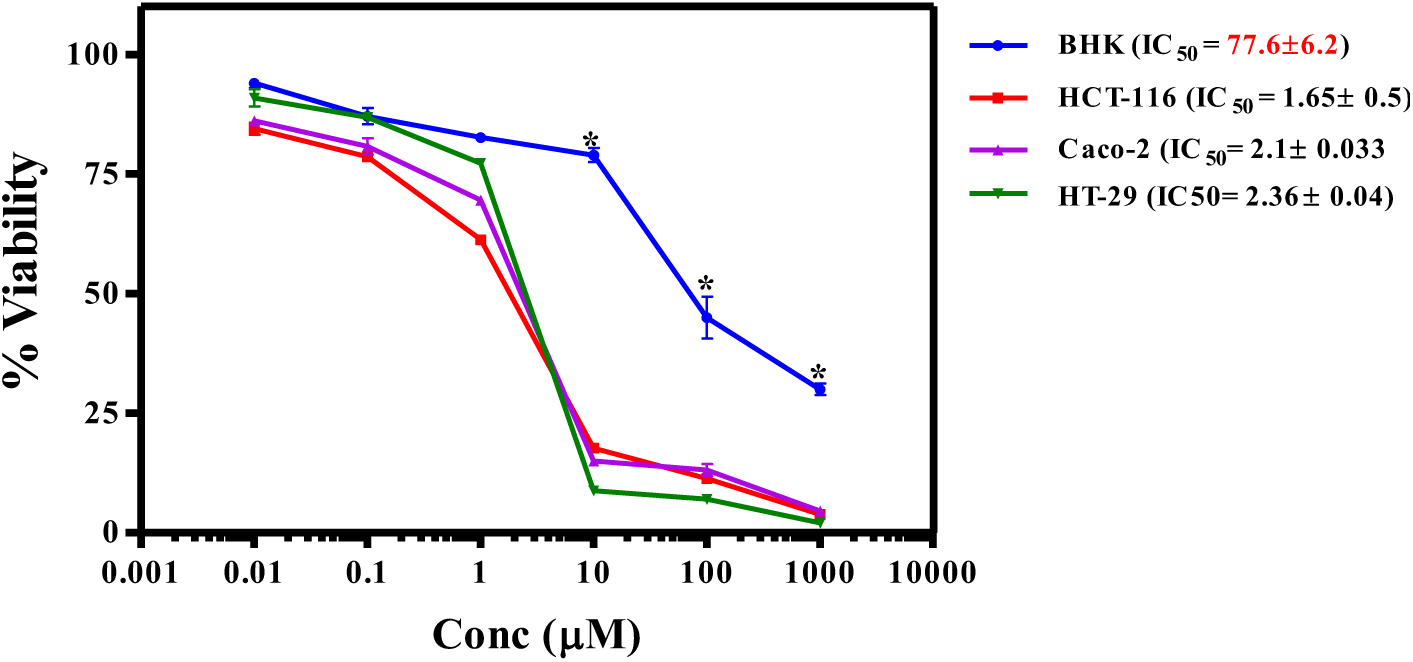
(A) Dose-response curve of compound 1512 on HCT-116 cell lines (blue) and C-166 cell line (brown). (B) Cytotoxic effect of 4112 against human colon adenocarcinoma cancer cells (HCT-116, Caco-2 and HT-29) vs. normal baby hamster kidney cells (BHK). All cells were exposed to different concentrations of 4112 for 72 h. The cell viability concentration curves were plotted, and the IC50 were determined. *denotes significance at P<0.001

Based on **1512** favorable pharmacologic and physicochemical properties, it was selected as a new lead for the second subset of compounds (Entries **23** to **56**). The compounds in this series contained variable groups at R^1^ and R^2^ (**Figure 2**) that were all tested against the three cancer cell lines (HCT-116, Caco-2 and HT-29). Some of the compounds showed high potencies similar to the reference drug doxorubicin (Table 1). For instance, entries **28**, **32**, **42**, **43**, **44**, **45** and **46** demonstrated excellent potencies against CRC cell lines (under 5 µM) with compound **4112** (Entry **46**) exhibiting the most significant inhibition of HCT-116, Caco-2, and HT-29 cells (IC50 of 2.85 ± 1.5, 0.89 ± 0.04 and 1.65 ± 0.07 µM, respectively) that are comparable to doxorubicin 0.6 ± 0.1 0.14 ± 0.05, and 0.3 ± 0.01 µM, respectively. Compound **4112** also had excellent selectivity against cancer cells versus baby hamster kidney (BHK) cell lines (IC_50_ = 77.6 µM). Also, notable is another potent compound **3712**, which exhibited IC_50_ of 1.65 ± 0.05, 2.1 ± 0.03 µM and 2.36 ± 0.04 against HCT-116, Caco-2, and HT-29 cells, respectively.

#### 2.2.3. Induction of apoptosis and apoptotic changes in HCT-116 cancer cell lines upon treatment with 1512 and 4112

Several assays were performed to determine the apoptotic effect of the lead **1512** as well as the derived **4112** on HCT-116 cells. First, the effect of **1512** on morphological changes of HCT-116 was monitored by staining the cell with the dye 4’,6’-diamidino-2-phenylindole (DAPI) after 48 hours of treatment with 32 µM of **1512** (IC_50_ concentration). We observed very strong blue fluorescence, indicative of increased cell membrane permeability and uptake of the dye, suggesting that **1512** is able to disrupt the integrity of the cell to induce apoptosis (**Figure 4A**).[17]

**Figure 4.**
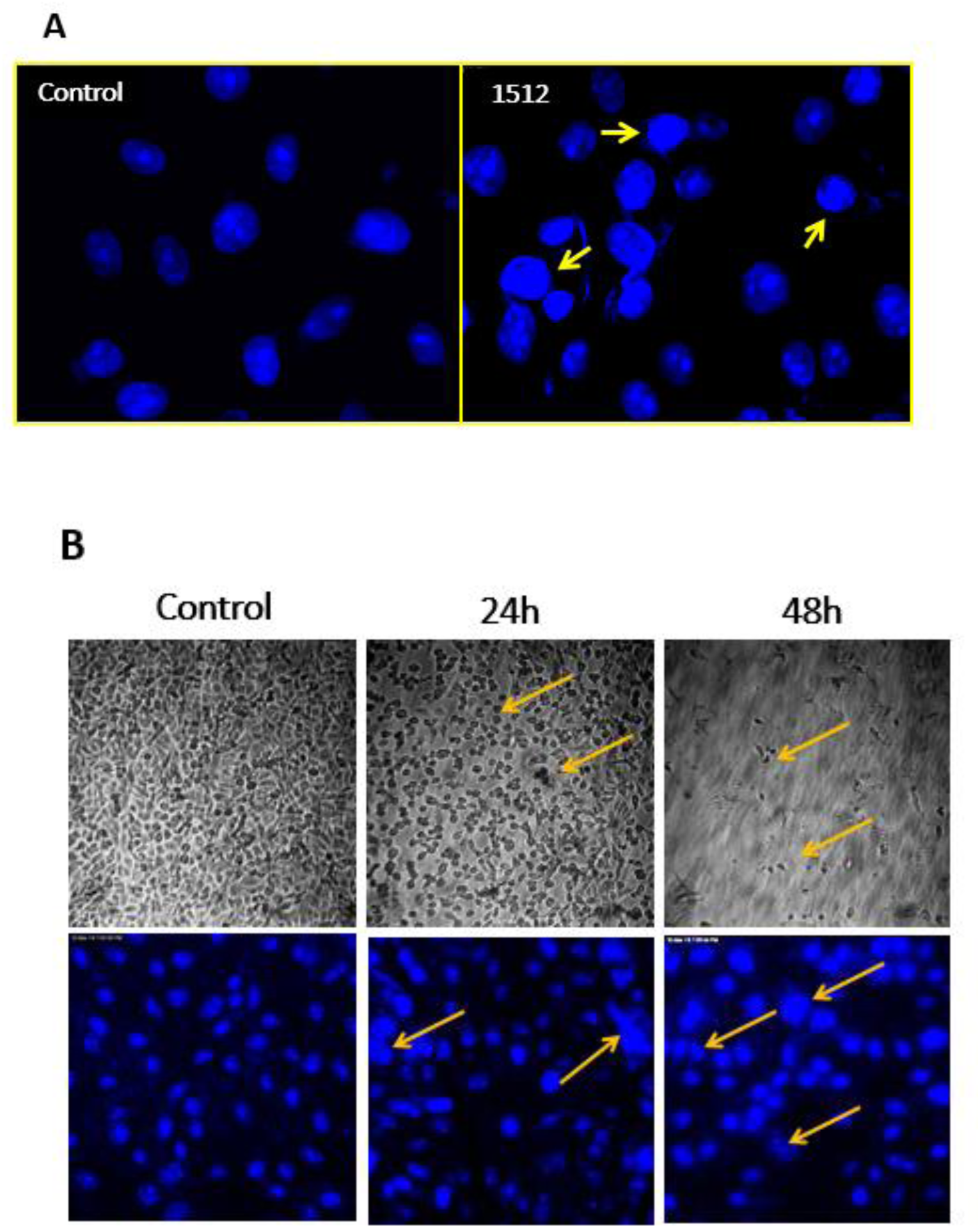
Effect of 1512 at 32 µM (IC_50_ concentration) on the nuclear structures of HCT-116 cells. Treated cells show nuclear condensation and fragmentation along with the condensed blue fluorescence of DAPI. (B) Morphological changes in HCT-116 cells following exposure to compound 4112. The cells were left untreated or treated with the IC_50_ for 24 and 48 h. Pictures at top show the morphological changes examined under light microscope. Images at the bottom show the nuclei stained with DAPI and visualized under fluorescence microscope. Arrows point to apoptotic cells.

DAPI staining test was also performed after incubation of HCT-116 cells with **4112** at 2.85 µM (the IC_50_ value), and the results at 24 and 48 h shown in figure 4B. Expectedly, the percentage of cells with fragmented DNA and condensed nuclei greatly increased after 48 h than 24 h.

#### 2.2.4. Cell cycle distribution upon treatment with 1512 and 4112

The effect of **1512** and **4112** on HCT-116 cell cycle distribution was examined at the respective IC_50_ concentrations for 24 h using a FACS Calibur flow cytometer. The exposure of the cells to **1512** led to a significant increase in the proportion of cells in pre-G1 phase (Up to 9-fold compared to the control) (**Figure 5A**). Accumulation of cells in pre-G1 phase, likely as a result of degradation or fragmentation of genetic material indicates a possible role for apoptosis through compound-induced growth inhibition.

**Figure 5.**
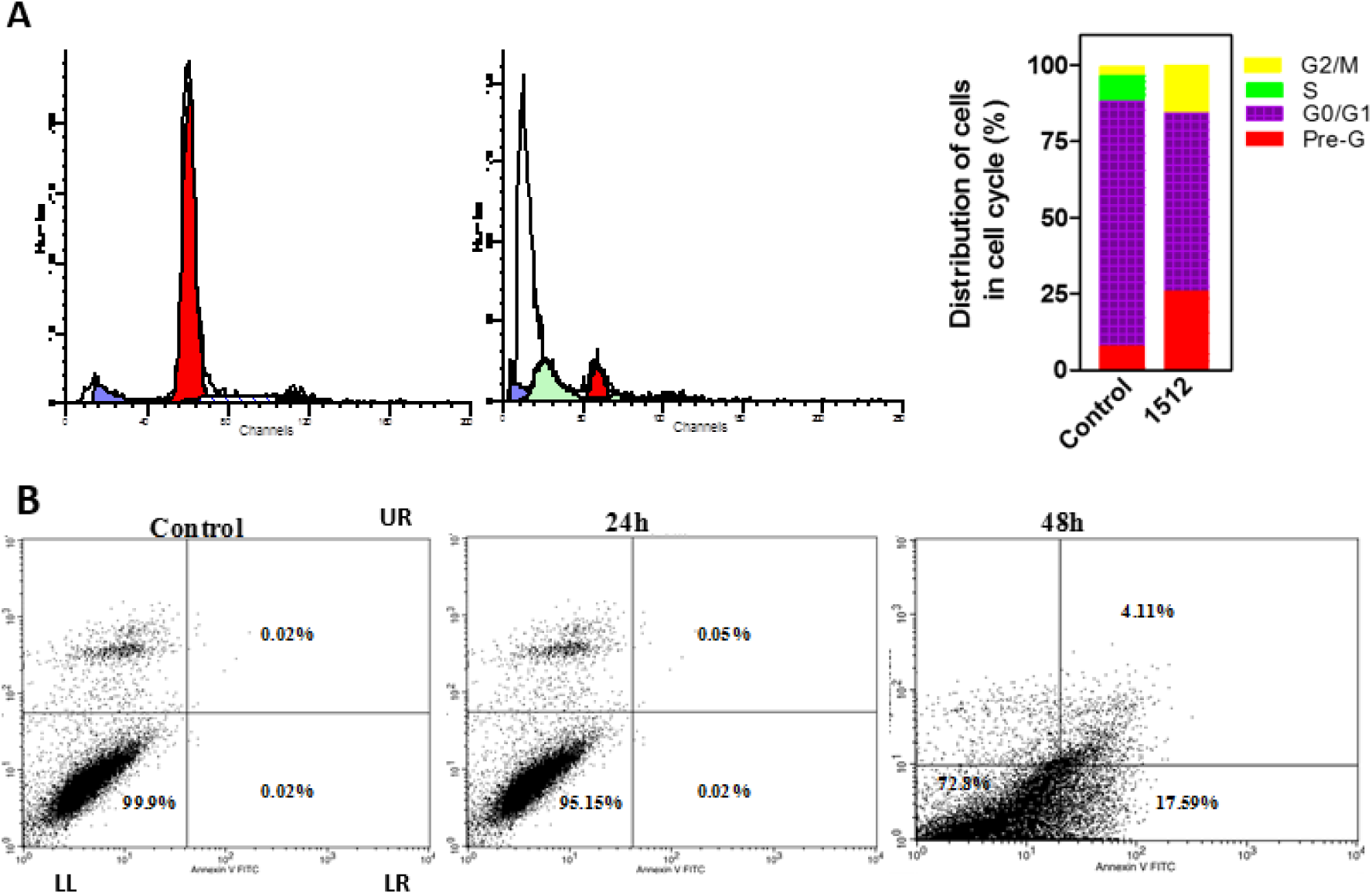
(A) Representative DNA histograms of HCT-116 cells following treatment with 1512 at 32 µM (IC_50_ concentration) for 24 h. (B) Effect of 4112 on HCT-116 cell cycle progression: Key LL= Viable cells, LR= Early apoptotic cells, UR= Late apoptosis.

In the subsequent similar experiment, **4112**-treated HCT-116 cells were monitored for apoptotic changes at 24 h and 48 h. Cell cycle populations for the treated cells were determined by flow cytometry analysis after staining with Annexin V/Propidium iodide (**Figure 5B**) and revealed a large time-dependent increase in the percentage of HCT-116 apoptotic cells. At 24 hours, there was very little change in cell viability, while after 48 hours, viable cells have decreased to 72% (down from 99.9%). In the context, cell populations in both early and late apoptosis increased from almost zero to 17.5% and 4.1%, respectively.

#### 2.2.5. Changes in marker proteins in apoptotic cells upon treatment with 1512 and 4112

Using antibody immunofluorescence assays, the apoptotic changes upon exposure to compound **1512** were confirmed by testing changes in cellular contents of cyclin B1 and cyclin D1, phosphohistone 3 and cleaved caspase 3 (**Figure 6**). As expected, all were observed to increase significantly compared to untreated cells.

**Figure 6.**
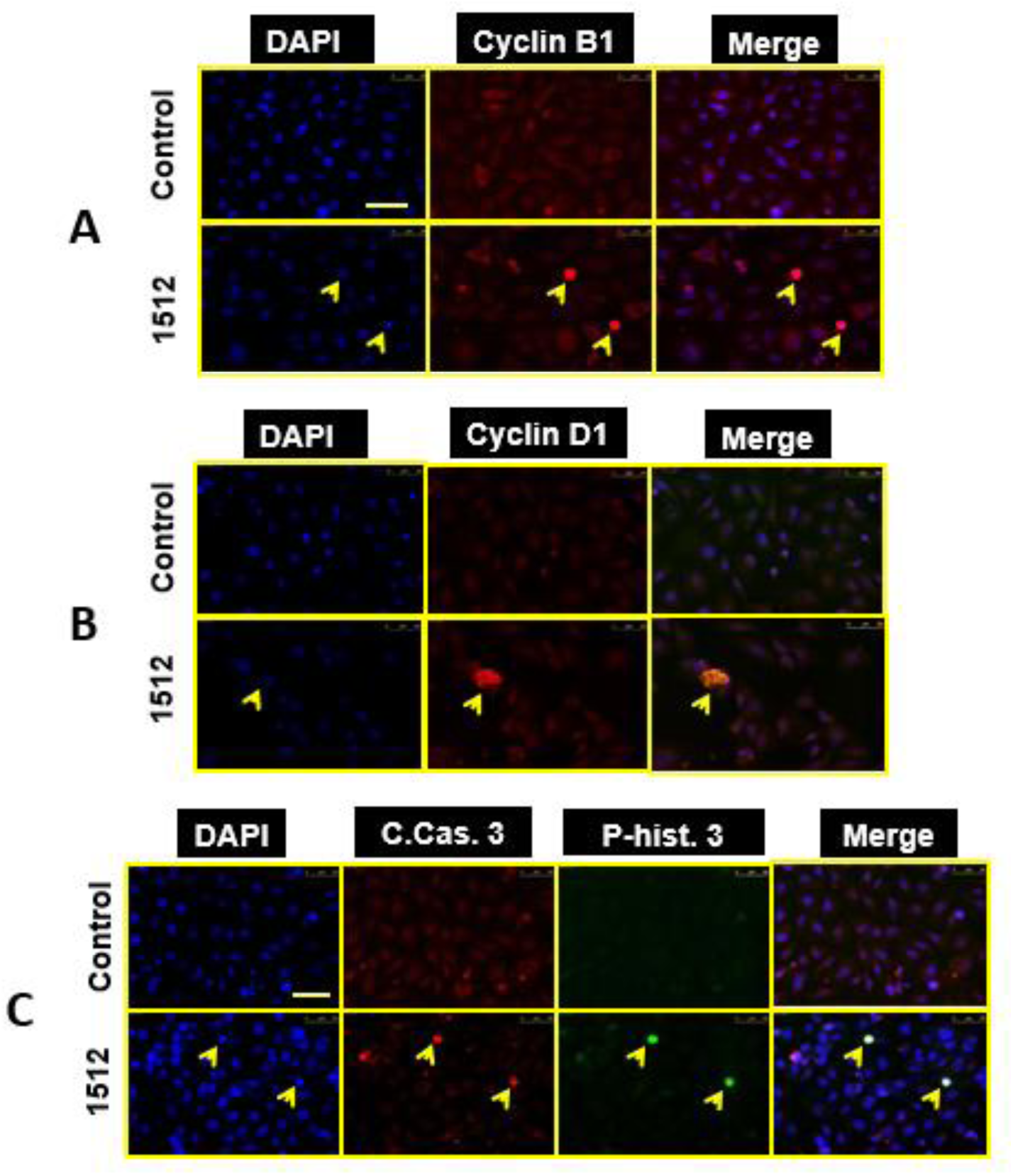
Effects of 1512 and on cellular expression of cyclin B1 (A) cyclin D1 (B) in HCT-116 after treatment for 24 h compared to control. Cells were stained with DAPI and incubated with a Cy3-coupled secondary antibody to visualize the distribution of cyclin B1 and cyclin D1 (red) proteins. Treatment of HCT-116 with 1512 increased the nuclear expression of cyclin B1 and cyclin D1 (arrow head) compared with the untreated cells. Scale bar, 50 µm. (C) Effect of 1512 on the cellular expression of caspase-3 (C.Cas. 3) and phospho-histone 3 (P-hist. 3) in human colon cancer cells (HCT-116). The cells were stained with DAPI to visualize nuclei (blue) and with Alexa Fluor 488- and Cy3-coupled secondary antibodies to visualize the distribution of caspase-3 (red) and phospho-histone-3 (green) proteins using immunofluorescence microscopy. The nuclear expression of the pro-apoptotic proteins, activated caspase 3 and phospho-histone 3 (yellow arrows) compared to the untreated cells. Scale bar, 50 µm.

A similar subsequent study with compound **4112** showed increased levels of cleaved caspase 3 in a time-dependent manner (24 and 48 h), indicating nuclear release of this apoptotic marker upon treatment with **4112**. (**Figure 7**).

**Figure 7.**
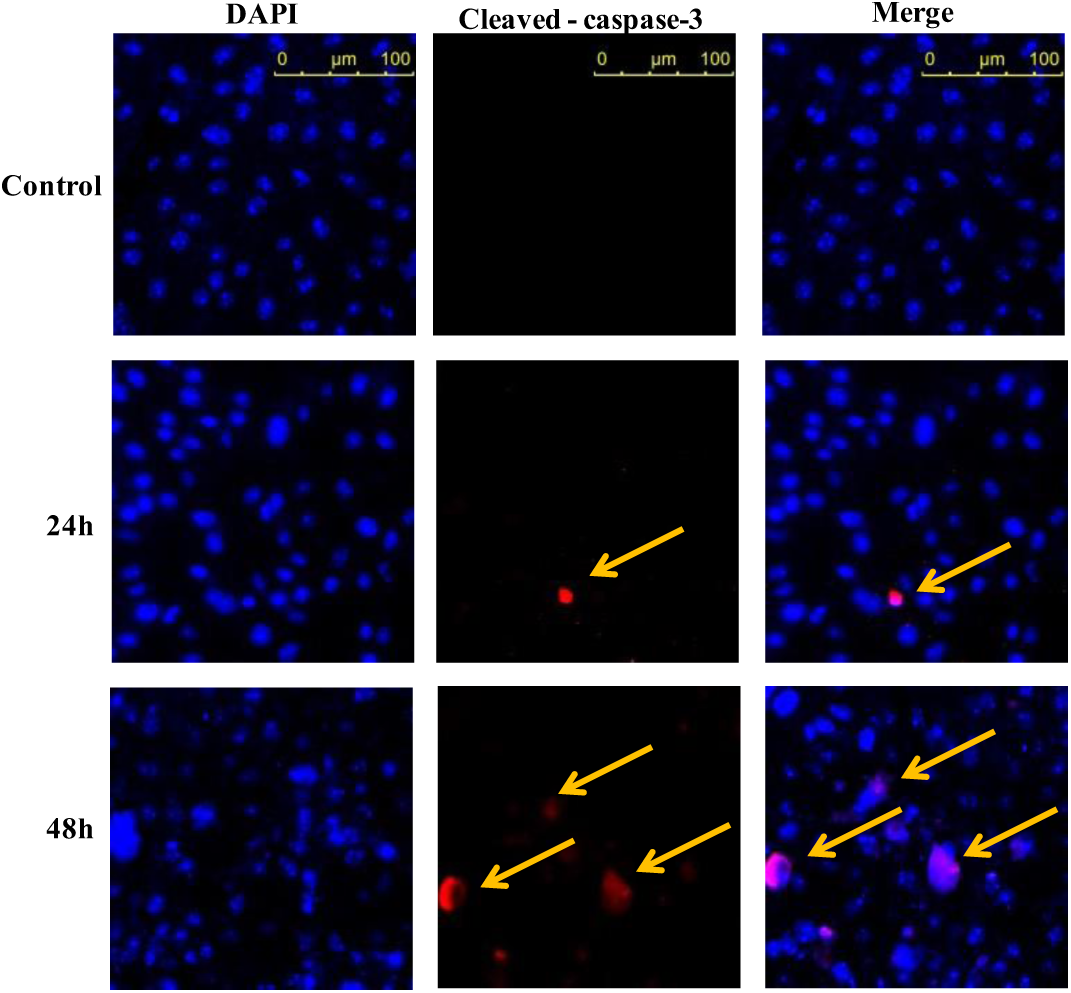
Effect of 4112 on cleaved caspase-3 expression in HCT-116 cells. The cells were stained with DAPI to visualize nuclei (blue) and with Alexa Fluor 488- and Cy3-coupled secondary antibodies to visualize the distribution of caspase-3 (red). Scale bar, 100 µm

#### 2.2.6. Elevation of oxidative stress indicators within HCT-116 cells upon treatment with 1512 and 4112

The underlying mechanism by which our compounds induced cancer cells to undergo apoptotic cell death was investigated by measuring changes in redox indicators within HCT-116 CRC cell lines. In this study, we first checked the influence of **1512** on cellular activities of the peroxide scavenging enzymes catalase and superoxide dismutase (SOD) (**Figure 8A**). Activities of both enzymes significantly diminished to 34.7% and 25.1% respectively, compared to untreated cells (control). However, the cellular contents of the non-enzymatic SH containing redox markers, such as reduced glutathione (GSH) and malondialdehyde (MDA, a lipid oxidation marker) were only slightly affected (95.2% and 103.4% respectively, compared to control).

**Figure 8.**
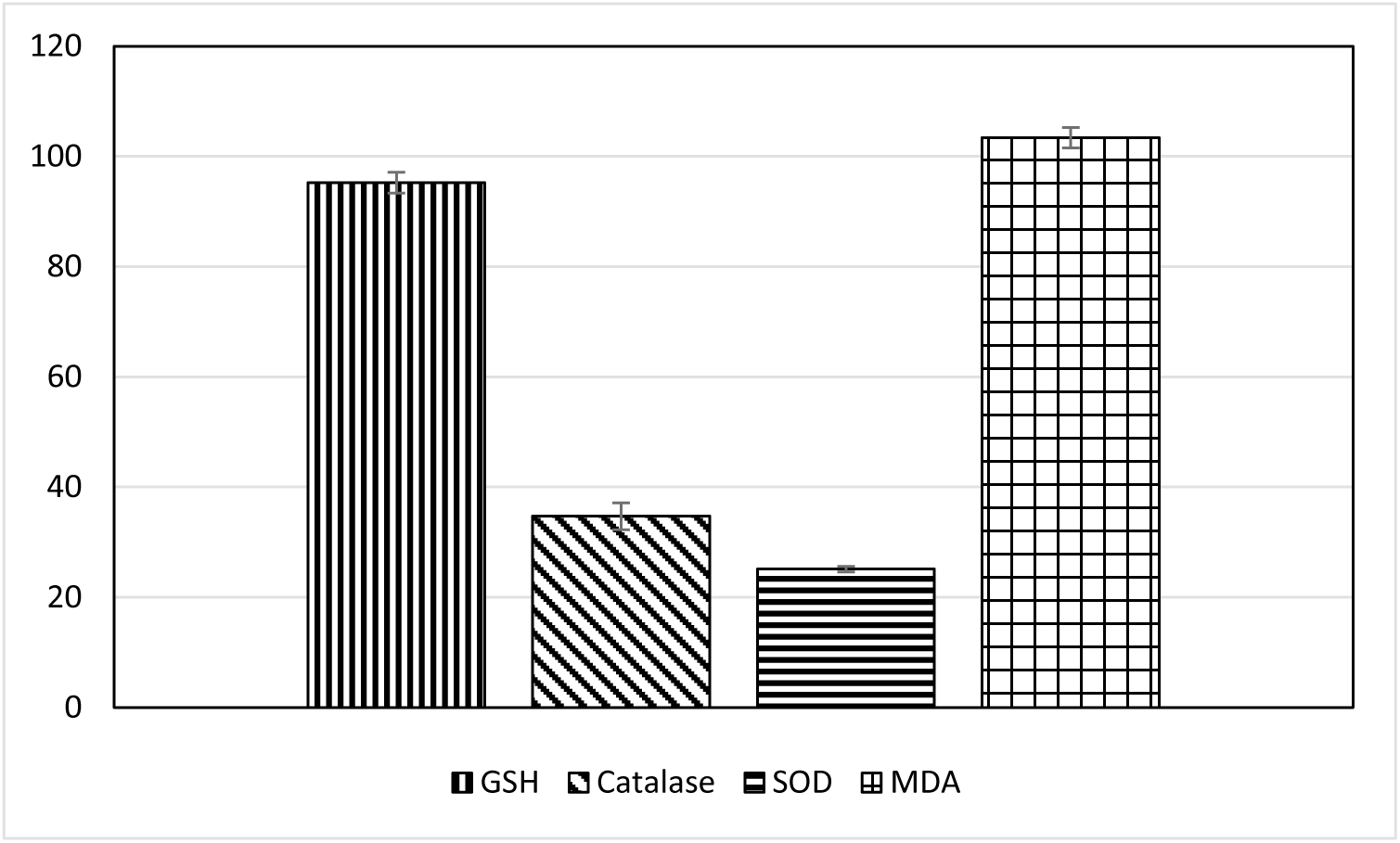
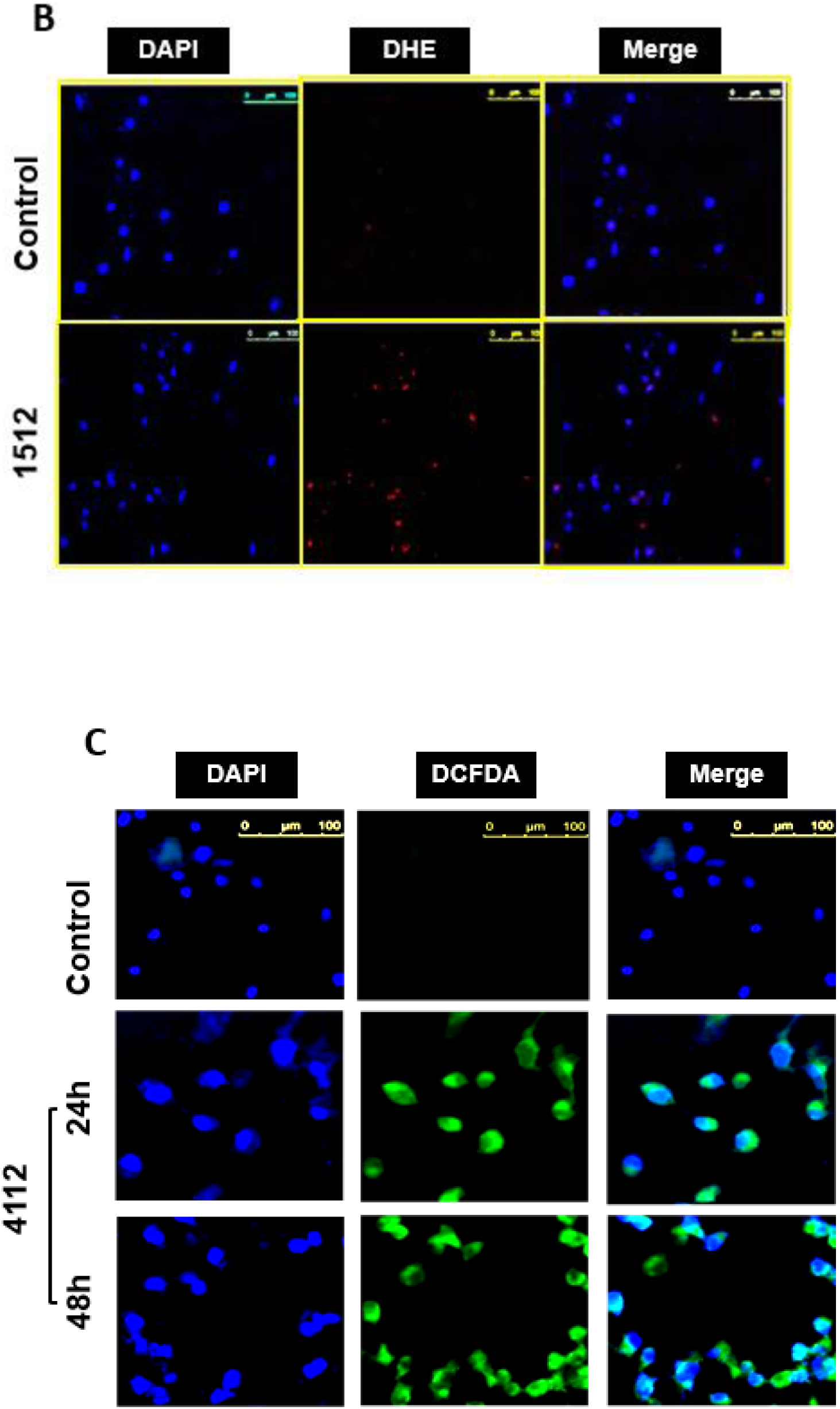
(A) Effect of 1512 (32 uM, 48 h) on redox indicators with HCT-116. The values are expressed as percentage of controls and calculated from the means of three independent experiments. (B) Effect of compound 1512 on superoxide anion generation in HCT-116 cells using DHE staining method. Red fluorescence represents DHE staining and DAPI was used as counter nuclear stain. (C) Effect of 4112 on ROS production from HCT-116 cells using DCFDA staining. Green fluorescence represents DCFDA staining and DAPI was used as counter nuclear staining. Scale bar, 100 µm.

**Figure 9.**
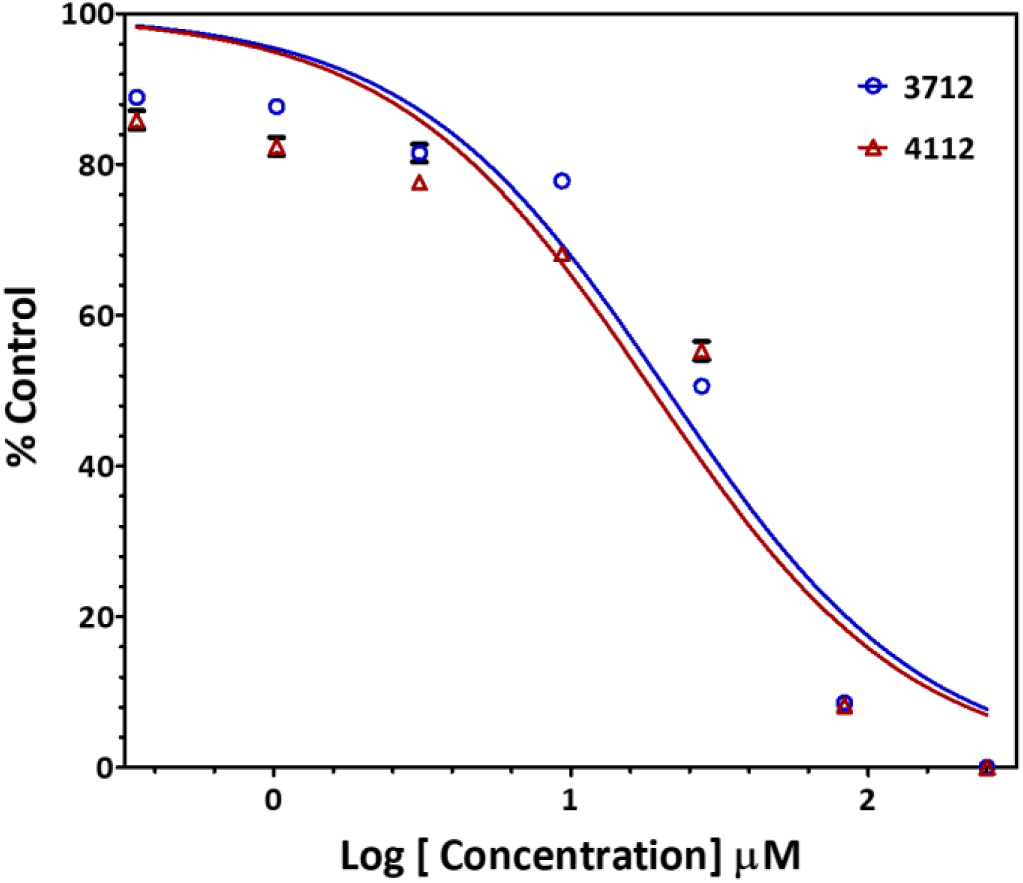
Dose-response curve of HT-29 cell spheroids (expressed as % of vehicle treated cells) by increasing concentration of 3712 (blue) and 4112 (red).

To confirm the increased intracellular ROS production, we performed dihydroethidium (DHE) staining test at which untreated HCT cancer cells were compared to **1512**-treated cells (at 32 µM, the IC value). As seen in figure 8B, the cells treated with **1512** attained more red fluorescence than the untreated cells indicating higher cellular peroxide content, which resulted from increased ROS production within the drug-treated cancer cells [18].

Regarding **4112**, ROS release was monitored over time (24 and 48 h) by using 2′,7′-dichlorofluorescin diacetate (DCFDA) dye which detect various ROS species and emit green fluorescence. It was clear that this compound causes accumulation of ROS and it is proportional to the time of exposure (**Figure 8C**).

#### 2.2.2. Inhibition of cancer stem cell proliferation in HT-29 cancer cell lines upon treatment with 3712 and 4112

We also evaluated **4112**, and another potent compound **3712** for their ability to inhibit cancer stem cell proliferation using a primary (1°) colonosphere formation assay. In this assay, HT-29 colon cancer cells were grown in low adhesion plates to form spheres as previously described.[19] Wells treated with our compounds were compared to vehicle treated cells for primary sphere growth inhibition after 5 days of incubation. Results, as illustrated in figure 4, showed a dose-dependent inhibition of colon cancer spheroids formation (> 50 µm). The IC_50_ of the two compounds were 21.12 ± 1.3µM and 18.85 ± 1.1 µM for **3712** and **4112** respectively, reflecting similar closeness in potency as in HT-29 monolayer cell line assay (2.36 ± 0.04 µM and 1.65 ± 0.07 µM, respectively). Of note, monolayer growth conditions are ideally suited for examining cellular proliferation, whereas spheroid growth conditions examine cancer stem cell growth and self-renewal properties. Hence, differential potency of the molecules in the two condition is reflective of their effects on two different phenotypes.

#### 2.2.7. Lethal dose toxicity test with compounds 1512, 3712 and 4112

Since compounds with electrophilic properties, e.g. MA are usually involved in variety of toxic effects on mammalians, we decided to investigate the general acute toxicity of **1512**, **3712** and **4112** by determining the *in vivo* lethal dose (LD_50_) according to the Globally Harmonized Classification System (GHS) and following the OECD guideline 423 (modified, adopted March 23, 2006). As observed in our cellular assay, the compound safety was confirmed, as the LD_50_ for all compounds were between 2000-5000 mg/kg (Category 5).[20]

## 3. DISCUSSION

We have continued interest in discovering new leads for chemotherapy against colorectal cancer [14, 21] as this fatal illness is the third most common diagnosed cancer in the world with nearly 1.4 million new cases diagnosed each year [22]. *Trans-*cinnamaldehyde and curcumin, commonly found ingredients in domestic condiments are two known MA compounds.[23, 24] These compounds have been studied for their antiproliferative and toxicity activities.[5, 8–13] We decided to take advantage of the pharmacologic and low toxicities of these two pharmacophores to develop other MA compounds with more potent anticancer properties but with similar low toxicity profile.

Erlenmeyer chemistry was utilized to synthesize the first batch of compounds (**1508** to **15EE**) varying only at the R^3^ substituent (see **Figure 2** and **Table 1**). Despite the moderate to weak anti-proliferative effects of this series, it was a good starting point to explore the SAR of bis-cinnamamides. In a cell line inhibition assay against the CRC cell line HCT-116, it was clear that hydrophobic R^3^ substituents (e.g. compounds **1505**, **1512**, **1556**, **1557** and **1559)** demonstrated superior activities when compared to their hydrophilic counterparts (e.g. compounds **1511**, **1530**, **1531**, **1532** and **1536) (Table 1)**. This SAR aspect is highlighted by comparing the activities of compounds **1532** and **1512**, where the more hydrophilic compound **1532** is relatively inactive while **1512** shows moderate activity despite differing from **1532** only by a terminal methyl instead of the terminal hydroxyl group of **1532**.

Interestingly, analogues bearing hydrophobic amides (e.g. compounds **1501**, **1502**, **1513**, **1529** and **1530)** showed lower anti-proliferative activities compared to aliphatic hydrophobic compounds (**Table 1**). It is quite possible that some compounds that show inactivity are likely due to poor cell membrane permeability, such as compound **1508**.

Despite the relatively moderate to weak activities of the synthesized compounds, they showed consistent SAR and provided a good starting point for developing more potent analogues. Although the *N*-adamantyl compound **1505** displayed higher potency (IC_50_ = 14.7 µM), the n-propyl amide analogue **1512 (**IC_50_ = 32 µM) was selected as a lead compound for further optimization due to solubility issue of the former. Importantly, **1512** showed significantly low cancer cell resistance with R-value of 8.2%.[14] (**Figure 3A**) Resistance of CRC tumors towards chemotherapeutic agents is a big problem in the treatment of this type of cancer.[25]

Prior to further optimization of **1512**, it was necessary to conduct a battery of tests to study its selectivity, safety and mechanism of action. Cell line inhibition assays were conducted for **1512** against the CRC cell lines Caco-2 and HT-29 where it displayed consistent potency (IC_50_ values 38 and 35.5 µM respectively), comparable to the previous result against HCT-116 (32.0 µM) (Table 1). Interestingly, this compound did not produce any significant cell growth inhibition against the highly proliferative non-cancerous mouse skin fibroblasts C-166 (**Figure 3A**). The selective inhibition against cancerous over non-cancerous cells, and along with its category 5 safety profile (LD50 > 2000 mg/kg on mice) clearly suggests **1512** as a good candidate for further optimization.

The design rationale of the tested compounds relied on oxidative stress induction within the cancerous cell to mimic the activity of tCA and its synthetic analogues. To establish a proof of this concept, the mechanism of action of compound **1512** was investigated. Several markers indicative of oxidative stress were observed within HCT-116 cells treated with **1512** at the IC_50_ concentration. The ROS-scavenging enzymes superoxide dismutase and catalase activities were found to be significantly diminished (25.1% and 32.7% respectively compared to control). However, the levels of reduced glutathione (GSH) and the lipid oxidation marker malondialdehyde (MDA) were not affected (94.2% and 103.4% respectively compared to the control). To confirm ROS elevation within **1512**-treated cells, DHE staining was utilized. Red fluorescence of DHE was observed, which, along with the previous oxidative stress markers, indicates a strong role of **1512** in the elevation of ROS levels (**Figure 8B**).

HCT-116 cells treated with **1512** at the IC_50_ concentration showed, as a consequence of elevated oxidative stress, apoptotic changes that encompassed loss of cell membrane integrity and fragmentation of the genetic material.[26] Furthermore, flow cytometry indicated a 9-fold accumulation of cells treated with **1512** in the pre-G1 phase compared to the control, also consistent with apoptosis due to genetic material fragmentation.[27] Interestingly, a 3-fold increase in cell accumulation in the G2/M phase and decrease in S-phase accumulation was also observed, indicating a G2/M cell cycle arrest (**Figure 5**). These findings are consistent with previous studies that reported a role for apoptosis in 2’-benzoyloxy-*trans-*cinnamaldehyde (BCA)-induced cancer cell death.[28]

Cyclins constitute critical components of the core cell cycle machinery in eukaryotic cells and their localization in part determines cellular decision to undergo apoptosis in response to DNA damage.[29] In an antibody immunofluorescent assay (**Figure 7**), we found that cyclin B1 protein accumulated in the nucleus of cells that were treated with **1512**. This finding suggests that our compound induced apoptosis that involves nuclear translocation of cyclin B1 and Cyclin D1, and highlights the fact that nuclear accumulation is necessary for cyclin B1- and cyclin D1–dependent apoptosis. This is consistent with findings with previous studies on increased cyclin B1 levels as a marker for apoptosis initiation.[30] Another study which is consistent with our result found that silencing cyclin D1 expression is necessary for the maintenance of the cell cycle exit, and suggests a mechanism that involves inhibition of M-phase entry.[31]

The DNA fragmentation and the nuclear condensation observed by DAPI staining upon treatment with **1512** (**Figure 4A**) led us to suspect that the compound is involved in the increased level of phospho-histone H3. In addition, the ROS elevation and increased oxidative stress observed upon treatment of HCT-116 with **1512** may have resulted in an increased level of cleaved caspase-3 that subsequently led to deactivation of caspase-3 and apoptotic induction of cancer cells (**Figure 7**).[32] These cellular changes indicate that our lead compound **1512** halted the HCT-116 cell cycle at the M phase and prevented cell proliferation.

The promising results from **1512** suggested this compound to be a good starting point for further optimization. We therefore explored the SAR of distance variation between the two aromatic rings by contracting and expanding this distance, resulting in compounds **1612** and **1712** respectively. Both compounds showed almost no activity at concentrations as high as 100 µM against HCT-116 cells, suggesting that the distance between the two aromatic rings in compound **1512** is crucial for activity.

Following, we investigated the SAR of R^1^ and R^2^ positions (**Figure** 2) to optimize the activity of the lead compound **1512**. The distance between both aromatic rings was held rigid and the R^3^ position was fixed as *N*-propyl based on previous results that showed this moiety to be optimal. Erlenmayer chemistry was also utilized to yield 34 chemically diverse oxazolones at positions R^1^ and R^2^ (Table 1, Compounds 1812 to 5112, Entries 23 to 56). These second-generation compounds showed a consistent SAR when tested against the CRC cell lines HCT-116, Caco-2 and HT-29. For instance, the variation in R^2^ position between electron-deficient and electron-withdrawing aromatic moieties demonstrated a strong correlation between electron density over the R^2^ ring and activity. Rings with higher electron density (4-Fluorophenyl, 4-dimethylaminophenyl, 4-hydroxy-3-methoxyphenyl and 3-indolyl) as exemplified by compounds **2312, 2712, 3312, 3712-4112** exerted potent anti-proliferative activities against cancer cells (0.89 to 6.9µM). Some of these compounds exhibited low activity against the non-cancerous cell line C-166 (Table 2). On the other hand, electron-deficient R^2^ rings showed almost no activity against the cancer cell lines, as in compounds **f** (R^2^ = 3-pyridyl) and **g** (R^2^ = 4-nitrophenyl). Notably, unlike R^2^, variations at the R^1^ position affected anticancer activities less significantly. In this regard, all compounds bearing 3-indolyl group at R^2^ were highly active regardless of the structure of the groups at R^1^. Meanwhile, all compounds with electron withdrawing groups at R^2^ (such as 3-pyridyl and 4-nitrophenyl) were much less active, and this was also irrelevant to the structure of R^1^.

**Table 2.**
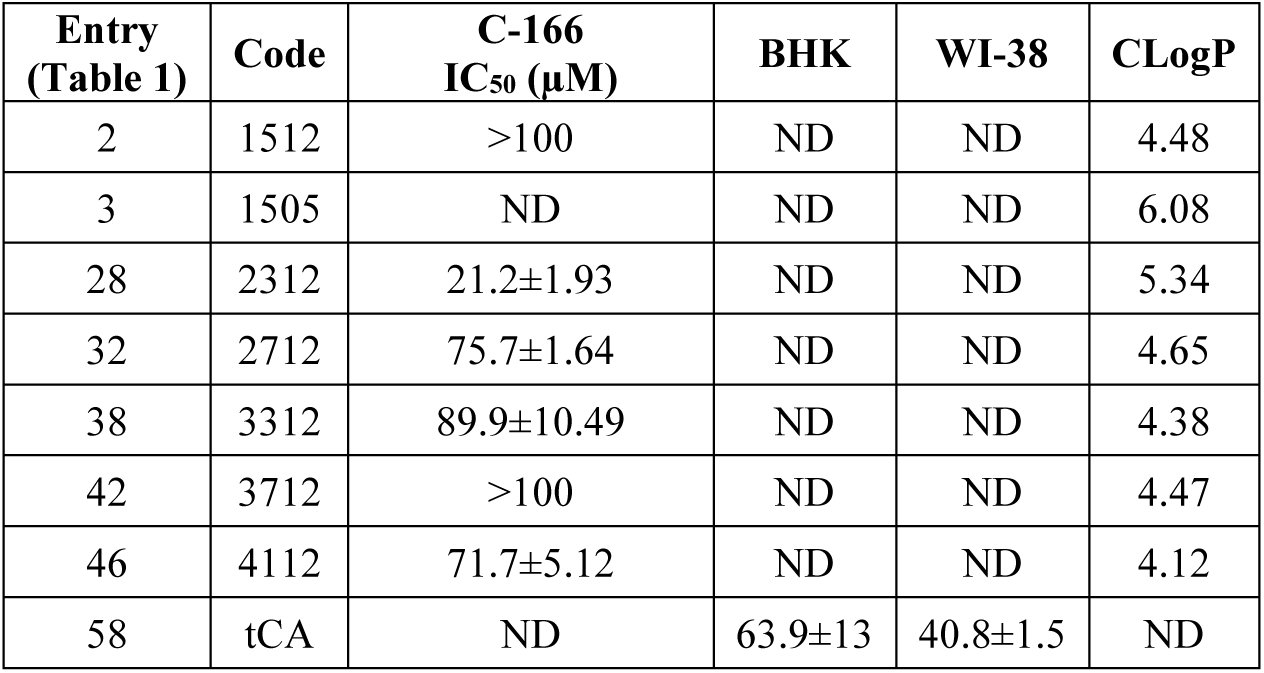
Cytotoxic activities of selected compounds against C-166 cell lines (Mouse skin fibroblast). Note that CLogP is calculated using SYBYL-X molecular modeling package licensed to King Abdulaziz University. ND = Not Determined

| Entry<br>(Table 1) | Code | C-166<br>IC<sub>50</sub> (μM) | BHK | WI-38 | CLogP |
| --- | --- | --- | --- | --- | --- |
| 2 | 1512 | >100 | ND | ND | 4.48 |
| 3 | 1505 | ND | ND | ND | 6.08 |
| 28 | 2312 | 21.2±1.93 | ND | ND | 5.34 |
| 32 | 2712 | 75.7±1.64 | ND | ND | 4.65 |
| 38 | 3312 | 89.9±10.49 | ND | ND | 4.38 |
| 42 | 3712 | >100 | ND | ND | 4.47 |
| 46 | 4112 | 71.7±5.12 | ND | ND | 4.12 |
| 58 | tCA | ND | 63.9±13 | 40.8±1.5 | ND |

The most potent compounds were the 3-indolyl derivatives (**3712** to **4112**) which proved to be the most active series against all the three cell lines. Compounds **3712** and **4112** showed almost comparable potency as doxorubicin (the positive control) on all of the tested cell lines. For instance, Caco-2 IC_50_ for **4112** was 0.89 µM compared to 0.14 µM for doxorubicin. The same compound exhibited about 100 times weaker anti-proliferative effect on C-166 (IC_50_ = 71 µM) compared to Caco2 cancer cell line (0.89 µM). These results give confidence that increasing the potency of these compounds primarily affects cancer cell killing with only marginal effect on normal cell killing.

As previously done for **1512**, compound **4112** and in some other experiments **3712** were used for further biological testing to understand the compounds mechanism of action and/or potential safety. Results were consistent with those of **1512**, while generally being more potent. The cell cycle assay using Annexin-V staining-flow cytometry method (**Figure 5B**) confirmed the apoptotic events upon exposure to **4112**. The compound induced time-dependent increase in cell population entrance to both early and late apoptosis. The apoptotic cell population were less significant after 24 h, compared to control. However, after 48 h, the population of early as well as late apoptotic cells increased from 0.02% and 0.5% to 17.59% and 4.11%, respectively. The cell apoptosis, induced by **4112**, was also confirmed by increased levels of cleaved caspase-3 and its release through the nuclear membrane (**Figure 7**). In agreement with the above mentioned Annexin-V staining assay result, the cleaved caspase 3 increase was time dependent, as revealed by comparing its levels at 24 to that at 48 h. The induction of apoptosis by **4112** was also attributed to increased ROS levels as suggested by DCFDA staining test (**Figure 8c**).

The potential safety of both **4112** and **3712** were evaluated *in vivo* according to OECD guidelines of LD_50_ test, and the results showed that these compounds, like 1512, reposed at category 5 safety (LD50 > 2000 mg/kg on mice).

Recently, models of anticancer activities are increasingly associated with activities of compounds on cancer cell aggregates. Similar to normal stem cells, cancer stem cells (CSCs) reside in niches characterized by hypoxia and low reactive oxygen species (ROS), both of which are critical for maintaining the potential for self-renewal and stemness.[33] Consistently, CSCs show low intracellular ROS levels, suggesting that maintaining a reduced intracellular environment is associated with an undifferentiated state.[34] We therefore tested the effect of **3712** and **4112** on CSCs in a cancer colonosphere formation assay, which is designed to screen for molecules that selectively target colorectal CSCs.[19] As expected, results showed clear concentration-dependent inhibition of sphere size; corroborating results from other experiments. Being, responsible for cancer recurrence after chemotherapy or radiotherapy, CSC targeting via ROS holds a great promise in cancer therapy.

In conclusion, this work presents a model of molecular discovery of promising MA compounds to be further optimized as anticancer agents especially for colorectal cancer. The compounds showed selective toxicity against cancer cell lines and CSC over noncancerous cells, low animal toxicity, and clear cell death mechanism. The two most potent compounds, **3712** and **4112** will be subjected to more SAR investigations as well as antitumor activities in future studies.

## 4. EXPERIMENTAL

All melting points were uncorrected and measured using the capillary melting point instrument BI 9100 (Barnstead Electrothermal, UK). Infrared spectra were recorded on a Thermo Scientific Niccolet iS10 FT-IR Spectrometer (King Fahd Center for Medical Research, King Abdulaziz University, Jeddah, Saudi Arabia). In this report, we only listed the important IR stretching bands, including NH, OH, CH, C=O, C=N and/or C=C. In FT-IR, all samples were measured neat. ^1^H NMR spectra were determined on an AVANCE-III 600 MHz and AVANCE-III HD 850 MHz spectrometers (Bruker, Germany), and chemical shifts are expressed as ppm against TMS as an internal reference (King Fahd Center for Medical Research and Faculty of Science, King Abdulaziz University, Jeddah, Saudi Arabia). LC/MS analyses were performed on an Agilent 6320 Ion Trap HPLC–ESI-MS/DAD (Santa Clara, CA, USA) with the following settings: The analytes were separated using an Macherey-Nagel Nucleodur-C18 column (150 mm length × 4.6 mm i.d., 5 μm) (Macherey-Nagel GMBH & Co. KG, Duren, Germany). Mobile phase; isocratic elution using a mixture of isopropanol and 0.01M ammonium acetate in water (65: 35, v/v). The flow rate was 0.4 mL/min; total run time = 20 min. Purities are reported according to percentage of Peak Areas at wavelength 280 nm. High-resolution mass spectrometry (HRMS) was performed in the Faculty of Science, King Abdulaziz University on Impact II™ Q-TOF spectrometer (Bruker, Germany). Microanalyses were operated using Vario, an Elmentar apparatus (Shimadzu, Japan), Organic Microanalysis Unit, Cairo University, Giza, Egypt. Column chromatography was performed on a silica gel 60 (particle size 0.06 mm - 0.20 mm).

### 4.1. Chemical Synthesis

The previously reported azlactone **3**[35], the final 2-aminopropenamide derivatives **1501**[36], **1502**[35], **1508**[37] and **15EE**[35, 38], and **1712**[14] were prepared according to procedures mentioned below, and their physical and spectral properties were confirmed. Purity of compounds were first assessed qualitatively using Thin Layer Chromatography (TLC), ^1^H NMR and quantitatively using LC/MS (UV detection). The compounds were screened only if purity was confirmed to be above 95%. The compounds subjected to all biological screenings were used as a single *Z* isomer as detected by TLC, LC/MS and NMR.

#### 4.1.1. (Z)-2-cinnamamido-N,3-diphenylacrylamide (1501)[35, 39]

In the reaction tube of the microwave reactor (Milestone™, SynthLab), oxazolone **3** (2 mmol, 0.554 g) and aniline (2 mmol, 0.193 mL) were added to 5 mL of DMF. The reaction was heated to 200°C with stirring for 10 min. After cooling, the mixture was added slowly to ice-cold 1 M HCl. The resulting solid was collected by filtration, washed with water and purified using silica gel chromatography (petroleum ether / DCM / MeOH, gradient). The final compound **1501** was collected as a white solid (0.520 g, 70.6%), Mp 129-132 °C. ^1^H NMR (600 MHz, acetone-*d*_6_) *δ*_H_ ppm 9.50 (1H, br. s), 9.11 (1H, br. s), 7.80 (d, *J* = 7.53 Hz, 2 H), 7.60 - 7.69 (m, 6 H), 7.38 - 7.49 (m, 6 H), 7.30 - 7.37 (m, 4 H), 7.17 (s, 1 H), 7.06 - 7.12 (m, 1 H), 7.02 (d, *J* = 15.81 Hz, 1 H).

#### 4.1.2. (Z)-N-benzyl-2-cinnamamido-3-phenylacrylamide (1502)[35, 38]

Oxazolone **3** (10 mmol, 2.773 g) was dissolved in 50 mL ethanol, benzylamine (20 mmol, 2.18 mL) was then added, and the mixture was stirred for 2 h. The solvent was removed under reduced pressure. The residue was treated with ice-cooled dilute HCl resulting in precipitation of a off-white solid product. The crude product was purified by crystallization from ethanol. The purified product **1502** was an off-white solid, (0.45 g, 60%), Mp. 198-199 °C. ^1^H NMR (600 MHz, Acetone-*d*_6_) δ_H_ ppm 8.08 (br. s., 1 H), 7.57 - 7.68 (m, 5 H), 7.36-7.48 (m, 8 H), 7.29-7.35 (m, 4 H), 7.24 (t, *J =* 7.34 Hz, 1 H), 6.96 (d, *J =* 15.43 Hz, 1 H), 4.52-4.58 (m, 2 H), LC-MS (ESI), RT = 4.4 min, m/z 383.1 [M + H]^+^.

#### 4.1.3. (Z)-2-cinnamamido-N-cyclopentyl-3-phenylacrylamide (1503)

This compound was prepared according to the procedure described for the synthesis of **1502** starting from oxazolone **3** (2 mmol, 0.554 g) and cyclopentylamine (2 mmol, 0.195 mL). The product **1503** was a white solid, (0.51 g, 73%), Mp 239 °C. ^1^H NMR (600 MHz, acetone-*d*_6_) *δ*_H_ ppm 8.85 (s, 1H), 7.59-7.67 (m, 2H), 7.55 (d, *J*=7.53 Hz, 2 H), 7.40-7.47 (m, 3H), 7.37 (t, *J*=7.53 Hz, 2H), 7.28-7.32 (m, 1H), 7.09 (br. s., 1H), 6.96 (d, *J* = 15.81 Hz, 1H), 4.24-4.32 (m, 1 H), 1.90-1.98 (m, 2 H), 1.71 (br. s., 2 H), 1.54-1.62 (m, 4H).

#### 4.1.4. (Z)-N-(1-adamantanyl)-2-cinnamamido-3-phenylacrylamide (1505)

This compound was prepared according to the procedure described for the synthesis of **1502** starting from oxazolone **3** (2 mmol, 0.554 g) and 1-aminoadamantane (2 mmol, 0.308 g). The product **1505** was a white solid (0.75 g, 88%), Mp 272 °C. ^1^H NMR (600 MHz, acetone-*d*_6_) *δ*_H_ ppm 8.86 (s, 1H), 7.62 - 7.68 (m, 3 H), 7.54 (d, *J*=7.53 Hz, 2 H), 7.40 - 7.47 (m, 3 H), 7.37 (t, *J*=7.53 Hz, 2 H), 7.27 - 7.32 (m, 1 H), 7.05 (s, 1 H), 6.98 (d, *J*=15.43 Hz, 1 H),2.17 (m, 1H), 2.13 (br. s., 6 H), 2.00 - 2.03 (m, 1 H), 1.95 - 1.97 (m, 1 H), 1.73 (br. s., 6 H), 1.57 - 1.59 (m, 1 H).

#### 4.1.5. (Z)-2-[(E)-cinnamamido]-3-phenyl-N-(3-pyridyl)acrylamide (1507)

In the reaction tube of a microwave reactor (SynthLab), oxazolone **3** (2 mmol, 0.554 g) and 3-aminopyridine (2 mmol, 0.188 g) were mixed with 5 mL DMF. The reaction was heated to 200°C while stirring in a Milestone microwave reactor for 10 min. After cooling, the mixture was added slowly to ice-cold water. The resulting solid was collected by filtration, washed with water and purified using silica gel chromatography (petroleum ether/ dichloromethane (DCM)/MeOH, gradient) to give **1507** as a white solid (0.206 g, 30%), Mp > 250°C (dec). ^1^H NMR (600 MHz, Acetone-*d*_6_) δ_H_ ppm 9.79 (1H, br s), 9.31 (1H, br s), 8.91 (1H, s), 8.31 (1H, dd, *J* = 6.52, 1.13 Hz), 8.27 (1H, d, *J* = 8.66 Hz), 7.69-7.61 (5H, m), 7.48-7.39 (5H, m), 7.38-7.32 (2H, m), 7.21 (1H, s), 7.03 (1H, d, *J* = 15.81 Hz), LC-MS (ESI), RT = 7.4 min, *m/z* 370.2 [M + H]^+^.

#### 4.1.6. N-[(Z)-3-amino-3-oxo-1-phenylprop-1-en-2-yl]-(E)-cinnamamide (1508)

Oxazolone **3** (2mmol, 0.554 g) was stirred in 5 mL solution of ammonia (2M) in ethanol for 2h. The product was precipitated as white powder which was filtered, washed with water several times followed by ethanol and dichloromethane. Crystallization from aqueous methanol provided a white solid of **1508** Mp >250 °C (dec.) ^1^H NMR (DMSO-*d*6) *d*_H_ ppm 10.11 (br. s., 1H), 7.75 (br. s., 1H), 7.61 (d, *J* = 6.78 Hz, 2H), 7.55 (d, *J* = 7.53 Hz, 2H), 7.35-7.53 (m, 6H), 7.31 (d, *J* = 7.15 Hz, 1H), 7.16 (br. s., 1H), 7.10 (s, 1 H), 6.91 (d, *J* = 15.81 Hz, 1 H).

#### 4.1.7. (Z)-2-[(E)-cinnamamido]-N-(4-methyl-1-piperazinyl)-3-phenyl-acrylamide (1511)

This compound was prepared according to the procedure described for the synthesis of **1502** starting from oxazolone **3** (2 mmol, 0.554 g) and 1-amino-4-methylpiperazine (4 mmol, 0.49 mL). The product **1511** was an off-white power, (305 mg, 39%), Mp. 190 °C; 1H NMR (600 MHz, DMSO-d6) *δ*_H_ ppm 9.83 (s, 1H), 9.58 (s, 1H), 7.62 (d, J = 7.53 Hz, 2H), 7.54 - 7.58 (m, J = 7.91 Hz, 2H), 7.52 (d, J = 15.81 Hz, 1H), 7.44 - 7.48 (m, 2H), 7.39 - 7.44 (m, 3H), 7.30 - 7.35 (m, 1H), 6.91 (d, J = 15.81 Hz, 1H), 6.75 (br. s., 1H), 3.43 (br. s., 2H), 3.14 - 3.22 (m, 4H), 3.10 (br. s., 2H), 2.77 (br. s., 3H).

#### 4.1.8. (Z)-2-[(E)-cinnamamido]-3-phenyl-N-propylacrylamide (1512)

This compound was prepared according to the procedure described for the synthesis of **1502** starting from oxazolone **3** (2 mmol, 0.554 g) and *n*-propylamine (2 mmol, 0.166 mL). The product **1512** was a white solid (542 mg, 81%), Mp 207 °C. ^1^H NMR (850 MHz, DMSO-*d*_6_) δ_H_ 9.69 (s, 1H), 8.12 (t, *J* = 5.97 Hz, 1H), 7.62 (d, *J* = 7.78 Hz, 2H), 7.55 (d, *J* = 7.78 Hz, 2H), 7.50 (d, *J* = 16.09 Hz, 1H), 7.46 (t, *J* = 7.79 Hz, 2H), 7.42 (t, *J* = 7.79 Hz, 1H), 7.38 (t, *J* = 7.79 Hz, 2H), 7.31 (t, *J* = 7.79 Hz, 1H), 7.00 (s, 1H), 6.87 (d, *J* = 16.09 Hz, 1H), 3.12 (q, *J* = 6.75 Hz, 2H), 1.49 (dt, *J* = 7.27 and 6.9 Hz, 2H), 0.88 (t, *J* = 7.53 Hz, 3H); ^13^C NMR (214 MHz, DMSO-*d_6_*) δ 165.1, 164.7, 140.0, 134.8, 134.3, 130.5, 129.9, 129.4, 129.1, 128.6, 128.6, 127.8, 126.9, 121.6, 41.0, 39.8, 39.7, 39.6, 39.5, 39.4, 39.3, 39.2, 22.3, 11.5; LC-MS (ESI), RT = 3.6 min, *m/z* 335.1 [M + H]^+^. Anal. Calcd for (C_21_H_22_N_2_O_2_): C, 75.42; H, 6.63; N, 8.38 Found C, 75.37; H, 6.59; N, 8.29.

#### 4.1.9. (Z)-2-[(E)-cinnamamido]-N-(3-cyanophenyl)-3-phenyl-acrylamide (1513)

This compound was prepared according to the procedure used in the synthesis of compound **1501** starting from the oxazolone **3** (2 mmol, 0.554 g) and 3-aminobenzonitrile (2 mmol, 0.236 g). The product **1513** as a white solid (0.352 g, 47%), Mp 235-237 °C. ^1^H NMR (600 MHz, DMSO-*d*_6_) *δ*_H_ 10.52 (1H, br s), 10.05 (1H, br s), 8.19 (s, 1H), 8.01 (1H, d, *J* = 7.78 Hz), 7.66-7.60 (m, 4H), 7.58-7.50 (m, 3H), 7.48-7.40 (m, 5H), 7.36 (m, 1H), 6.98 (s, 1H), 6.93 (1H, d, *J* = 16.09 Hz).

#### 4.1.10. (Z)-2-[(E)-cinnamamido]-N-(4-fluorophenyl)-3-phenylacrylamide (1528)

This compound was prepared according to the procedure used in the synthesis of compound **1501** starting from the oxazolone **3** (2 mmol, 0.554 g) and 3-fluoroaniline (2 mmol, 0.192 mL). The final compound **1528** was collected as a white solid (0.471 g, 65%), Mp 200°C. ^1^H NMR (600 MHz, DMSO-*d*_6_) *δ*_H_ ppm 10.18 (1H, s), 9.92 (1H, s), 7.74-7.68 (2H, m), 7.62 (4H, t, *J* = 6.96 Hz), 7.51(1H, d, *J* = 15.81 Hz), 7.48-7.39 (7H, m), 7.35 (1H, m,), 7.17 (2H, m), 6.96 (s, 1H), 6.91 (1H, d, *J* = 15.81 Hz); IR (FT-IR, cm^−1^): 3214.5, 3059.3, 2979.2, 1647.5, 1614.2, 1505.4.

#### 4.1.11. (Z)-2-[(E)-cinnamamido]-N-(2-furylmethyl)-3-phenyl-acrylamide (1530)

This compound was prepared according to the procedure described for the synthesis of **1502** starting from oxazolone **3** (2 mmol, 0.554 g) and furfurylamine (2 mmol, 0.176 mL). The product **1530** was an off-white solid, (0.45 g, 60%), Mp 198-199 °C. ^1^H NMR (600 MHz, DMSO-*d*_6_) *δ*_H_ 9.76 (1H, s), 8.62 (1H, t, *J* = 5.8 Hz), 7.63 (2H, m), 7.55-7.60 (3H, m), 7.53 (1H, d, *J* = 15.8 Hz), 7.46 (2H, m), 7.41 (1H, d, *J =* 7.5 Hz), & 0.39 (2H, t, *J =* 7.7 Hz), 7.32 (1H, m), 7.10 (1H, s), 6.88 (1H, d, *J* = 15.8 Hz), 6.41 (1H, m), 6.32 (1H, m), 4.37 (2H, d, *J* = 5.65 Hz); IR (FT-IR, cm^−1^): 3394.3, 3065.2, 2955.9, 1705.5, 1625.7, 1508.2; LC-MS (ESI) RT = 3.6 min, *m/z* 373.1[M + H]^+^. Anal. Calcd for (C_23_H_20_N_2_O_3_): C, 74.18; H, 5.41; N, 7.52; Found C, 73.79; H, 5.08; N, 7.65.

#### 4.1.12. (Z)-2-[(E)-cinnamamido]-N-(2-morpholinoethyl)-3-phenyl-acrylamide (1531)

This compound was prepared according to the procedure described for the synthesis of **1502** starting from oxazolone **3** (2 mmol, 0.554 g) and 2-(4-morpholinyl)ethylamine (5 mmol, 0.65 mL). The product **1531** was an off-white solid, (0.61 g, 75%), Mp 169 °C. ^1^H NMR (600 MHz, DMSO-*d_6_*) *δ*_H_ 9.74 (s, 1H), 7.96 (br s, 1H), 7.63 (d, *J* = 7.15 Hz, 2H), 7.56 (d, *J* = 7.53 Hz, 2H), 7.53 (d, *J* = 15.81 Hz, 1H), 7.37-7.50 (m, 5H), 7.29-7.36 (m, 1H), 7.07 (s, 1H), 6.88 (d, *J* = 16.19 Hz, 1H), 3.52-3.61 (m, 4 H), 3.25 - 3.32 (m, 2 H), 2.37 - 2.47 (m, 6 H). LC-MS (ESI) RT = 3.7 min, *m/z* 406.2 [M + H]^+^.

#### 4.1.13. (Z)-2-[(E)-cinnamamido]-N-(2-hydroxyethyl)-3-phenyl-acrylamide (1532)

Ethanolamine (10 mmol, 0.6 m) was placed in a conical flask and stirred, added Oxazolone **3** (2 mmol, 0.554 g) portion wise while stirring. Reaction was left to go to completion for two hours. The mixture was treated with ice-cooled water containing 10 mL 1M HCl and the precipitated product was collected by filtration. Purification was carried out by crystallization in ethanol to furnish **1532** as a white fluffy solid (0.43 g, 63.9%), Mp 204 °C, ^1^H NMR (850 MHz, DMSO-*d_6_*) δ_H_ 9.72 (s, 1H), 8.06 (t, *J* = 5.71 Hz, 1H), 7.62 (d, *J* = 7.79 Hz, 2H), 7.55 (d, *J* = 7.79 Hz, 2H), 7.51 (d, *J* = 15.57 Hz, 1H), 7.49-7.44 (m, 2 H), 7.42 (d, *J* = 7.27 Hz, 1H), 7.39 (t, *J* = 7.79 Hz, 2H), 7.29-7.34 (m, 1 H), 6.88 (d, *J* = 16.09 Hz, 1 H) 7.05 (s, 1H), 4.65 (t, *J* = 5.71 Hz, 1 H), 3.47 (q, *J* = 5.88 Hz, 2 H), 3.24 (q, *J* = 6.23 Hz, 2 H). LC-MS (ESI) RT = 3.48 min, *m/z* 337.0 [M + H]^+^.

#### 4.1.14. N-[(Z)-3-morpholino-3-oxo-1-phenylprop-1-en-2-yl]cinnamamide (1536)

This compound was prepared according to the procedure described for the synthesis of **1502** starting from oxazolone **3** (2 mmol, 0.554 g) and morpholine (4 mmol, 0.39 mL). The product **1536** was an off-white power, (.514 g, 71%), Mp. 185 °C; 1H NMR (600 MHz, DMSO-d6) δ_H_ 10.07 (s, 1H), 7.56 - 7.63 (m, 6H), 7.40 - 7.48 (m, 6H), 7.27 - 7.36 (m, 1H), 6.93 (d, J = 15.81 Hz, 1H), 6.19 (s, 1H), 3.53 (br. s., 4H), 3.35 (br. s., 4H).

#### 4.1.15. N-[(Z)-3-(1-pyyrrolidinyl)-3-oxo-1-phenylprop-1-en-2-yl]cinnamamide (1555)

This compound was prepared according to the procedure described for the synthesis of **1502** starting from oxazolone **3** (2 mmol, 0.554 g) and pyrrolidine (2mmol, 0.165 mL). The product **1555** was a white power, (0.580 g, 84%), Mp. 194 °C; ^1^H NMR (600 MHz, DMSO-*d*_6_) δ_H_ 10.00 (s, 1H), 7.52 - 7.64 (m, 6H), 7.37 - 7.49 (m, 6H), 7.30 - 7.33 (m, 1H), 6.91 (d, *J* = 15.81 Hz, 1H), 6.32 (s, 1H), 3.58 (t, *J* = 6.02 Hz, 2H), 3.37 (t, *J* = 6.59 Hz, 2H), 1.82 - 1.87 (m, 4H).

#### 4.1.16. (Z)-N-(n-butyl)-2-[(E)-cinnamamido]-3-phenylacrylamide (1556)

This compound was prepared according to the procedure described for the synthesis of **1502** starting from oxazolone **3** (2 mmol, 0.554 g) and *n*-butylamine (2 mmol, 0.176 mL). The product **1556** was an off-white solid, (0.45 g, 60%), Mp 198-199 °C. ^1^H NMR (600 MHz, DMSO-*d_6_*) *δ*_H_ 9.69 (s, 1H), 8.10 (t, *J* = 5.83 Hz, 1H), 7.62 (d, *J* = 7.15 Hz, 2H), 7.56 (d, *J* = 7.53 Hz, 2H), 7.51 (d, *J* = 15.81 Hz, 1H), 7.44 - 7.48 (m, 2H), 7.36 - 7.44 (m, 3H), 7.31 (t, *J* = 7.34 Hz, 1H), 7.01 (s, 1H), 6.88 (d, *J* = 16.19 Hz, 1H), 3.17 (q, *J* = 6.78 Hz, 2H), 1.47 (quin, *J* = 7.25 Hz, 2H), 1.32 (sxt, *J* = 7.38 Hz, 2H), 0.90 (t, *J* = 7.34 Hz, 3H).

#### 4.1.17. (Z)-N-(sec-butyl)-2-cinnamamido-3-phenylacrylamide (1557)

This compound was prepared according to the procedure described for the synthesis of **1502** starting from oxazolone **3** (2 mmol, 0.554 g) and *sec*-butylamine (2 mmol, 0.202 mL). The product **1557** was an off-white solid, (0.51 g, 73%), Mp 229 °C. ^1^H NMR (600 MHz, DMSO-*d_6_*) *δ*_H_ ppm 9.66 (s, 1H), 7.84 (d, *J* = 8.66 Hz, 1H), 7.62 (d, *J* = 7.15 Hz, 2 H), 7.56 (d, *J* = 7.91 Hz, 2H), 7.52 (d, *J* = 15.81 Hz, 1H), 7.49-7.41 (m, 3H), 7.39 (t, *J* = 7.72 Hz, 2H), 7.34-7.28 (m, 1H), 6.94-6.86 (m, 2 H), 3.81 (dt, *J* = 13.93, 7.34 Hz, 1H), 1.39-1.59 (m, 2H), 1.11 (d, *J =* 6.78 Hz, 3H), 0.84-0.93 (m, 3 H).

#### 4.1.18. (Z)-2-[(E)-cinnamamido]-N-ethyl-3-phenylacrylamide (1558)

This compound was prepared according to the procedure described for the synthesis of **1502** starting from oxazolone **3** (2 mmol, 0.554 g) and ethylamine (1 mL of 40% in water). The product **1558** was an off-white solid, (0.39 g, 61%), Mp 190 °C.^1^H NMR (600 MHz, DMSO-*d*_6_) *δ*_H_ ppm 1.08 (t, *J*=7.34 Hz, 3 H) 3.19 (dd, *J*=7.15, 6.02 Hz, 2 H) 6.87 (d, *J*=15.81 Hz, 1 H) 7.02 (s, 1 H) 7.28 - 7.33 (m, 1 H) 7.39 (t, *J*=7.72 Hz, 2 H) 7.43 (d, *J*=7.15 Hz, 1 H) 7.44 - 7.49 (m, 2 H) 7.51 (d, *J*=16.19 Hz, 1 H) 7.55 (d, *J*=7.53 Hz, 2 H) 7.62 (d, *J*=7.15 Hz, 2 H) 8.11 - 8.17 (m, 1 H), 9.68 (s, 1 H).

#### 4.1.19. (Z)-2-cinnamamido-N-ethyl-N-methyl-3-phenylacrylamide (1559)

This compound was prepared according to the procedure described for the synthesis of **1502** starting from oxazolone **3** (2 mmol, 0.554 g) and *N*-ethyl-*N*-Methylamine (2 mmol, 0.171 mL). The product **1559** was a white solid (0.61 g, 91%), Mp 169 °C. ^1^H NMR (600 MHz, DMSO-*d*_6_) *δ*_H_ ppm 10.01 (br. s., 1 H), 7.61 (d, *J*=7.15 Hz, 2 H), 7.52 - 7.59 (m, 3 H), 7.37 - 7.49 (m, 5 H), 7.27 - 7.34 (m, 1 H), 6.94 (d, *J*=15.81 Hz, 1 H), 6.09 - 6.20 (s, 1 H), 3.08 (s, 3H), 2.85 (q, *J* = 7.1 Hz, 2 H), 1.12 (t, *J=* 6.9 Hz, 3H).

#### 4.1.20. Ethyl (Z)-2-cinnamamido-3-phenylacrylate (15EE)

The oxazolone **3** (2 mmol, 0.554 g) was heated under reflux in absolute ethanol in presence of 5 mg of 4-dimethylaminopyridine (DMAP) for 3 hours. The solvent was removed by vacuum evaporation and residue was partitioned between dichloromethane and 1M HCl. The organic layer was washed with water then brine, dried with sodium sulfated and evaporated under vacuum to give the product **15EE** as an off-white solid (0.51 g, 79%), Mp 153 °C. ^1^H NMR (600 MHz, CDCl_3_) *δ*_H_ 8.97 (br. s., 1 H), 7.62 - 7.69 (m, 5 H), 7.34 - 7.50 (m, 6 H), 7.30 (s, 1 H), 6.99 (d, *J* = 15.81 Hz, 1 H), 4.27 (q, *J* = 7.03 Hz, 2 H), 1.31(t, *J* = 7.15 Hz, 3 H).

#### 4.1.21. (Z)-N-(n-propyl)-2-(benzoylamino)-3-phenylacrylamide (1612)

This compound was prepared according to the procedure described for the synthesis of **1502** starting from 4-benzylidene-2-phenyloxazol-5(4H)-one (0.25 g, 1 mmol) and *n*-propylamine (0.16 mL, 2 mmol). The product **1612** was an off-white solid, (0.27 g, 81%), Mp 169 °C. ^1^H NMR (850 MHz, DMSO-*d_6_*) *δ*_H_ 10.17 (br s, 1 H), 8.32 (br s, 1H), 8.00 (d, *J* = 7.27 Hz, 1H), 7.88-7.86 (m, 1H), 7.61-7.55 (m, 1H), 7.53-7.47 (m, 1H), 7.40 (d, *J* = 8.30 Hz, 1H), 7.20-7.28 (m, 2H), 3.11 (m, *J* = 6.75 Hz, 2H), 1.47 (m, 1H), 0.86 (t, *J* = 7.27 Hz, 3H). LC-MS (ESI) RT = 4.52 min, *m/z* 309.0 [M + H]+.

#### 4.1.22. (2Z,4E)-2-cinnamamido-5-phenyl-N-propylpenta-2,4-dienamide (1712)[14]

This compound was prepared according to the procedure described for the synthesis of **1502** starting from (Z)-4-[(E)-3-(phenylallylidene)-2-(E)-styryl]oxazol-5(4H)-one (0.3 g, 1 mmol) and n-propylamine (0.16 mL, 2 mmol). The product **1712** was a light yellow solid, (0.314 g, 87%), Mp 186 °C. ^1^H NMR (850 MHz, DMSO-d_6_) δ_H_ 9.69 (s, 1 H), 8.02 (t, J = 5.71 Hz, 1H), 7.64 (d, J = 7.27 Hz, 2H), 7.55-7.52 (m, 3H), 7.47 - 7.45 (m, 2H), 7.42-7.40 (m, 1H), 7.38-7.35 (m, 2H), 7.31-7.28 (t, J = 7.26 Hz, 1H), 7.05-7.00 (dd, 1H, J = 15.8 and 10.7 Hz), 6.92-6.87 (m, 2H), 6.73 (d, J = 10.90 Hz, 1H), 3.12-3.08 (q, J = 8.6 Hz, 2H), 1.50-1.44 (m, 2H), 0.94 (t, J = 7.53 Hz, 1H), 0.87 (t, J = 7.27 Hz, 3 H), LC-MS (ESI) RT = 5.25 min, m/z 361.1 [M + H]+ (The compound contained 24.2% of E-isomer (2E,4E)-2-cinnamamido-5-phenyl-N-propylpenta-2,4-dienamide according to LC-MS (UV) determination. All data are reported for the Z isomer.

#### 4.1.23. (Z)-2-[(E)-3-(4-chlorophenyl)acrylamido]-3-phenyl-N-propylacrylamide (1812)

This compound was prepared according to the procedure described for the synthesis of **1502** starting from 4-[(*Z*)-benzylidene]-2-[(*E*)-4-chlorostyryl]oxazol-5(4H)-one (0.310 g, 1 mmol) and *n*-propylamine (0.16 mL, 2 mmol). The product **1812** was a white solid (325 mg, 88%), Mp 190-191 °C. IR (KBr, ν_max_ cm^-1^) 3281, 2958, 1651, 1611; ^1^H NMR (850 MHz, DMSO-*d_6_*) δ_H_ 9.71 (br. s., 1H), 8.12 (t, *J* = 5.71 Hz, 1H), 7.64 (d, *J* = 8.30 Hz, 2H), 7.46 - 7.58 (m, 4H), 7.38 (t, *J* = 7.79 Hz, 2H), 7.31 (t, *J* = 7.79 Hz, 1H), 7.00 (s, 1H), 6.87 (d, *J* = 16.09 Hz, 1H), 3.12 (q, *J* = 6.75 Hz, 2H), 1.49 (sxt, *J* = 7.27 Hz, 2H), 0.88 (t, *J* = 7.27 Hz, 3H); ^13^C NMR (214 MHz, DMSO-*d_6_*) δ_C_ 165.1, 164.5, 138.7, 134.3, 133.8, 130.4, 129.5, 129.4, 129.2, 128.7, 128.6, 127.0, 122.4, 41.1, 39.8, 39.7, 39.6, 39.5, 39.4, 39.3, 39.2, 22.4, 11.5; LC/MS (ESI), RT = 4.8 min, m/z 369 (M+1), 370 (M+2).

#### 4.1.24. (Z)-2-[(E)-3-(4-methoxyphenyl)acrylamido]-3-phenyl-N-propylacrylamide (1912)

This compound was prepared according to the procedure described for the synthesis of **1502** starting from 4-[(*Z*)-benzylidene]-2-[(*E*)-4-methoxystyryl]oxazol-5(4H)-one (0.305 g, 1 mmol) and *n*-propylamine (0.16 mL, 2 mmol). The product **1912** was a white solid (303 mg, 83%), Mp 184-185 °C. IR (KBr, ν_max_ cm^-1^) 3229, 2964, 1650, 1625; ^1^H NMR (600 MHz, DMSO-*d_6_*) δ_H_ 9.62 (s, 1H), 8.12 (t, *J* = 5.65 Hz, 1H), 7.51 - 7.62 (m, 4H), 7.45 (d, *J* = 15.43 Hz, 1H), 7.38 (t, *J* = 7.72 Hz, 2H), 7.31 (t, *J* = 7.79 Hz, 1H), 7.02 (d, *J* = 8.66 Hz, 2H), 6.97 (s, 1H), 6.72 (d, *J* = 15.81 Hz, 1H), 3.81 (s, 3H), 3.11 (q, *J* = 6.40 Hz, 2H), 1.48 (sxt, *J* = 7.30 Hz, 2H), 0.87 (t, *J* = 7.34 Hz, 3H); ^13^C NMR (214 MHz, DMSO-*d_6_*) δ_C_ 165.1, 164.9, 160.6, 139.7, 134.4, 130.7, 129.4, 129.3, 128.6, 128.5, 127.3, 126.6, 119.0, 114.5, 55.4, 41.0, 39.8, 39.7, 39.6, 39.5, 39.4, 39.3, 39.2, 22.4, 11.5; LC/MS (ESI), RT = 3.5 min, m/z 365 (M+1).

#### 4.1.25. (Z)-3-phenyl-N-propyl-2-[(E)-3-(p-tolyl)acrylamido]acrylamide (2012)

This compound was prepared according to the procedure described for the synthesis of **1502** starting from 4-[(Z)-benzylidene]-2-[(E)-4-methylstyryl]oxazol-5(4H)-one (0.289 g, 1 mmol) and *n*-propylamine (0.16 mL, 2 mmol). The product **2012** was an off-white solid (321 mg, 92%), Mp 175-176 °C. IR (KBr, ν_max_ cm^-1^) 3296, 3172, 2967, 1646, 1610; ^1^H NMR (850 MHz, DMSO-*d_6_*) δ_H_ 9.65 (br. s., 1H), 8.11 (t, *J* = 5.45 Hz, 1H), 7.48 - 7.57 (m, 4H), 7.46 (d, *J* = 15.57 Hz, 1H), 7.37 - 7.40 (m, 2H), 7.27 - 7.34 (m, 2H), 7.26 (d, *J* = 7.78 Hz, 1H), 6.98 - 7.01 (m, 1H), 6.82 (d, *J* = 15.57 Hz, 1H), 3.09 - 3.13 (m, 2H), 1.48 (td, *J* = 7.20, 14.14 Hz, 2H), 0.87 (t, *J* = 7.27 Hz, 3H); ^13^C NMR (214 MHz, DMSO-*d_6_*) δ_C_ 165.2, 164.9, 140.1, 139.8, 134.4, 132.1, 130.6, 129.8, 129.4, 128.7, 127.9, 127.5, 126.9, 120.6, 41.1, 39.8, 39.7, 39.6, 39.5, 39.4, 39.3, 39.2, 22.5, 21.1, 11.6; LC/MS (ESI), RT = 4.3 min, m/z 349 (M+1).

#### 4.1.26. (Z)-3-phenyl-N-propyl-2-[(E)-3-(thien-2-yl)acrylamido]acrylamide (2112)

This compound was prepared according to the procedure described for the synthesis of **1502** starting from 4-[(Z)-benzylidene]-2-[(E)-2-(thien-3-yl)vinyl]oxazol-5(4H)-one (0.281 g, 1 mmol) and *n*-propylamine (0.16 mL, 2 mmol). The product **2112** was an off-white solid (263 mg, 77%), Mp 195-196 °C. IR (KBr, ν_max_ cm^-1^) 3296, 3172, 2967, 1646, 1610; ^1^H NMR (850 MHz, DMSO-*d_6_*) δ_H_ 9.65 (s, 1H), 8.10 (t, *J* = 5.71 Hz, 1H), 7.61 - 7.68 (m, 2H), 7.53 (d, *J* = 7.79 Hz, 2H), 7.43 (d, *J* = 3.11 Hz, 1H), 7.39 (t, *J* = 7.79 Hz, 2H), 7.31 (t, *J* = 7.79 Hz, 1H), 7.14 (dd, *J* = 3.37, 4.93 Hz, 1H), 6.99 (s, 1H), 6.62 (d, *J* = 15.57 Hz, 1H), 3.11 (q, *J* = 6.75 Hz, 2H), 1.48 (sxt, *J* = 7.27 Hz, 2H), 0.87 (t, *J* = 7.53 Hz, 3H); ^13^C NMR (214 MHz, DMSO-*d_6_*) δ_C_ 165.0, 164.4, 139.8, 134.3, 133.0, 131.3, 130.5, 129.3, 128.6, 128.6, 128.5, 128.5, 126.9, 120.3, 41.0, 39.8, 39.7, 39.6, 39.5, 39.4, 39.3, 39.2, 22.4, 11.5; LC/MS (ESI), RT = 3.4 min, m/z 341 (M+1).

#### 4.1.27. (Z)-2-cinnamamido-3-(4-fluorophenyl)-N-propylacrylamide (2212)

This compound was prepared according to the procedure described for the synthesis of **1502** starting from 4-[(*Z*)-4-fluorobenzylidene]-2-[(*E*)-styryl]oxazol-5(4*H*)-(0.293 g, 1 mmol) and *n*-propylamine (0.16 mL, 2 mmol). The product **2212** was an off-white solid (290 mg, 82%), Mp 192-193 °C. IR (KBr, ν_max_ cm^-1^) 3360, 3026, 2950, 1650, 1610; ^1^H NMR (DMSO-*d*_6_) δ_H_ ppm 9.68 (s, 1 H), 8.12 (s, 1 H), 7.56 - 7.67 (m, 4 H), 7.50 (d, *J*=16.09 Hz, 1 H), 7.44 - 7.47 (m, 2 H), 7.42 (d, *J*=7.78 Hz, 1 H), 7.23 (t, *J* = 8.82 Hz, 2 H), 7.01 (s, 1 H), 6.86 (d, *J* = 16.09 Hz, 1 H), 3.12 (q, *J* = 6.75 Hz, 2 H), 1.45 - 1.52 (m, 2 H), 0.87 (t, *J*=7.53 Hz, 3 H); LC/MS (ESI), RT = 3.9 min, m/z 353 (M+1).

#### 4.1.28. (Z)-2-[(E)-3-(4-chlorophenyl)acrylamido]-3-(4-fluorophenyl)-N- propylacrylamide (2312)

This compound was prepared according to the procedure described for the synthesis of **1502** starting from 2-[(E)-4-chlorostyryl]-4-[(Z)-4-fluorobenzylidene]oxazol-5(4H)-one (0.327 g, 1 mmol) and *n*-propylamine (0.16 mL, 2 mmol). The product **2312** was an off-white solid (341 mg, 88%), Mp 187-188 °C. IR (KBr, ν_max_ cm^-1^) 3360, 3002, 2949, 1650, 1610; ^1^H NMR (850 MHz, DMSO-*d_6_*) δ_H_ 9.69 (s, 1H), 8.12 (t, *J* = 5.71 Hz, 1H), 7.64 (d, *J* = 8.30 Hz, 2H), 7.60 (dd, *J* = 5.71, 8.82 Hz, 2H), 7.46 - 7.55 (m, 3H), 7.20 - 7.26 (m, 2H), 7.01 (s, 1H), 6.85 (d, *J* = 16.09 Hz, 1H), 3.11 (q, *J* = 6.40 Hz, 2H), 1.48 (sxt, *J* = 7.27 Hz, 2H), 0.87 (t, *J* = 7.27 Hz, 3H); ^13^C NMR (214 MHz, DMSO-*d_6_*) δ_C_ 164.9, 164.4, 138.7, 134.3, 133.7, 131.5, 130.9, 130.1, 129.5, 129.2, 125.9, 122.4, 115.6, 115.5, 41.0, 39.8, 39.7, 39.6, 39.5, 39.4, 39.3, 39.2, 22.4, 11.5; LC/MS (ESI), RT = 5.1 min, m/z 387 (M+1), 388 (M+2).

#### 4.1.29. (Z)-3-(4-fluorophenyl)-2-[(E)-3-(4-methoxyphenyl)acrylamido]-N-propylacrylamide (2412)

This compound was prepared according to the procedure described for the synthesis of **1502** starting from 4-[(Z)-4-fluorobenzylidene)]-2-[(E)-4-methoxystyryl]oxazol-5(4H)- one (0.323 g, 1 mmol) and *n*-propylamine (0.16 mL, 2 mmol). The product **2412** was a white solid (355 mg, 93%), Mp 199-200 °C. IR (KBr, ν_max_ cm^-1^) 3285, 2961, 1649, 1608; ^1^H NMR (600 MHz, DMSO-*d_6_*) δ_H_ 9.61 (s, 1H), 8.12 (t, *J* = 5.65 Hz, 1H), 7.56 - 7.61 (m, 4H), 7.45 (d, *J* = 15.81 Hz, 1H), 7.20 - 7.26 (m, 2H), 7.02 (d, *J* = 8.66 Hz, 2H), 6.98 (s, 1H), 6.71 (d, *J* = 15.81 Hz, 1H), 3.81 (s, 3H), 3.11 (q, *J* = 6.65 Hz, 2H), 1.48 (sxt, *J* = 7.30 Hz, 2H), 0.87 (t, *J* = 7.53 Hz, 3H); ^13^C NMR (214 MHz, DMSO-*d_6_*) δ_C_ 165.1, 164.9, 160.7, 139.9, 131.5, 131.0, 130.4, 129.5, 127.3, 125.6, 119.0, 115.6, 115.5, 114.6, 55.4, 41.0, 39.8, 39.7, 39.6, 39.5, 39.4, 39.3, 39.2, 22.4, 11.5; LC/MS (ESI), RT = 3.8 min, m/z 383 (M+1).

#### 4.1.30. (Z)-3-(4-fluorophenyl)-N-propyl-2-[(E)-3-(p-tolyl)acrylamido]acrylamide (2512)

This compound was prepared according to the procedure described for the synthesis of **1502** starting from 4-[(Z)-4-fluorobenzylidene]-2-[(E)-4-methylstyryl]oxazol-5(4H)-one (0.307 g, 1 mmol) and *n*-propylamine (0.16 mL, 2 mmol). The product **2512** was an off-white solid (289 mg, 79%), Mp 164-165 °C. IR (KBr, ν_max_ cm^-1^) 3302, 3208, 2970, 1649, 1640; ^1^H NMR (850 MHz, DMSO-*d_6_*) δ_H_ 9.64 (s, 1H), 8.11 (t, *J* = 5.71 Hz, 1H), 7.60 (dd, *J* = 5.71, 8.82 Hz, 2H), 7.50 (d, *J* = 8.30 Hz, 2H), 7.46 (d, *J* = 16.09 Hz, 1H), 7.26 d, *J* = 8.30 Hz, 2H), 7.20 - 7.25 (m, 2H), 7.00 (s, 1H), 6.81 (d, *J* = 15.57 Hz, 1H), 3.12 (q, *J* = 6.75 Hz, 2H), 2.35 (s, 3H), 1.48 (sxt, *J* = 7.27 Hz, 2H), 0.87 (t, *J* = 7.27 Hz, 3H); ^13^C NMR (214 MHz, DMSO-*d_6_*) δ_C_ 165.0, 164.8, 140.0, 139.7, 132.0, 131.5, 131.0, 130.3, 129.7, 127.8, 125.7, 120.5, 115.6, 115.5, 41.0, 39.8, 39.7, 39.6, 39.5, 39.4, 39.3, 39.2, 22.4, 21.1, 11.5; LC/MS (ESI), RT = 3.7 min, m/z 367 (M+1).

#### 4.1.31. (Z)-3-(4-fluorophenyl)-N-propyl-2-[(E)-3-(thien-2-yl)acrylamido]acrylamide (2612)

This compound was prepared according to the procedure described for the synthesis of **1502** starting from 4-([(Z)-4-fluorobenzylidene]-2-[(E)-2-(thien-3-yl)vinyl]oxazol-5(4H)-one (0.299 g, 1 mmol) and *n*-propylamine (0.16 mL, 2 mmol). The product **2612** was a yellow solid (279 mg, 78%), Mp 142-143 °C. IR (KBr, ν_max_ cm^-1^) 3302, 3143, 2961, 1650, 1610; ^1^H NMR (850 MHz, DMSO-*d*_6_) δ ppm 9.63 (s, 1 H) 8.11 (t, *J*=5.71 Hz, 1 H) 7.61 - 7.68 (m, 2 H) 7.58 (dd, *J*=8.04, 5.97 Hz, 2 H) 7.43 (d, *J*=3.11 Hz, 1 H) 7.23 (t, *J*=8.56 Hz, 2 H) 7.11 - 7.16 (m, 1 H) 7.00 (s, 1 H) 6.60 (d, *J*=15.57 Hz, 1 H) 3.11 (q, *J*=6.40 Hz, 2 H) 1.48 (sxt, *J*=7.27 Hz, 2 H) 0.87 (t, *J*=7.53 Hz, 3 H); ^13^C NMR (214 MHz, DMSO-*d_6_*) δ_C_ 165.0, 164.4, 139.8, 133.1, 131.6, 131.5, 131.4, 130.9, 130.2, 128.6, 128.5, 126.3, 125.9, 120.3, 41.0, 39.8, 39.7, 39.6, 39.5, 39.4, 39.3, 39.2, 22.4, 11.5; LC/MS (ESI), RT = 3.5 min, m/z 359 (M+1).

#### 4.1.32. (Z)-2-cinnamamido-3-[4-(dimethylamino)phenyl]-N-propylacrylamide (2712)

This compound was prepared according to the procedure described for the synthesis of **1502** starting from 4-[(Z)-4-(dimethylamino)benzylidene]-2-[(E)-styryl]oxazol-5(4H)-one (0.318 g, 1 mmol) and *n*-propylamine (0.16 mL, 2 mmol). The product **2712** was an off-white solid (328 mg, 87%), Mp 207-208 °C. IR (KBr, ν_max_ cm^-1^) 3087, 2933, 1659; ^1^H NMR (600 MHz, DMSO-*d_6_*) δ_H_ 9.51 (s, 1H), 7.91 (t, *J* = 5.84 Hz, 1H), 7.63 (d, *J* = 7.15 Hz, 2H), 7.45 - 7.52 (m, 3H), 7.40 - 7.44 (m, 3H), 7.04 (s, 1H), 6.90 (d, *J* = 15.81 Hz, 1H), 6.70 (d, *J* = 9.03 Hz, 2H), 3.11 (q, *J* = 6.75 Hz, 2H), 2.92 (s, 6H), 1.46 (sxt, *J* = 7.23 Hz, 2H), 0.86 (t, *J* = 7.34 Hz, 3H); ^13^C NMR (214 MHz, DMSO-*d_6_*) δ_C_ 169.6, 168.0, 142.5, 141.1, 135.8, 134.7, 131.2, 129.3, 129.0, 128.7, 128.1, 127.5, 126.3, 111.0, 41.6, 40.7, 39.8, 39.7, 39.6, 39.5, 39.4, 39.3, 39.2, 22.3, 11.4; LC/MS (ESI), RT = 3.8 min, m/z 378 (M+1).

#### 4.1.33. (Z)-2-[(E)-3-(4-chlorophenyl)acrylamido]-3-[4-(dimethylamino)phenyl]-N-propylacrylamide (2812)

This compound was prepared according to the procedure described for the synthesis of **1502** starting from 2-[(*E*)-4-chlorostyryl]-4-[(*Z*)-4-(dimethylamino)benzylidene]oxazol- 5(4H)-one (0.353 g, 1 mmol) and *n*-propylamine (0.16 mL, 2 mmol). The product **2812** was a yellow solid (371 mg, 90%), Mp 243-244 °C. IR (KBr, ν_max_ cm^-1^) 3067, 2912, 1648, 1609; ^1^H NMR (600 MHz, DMSO-*d_6_*) δ_H_ 9.53 (s, 1H), 7.92 (t, *J* = 6.02 Hz, 1H), 7.65 (d, *J* = 8.66 Hz, 2H), 7.53 (d, *J* = 8.28 Hz, 2H), 7.50 (d, *J* = 15.81 Hz, 1H), 7.42 (d, *J* = 9.03 Hz, 2H), 7.04 (s, 1H), 6.90 (d, *J* = 15.81 Hz, 1H), 6.69 (d, *J* = 9.03 Hz, 2H), 3.10 (q, *J* = 6.40 Hz, 2H), 2.93 (s, 6H), 1.46 (sxt, *J* = 7.23 Hz, 2H), 0.86 (t, *J* = 7.34 Hz, 3H); ^13^C NMR (214 MHz, DMSO-*d_6_*) δ_C_ 166.5, 163.1, 140.1, 137.9, 134.6, 133.4, 131.2, 129.4, 129.3, 128.8, 128.6, 128.2, 127.8, 127.5, 41.3, 40.5, 39.8, 39.7, 39.6, 39.5, 39.4, 39.3, 39.2, 22.4, 11.4; LC/MS (ESI), RT = 5.9 min, m/z 412 (M+1), 413 (M+2).

#### 4.1.34. (Z)-3-[4-(dimethylamino)phenyl]-2-[(E)-3-(4-methoxyphenyl)acrylamido]-N-propylacrylamide (2912)

This compound was prepared according to the procedure described for the synthesis of **1502** starting from 4-[(Z)-4-(dimethylamino)benzylidene]-2-[(E)-4-methoxystyryl]-oxazol-5(4H)-one (0.348 g, 1 mmol) and *n*-propylamine (0.16 mL, 2 mmol). The product **2912** was a yellow solid (346 mg, 85%), Mp 236-237 °C. IR (KBr, ν_max_ cm^-1^) 3070, 2952, 1647, 1603; ^1^H NMR (600 MHz, DMSO-*d_6_*) δ_H_ 9.41 (s, 1H), 7.88 (t, *J* = 5.83 Hz, 1H), 7.58 (d, *J* = 9.04 Hz, 2H), 7.40 - 7.47 (m, 3H), 7.01 - 7.04 (m, 3H), 6.75 (d, *J* = 15.81 Hz, 1H), 6.69 (d, *J* = 9.03 Hz, 2H), 3.81 (s, 3H), 3.10 (q, *J* = 6.40 Hz, 2H), 2.92 (s, 6H), 1.46 (sxt, *J* = 7.30 Hz, 2H), 0.86 (t, *J* = 7.34 Hz, 3H); ^13^C NMR (214 MHz, DMSO-*d_6_*) δ_C_ 164.3, 163.1, 158.6, 142.3, 134.5, 133.1, 131.9, 131.4, 129.4, 128.6, 119.0, 114.6, 113.9, 113.2, 55.3, 41.5, 40.5, 39.8, 39.7, 39.6, 39.5, 39.4, 39.3, 39.2, 22.1, 11.3; LC/MS (ESI), RT = 3.7 min, m/z 408 (M+1).

#### 4.1.35. (Z)-3-[4-(dimethylamino)phenyl]-N-propyl-2-[(E)-3-(p-tolyl)acrylamido]acrylamide (3012)

This compound was prepared according to the procedure described for the synthesis of **1502** starting from 4-[(Z)-4-(dimethylamino)benzylidene]-2-[(E)-4-methylstyryl]oxazol-5(4H)-one (0.332 g, 1 mmol) and *n*-propylamine (0.16 mL, 2 mmol). The product **3012** was a yellow solid (341 mg, 87%), Mp 218-219 °C. IR (KBr, ν_max_ cm^-1^) 3268, 2952, 1647, 1601; ^1^H NMR (850 MHz, DMSO-*d*_6_) δ ppm 9.51 (br. s., 1 H) 7.96 (t, *J*=5.97 Hz, 1 H) 7.45 - 7.52 (m, 3 H) 7.26 (m, *J*=7.78 Hz, 2 H) 7.21 (s, 1 H) 7.04 (s, 1 H) 6.99 (d, *J*=8.30 Hz, 1 H) 6.84 (d, *J*=15.57 Hz, 1 H) 6.76 (d, *J*=8.30 Hz, 1 H) 3.11 (q, *J*=6.57 Hz, 2 H) 2.51 (br. s., 6 H) 2.34 (s, 3H) 1.47 (sxt, *J*=7.16 Hz, 2 H) 0.85 - 0.87 (m, 3 H); LC/MS (ESI), RT = 5.0 min, m/z 392 (M+1).

#### 4.1.36. (Z)-3-[4-(dimethylamino)phenyl]-N-propyl-2-[(E)-3-(thien-2-yl)acrylamido]acrylamide (3112)

This compound was prepared according to the procedure described for the synthesis of **1502** starting from 4-[(Z)-4-(dimethylamino)benzylidene]-2-[(E)-2-(thien-3-yl)vinyl]-oxazol-5(4H)-one (0.324 g, 1 mmol) and *n*-propylamine (0.16 mL, 2 mmol). The product **3112** was an off-white solid (352 mg, 92%), Mp 208-209 °C. IR (KBr, ν_max_ cm^-1^) 3296, 2948, 1643, 1605; ^1^H NMR (600 MHz, DMSO-*d_6_*) δ_H_ 9.47 (s, 1H), 7.90 (t, *J* = 5.83 Hz, 1H), 7.61 - 7.68 (m, 2H), 7.44 (d, *J* = 3.39 Hz, 1H), 7.40 (d, *J* = 9.04 Hz, 2H), 7.15 (dd, *J* = 3.58, 5.08 Hz, 1H), 7.03 (s, 1H), 6.70 (d, *J* = 9.03 Hz, 2H), 6.64 (d, *J* = 15.81 Hz, 1H), 3.09 (q, *J* = 6.40 Hz, 2H), 2.93 (s, 6H), 1.46 (sxt, *J* = 7.30 Hz, 2H), 0.85 (t, *J* = 7.34 Hz, 3H); LC/MS (ESI), RT = 3.5 min, m/z 384 (M+1).

#### 4.1.37. (Z)-2-cinnamamido-3-(4-hydroxy-3-methoxyphenyl)-N-propylacrylamide (3212)

This compound was prepared according to the procedure described for the synthesis of **1502** starting from 4-[(Z)-4-hydroxy-3-methoxybenzylidene]-2-[(E)-styryl]oxazol-5(4H)-one (0.321 g, 1 mmol) and *n*-propylamine (0.16 mL, 2 mmol). The product **3212** was an off-white solid (323 mg, 85%), Mp 199-200 °C. IR (KBr, ν_max_ cm^-1^) 3292, 2971, 1650, 1610; ^1^H NMR (600 MHz, DMSO-*d_6_*) δ_H_ 8.02 (br. s., 1H), 7.61 (d, *J* = 7.15 Hz, 2H), 7.39 - 7.55 (m, 4H), 7.21 (s, 1H), 7.04 (s, 1H), 6.91 (d, *J* = 16.19 Hz, 1H), 6.77 (d, *J* = 8.28 Hz, 1H), 3.69 (s, 3H), 3.11 (q, *J* = 6.65 Hz, 2H), 1.47 (sxt, *J* = 7.15 Hz, 2H), 0.86 (t, *J* = 7.34 Hz, 3H); ^13^C NMR (214 MHz, DMSO-*d_6_*) δ_C_ 165.2, 164.8, 147.5, 147.3, 139.8, 134.9, 129.9, 129.2, 128.6, 127.7, 127.2, 125.5, 123.8, 121.8, 115.5, 112.9, 55.4, 41.0, 39.8, 39.7, 39.6, 39.5, 39.4, 39.3, 39.2, 22.5, 11.5; LC/MS (ESI), RT = 2.9 min, m/z 381 (M+1).

#### 4.1.38. (Z)-2-[(E)-3-(4-chlorophenyl)acrylamido]-3-(4-hydroxy-3-methoxyphenyl)-N-propylacrylamide (3312)

This compound was prepared according to the procedure described for the synthesis of **1502** starting from 2-[(E)-4-chlorostyryl]-4-[(Z)-4-hydroxy-3-methoxybenzylidene]-oxazol-5(4H)-one (0.355 g, 1 mmol) and *n*-propylamine (0.16 mL, 2 mmol). The product **3312** was a yellow solid (356 mg, 86%), Mp 136-137 °C. IR (KBr, ν_max_ cm^-1^) 3566, 2988, 2902, 1650; ^1^H NMR (600 MHz, DMSO-*d_6_*) δ_H_ 9.61 (br. s., 1H), 8.01 (t, *J* = 5.83 Hz, 1H), 7.64 (d, *J* = 8.28 Hz, 2H), 7.48 - 7.56 (m, 3H), 7.20 (d, *J* = 2.26 Hz, 1H), 7.04 (s, 1H), 6.99 (dd, *J* = 1.88, 8.28 Hz, 1H), 6.90 (d, *J* = 16.19 Hz, 1H), 6.76 (d, *J* = 8.28 Hz, 1H), 3.68 (s, 3H), 3.11 (q, *J* = 6.65 Hz, 2H), 1.47 (sxt, *J* = 7.23 Hz, 2H), 0.86 (t, *J* = 7.34 Hz, 3H); LC/MS (ESI), RT = 3.4 min, m/z 415 (M+1), 416 (M+2).

#### 4.1.39. (Z)-3-(4-hydroxy-3-methoxyphenyl)-2-[(E)-3-(4-methoxyphenyl)acrylamido]-N-propylacrylamide (3412)

This compound was prepared according to the procedure described for the synthesis of **1502** starting from 4-[(Z)-4-hydroxy-3-methoxybenzylidene]-2-[(E)-4-methoxystyryl]-oxazol-5(4H)-one (0.351 g, 1 mmol) and *n*-propylamine (0.16 mL, 2 mmol). The product **3412** was an orange solid (332 mg, 81%), Mp 180-181 °C. IR (KBr, ν_max_ cm^-1^) 3567, 2989, 2955, 1655; ^1^H NMR (600 MHz, DMSO-*d_6_*) δ_H_ 9.49 (br. s., 1H), 7.97 (t, *J* = 6.02 Hz, 1H), 7.56 (d, *J* = 8.28 Hz, 2H), 7.46 (d, *J* = 15.81 Hz, 1H), 7.21 (d, *J* = 1.88 Hz, 1H), 7.00 - 7.03 (m, 3H), 6.99 (dd, *J* = 1.88, 8.28 Hz, 1H), 6.77 (d, *J* = 6.78 Hz, 1H), 6.75 (s, 1H), 3.81 (s, 3H), 3.68 (s, 3H), 3.11 (q, *J* = 6.65 Hz, 2H), 1.47 (sxt, *J* = 7.23 Hz, 2H), 0.86 (t, *J* = 7.34 Hz, 3H); LC/MS (ESI), RT = 2.7 min, m/z 411 (M+1).

#### 4.1.40. (Z)-3-(4-hydroxy-3-methoxyphenyl)-N-propyl-2-[(E)-3-(p-tolyl)acrylamido]acrylamide (3512)

This compound was prepared according to the procedure described for the synthesis of **1502** starting from 4-[(Z)-4-hydroxy-3-methoxybenzylidene]-2-[(E)-4-methylstyryl]-oxazol-5(4H)-one (0.335 g, 1 mmol) and *n*-propylamine (0.16 mL, 2 mmol). The product **3512** was a yellow solid (296 mg, 75%), Mp 183-184 °C. IR (KBr, ν_max_ cm^-1^) 3566, 3213, 2947, 1663; ^1^H NMR (600 MHz, DMSO-*d_6_*) δ_H_ 9.54 (s, 1H), 9.39 (br. s., 1H), 8.00 (t, *J* = 5.65 Hz, 1H), 7.44 - 7.53 (m, 3H), 7.27 (d, *J* = 7.91 Hz, 2H), 7.21 (s, 1H), 7.03 (s, 1H), 6.99 (dd, *J* = 1.88, 8.28 Hz, 1H), 6.85 (d, *J* = 15.81 Hz, 1H), 6.76 (d, *J* = 8.28 Hz, 1H), 3.68 (s, 3H), 3.10 (q, *J* = 6.65 Hz, 2H), 2.35 (s, 3H), 1.47 (sxt, *J* = 7.23 Hz, 2H), 0.86 (t, *J* = 7.34 Hz, 3H); ^13^C NMR (214 MHz, DMSO-*d_6_*) δ_C_ 165.1, 164.9, 147.5, 147.3, 139.7, 139.6, 132.1, 129.7, 128.5, 127.7, 127.3, 125.5, 123.7, 120.8, 115.4, 112.9, 55.3, 41.0, 39.8, 39.7, 39.6, 39.5, 39.4, 39.3, 39.2, 22.5, 11.5; LC/MS (ESI), RT = 3.3 min, m/z 395 (M+1).

#### 4.1.41. (Z)-3-(4-hydroxy-3-methoxyphenyl)-N-propyl-2-[(E)-3-(thien-2-yl)acrylamido]acrylamide (3612)

This compound was prepared according to the procedure described for the synthesis of **1502** starting from 4-[(Z)-4-hydroxy-3-methoxybenzylidene]-2-[(E)-2-(thien-3-yl)vinyl]-oxazol-5(4H)-one (0.327 g, 1 mmol) and *n*-propylamine (0.16 mL, 2 mmol). The product **3612** was a yellow solid (340 mg, 88%), Mp 192-93 °C. IR (KBr, ν_max_ cm^-1^) 3560, 3105, 2954, 1648, 1603; ^1^H NMR (600 MHz, DMSO-*d_6_*) δ_H_ 9.54 (br. s., 1H), 7.99 (t, *J* = 5.83 Hz, 1H), 7.63 - 7.68 (m, 2H), 7.43 (d, *J* = 3.39 Hz, 1H), 7.19 (d, *J* = 1.88 Hz, 1H), 7.13 - 7.16 (m, 1H), 7.04 (s, 1H), 6.98 (dd, *J* = 1.88, 8.28 Hz, 1H), 6.76 (d, *J* = 7.91 Hz, 1H), 6.66 (d, *J* = 15.81 Hz, 1H), 3.69 (s, 3H), 3.11 (q, *J* = 6.65 Hz, 2H), 1.46 (sxt, *J* = 7.23 Hz, 2H), 0.86 (t, *J* = 7.34 Hz, 3H); LC/MS (ESI), RT = 2.7 min, m/z 387 (M+1).

#### 4.1.42. (Z)-2-cinnamamido-3-(1H-indol-3-yl)-N-propylacrylamide (3712)

This compound was prepared according to the procedure described for the synthesis of **1502** starting from (Z)-4-[(1H-indol-3-yl)methylene]-2-[(E)-styryl]oxazol-5(4H)-one (0.314 g, 1 mmol) and *n*-propylamine (0.16 mL, 2 mmol). The product **3712** was a white solid (287 mg, 77%), Mp 249-250 °C. IR (KBr, ν_max_ cm^-1^) 3401, 3057, 2956, 1650, 1608; ^1^H NMR (600 MHz, DMSO-*d_6_*) δ_H_ 11.56 (br. s., 1H), 9.46 (s, 1H), 7.98 (t, *J* = 5.83 Hz, 1H), 7.74 (d, *J* = 7.91 Hz, 1H), 7.63 - 7.67 (m, 3H), 7.42 - 7.55 (m, 6H), 7.17 (t, *J* = 7.53 Hz, 1H), 7.12 (t, *J* = 7.53 Hz, 1H), 6.98 (d, *J* = 15.81 Hz, 1H), 3.14 (q, *J* = 6.53 Hz, 2H), 1.50 (sxt, *J* = 7.23 Hz, 2H), 0.88 (t, *J* = 7.34 Hz, 3H); ^13^C NMR (214 MHz, DMSO-*d_6_*) δ_C_ 164.5, 164.2, 141.5, 135.5, 134.8, 129.6, 129.3, 129.1, 129.0, 128.7, 127.9, 124.9, 124.6, 122.1, 120.0, 118.2, 111.9, 110.0, 41.0, 39.8, 39.7, 39.6, 39.5, 39.4, 39.3, 39.2, 22.3, 11.5; LC/MS (ESI), RT = 3.3 min, m/z 374 (M+1).

#### 4.1.43. (Z)-2-[(E)-3-(4-chlorophenyl)acrylamido]-3-(1H-indol-3-yl)-N-propylacrylamide (3812)

This compound was prepared according to the procedure described for the synthesis of **1502** starting from (Z)-4-[(1H-indol-3-yl)methylene]-2-[(E)-4-chlorostyryl]oxazol-5(4H)-one (0.348 g, 1 mmol) and *n*-propylamine (0.16 mL, 2 mmol). The product **3812** was an orange solid (330 mg, 81%), Mp 210-211 °C. IR (KBr, ν_max_ cm^-1^) 3208, 1650, 1615; ^1^H NMR (600 MHz, DMSO-*d_6_*) δ_H_ 11.56 (br. s., 1H), 9.47 (s, 1H), 7.98 (t, *J* = 5.83 Hz, 1H), 7.73 (d, *J* = 7.91 Hz, 1H), 7.68 (d, *J* = 8.66 Hz, 2H), 7.65 (d, *J* = 2.63 Hz, 1H), 7.50 - 7.56 (m, 4H), 7.43 (d, *J* = 7.91 Hz, 1H), 7.17 (t, *J* = 7.15 Hz, 1H), 7.12 (t, *J* = 7.34 Hz, 1H), 6.98 (d, *J* = 15.81 Hz, 1H), 3.14 (q, *J* = 6.53 Hz, 2H), 1.50 (sxt, *J* = 7.30 Hz, 2H), 0.88 (t, *J* = 7.53 Hz, 3H); ^13^C NMR (214 MHz, DMSO-*d_6_*) δ_C_ 165.7, 165.3, 140.5, 138.1, 135.7, 134.2, 132.3, 132.2, 130.7, 130.6, 130.5, 130.1, 129.8, 125.1, 123.9, 122.7, 113.7, 111.5, 110.1, 41.6, 39.8, 39.7, 39.6, 39.5, 39.4, 39.3, 39.2, 23.0, 11.6; LC/MS (ESI), RT = 4.2 min, m/z 408 (M+1), 409 (M+2).

#### 4.1.44. (Z)-3-(1H-indol-3-yl)-2-[(E)-3-(4-methoxyphenyl)acrylamido]-N-propylacrylamide (3912)

This compound was prepared according to the procedure described for the synthesis of **1502** starting from (Z)-4-[(1H-indol-3-yl)methylene]-2-[(E)-4-methoxystyryl]oxazol-5(4H)-one (0.344 g, 1 mmol) and *n*-propylamine (0.16 mL, 2 mmol). The product **3912** was an orange solid (335 mg, 83%), Mp 225-226 °C. IR (KBr, ν_max_ cm^-1^) 3181, 1650, 1615; ^1^H NMR (600 MHz, DMSO-*d_6_*) δ_H_ 11.55 (br. s., 1H), 9.36 (s, 1H), 7.95 (t, *J* = 5.83 Hz, 1H), 7.73 (d, *J* = 7.91 Hz, 1H), 7.64 (d, *J* = 2.26 Hz, 1H), 7.60 (d, *J* = 8.66 Hz, 2H), 7.46 - 7.50 (m, 2H), 7.43 (d, *J* = 7.91 Hz, 1H), 7.17 (t, *J* = 7.53 Hz, 1H), 7.12 (t, *J* = 7.53 Hz, 1H), 7.03 (d, *J* = 8.66 Hz, 2H), 6.84 (d, *J* = 15.81 Hz, 1H), 3.82 (s, 3H), 3.14 (q, *J* = 6.27 Hz, 2H), 1.49 (sxt, *J* = 7.23 Hz, 2H), 0.88 (t, *J* = 7.34 Hz, 3H); ^13^C NMR (214 MHz, DMSO-*d_6_*) δ_C_ 166.0, 164.3, 160.6, 141.5, 138.5, 137.1, 132.3, 131.9, 131.4, 129.2, 123.5, 122.2, 121.0, 120.6, 114.6, 113.9, 112.5, 112.3, 55.8, 41.5, 39.8, 39.7, 39.6, 39.5, 39.4, 39.3, 39.2, 22.6, 11.5; LC/MS (ESI), RT = 3.2 min, m/z 404 (M+1).

#### 4.1.45. (Z)-3-(1H-indol-3-yl)-N-propyl-2-[(E)-3-(p-tolyl)acrylamido]acrylamide (4012)

This compound was prepared according to the procedure described for the synthesis of **1502** starting from (Z)-4-[(1H-indol-3-yl)methylene]-2-[(E)-4-methylstyryl]oxazol-5(4H)-one (0.328 g, 1 mmol) and *n*-propylamine (0.16 mL, 2 mmol). The product **4012** was a yellow solid (345 mg, 89%), Mp 192-193 °C. IR (KBr, ν_max_ cm^-1^) 3220, 2950, 1651, 1595; ^1^H NMR (600 MHz, DMSO-*d_6_*) δ_H_ 11.56 (br. s., 1H), 9.42 (s, 1H), 7.97 (t, *J* = 5.83 Hz, 1H), 7.73 (d, *J* = 7.91 Hz, 1H), 7.65 (s, 1H), 7.54 (d, *J* = 7.91 Hz, 2H), 7.47 - 7.52 (m, 2H), 7.43 (d, *J* = 7.91 Hz, 1H), 7.28 (d, *J* = 7.91 Hz, 2H), 7.17 (t, *J* = 7.53 Hz, 1H), 7.12 (t, *J* = 7.53 Hz, 1H), 6.93 (d, *J* = 15.81 Hz, 1H), 3.14 (q, *J* = 6.40 Hz, 2H), 2.36 (s, 3H), 1.50 (sxt, *J* = 7.30 Hz, 2H), 0.88 (t, *J* = 7.53 Hz, 3H); ^13^C NMR (214 MHz, DMSO-*d_6_*) δ_C_ 165.1, 164.6, 141.5, 139.6, 135.5, 129.8, 129.6, 129.2, 127.9, 127.7, 127.6, 122.2, 121.3, 121.1, 118.2, 112.5, 111.9, 110.0, 40.9, 39.8, 39.7, 39.6, 39.5, 39.4, 39.3, 39.2, 22.6, 21.0, 11.5; LC/MS (ESI), RT = 3.7 min, m/z 388 (M+1).

#### 4.1.46. (Z)-3-(1H-indol-3-yl)-N-propyl-2-[(E)-3-(thien-2-yl)acrylamido]acrylamide (4112)

This compound was prepared according to the procedure described for the synthesis of **1502** starting from (Z)-4-[(1H-indol-3-yl)methylene]-2-[(E)-2-(thien-3-yl)vinyl]oxazol-5(4H)-one (0.320 g, 1 mmol) and *n*-propylamine (0.16 mL, 2 mmol). The product **4112** was a yellow solid (326 mg, 86%), Mp 244-245 °C. IR (KBr, ν_max_ cm^-1^) 3567, 3997, 2950, 1650; ^1^H NMR (600 MHz, DMSO-*d_6_*) δ_H_ 11.56 (br. s., 1H), 9.41 (s, 1H), 7.96 (t, *J* = 5.83 Hz, 1H), 7.72 (d, *J* = 7.91 Hz, 1H), 7.65 - 7.70 (m, 2H), 7.64 (d, *J* = 1.88 Hz, 1H), 7.50 (s, 1H), 7.42 - 7.46 (m, 2H), 7.14 - 7.19 (m, 2H), 7.12 (t, *J* = 7.53 Hz, 1H), 6.73 (d, *J* = 15.43 Hz, 1H), 3.13 (q, *J* = 6.40 Hz, 2H), 1.49 (sxt, *J* = 7.23 Hz, 2H), 0.88 (t, *J* = 7.53 Hz, 3H); ^13^C NMR (214 MHz, DMSO-*d_6_*) δ_C_ 163.8, 163.2, 139.7, 138.1, 136.5, 136.2, 133.4, 133.3, 132.4, 129.1, 128.3, 123.6, 122.9, 122.2, 121.1, 120.6, 112.3, 107.4, 41.4, 39.8, 39.7, 39.6, 39.5, 39.4, 39.3, 39.2, 22.3, 11.3; LC/MS (ESI), RT = 3.1 min, m/z 380 (M+1).

#### 4.1.47. (Z)-2-cinnamamido-N-propyl-3-(pyridin-3-yl)acrylamide (4212)

This compound was prepared according to the procedure described for the synthesis of **1502** starting from (Z)-4-(pyridin-3-ylmethylene)-2-[(E)-styryl]oxazol-5(4H)-one (0.276 g, 1 mmol) and *n*-propylamine (0.16 mL, 2 mmol). The product **4212** was a white solid (248 mg, 74%), Mp 189-190 °C. IR (KBr, ν_max_ cm^-1^) 3258, 2964, 1649, 1610; ^1^H NMR (600 MHz, DMSO-*d_6_*) δ_H_ 9.82 (s, 1H), 8.70 (d, *J* = 2.26 Hz, 1H), 8.47 (dd, *J* = 1.69, 4.71 Hz, 1H), 8.25 (t, *J* = 5.65 Hz, 1H), 7.92 (td, *J* = 1.74, 8.19 Hz, 1H), 7.60 - 7.66 (m, 2H), 7.51 (d, *J* = 15.81 Hz, 1H), 7.39 - 7.49 (m, 4H), 7.00 (s, 1H), 6.86 (d, *J* = 16.19 Hz, 1H), 3.13 (q, *J* = 6.40 Hz, 2H), 1.49 (sxt, *J* = 7.30 Hz, 2H), 0.88 (t, *J* = 7.34 Hz, 3H); ^13^C NMR (214 MHz, DMSO-*d_6_*) δ_C_ 164.7, 164.5, 150.1, 148.9, 140.3, 135.9, 134.7, 132.2, 130.5, 129.9, 129.1, 127.8, 123.7, 123.2, 121.4, 41.0, 39.8, 39.7, 39.6, 39.5, 39.4, 39.3, 39.2, 22.4, 11.5; LC/MS (ESI), RT = 2.7 min, m/z 336 (M+1).

#### 4.1.48. (Z)-2-[(E)-3-(4-chlorophenyl)acrylamido]-N-propyl-3-(pyridin-3-yl)acrylamide (4312)

This compound was prepared according to the procedure described for the synthesis of **1502** starting from (Z)-2-[(E)-4-chlorostyryl]-4-(pyridin-3-ylmethylene)oxazol-5(4H)-one (0.310 g, 1 mmol) and *n*-propylamine (0.16 mL, 2 mmol). The product **4312** was a white solid (288 mg, 78%), Mp 191-192 °C. IR (KBr, ν_max_ cm^-1^) 3261, 2964, 1649, 1609; ^1^H NMR (600 MHz, DMSO-*d_6_*) δ_H_ 9.83 (br. s., 1H), 8.70 (d, *J* = 2.26 Hz, 1H), 8.47 (dd, *J* = 1.69, 4.71 Hz, 1H), 8.25 (t, *J* = 5.46 Hz, 1H), 7.92 (d, *J* = 7.91 Hz, 1H), 7.63 (d, *J* = 7.15 Hz, 2H), 7.51 (d, *J* = 15.81 Hz, 1H), 7.44 - 7.48 (m, 2H), 7.40 - 7.44 (m, 2H), 7.00 (s, 1H), 6.86 (d, *J* = 15.81 Hz, 1H), 3.13 (q, *J* = 6.53 Hz, 2H), 1.49 (sxt, *J* = 7.30 Hz, 2H), 0.88 (t, *J* = 7.34 Hz, 3H).

#### 4.1.49. (Z)-2-[(E)-3-(4-methoxyphenyl)acrylamido]-N-propyl-3-(pyridin-3- yl)acrylamide (4412)

This compound was prepared according to the procedure described for the synthesis of **1502** starting from (Z)-2-[(E)-4-methoxystyryl]-4-(pyridin-3-ylmethylene)oxazol-5(4H)-one (0.306 g, 1 mmol) and *n*-propylamine (0.16 mL, 2 mmol). The product **4412** was a white solid (300 mg, 82%), Mp 185-186 °C. IR (KBr, ν_max_ cm^-1^) 3273, 3220, 2964, 1648, 1616; ^1^H NMR (600 MHz, DMSO-*d_6_*) δ_H_ 9.72 (s, 1H), 8.69 (s, 1H), 8.47 (dd, *J* = 1.69, 4.71 Hz, 1H), 8.22 (t, *J* = 5.27 Hz, 1H), 7.91 (dd, *J* = 1.32, 8.09 Hz, 1H), 7.57 (d, *J* = 8.66 Hz, 2H), 7.46 (d, *J* = 15.43 Hz, 1H), 7.41 (dd, *J* = 4.71, 8.09 Hz, 1H), 7.02 (d, *J* = 8.66 Hz, 2H), 6.97 (s, 1H), 6.71 (d, *J* = 15.81 Hz, 1H), 3.81 (s, 3H), 3.13 (q, *J* = 6.53 Hz, 2H), 1.49 (sxt, *J* = 7.23 Hz, 2H), 0.88 (t, *J* = 7.34 Hz, 3H), ^13^C NMR (214 MHz, DMSO-*d_6_*) δ_C_ 164.8, 164.4, 160.7, 150.1, 148.9, 140.1, 135.9, 132.3, 130.6, 129.5, 127.3, 123.7, 123.0, 118.8, 114.6, 55.4, 41.0, 39.8, 39.7, 39.6, 39.5, 39.4, 39.3, 39.2, 22.4, 11.5; LC/MS (ESI), RT = 2.7 min, m/z 366 (M+1).

#### 4.1.50. (Z)-N-propyl-3-(pyridin-3-yl)-2-[(E)-3-(p-tolyl)acrylamido]acrylamide (4512)

This compound was prepared according to the procedure described for the synthesis of **1502** starting from (Z)-2-[(E)-4-methylstyryl]-4-(pyridin-3-ylmethylene)oxazol-5(4H)-one (0.290 g, 1 mmol) and *n*-propylamine (0.16 mL, 2 mmol). The product **4512** was a white solid (272 mg, 78%), Mp 190-191 °C. IR (KBr, ν_max_ cm^-1^) 3270, 2967, 1648, 1608; ^1^H NMR (600 MHz, DMSO-*d_6_*) δ_H_ 9.77 (s, 1H), 8.69 (d, *J* = 1.88 Hz, 1H), 8.47 (dd, *J* = 1.69, 4.71 Hz, 1H), 8.23 (t, *J* = 5.83 Hz, 1H), 7.92 (td, *J* = 1.60, 8.09 Hz, 1H), 7.52 (d, *J* = 7.91 Hz, 2H), 7.47 (d, *J* = 15.81 Hz, 1H), 7.41 (dd, *J* = 4.33, 8.09 Hz, 1H), 7.27 (d, *J* = 7.91 Hz, 2H), 6.98 (s, 1H), 6.80 (d, *J* = 15.81 Hz, 1H), 3.12 (q, *J* = 6.65 Hz, 2H), 2.35 (s, 3H), 1.49 (sxt, *J* = 7.30 Hz, 2H), 0.88 (t, *J* = 7.34 Hz, 3H); ^13^C NMR (214 MHz, DMSO-*d_6_*) δ_C_ 164.7, 164.6, 150.1, 148.9, 140.3, 139.8, 135.9, 132.3, 132.0, 130.6, 129.7, 127.8, 123.7, 123.1, 120.4, 41.1, 39.8, 39.7, 39.6, 39.5, 39.4, 39.3, 39.2, 22.4, 21.1, 11.5; LC/MS (ESI), RT = 3.1 min, m/z 350 (M+1).

#### 4.1.51. (Z)-N-propyl-3-(pyridin-3-yl)-2-[(E)-3-(thien-2-yl)acrylamido]acrylamide (4612)

This compound was prepared according to the procedure described for the synthesis of **1502** starting from (Z)-4-(pyridin-3-ylmethylene)-2-[(E)-2-(thien-3-yl)vinyl]oxazol-5(4H)-one (0.282 g, 1 mmol) and *n*-propylamine (0.16 mL, 2 mmol). The product **4612** was an off-white solid (300 mg, 88%), Mp 182-183 °C. IR (KBr, ν_max_ cm^-1^) 3270, 2967, 1647, 1605; ^1^H NMR (600 MHz, DMSO-*d_6_*) δ_H_ 9.78 (br. s., 1H), 8.68 (d, *J* = 1.88 Hz, 1H), 8.47 (dd, *J* = 1.51, 4.89 Hz, 1H), 8.23 (t, *J* = 5.84 Hz, 1H), 7.91 (td, *J* = 1.79, 8.09 Hz, 1H), 7.62 - 7.70 (m, 2H), 7.39 - 7.47 (m, 2H), 7.15 (dd, *J* = 3.58, 5.08 Hz, 1H), 6.99 (s, 1H), 6.60 (d, *J* = 15.81 Hz, 1H), 3.12 (q, *J* = 6.65 Hz, 2H), 1.49 (sxt, *J* = 7.23 Hz, 2H), 0.88 (t, *J* = 7.53 Hz, 3H); ^13^C NMR (214 MHz, DMSO-*d_6_*) δ_C_ 164.6, 164.2, 150.1, 148.9, 139.7, 135.9, 133.3, 132.1, 131.5, 131.5, 130.5, 128.6, 123.7, 123.2, 120.1, 41.0, 39.8, 39.7, 39.6, 39.5, 39.4, 39.3, 39.2, 22.3, 11.5; LC/MS (ESI), RT = 2.8 min, m/z 342 (M+1).

#### 4.1.52. (Z)-2-cinnamamido-3-(4-nitrophenyl)-N-propylacrylamide (4712)

This compound was prepared according to the procedure described for the synthesis of **1502** starting from 4-[(Z)-4-nitrobenzylidene]-2-[(E)-styryl]oxazol-5(4H)-one (0.320 g, 1 mmol) and *n*-propylamine (0.16 mL, 2 mmol). The product **4712** was a white solid (341 mg, 90%), Mp 211-212 °C. IR (KBr, ν_max_ cm^-1^) 3294, 2958, 1647, 1619; ^1^H NMR (600 MHz, DMSO-*d_6_*) δ_H_ 9.94 (br. s., 1H), 8.34 (t, *J* = 5.84 Hz, 1H), 8.23 (d, *J* = 8.66 Hz, 2H), 7.77 (d, *J* = 9.03 Hz, 2H), 7.62 (d, *J* = 7.15 Hz, 2H), 7.51 (d, *J* = 15.81 Hz, 1H), 7.43 - 7.48 (m, 3H), 6.96 (s, 1H), 6.86 (d, *J* = 16.19 Hz, 1H), 3.13 (q, *J* = 6.65 Hz, 2H), 1.50 (sxt, *J* = 7.30 Hz, 2H), 0.89 (t, *J* = 7.53 Hz, 3H); ^13^C NMR (214 MHz, DMSO-*d_6_*) δ_C_ 164.7, 164.3, 146.4, 141.7, 140.5, 134.7, 133.8, 130.2, 130.0, 129.1, 127.8, 123.7, 123.0, 121.3, 41.1, 39.8, 39.7, 39.6, 39.5, 39.4, 39.3, 39.2, 22.3, 11.5; LC/MS (ESI), RT = 3.7 min, m/z 380 (M+1).

#### 4.1.53. (Z)-2-[(E)-3-(4-chlorophenyl)acrylamido]-3-(4-nitrophenyl)-N-propylacrylamide (4812)

This compound was prepared according to the procedure described for the synthesis of **1502** starting from 2-[(E)-4-chlorostyryl]-4-[(Z)-4-nitrobenzylidene]oxazol-5(4H)-one (0.355 g, 1 mmol) and *n*-propylamine (0.16 mL, 2 mmol). The product **4812** was a yellow solid (363 mg, 88%), Mp 199-200 °C. IR (KBr, ν_max_ cm^-1^) 3070, 2967, 1648, 1622, 1530, 1350; ^1^H NMR (600 MHz, DMSO-*d_6_*) δ_H_ 9.95 (s, 1H), 8.34 (t, *J* = 5.46 Hz, 1H), 8.23 (d, *J* = 9.04 Hz, 2H), 7.77 (d, *J* = 9.04 Hz, 2H), 7.65 (d, *J* = 8.66 Hz, 2H), 7.50 - 7.55 (m, 3H), 6.97 (s, 1H), 6.86 (d, *J* = 15.81 Hz, 1H), 3.13 (q, *J* = 6.53 Hz, 2H), 1.50 (sxt, *J* = 7.23 Hz, 2H), 0.89 (t, *J* = 7.53 Hz, 3H); ^13^C NMR (214 MHz, DMSO-*d_6_*) δ_C_ 164.7, 164.1, 146.4, 141.7, 139.1, 134.4, 133.7, 133.7, 130.2, 129.5, 129.2, 123.7, 123.1, 122.1, 41.1, 39.8, 39.7, 39.6, 39.5, 39.4, 39.3, 39.2, 22.3, 11.5; LC/MS (ESI), RT = 4.9 min, m/z 414 (M+1), 415 (M+2).

#### 4.1.54. (Z)-2-[(E)-3-(4-methoxyphenyl)acrylamido]-3-(4-nitrophenyl)-N-propylacrylamide (4912)

This compound was prepared according to the procedure described for the synthesis of **1502** starting from 2-[(E)-4-methoxystyryl]-4-[(Z)-4-nitrobenzylidene]oxazol-5(4H)-one (0.350 g, 1 mmol) and *n*-propylamine (0.16 mL, 2 mmol). The product **4912** was a yellow solid (348 mg, 85%), Mp 235-236 °C. IR (KBr, ν_max_ cm^-1^) 3264, 2967, 1644, 1608, 1520, 1335; ^1^H NMR (600 MHz, DMSO-*d_6_*) δ_H_ 9.84 (br. s., 1H), 8.31 (t, *J* = 5.65 Hz, 1H), 8.23 (d, *J* = 9.04 Hz, 2H), 7.77 (d, *J* = 8.66 Hz, 2H), 7.57 (d, *J* = 8.66 Hz, 2H), 7.46 (d, *J* = 15.81 Hz, 1H), 7.02 (d, *J* = 8.66 Hz, 2H), 6.93 (s, 1H), 6.71 (d, *J* = 15.81 Hz, 1H), 3.81 (s, 3H), 3.13 (q, *J* = 6.40 Hz, 2H), 1.50 (sxt, *J* = 7.23 Hz, 2H), 0.89 (t, *J* = 7.53 Hz, 3H); ^13^C NMR (214 MHz, DMSO-*d_6_*) δ_C_ 164.8, 164.6, 160.7, 146.3, 141.8, 140.3, 134.0, 130.1, 129.5, 127.2, 123.7, 122.6, 118.7, 114.5, 55.4, 41.1, 39.8, 39.7, 39.6, 39.5, 39.4, 39.3, 39.2, 22.3, 11.5.

#### 4.1.55. (Z)-3-(4-nitrophenyl)-N-propyl-2-[(E)-3-(p-tolyl)acrylamido]acrylamide (5012)

This compound was prepared according to the procedure described for the synthesis of **1502** starting from 2-[(E)-4-methylstyryl]-4-[(Z)-4-nitrobenzylidene]oxazol-5(4H)-one (0.334 g, 1 mmol) and *n*-propylamine (0.16 mL, 2 mmol). The product **5012** was a yellow solid (350 mg, 89%), Mp 206-207 °C. IR (KBr, ν_max_ cm^-1^) 3257, 2964, 1647, 1621, 1515, 1340; ^1^H NMR (600 MHz, DMSO-*d_6_*) δ_H_ 9.89 (s, 1H), 8.33 (t, *J* = 5.65 Hz, 1H), 8.23 (d, *J* = 9.04 Hz, 2H), 7.77 (d, *J* = 9.04 Hz, 2H), 7.45 - 7.52 (m, 3H), 7.27 (d, *J* = 7.91 Hz, 2H), 6.95 (s, 1H), 6.80 (d, *J* = 15.81 Hz, 1H), 3.13 (q, *J* = 6.53 Hz, 2H), 2.35 (s, 3H), 1.50 (sxt, *J* = 7.30 Hz, 2H), 0.89 (t, *J* = 7.34 Hz, 3H); ^13^C NMR (214 MHz, DMSO-*d_6_*) δ_C_ 164.8, 164.5, 146.4, 141.8, 140.5, 139.9, 133.9, 131.9, 130.1, 129.7, 127.9, 123.7, 122.8, 120.3, 41.1, 39.8, 39.7, 39.6, 39.5, 39.4, 39.3, 39.2, 22.3, 21.1, 11.5; LC/MS (ESI), RT = 5.8 min, m/z 394 (M+1).

#### 4.1.56. (Z)-3-(4-nitrophenyl)-N-propyl-2-[(E)-3-(thien-2-yl)acrylamido]acrylamide (5112)

This compound was prepared according to the procedure described for the synthesis of **1502** starting from 4-[(Z)-4-nitrobenzylidene]-2-[(E)-2-(thien-3-yl)vinyl]oxazol-5(4H)-one (0.326 g, 1 mmol) and *n*-propylamine (0.16 mL, 2 mmol). The product **5112** was a yellow solid (320 mg, 83%), Mp 187-188 °C. IR (KBr, ν_max_ cm^-1^) 3272, 2964, 1650, 1609, 1510, 1335; ^1^H NMR (600 MHz, DMSO-*d_6_*) δ_H_ 9.89 (s, 1H), 8.32 (t, *J* = 5.65 Hz, 1H), 8.24 (d, *J* = 9.04 Hz, 2H), 7.76 (d, *J* = 9.03 Hz, 2H), 7.64 - 7.69 (m, 2H), 7.45 (d, *J* = 3.39 Hz, 1H), 7.15 (dd, *J* = 3.58, 5.08 Hz, 1H), 6.95 (s, 1H), 6.60 (d, *J* = 15.43 Hz, 1H), 3.12 (q, *J* = 6.40 Hz, 2H), 1.50 (sxt, *J* = 7.30 Hz, 2H), 0.89 (t, *J* = 7.53 Hz, 3H); ^13^C NMR (214 MHz, DMSO-*d_6_*) δ_C_ 164.7, 164.1, 146.4, 141.7, 139.7, 133.7, 133.6, 131.6, 130.1, 128.7, 128.6, 123.7, 123.0, 120.0, 41.1, 39.8, 39.7, 39.6, 39.5, 39.4, 39.3, 39.2, 22.3, 11.5; LC/MS (ESI), RT = 4.9 min, m/z 386 (M+1).

### 4.2. Biological screening

#### 4.2.1. Materials for Cell culture

Different cell lines were originally purchased from American type culture collection (ATCC, Wesel, Germany) and grown in the tissue culture lab of the Egyptian company for production of vaccines, sera and drugs (Vacsera, Giza, Egypt). The cells were transferred to our laboratory and maintained in the appropriate media as following. HCT-116, CACO-2, and HT-29 (human colon cancer cell lines) were maintained in Roswell Park Memorial Institute medium (RPMI1640) (Invitrogen, Carlsbad, CA). The mouse skin fibroblasts (C-166) and Baby Hamster Kidney fibroblasts (BHK) were grown in Dulbecco Modified Eagle’s medium (DMEM). Both media were supplemented with 1% of 100 mg/ mL of streptomycin, 100 units/ mL of penicillin and 10% of heat-inactivated fetal bovine serum (Invitrogen, Carlsbad, CA) in a humidified, 5% (v/v) CO2 atmosphere at 37 °C.

#### 4.2.2. Cytotoxicity assay

The sulforhodamine B (SRB) assays were performed according to Skehan et al.^19^ Briefly, exponentially growing cells were trypsinized, counted and seeded at the appropriate densities (5000 cells/100 µL/ well) into 96-well microtiter plates. Cells were incubated in a humidified atmosphere at 37°C for 24 h. Then, the cells were exposed to different compounds at the desired concentrations, (0.01, 0.1, 1, 10, and 100 µM) or to 1% dimethyl sulfoxide (DMSO) for 72 h. At the end of the treatment period, the media were removed, and the cells were fixed with 10% trichloroacetic acid at 4°C for 1 hr. Following, the cells were washed with tap water four times and incubated with SRB 0.4% for 30 min. Excess dye was removed by washing repeatedly with 1% (vol/vol) acetic acid. The protein-bound dye was dissolved in 10 mM Tris base solution for (optical density) OD determination at 510 nm using a SpectraMax plus Microplate Reader (Molecular Devices, CA). Cell viability was expressed relative to the untreated control cells.

#### 4.2.3. Nuclear fragmentation by DAPI staining

Cells were cultured on sterile 22 mm^2^ cover slips (Harvard Apparatus, MA, USA) in sterile six well plates at a density of 2×10^5^ cells/well. 24 h after seeding, cells were exposed to IC_50_ of the tested compound in fresh medium for 24 h. At the end of the exposure, cells attached to cover slips were washed with PBS and fixed with 3.7% paraformaldhyde for 10 min, permeabilized with 0.25% Triton X-100 in TBST containing 0.01% Tween 20 for 10 min, and blocked for 1 hr with 5% goat serum in TBST. The fixed and permeabilized cells nuclei were denatured with 2 N HCl (300 μl) for 10 min, washed three times more, and treated with 0.1 μg/ml 4’,6’-Diamidino-2-Phenylindole, dihydrochloride (DAPI) (Sigma– Aldrich, St. Louis, MO, USA) (1:1000) in PBST for 1 hr. After staining, the cells were washed twice with PBS. The cover slips were then mounted on a glass slide with anti-fade mounting medium and viewed with an epifluorescence microscope, Leica, DM 5500 B (Leica, Buffalo Grove, IL, USA) at a magnification of 60×, and data were captured digitally and quantified using the microscope provided software.

#### 4.2.4. Cell morphology

Cells were cultured on sterile 22 mm^2^ cover slips (Harvard Apparatus, MA, USA) in sterile six well plates at a density of 2×10^5^ cells/well. 24 h after seeding, cells were exposed to IC_50_ of the tested compounds in fresh medium for 24 h. At the end of the exposure, cells attached to cover slips were washed with PBS and visulized under leica light microscope (Leica, Buffalo Grove, IL, USA).

#### 4.2.5. Cell Cycle Analysis

To analyze the DNA content by flow cytometry, HCT-116 cells were seeded at a density of 3×10^6^ cell/ T 75 flask for 24 h and then exposed to different compounds at their IC_50_ values for 24 h. The cells were collected by trypsinization, washed with phosphate buffered saline (PBS) and fixed in ice-cold absolute alcohol. Thereafter, cells were stained using Cycletest^TM^ Plus DNA Reagent Kit (BD Biosciences, San Jose, CA) according to the manufacturer’s instructions. Cell cycle distribution was determined using a FACS Calibur flow cytometer (BD Biosciences, San Jose, CA).

#### 4.2.6. Primary (1°) Colonosphere Formation Assay

For primary sphere formation, cells were plated in nontreated, low adhesion, 96 wells plate at the concentration of 100 cells/100 μL/well in stem cell media (SCM) that consisted of DMEM:F12:AA (Gibco), supplemented with 1× B27 (Gibco), 20 ng/mL epidermal growth factor, and 10 ng/mL fibroblast growth factor (Sigma). After 4 h of incubation, vehicle (control) or **3712** and **4112** at the desired concentrations were added to each well (at least in triplicates for each sample). On day five, numbers of spheres ranging from 50 to 150 mm in diameter were counted using phase contrast microscope and percent inhibition was calculated compared to control.

#### 4.2.7. Determination of ROS accumulation

To determine the effect of the newly synthesized compounds on the cellular redox status, two different free radical sensitive props, dihydroethidium (DHE) and dichlorofluorescin diacetate (DCFDA) were used. Moreover, the activity of two intracellular antioxidant enzymes, superoxide dismutase (SOD) and catalase (CAT), reduced glutathione (GSH) level and a lipid peroxidation product, Malone di aldehyde (MDA) were assessed. For DHE and DCFDA, cells were cultured on sterile 22 mm2 cover slips (Harvard Apparatus, MA, USA) in sterile six well plates at a density of 2×105 cells/well. 24 h after seeding, cells were exposed to IC_50_ of the tested compounds in fresh medium for 24 h. At the end of the exposure, cells attached to cover slips were washed thrice with PBS and incubated with DHE 10µM or DCFDA 10 µM for 30 min at 37 °C in the dark. Thereafter, cells were washed thrice with PBS and the cover slips were then mounted on a glass slide with anti-fade mounting medium containing 4’,6’-Diamidino-2-Phenylindole, dihydrochloride (DAPI) (Sigma– Aldrich, St. Louis, MO, USA), which was used as counter stain and viewed with an epifluorescence microscope, Leica, DM 5500 B (Leica, Buffalo Grove, IL, USA) at a magnification of 60×. Data were captured digitally and quantified using the microscope provided software.

For assessment of SOD and CAT activities as well as SOD and MDA levels, 4 x10^6^ cell/ T 75 flask were exposed to the IC_50_ of tested compound for 24 h. The cells were collected by trypsinization and washed twice with PBS. Cells were directly homogenized in PBS on ice with a Dounce homogenizer 3 times (each 25 strokes) at 10-min intervals, then centrifuged at 15000 rpm for 15 min at 4 °C. An aliquot was kept to determine the protein concentration using a BioRad protein assay DC kit (Bio Rad Laboratories, CA). Different parameters were then assessed using equal protein amounts in all samples and employing the specified kit according to the manufacturer’s instructions. (3, 4, 5 and 6).

#### 4.2.8. Accumulation and translocation of cyclin B1and cyclin D1

Cells were cultured on sterile 22 mm^2^ cover slips (Harvard Apparatus, MA, USA) in sterile six well plates at a density of 2×10^5^ cells/well. 24 h after seeding, cells were exposed to IC_50_ of the tested compound in fresh medium for 24 h. At the end of the exposure, cells attached to cover slips were washed with PBS and fixed with 3.7% paraformaldhyde for 10 min, permeabilized with 0.25% Triton X-100 in TBST containing 0.01% Tween 20 for 10 min, and blocked for 1 hr with 5% goat serum in TBST. The fixed and permeabilized cells were incubated with Cyclin B1 rabbit mAb and Cyclin D1 rabbit mAb (Cell signaling technology, MA, USA) at a dilution of 1:500 in blocking solution overnight at 4°C, followed by secondary anti-mouse Alexa fluor-488-(Invitrogen, Carlsbad, CA) and Cy3-goat anti-Rabbit antibody (Jackson Immuno Research, West Grove, PA, USA) in 1:1000 dilution in the blocking solution for 1 hr at room temperature in the dark. 4’,6’-Diamidino-2-Phenylindole, dihydrochloride (DAPI) (Sigma– Aldrich, St. Louis, MO, USA) was used as counter stain to stain the DNA. The cover slips were then mounted on a glass slide with anti-fade mounting medium and viewed with an epifluorescence microscope, Leica, DM 5500 B (Leica, Buffalo Grove, IL, USA) at a magnification of 60×, and data were captured digitally and quantified using the microscope provided software.

#### 4.2.9. Phosphohistone H3 and Caspase-3 activity assay

Cells were cultured on sterile 22 mm^2^ cover slips (Harvard Apparatus, MA, USA) in sterile six well plates at a density of 2×10^5^ cells/well. 24 h after seeding, cells were exposed to IC_50_ of the tested compound in fresh medium for 24 h. At the end of the exposure, cells attached to cover slips were washed with PBS and fixed with 3.7% paraformaldhyde for 10 min, permeabilized with 0.25% Triton X-100 in TBST containing 0.01% Tween 20 for 10 min, and blocked for 1 hr with 5% goat serum in TBST. The fixed and permeabilized cells were incubated with Phosphohistone-H3 mouse mAb, and cleaved caspase-3 rabbit mAb (Cell signaling technology, MA, USA) at a dilution of 1:500 in blocking solution overnight at 4°C, followed by secondary anti-mouse Alexa fluor-488-(Invitrogen, Carlsbad, CA) and Cy3-goat anti-Rabbit antibody (Jackson Immuno Research, West Grove, PA, USA) in 1:1000 dilution in the blocking solution for 1 hr at room temperature in the dark. 4’,6’-Diamidino-2-Phenylindole, dihydrochloride (DAPI) (Sigma– Aldrich, St. Louis, MO, USA) was used as counter stain to stain the DNA. The cover slips were then mounted on a glass slide with anti-fade mounting medium and viewed with an epifluorescence microscope, Leica, DM 5500 B (Leica, Buffalo Grove, IL, USA) at a magnification of 60×, and data were captured digitally and quantified using the microscope provided software.

#### 4.2.10. Apoptotic cell determination

Apoptosis was determined by staining cells with Annexin V–fluorescein isothiocyanate (FITC) and counterstaining with propidium iodide (PI) using the Annexin V–FITC/PI apoptosis detection kit (BD Biosciences, San Diego, CA, USA) according to the manufacturer’s instructions. Briefly, 4 x10^6^ cell/ T 75 flask were exposed to the IC_50_ of tested compound for 24 and 48 h. The cells were collected by trypsinization and 0.5 × 10^6^ cells were washed twice with phosphate-buffered saline (PBS) and stained with 5 μl Annexin V–FITC and 5 μl PI in 1× binding buffer (BD Biosciences, San Jose, CA, USA) for 15 min at room temperature in the dark. Analyses were performed using FACS Calibur flow cytometer (BD Biosciences, San Jose, CA, USA).

#### 4.2.11. Determination of acute oral toxicity (LD_50_)[40]

##### A. Animals and Compounds

The experiment was conducted on 12 healthy Swiss albino mice (males and females) weighing 22-27 g and aged 8 to 10 weeks obtained from the Animal Station, Pharmacology Dept, Faculty of Pharmacy, King Abdulaziz University, Jeddah. All animals were kept at the regulated temperature (average 23 °C), air quality (Central air conditioning) and light (12-h light/dark cycles). Animals were provided free access to food pellets *ad libitum* and water. The experimental procedure was approved by the Research Ethics Committee, Faculty of Pharmacy, King Abdulaziz University prior to starting the laboratory work. Animals were humanely treated according to international and scientific principles. Changes other than vitality of the animals such as behavioral and food consumption habits were not observed in this study.

The compounds **1512**, **3712** and **4112** of purity 95% or more (LC/MS) were prepared for this study as 10% suspension in water containing 0.5% tween 80.

##### B. Acute oral toxicity test

The animals were distributed randomly into four groups (3 mice at each group). One group did not receive any drug (control group). The second group received an oral dose of 2000 mg/Kg of the drug and observed after 24 h to count the deceased animals. According to the Guideline 423, when all animals were found alive, the same test was repeated (dose 2000 mg/Kg) were give to the third group (3 animals) and observed for 24 h. All animals survived and therefore, a dose of 5000 mg/Kg were administered to the fourth and final group.

## Supporting information

Spectral data of compounds and Details of LD50 protocol

## 5. ACKNOWLEDGEMENT

This project was funded by the National Plan for Science, Technology and Innovation (MAARIFAH), King Abdulaziz City for Science and Technology, the Kingdom of Saudi Arabia; Award number 12-BIO3193-03. The authors also, acknowledge with thanks Science and Technology Unit, King Abdulaziz University for technical support.

Authors also would like to thank Professor Alaa Khedr (KAU) and Dr. Mohini Ghatge (VCU) for their valuable help in LC/MS analyses. We also express gratitude to Professor Ashraf Abdel-Naim, Pharmacology and Toxicology Dept, KAU for his guidance on the acute toxicity studies.

**Note:** Virginia Commonwealth University and King Abdul Aziz University have filed provisional patents related to the 2-cinnamamido-N-substituted-cinnamamide (bis-cinnamamide) compounds.

---

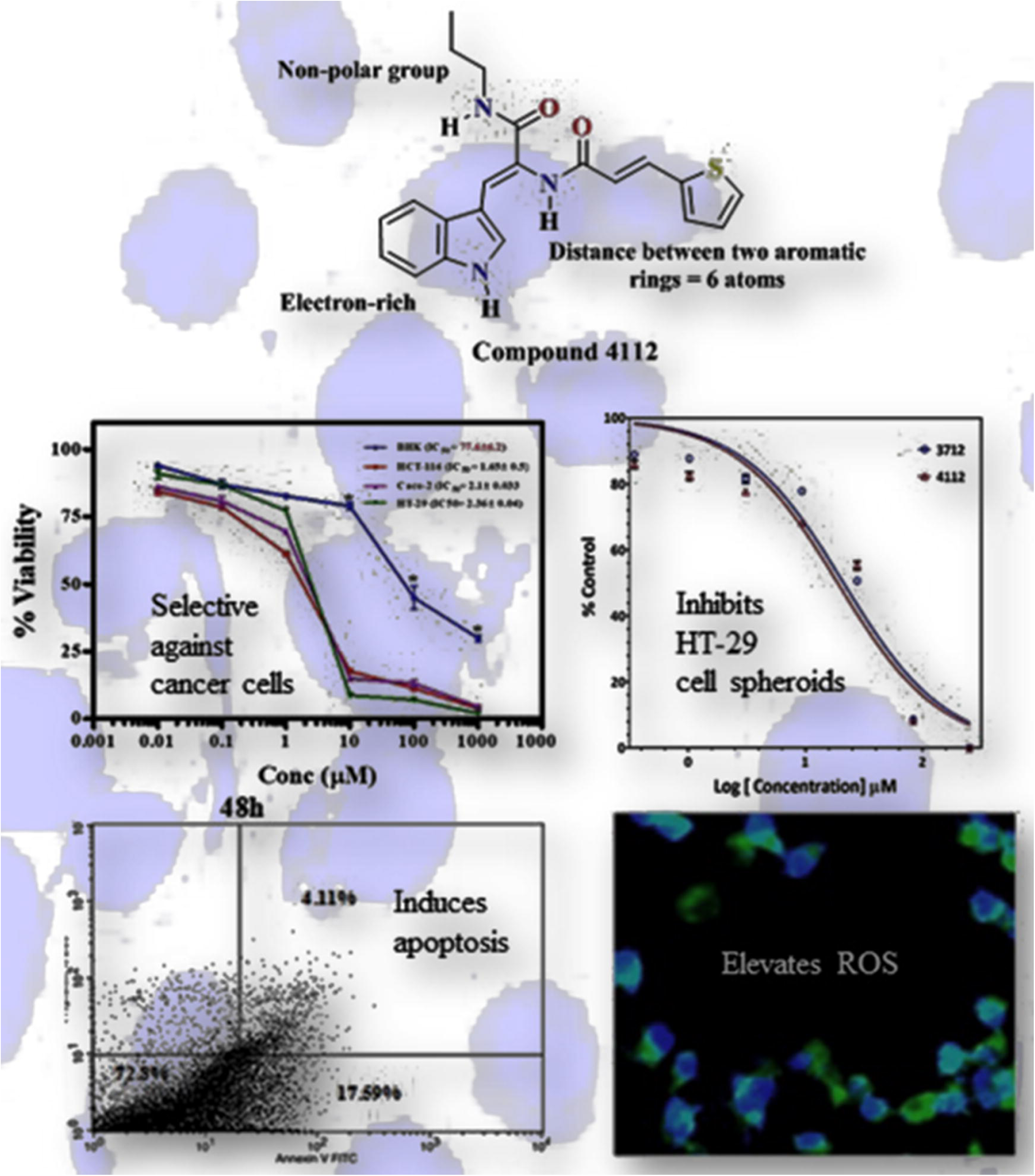

