## Supplementary material for "Discovery, optimization, and cellular activities of 2-(aroylamino)cinnamamide derivatives against colon cancer cells": Spectral data of compounds and Details of LD50 protocol

### I. Spectra of Compounds

#### Spectra of (1501)[1-2]

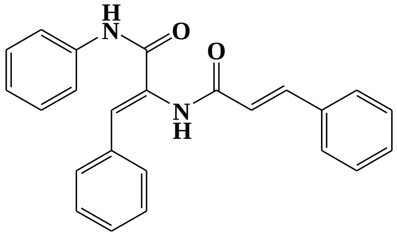


##### $^1$H NMR

This report was created by ACD/NMR Processor Academic Edition. For more information go to www.acdlabs.com/nmrproc/

**1501A**

4/23/2017 10:06:13 PM

| | | | | | | | |
|---|---|---|---|---|---|---|---|
| ***Acquisition Time (sec)*** | 2.6564 | ***Comment*** | Dr.A.Omar DMSO Cin 1501A | | | ***Date*** | 30 Mar 2013 11:51:12 |
| ***Date Stamp*** | 30 Mar 2013 11:51:12 | | | | | | |
| ***File Name*** | E:\Google Drive\AAA AAA Research\AA AA Projects\CIN\Safo-Cinamamide\SupportingInformation\Spectra\Cin 1501A dmso\1\fid | | | | | | |
| ***Frequency (MHz)*** | 600.13 | ***Nucleus*** | 1H | ***Number of Transients*** | 16 | ***Origin*** | spect |
| ***Original Points Count*** | 32768 | ***Owner*** | nmr | ***Points Count*** | 32768 | ***Pulse Sequence*** | zg30 |
| ***Receiver Gain*** | 83.44 | ***SW(cyclical) (Hz)*** | 12335.53 | ***Solvent*** | DMSO-d6 | ***Spectrum Offset (Hz)*** | 3706.0493 |
| ***Spectrum Type*** | STANDARD | ***Sweep Width (Hz)*** | 12335.15 | ***Temperature (degree C)*** | 25.014 | | |

$^1$H NMR (DMSO-$d_6$) δ ppm 9.95 (br. s., 1 H), 7.61 - 7.68 (m, 5 H), 7.55 (d, $J$=15.81 Hz, 1 H), 7.41 - 7.49 (m, 7 H), 7.36 - 7.41 (m, 1 H), 7.23 (s, 1 H), 6.89 (d, $J$=15.81 Hz, 1 H)

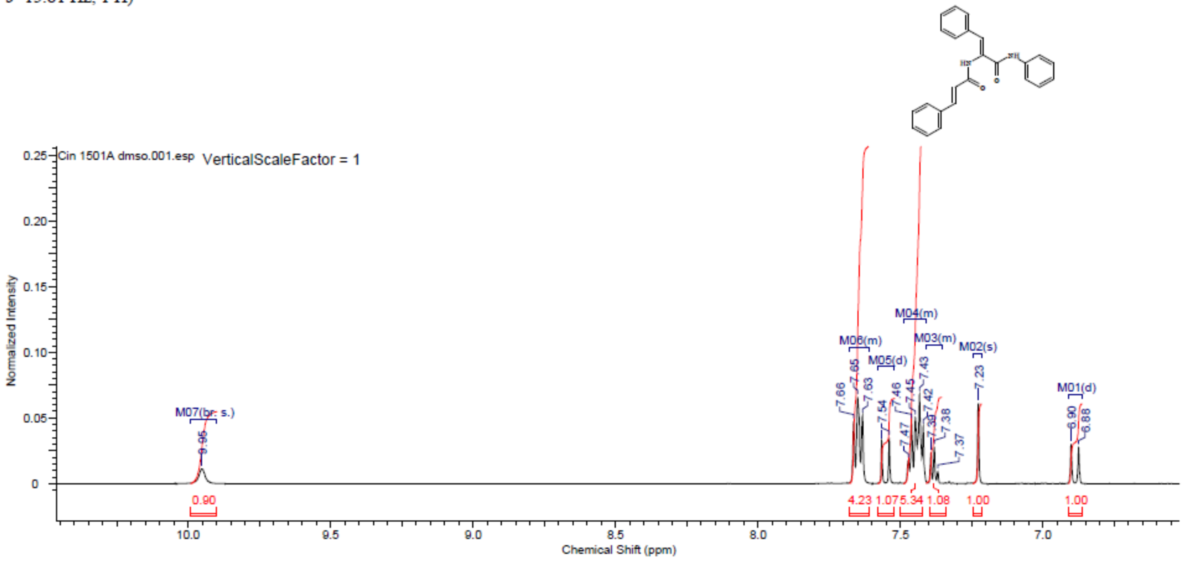


#### Spectra of (1502)[1, 3]

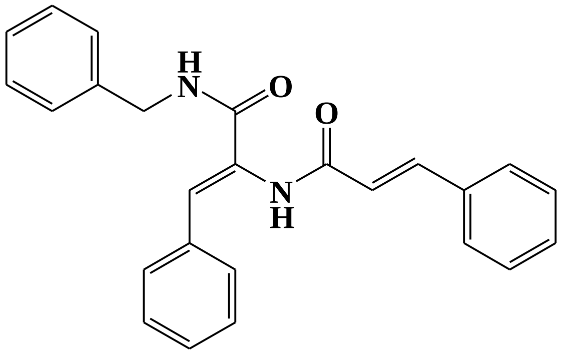
H
N
O
O
N
H

**LC/MS**

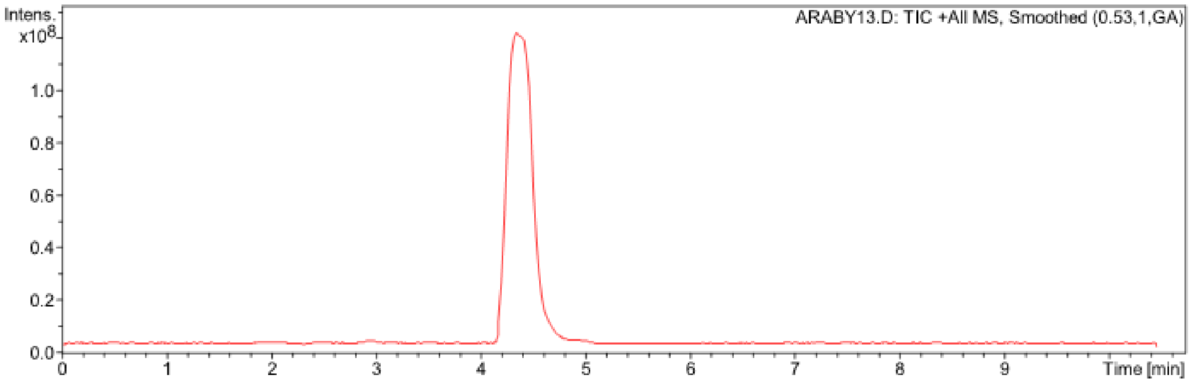
Intens.
x10^8
1.0
0.8
0.6
0.4
0.2
0.0
ARABY13.D: TIC +All MS, Smoothed (0.53,1,GA)
0
1
2
3
4
5
6
7
8
9
Time [min]

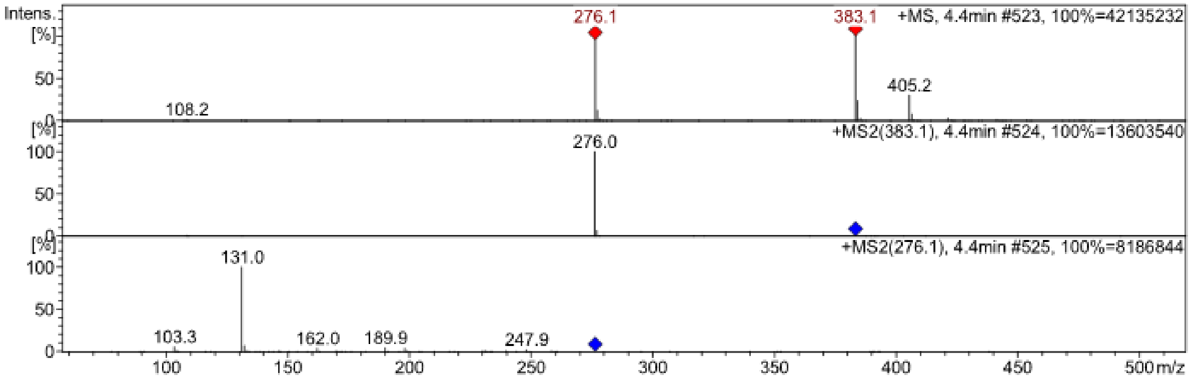
Intens.
[%]
50
0
276.1
383.1
+MS, 4.4min #523, 100%=42135232
405.2
108.2
[%]
100
50
0
276.0
+MS2(383.1), 4.4min #524, 100%=13603540
[%]
100
50
0
131.0
+MS2(276.1), 4.4min #525, 100%=8186844
103.3
162.0
189.9
247.9
100
150
200
250
300
350
400
450
500 m/z

**FT-IR**

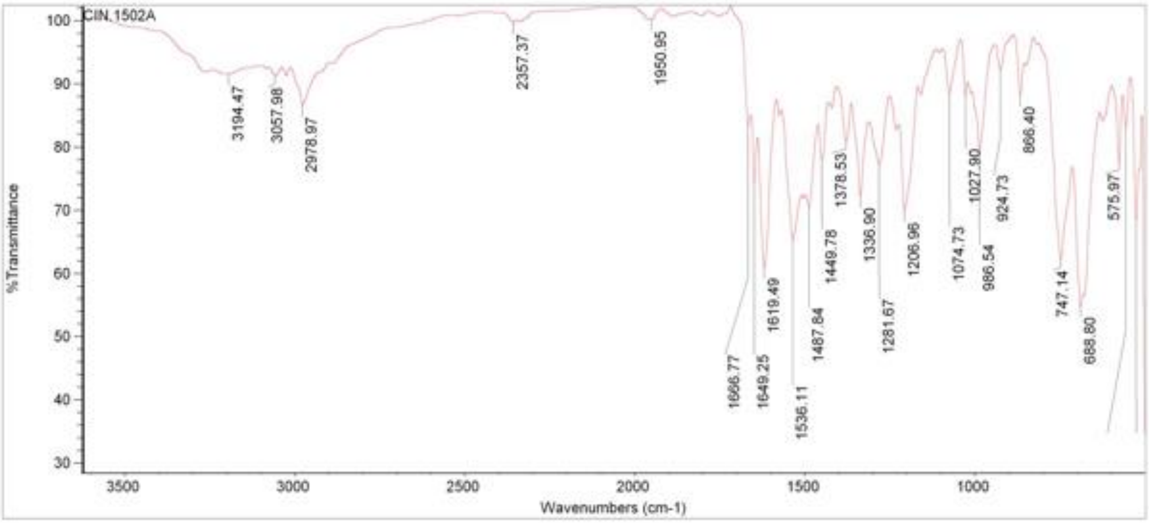


#### $^1$H NMR

This report was created by ACD/NMR Processor Academic Edition. For more information go to www.acdlabs.com/nmrproc/

8/25/2013 6:50:15 PM

| | | | | | | | |
|---|---|---|---|---|---|---|---|
| ***Acquisition Time (sec)*** | 2.6564 | ***Comment*** | Dr A.Mansour Acetone Cin-1502A 27-3-2013 | | | ***Date*** | 27 Mar 2013 14:01:20 |
| ***Date Stamp*** | 27 Mar 2013 14:01:20 | | | | | | |
| ***File Name*** | D:\AAA Research\AAA Ongoing\CINN\Spectra\NMR\CIN15Series NMR 20130707\CIN15series 1501to12 20130707\Cin-1502A\30\fid | | | | | | |
| ***Frequency (MHz)*** | 600.13 | ***Nucleus*** | 1H | ***Number of Transients*** | 16 | ***Origin*** | spect |
| ***Original Points Count*** | 32768 | ***Owner*** | nmr | ***Points Count*** | 32768 | ***Pulse Sequence*** | zg30 |
| ***Receiver Gain*** | 173.48 | ***SW(cyclical) (Hz)*** | 12335.53 | ***Solvent*** | Acetone | ***Spectrum Offset (Hz)*** | 3706.0493 |
| ***Spectrum Type*** | STANDARD | ***Sweep Width (Hz)*** | 12335.15 | ***Temperature (degree C)*** | 25.015 | | |

$^1$H NMR (600 MHz, Acetone) δ ppm 4.52 - 4.58 (m, 2 H) 6.96 (d, $J$=15.43 Hz, 1 H) 7.24 (t, $J$=7.34 Hz, 1 H) 7.29 - 7.35 (m, 4 H) 7.36 - 7.48 (m, 8 H) 7.57 - 7.68 (m, 5 H) 8.08 (br. s., 1 H)

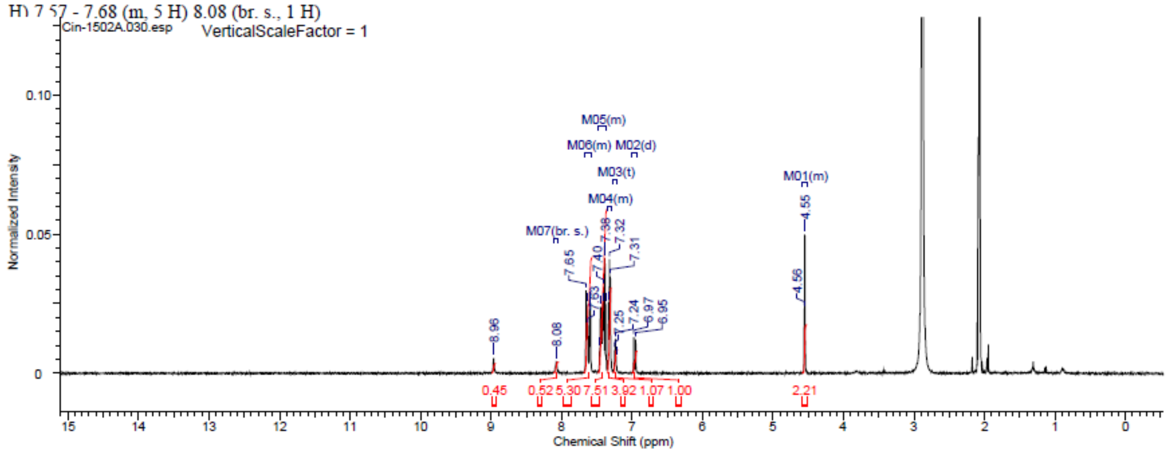

### Spectra of (1503)

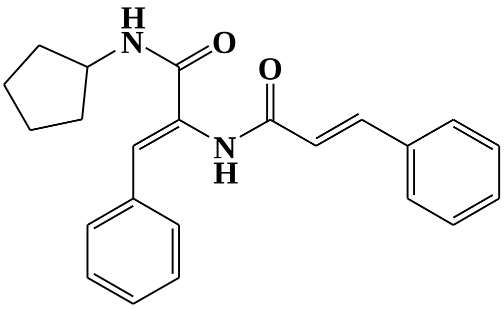

#### [1]H NMR

This report was created by ACD/NMR Processor Academic Edition. For more information go to www.acdlabs.com/nmrproc/

**1503**

2018-10-01 7:54:52 AM

| Formula $C_{23}H_{24}N_2O_2$ | FW 360.4489 |
|---|---|

| | | | | | | | |
|---|---|---|---|---|---|---|---|
| Acquisition Time (sec) | 2.6564 | Comment | Dr A.Mansour Acetone Cln-1503A 27-3-2013 | | | Date | 27 Mar 2013 14:07:44 |
| Date Stamp | 27 Mar 2013 14:07:44 | | | | | | |
| File Name | D:\AAA_Research\AAA_Ongoing\CINN\Spectra\NMR\CIN15Series_NMR_20130707\CIN15series_1501to12_20130707\Cln-1503A\50\fid | | | | | | |
| Frequency (MHz) | 600.13 | Nucleus | 1H | Number of Transients | 16 | Origin | spect |
| Original Points Count | 32768 | Owner | nmr | Points Count | 32768 | Pulse Sequence | zg30 |
| Receiver Gain | 173.48 | SW(cyclical) (Hz) | 12335.53 | Solvent | Acetone | Spectrum Offset (Hz) | 3706.0493 |
| Spectrum Type | STANDARD | Sweep Width (Hz) | 12335.15 | Temperature (degree C) | 25.012 | | |

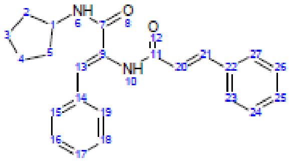

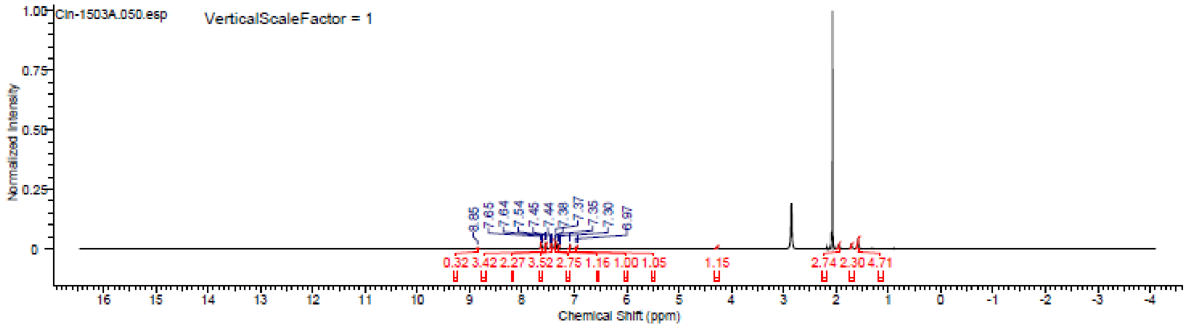

### Spectra of (1505)

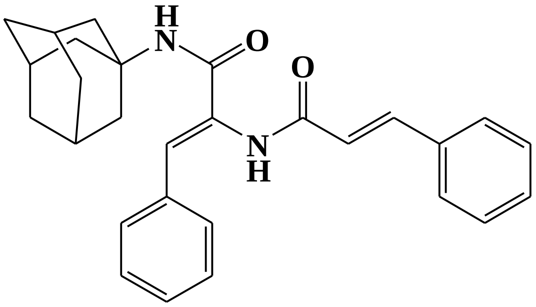

This report was created by ACD/NMR Processor Academic Edition. For more information go to www.acdlabs.com/nmrproc/

#### CIN-1505A-4m

1/6/2017 09:14:03 AM

| | | | | | | | |
|---|---|---|---|---|---|---|---|
| Acquisition Time (sec) | 2.6564 | Comment | Dr A.Mansour Acetone Cin-1505A 27-3-2013 | | | Date | 27 Mar 2013 13:54:56 |
| Date Stamp | 27 Mar 2013 13:54:56 | | | | | | |
| File Name | E:\Google Drive\Projects\CIN\Safo-Cinamamide\Spectra Bis-Cinnamamide\1H NMR\SecondBatch-Table4l-4t\ConfirmedCpdsNMR\Cin-1505A\10\fid | | | | | | |
| Frequency (MHz) | 600.13 | Nucleus | 1H | Number of Transients | 16 | Origin | spect |
| Original Points Count | 32768 | Owner | nmr | Points Count | 32768 | Pulse Sequence | zg30 |
| Receiver Gain | 140.47 | SW(cyclical) (Hz) | 12335.53 | Solvent | Acetone | Spectrum Offset (Hz) | 3706.0493 |
| Spectrum Type | STANDARD | Sweep Width (Hz) | 12335.15 | Temperature (degree C) | 25.012 | | |

$^1$H NMR (Acetone) δ ppm 7.61 - 7.68 (m, 3 H), 7.54 (d, *J*=7.53 Hz, 2 H), 7.40 - 7.48 (m, 3 H), 7.37 (t, *J*=7.53 Hz, 2 H), 7.27 - 7.32 (m, 1 H), 7.05 (br. s., 1 H), 6.98 (d, *J*=15.43 Hz, 1 H), 6.76 (br. s., 1 H), 2.10 - 2.15 (m, 6 H), 1.99 - 2.03 (m, 1 H), 1.73 (br. s., 6 H), 1.67 (d, *J*=12.05 Hz, 1 H), 1.61 (d, *J*=11.29 Hz, 1 H), 1.55 - 1.59 (m, 1 H)

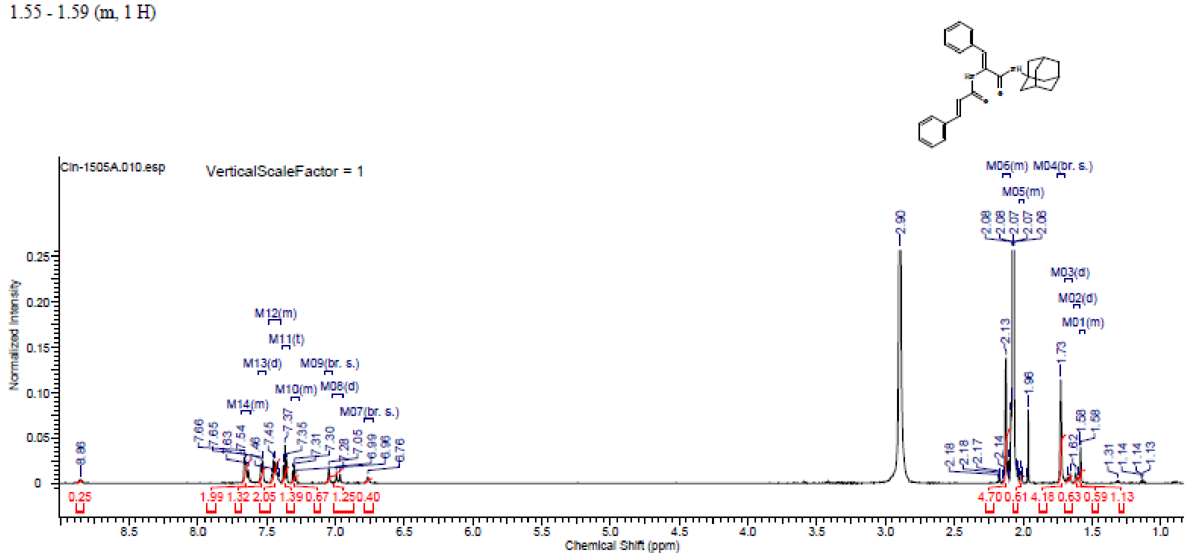

### Spectra of (1507)

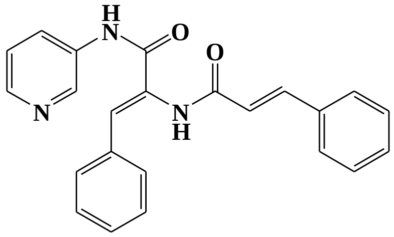


## LC/MS

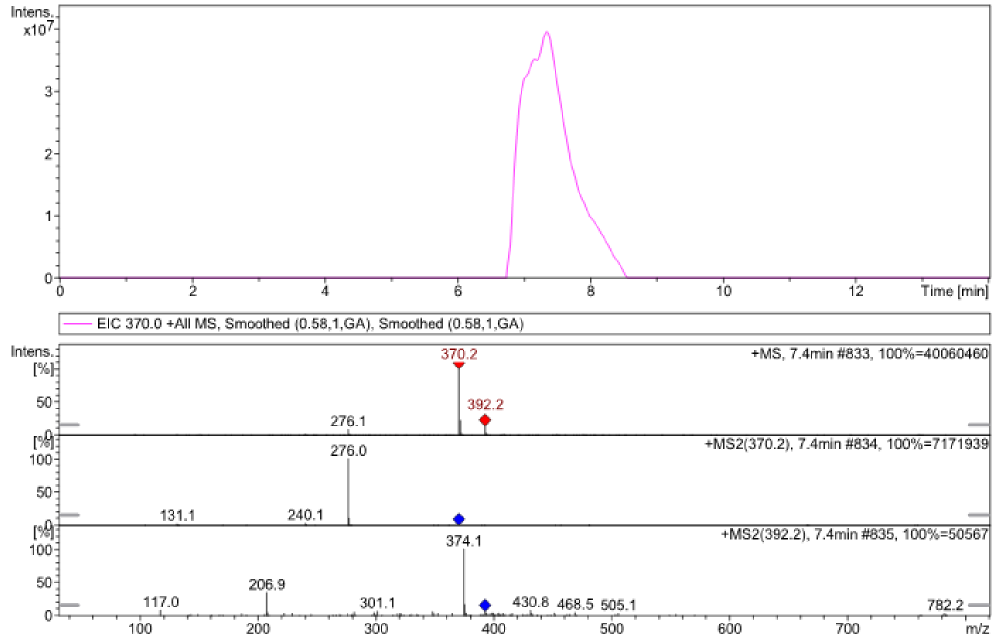


#### $^{1}$H NMR

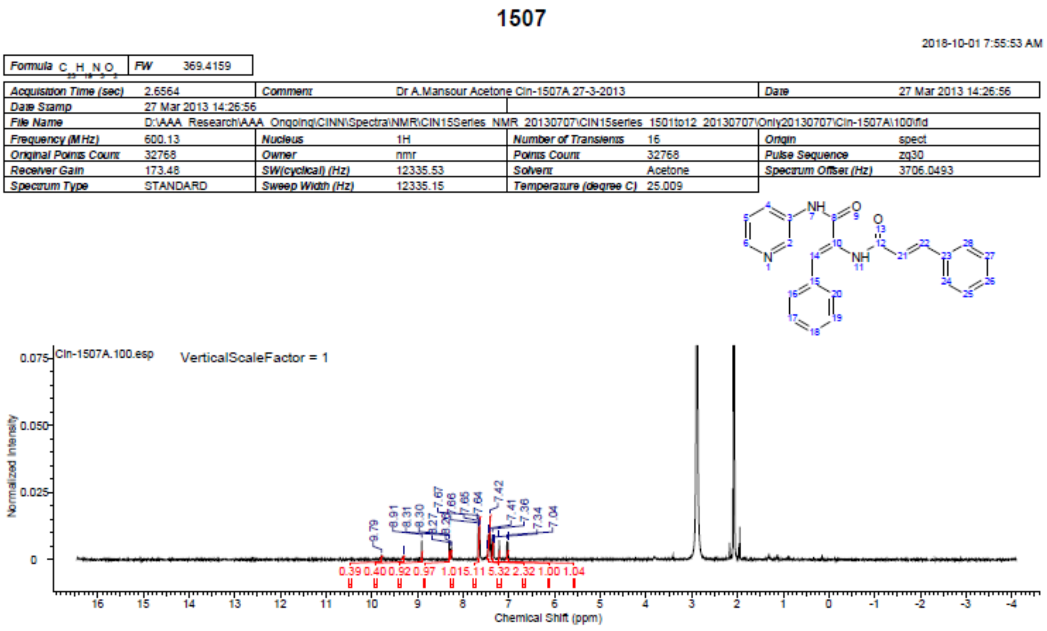

### Spectra of (1508)[3]

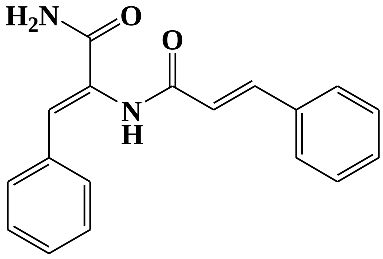


#### [1]H NMR

This report was created by ACD/NMR Processor Academic Edition. For more information go to www.acdlabs.com/nmrproc/

**1508**

10/6/2018 12:11:26 AM
Dr.Mansour DMSO CIN-1508A

| | | | | | | | |
|---|---|---|---|---|---|---|---|
| *Acquisition Time (sec)* | 2.6564 | *Comment* | Dr.Mansour DMSO CIN-1508A | | | *Date* | 07 Jul 2013 14:07:44 |
| *Date Stamp* | 07 Jul 2013 14:07:44 | | | | | | |
| *File Name* | D:\AAA_Research\AAA_Ongoing\CINN\Spectra\NMR\CIN15Series_NMR_20130707\CIN15series_1501to12_20130707\Only20130707\Cin-1508A\70\fid | | | | | | |
| *Frequency (MHz)* | 600.13 | *Nucleus* | 1H | *Number of Transients* | 16 | *Origin* | spect |
| *Original Points Count* | 32768 | *Owner* | nmr | *Points Count* | 32768 | *Pulse Sequence* | zg30 |
| *Receiver Gain* | 37.66 | *SW(cyclical) (Hz)* | 12335.53 | *Solvent* | DMSO-d6 | *Spectrum Offset (Hz)* | 3706.0493 |
| *Spectrum Type* | STANDARD | *Sweep Width (Hz)* | 12335.15 | *Temperature (degree C)* | 25.000 | | |

[1]H NMR (DMSO-$d_6$) δ ppm 10.11 (br. s., 1 H), 7.75 (br. s., 1 H), 7.61 (d, $J$=6.78 Hz, 2 H), 7.55 (d, $J$=7.53 Hz, 2 H), 7.35 - 7.53 (m, 6 H), 7.31 (d, $J$=7.15 Hz, 1 H), 7.16 (br. s., 1 H), 7.10 (s, 1 H), 6.91 (d, $J$=15.81 Hz, 1 H)

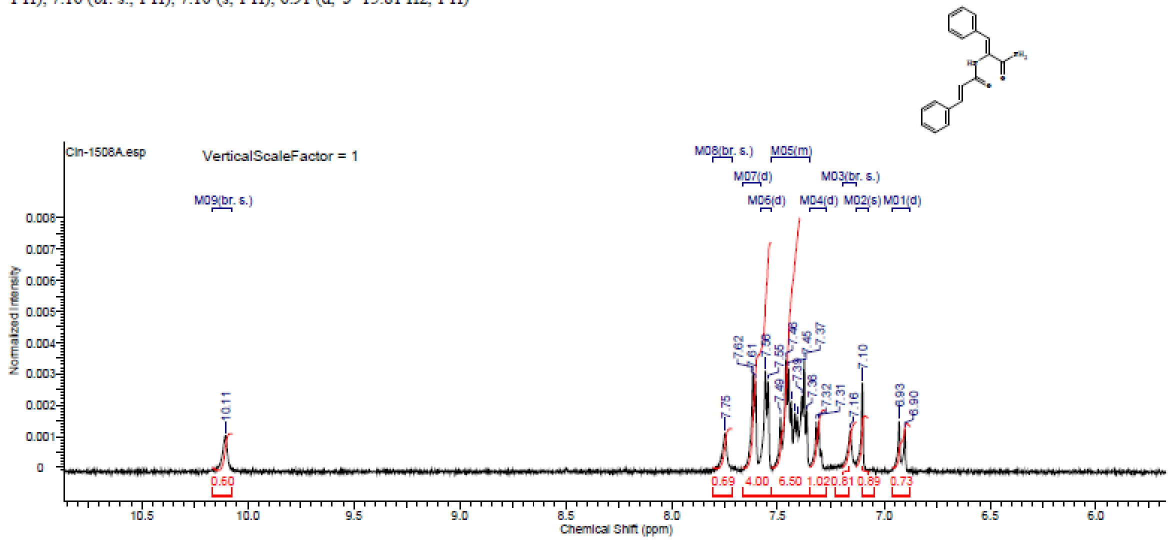

### Spectra of (1511)

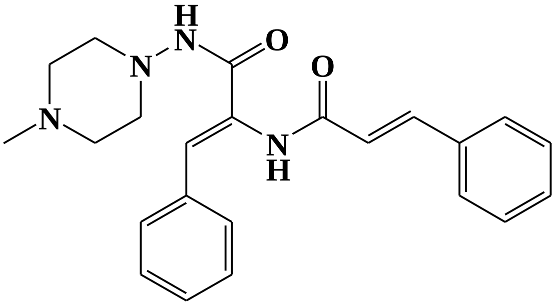

#### [1]H NMR

This report was created by ACD/NMR Processor Academic Edition. For more information go to www.acdlabs.com/nmrproc/

**Cin-1511A**

4/7/2015 04:15:42 م

| | | | | | |
|---|---|---|---|---|---|
| *Acquisition Time (sec)* | 2.6564 | *Comment* | Dr.A.Mansour DMSO RAKAN 1511A | *Date* | 25 Feb 2015 14:44:00 |
| *Date Stamp* | 25 Feb 2015 14:44:00 | *File Name* | E:\Google Drive\Projects\CIN-Extension\Spectra\Rakan\RAKAN 1511A\1\fid | | |
| *Frequency (MHz)* | 600.13 | *Nucleus* | 1H | *Number of Transients* | 16 |
| *Origin* | spect | | | | |
| *Original Points Count* | 32768 | *Owner* | nmr | *Points Count* | 32768 |
| *Pulse Sequence* | zg30 | | | | |
| *Receiver Gain* | 173.48 | *SW(cyclical) (Hz)* | 12335.53 | *Solvent* | DMSO-d6 |
| *Spectrum Offset (Hz)* | 3706.0500 | | | | |
| *Spectrum Type* | STANDARD | *Sweep Width (Hz)* | 12335.15 | *Temperature (degree C)* | 25.001 |

[1]H NMR (600 MHz, DMSO-$d_6$) δ ppm 2.77 (br. s., 2 H) 3.10 (br. s., 2 H) 3.14 - 3.22 (m, 4 H) 3.43 (br. s., 2 H) 6.75 (br. s., 1 H) 6.91 (d, J=15.81 Hz, 1 H) 7.30 - 7.35 (m, 1 H) 7.39 - 7.44 (m, 3 H) 7.44 - 7.48 (m, 2 H) 7.52 (d, J=15.81 Hz, 1 H) 7.56 (m, J=7.91 Hz, 2 H) 7.62 (d, J=7.53 Hz, 2 H) 9.58 (s, 1 H) 9.83 (s, 1 H)

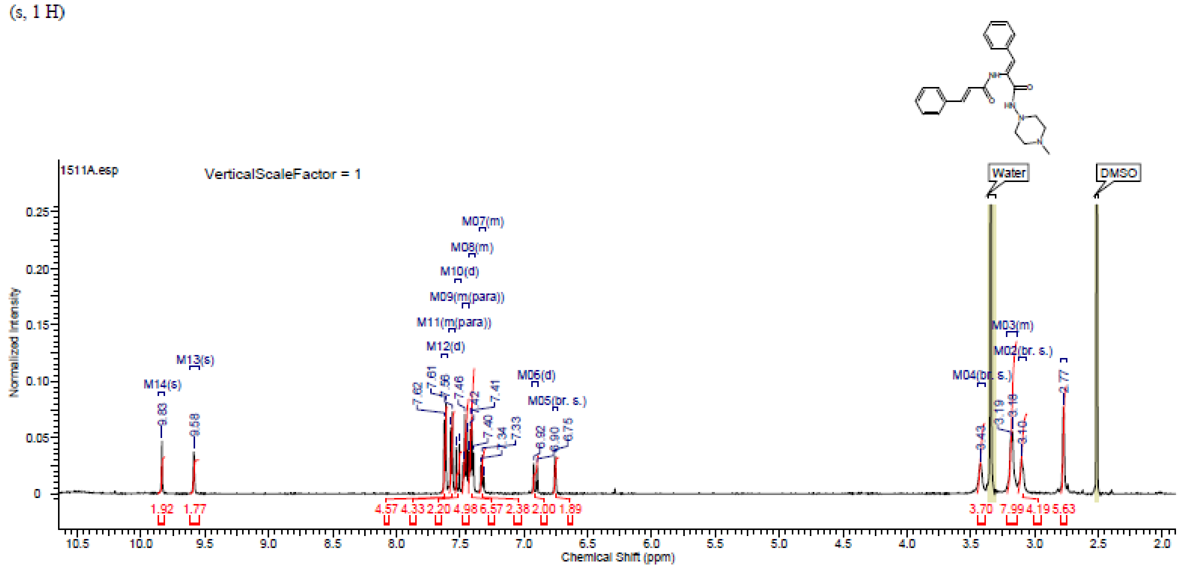

#### Spectra of (1512)

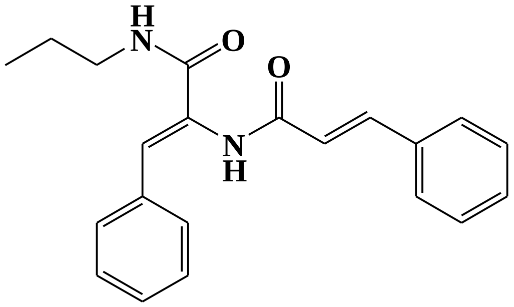


### LC/MS

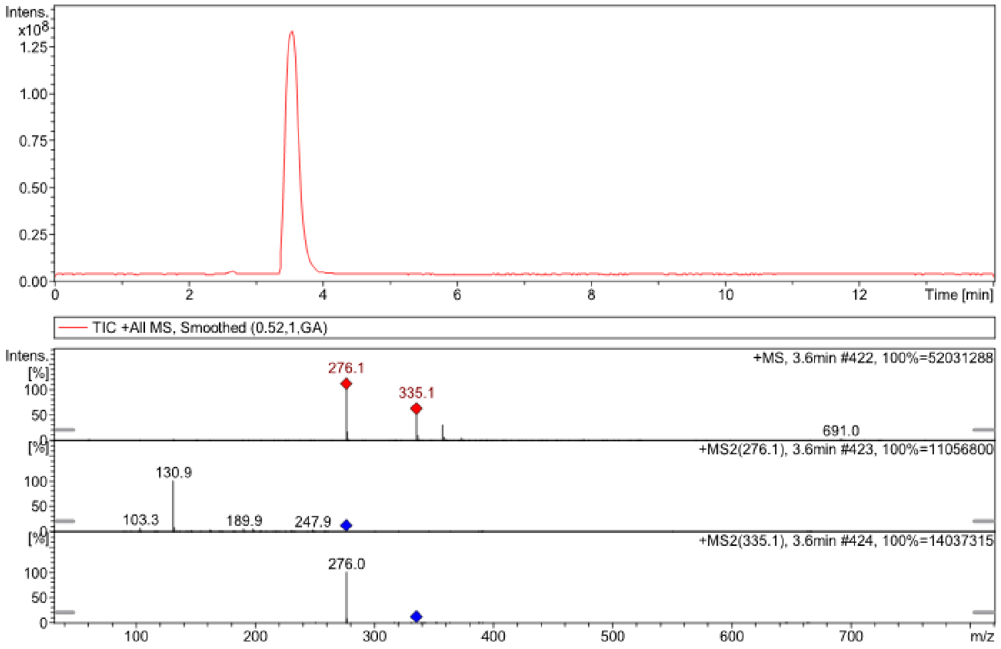


##### $^{1}$H NMR

This report was created by ACD/NMR Processor Academic Edition. For more information go to www.acdlabs.com/nmrproc/

**1512**

2018-10-01 7:58:31 AM

| Formula | $C_{21}H_{22}N_2O_2$ | FW | 334.4116 |
|---|---|---|---|

| | | | | | | | |
|---|---|---|---|---|---|---|---|
| Acquisition Time (sec) | 2.6564 | Comment | Dr.Mansour DMSO CIN-1512A | | | Date | 07 Jul 2013 14:22:40 |
| Date Stamp | 07 Jul 2013 14:22:40 | | | | | | |
| File Name | D:\AAA Research\AAA Ongoing\CINN\Spectra\NMR\CIN15Series NMR 20130707\CIN15series 1501to12 20130707\Garbage\CIN-1512A\90\fid | | | | | | |
| Frequency (MHz) | 600.13 | Nucleus | 1H | Number of Transients | 16 | Origin | spect |
| Original Points Count | 32768 | Owner | nmr | Points Count | 32768 | Pulse Sequence | zg30 |
| Receiver Gain | 34.94 | SW(cyclical) (Hz) | 12335.53 | Solvent | DMSO-d6 | Spectrum Offset (Hz) | 3706.0483 |
| Spectrum Type | STANDARD | Sweep Width (Hz) | 12335.15 | Temperature (degree C) | 25.001 | | |

$^{1}$H NMR (600 MHz, DMSO-$d_6$) δ ppm 0.84 - 0.91 (m, 3 H) 1.44 - 1.53 (m, 2 H) 3.09 - 3.16 (m, 2 H) 6.90 (dd, $J$=15.81, 2.64 Hz, 1 H) 6.99 - 7.04 (m, 1 H) 7.28 - 7.34 (m, 1 H) 7.34 - 7.52 (m, 7 H) 7.56 (d, $J$=6.78 Hz, 2 H) 7.62 (d, $J$=7.15 Hz, 2 H) 8.15 - 8.22 (m, 1 H) 9.86 (br. s., 1 H)

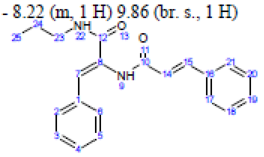

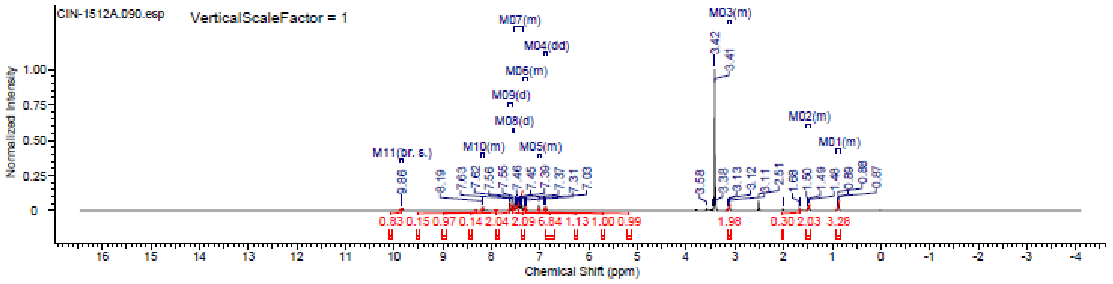

### Spectra of (1513)

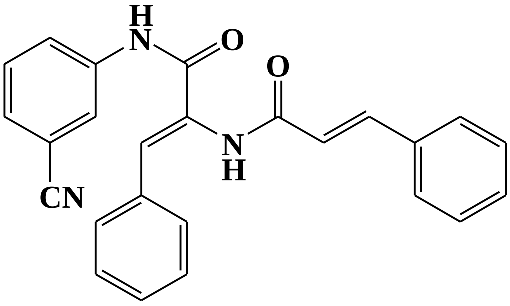

## LC/MS

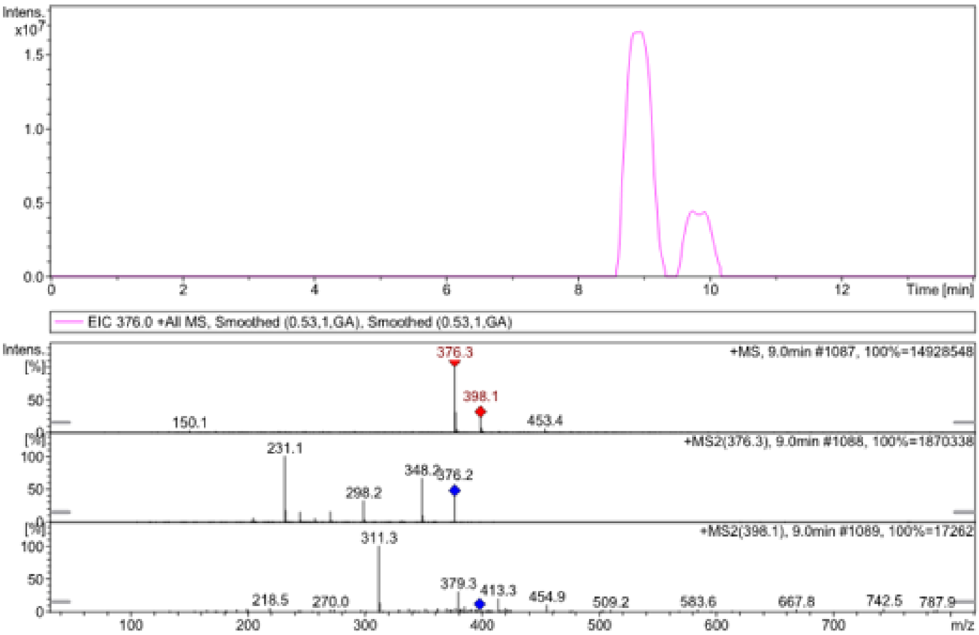


#### $^1$H NMR

This report was created by ACD/NMR Processor Academic Edition. For more information go to www.acdlabs.com/nmrproc/

**1513**

2018-10-06 5:28:54 PM

| Formula | $C_{25}H_{19}N_3O_2$ | FW | 393.4373 |
|---|---|---|---|

| | | | | | | | |
|---|---|---|---|---|---|---|---|
| Acquisition Time (sec) | 1.9268 | Comment | Dr.Abdelsattar Omar Sample: CINB-12A DMSO | | | Date | 22 Dec 2014 14:18:24 |
| Date Stamp | 22 Dec 2014 14:18:24 | | | | | | |
| File Name | E:\AAA Research\Spectra Reservoir\NMR 20141223\Dr.Omar CIN-13A 22-12-2014\20\fid | | | | | Frequency (MHz) | 850.15 |
| Nucleus | 1H | Number of Transients | 16 | Origin | spect | Original Points Count | 32768 |
| Owner | nmr | Points Count | 32768 | Pulse Sequence | zg30 | Receiver Gain | 7.13 |
| SW(cyclical) (Hz) | 17006.80 | Solvent | DMSO-d6 | Spectrum Offset (Hz) | 5250.0283 | Spectrum Type | STANDARD |
| Sweep Width (Hz) | 17006.28 | Temperature (degree C) | 25.000 | | | | |

$^1$H NMR (850 MHz, DMSO-$d_6$) δ ppm 10.52 (br. s., 1 H) 10.05 (br. s., 1 H) 8.19 (s, 1 H) 8.01 (d, $J$=7.78 Hz, 1 H) 7.60 - 7.66 (m, 4 H) 7.50 - 7.58 (m, 3 H) 7.40 - 7.48 (m, 5 H) 7.34 - 7.39 (m, 1 H) 6.98 (s, 1 H) 6.93 (d, $J$=16.09 Hz, 1 H)

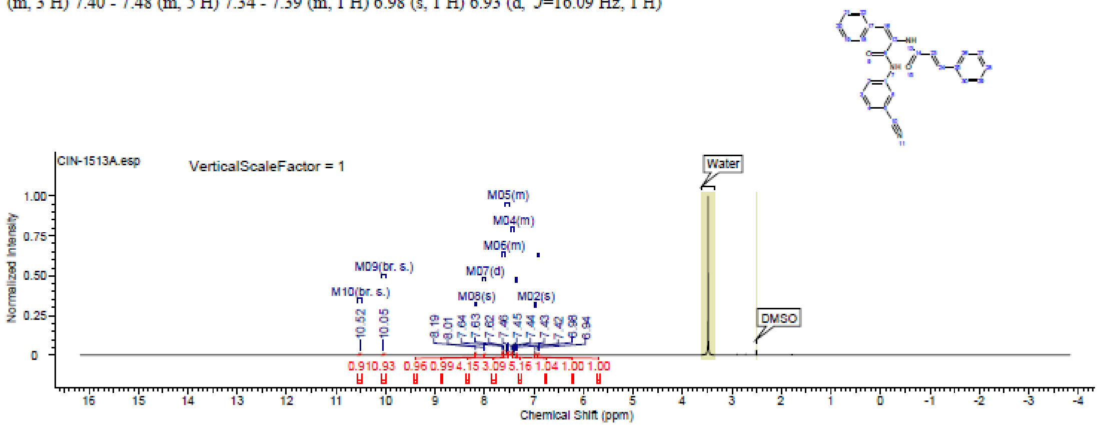

### Spectra of (1528)

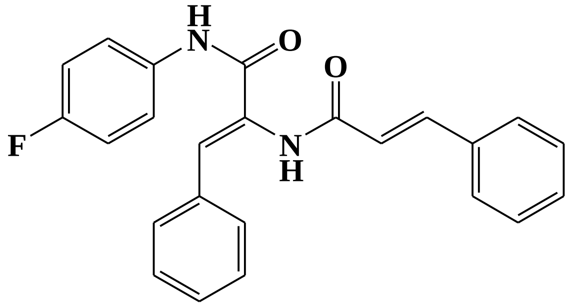


## FT-IR

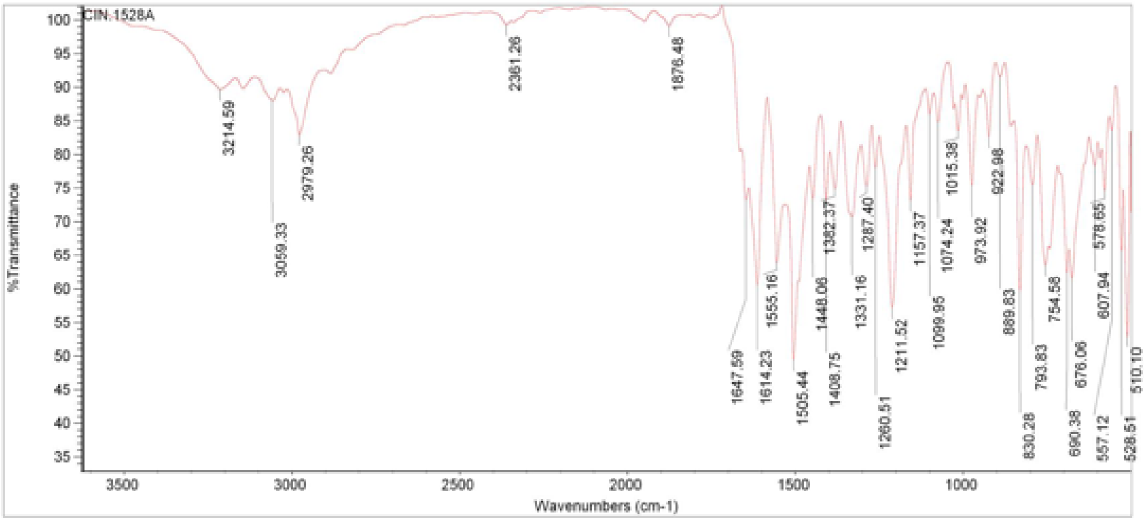


## LC/MS

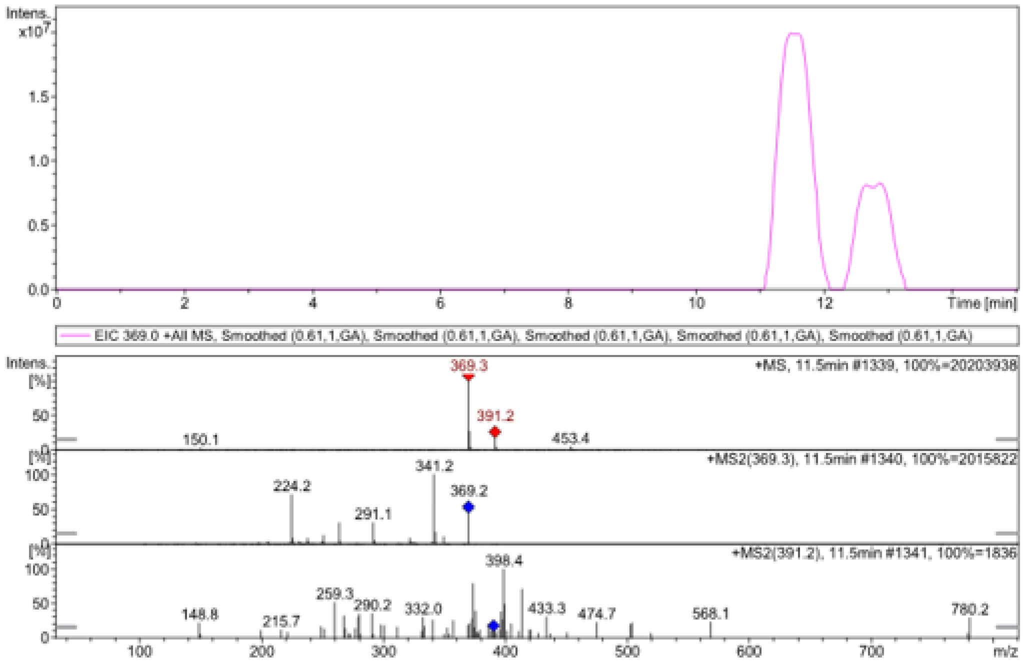

#### $^{1}$H NMR

This report was created by ACD/NMR Processor Academic Edition. For more information go to www.acdlabs.com/nmrproc/

### 1528

2018-10-01 8:02:14 AM

| | |
|---|---|
| Formula $C_{24}H_{19}FN_2O_2$ | FW 386.4183 |

| | | | | | | | |
|---|---|---|---|---|---|---|---|
| Acquisition Time (sec) | 2.6564 | Comment | Dr A Mansour DMSO CIN-1528A | | | Date | 09 Jun 2013 12:06:08 |
| Date Stamp | 09 Jun 2013 12:06:08 | | | | | | |
| File Name | D:\AAA_Research\AAA_Ongoing\CINN\Spectra\NMR\CIN15Series_NMR_20130707\CIN15series_1513to31_20130707\Cin-1528A\70\fid | | | | | | |
| Frequency (MHz) | 600.13 | Nucleus | 1H | Number of Transients | 16 | Origin | spect |
| Original Points Count | 32768 | Owner | nmr | Points Count | 32768 | Pulse Sequence | zg30 |
| Receiver Gain | 15.76 | SW(cyclical) (Hz) | 12335.53 | Solvent | DMSO-d6 | Spectrum Offset (Hz) | 3706.0483 |
| Spectrum Type | STANDARD | Sweep Width (Hz) | 12335.15 | Temperature (degree C) | 25.008 | | |

$^{1}$H NMR (600 MHz, DMSO-$d_6$) δ ppm 6.91 (d, $J$=15.81 Hz, 1 H) 6.96 (s, 1 H) 7.13 - 7.20 (m, 3 H) 7.33 - 7.37 (m, 1 H) 7.39 - 7.48 (m, 8 H) 7.51 (d, $J$=15.81 Hz, 1 H) 7.62 (t, $J$=6.96 Hz, 5 H) 7.68 - 7.74 (m, 2 H) 9.92 (s, 1 H) 10.18 (s, 1 H)

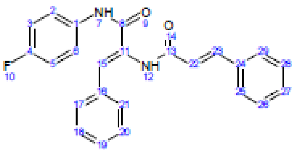


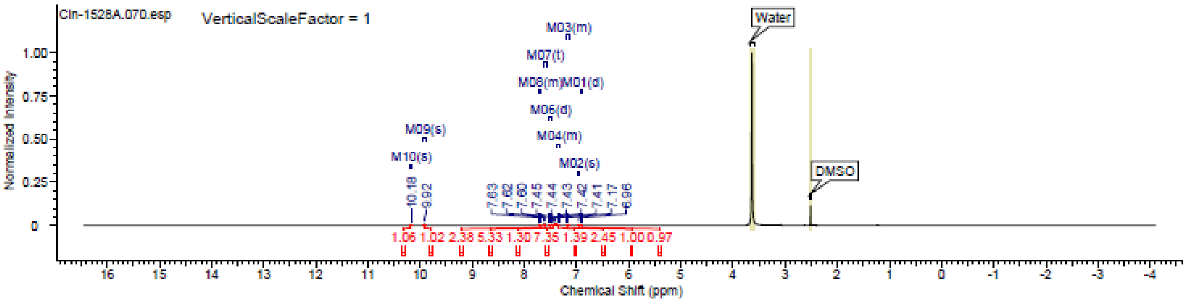

#### Spectra of (1530)

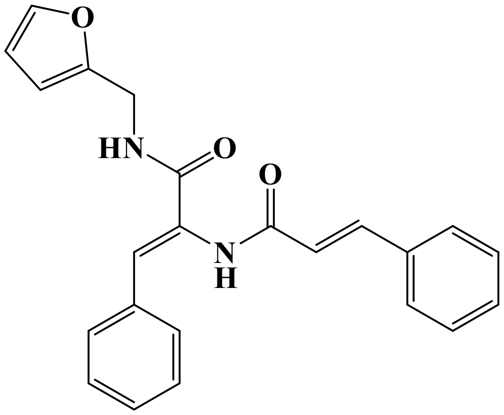

### LC/MS

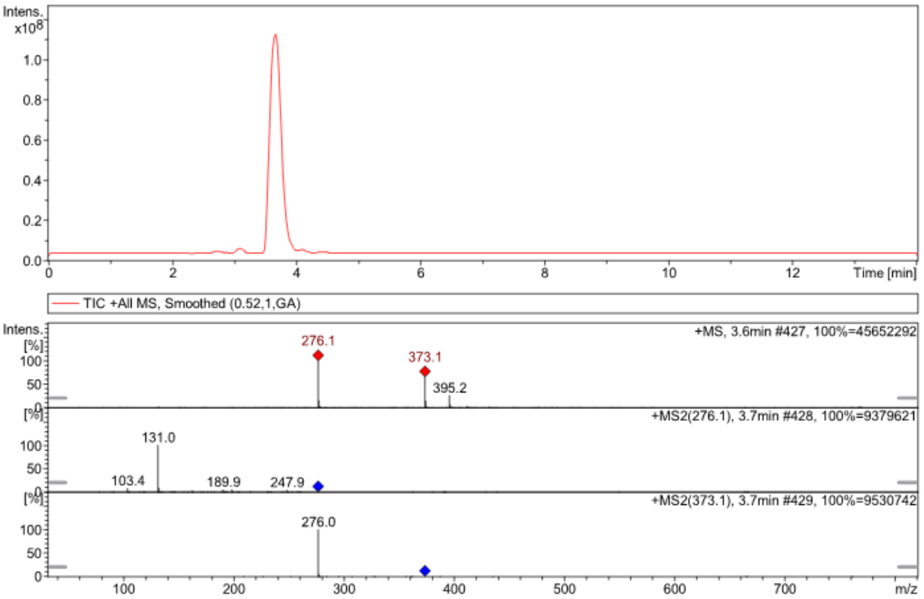


##### $^1$H NMR

**Cin-1530A**

1/29/2015 10:48:51 م

| | | | | | | | |
|---|---|---|---|---|---|---|---|
| ***Acquisition Time (sec)*** | 2.6564 | ***Comment*** | Dr.Mansour DMSO CIN-1530A | | | ***Date*** | 07 Jul 2013 14:16:16 |
| ***Date Stamp*** | 07 Jul 2013 14:16:16 | | | | | | |
| ***File Name*** | D:\AAA_Research\AAA_Ongoing\CINN\Spectra\NMR\CIN15Series_NMR_20130707\CIN15series_1513to31_20130707\CIN-1530A\80\fid | | | | | | |
| ***Frequency (MHz)*** | 600.13 | ***Nucleus*** | 1H | ***Number of Transients*** | 16 | ***Origin*** | spect |
| ***Original Points Count*** | 32768 | ***Owner*** | nmr | ***Points Count*** | 32768 | ***Pulse Sequence*** | zg30 |
| ***Receiver Gain*** | 34.94 | ***SW(cyclical) (Hz)*** | 12335.53 | ***Solvent*** | DMSO-d6 | ***Spectrum Offset (Hz)*** | 3706.0493 |
| ***Spectrum Type*** | STANDARD | ***Sweep Width (Hz)*** | 12335.15 | ***Temperature (degree C)*** | 25.002 | | |

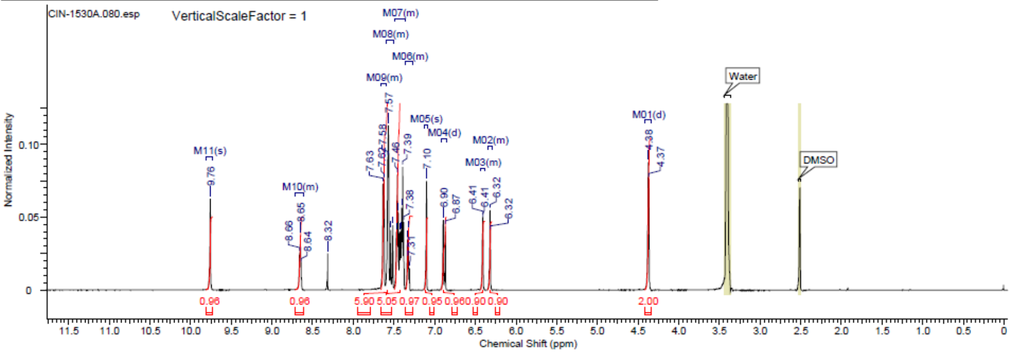


| No. | (ppm) | (Hz) | Height |
|---|---|---|---|
| 1 | 4.37 | 2620.9 | 0.0761 |
| 2 | 4.38 | 2626.6 | 0.0877 |
| 3 | 6.32 | 3790.6 | 0.0401 |
| 4 | 6.32 | 3792.8 | 0.0547 |
| 5 | 6.32 | 3793.6 | 0.0547 |
| 6 | 6.41 | 3845.5 | 0.0484 |
| 7 | 6.41 | 3846.7 | 0.0502 |
| 8 | 6.41 | 3848.5 | 0.0496 |
| 9 | 6.87 | 4122.6 | 0.0436 |
| 10 | 6.90 | 4138.4 | 0.0481 |
| 11 | 7.10 | 4262.3 | 0.0748 |
| 12 | 7.31 | 4387.6 | 0.0148 |
| 13 | 7.31 | 4388.7 | 0.0135 |
| 14 | 7.32 | 4395.1 | 0.0414 |
| 15 | 7.34 | 4402.3 | 0.0348 |
| 16 | 7.34 | 4403.4 | 0.0308 |
| 17 | 7.38 | 4429.4 | 0.0465 |
| 18 | 7.39 | 4437.3 | 0.0846 |
| 19 | 7.41 | 4444.8 | 0.0565 |
| 20 | 7.42 | 4452.4 | 0.0406 |
| 21 | 7.43 | 4459.9 | 0.0399 |
| 22 | 7.45 | 4469.7 | 0.0533 |
| 23 | 7.46 | 4477.2 | 0.0790 |
| 24 | 7.47 | 4484.0 | 0.0399 |
| 25 | 7.51 | 4510.0 | 0.0444 |
| 26 | 7.54 | 4525.8 | 0.0412 |
| 27 | 7.57 | 4542.3 | 0.1130 |
| 28 | 7.57 | 4543.1 | 0.1103 |
| 29 | 7.58 | 4547.6 | 0.0913 |
| 30 | 7.62 | 4574.7 | 0.0756 |
| 31 | 7.63 | 4581.9 | 0.0728 |
| 32 | 7.64 | 4582.6 | 0.0726 |
| 33 | 8.32 | 4991.8 | 0.0249 |
| 34 | 8.64 | 5186.4 | 0.0177 |
| 35 | 8.65 | 5192.1 | 0.0376 |
| 36 | 8.66 | 5197.7 | 0.0252 |
| 37 | 9.76 | 5856.5 | 0.0626 |

#### Spectra of (1531)

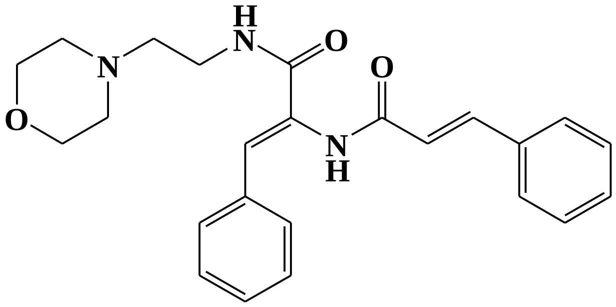


**LC/MS**

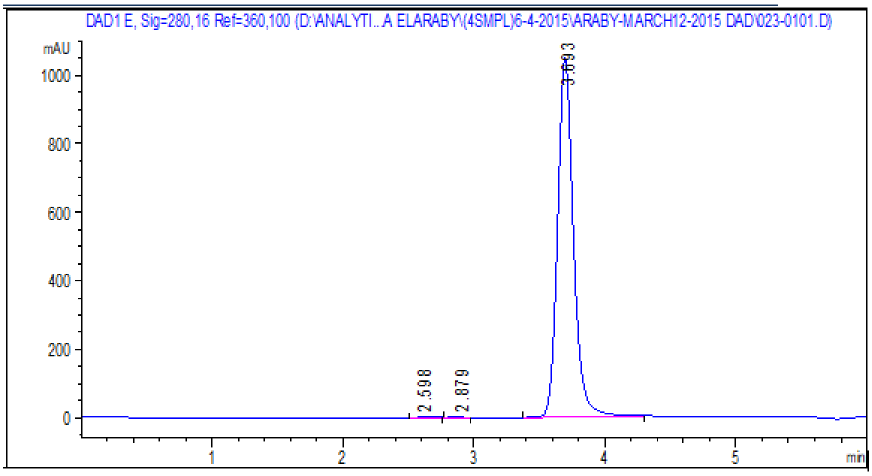


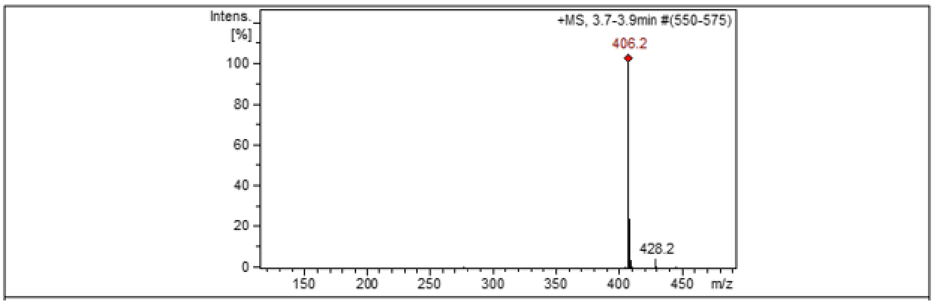


**¹H NMR**

This report was created by ACD/NMR Processor Academic Edition. For more information go to www.acdlabs.com/nmrproc/

#### Cin-1531A

4/6/2015 02:51:29 م

| | | | | | | | |
|---|---|---|---|---|---|---|---|
| *Acquisition Time (sec)* | 2.6564 | *Comment* | Dr.A.Mansour DMSO RAKAN 51A | | | *Date* | 25 Feb 2015 14:29:04 |
| *Date Stamp* | 25 Feb 2015 14:29:04 | | | *File Name* | E:\Google Drive\Projects\CIN-Extension\Spectra\Rakan\RAKAN 51A\1\fid | | |
| *Frequency (MHz)* | 600.13 | *Nucleus* | 1H | *Number of Transients* | 16 | *Origin* | spect |
| *Original Points Count* | 32768 | *Owner* | nmr | *Points Count* | 32768 | *Pulse Sequence* | zg30 |
| *Receiver Gain* | 99.00 | *SW(cyclical) (Hz)* | 12335.53 | *Solvent* | DMSO-d6 | *Spectrum Offset (Hz)* | 3706.0500 |
| *Spectrum Type* | STANDARD | *Sweep Width (Hz)* | 12335.15 | *Temperature (degree C)* | 25.021 | | |

$^1$H NMR (600 MHz, DMSO-$d_6$) δ ppm 2.37 - 2.47 (m, 6 H) 3.25 - 3.32 (m, 2 H) 3.52 - 3.61 (m, 4 H) 6.88 (d, $J$=16.19 Hz, 1 H) 7.07 (s, 1 H) 7.29 - 7.36 (m, 1 H) 7.37 - 7.50 (m, 5 H) 7.53 (d, $J$=15.81 Hz, 1 H) 7.56 (d, $J$=7.53 Hz, 2 H) 7.63 (d, $J$=7.15 Hz, 2 H) 7.96 (br. s., 1 H) 9.74 (s, 1 H)

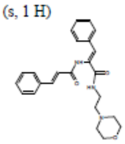

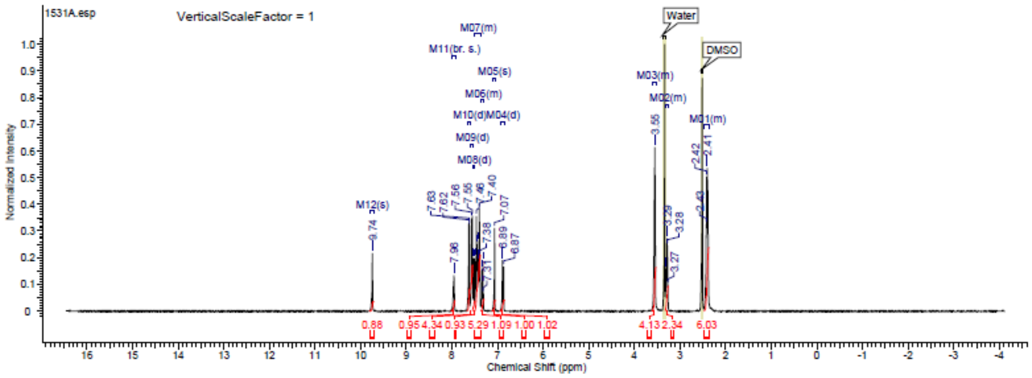

#### Spectra of (1532)

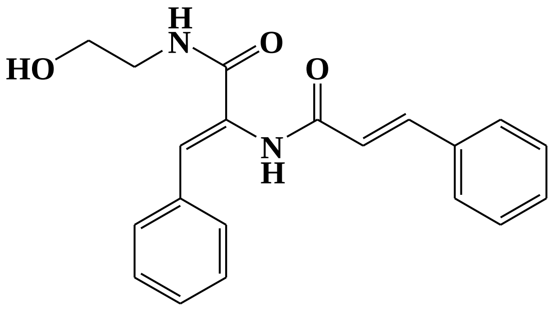


**LC/MS**

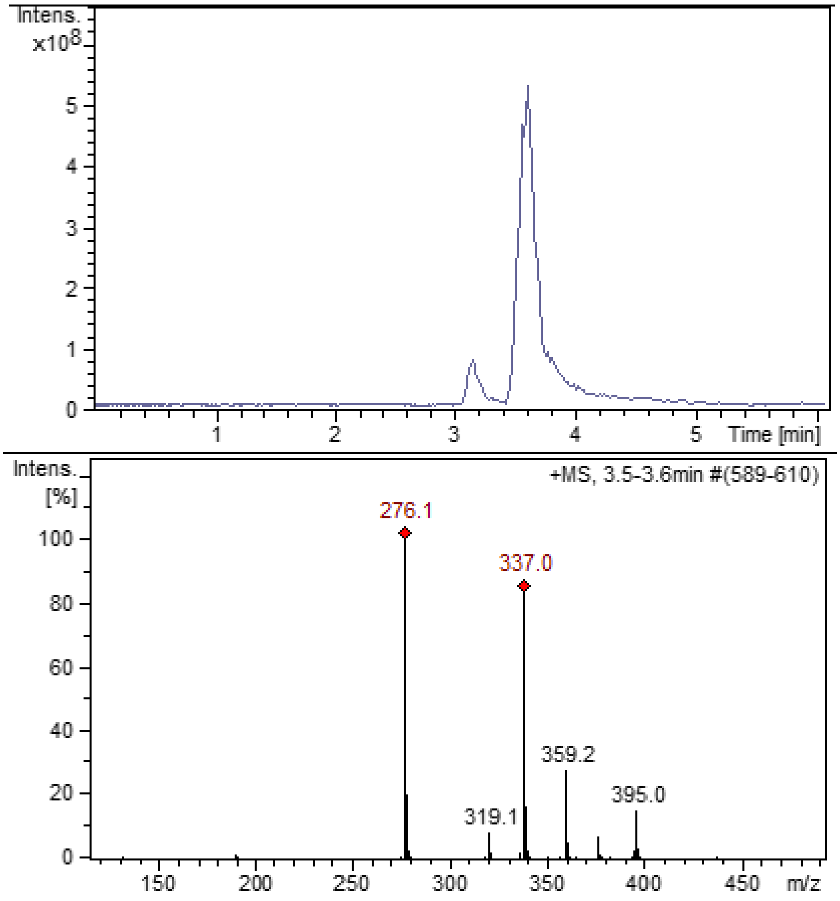

#### $^{1}$H NMR

This report was created by ACD/NMR Processor Academic Edition. For more information go to www.acdlabs.com/nmrproc/

##### Cin-1532A

4/7/2015 05:03:02 م

| | | | | | | | |
|---|---|---|---|---|---|---|---|
| *Acquisition Time (sec)* | 2.6564 | *Comment* | Dr.A.Mansour DMSO RAKAN 1532A | | | *Date* | 25 Feb 2015 14:41:52 |
| *Date Stamp* | 25 Feb 2015 14:41:52 | | | *File Name* | E:\Google Drive\Projects\CIN-Extension\Spectra\Rakan\RAKAN 1532A\1\fid | | |
| *Frequency (MHz)* | 600.13 | *Nucleus* | 1H | *Number of Transients* | 16 | *Origin* | spect |
| *Original Points Count* | 32768 | *Owner* | nmr | *Points Count* | 32768 | *Pulse Sequence* | zg30 |
| *Receiver Gain* | 34.94 | *SW(cyclical) (Hz)* | 12335.53 | *Solvent* | DMSO-d6 | *Spectrum Offset (Hz)* | 3706.0500 |
| *Spectrum Type* | STANDARD | *Sweep Width (Hz)* | 12335.15 | *Temperature (degree C)* | 25.000 | | |

$^{1}$H NMR (600 MHz, DMSO-$d_6$) δ ppm 2.58 (t, J=5.65 Hz, 1 H) 3.26 (q, J=6.02 Hz, 2 H) 3.36 (t, J=5.83 Hz, 2 H) 3.48 (t, J=6.21 Hz, 2 H) 6.89 (d, J=15.81 Hz, 1 H) 7.07 (s, 1 H) 7.32 (d, J=7.53 Hz, 1 H) 7.39 (t, J=7.53 Hz, 2 H) 7.43 (d, J=7.15 Hz, 1 H) 7.44 - 7.48 (m, 1 H) 7.52 (d, J=15.81 Hz, 1 H) 7.56 (d, J=7.91 Hz, 1 H) 7.63 (d, J=7.53 Hz, 1 H) 8.10 (s, 1 H)

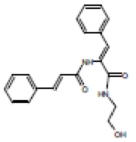


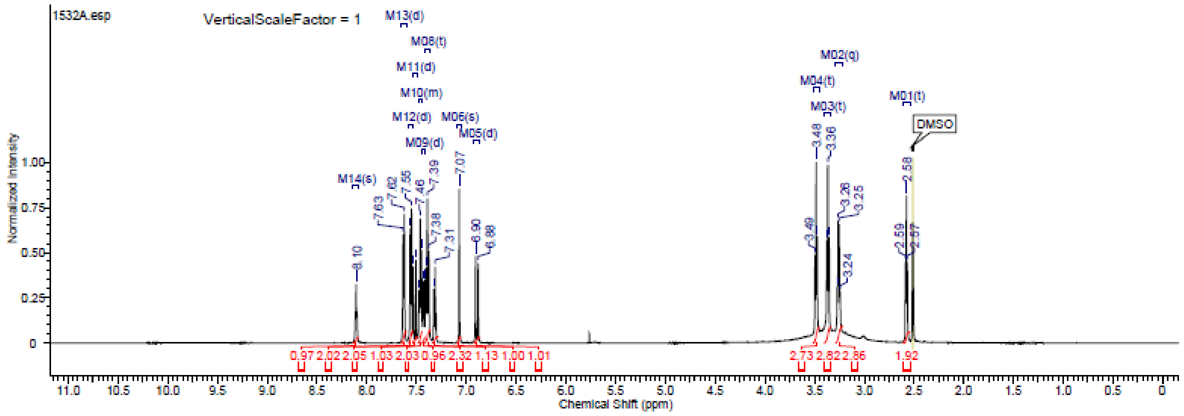

### Spectra of (1535)

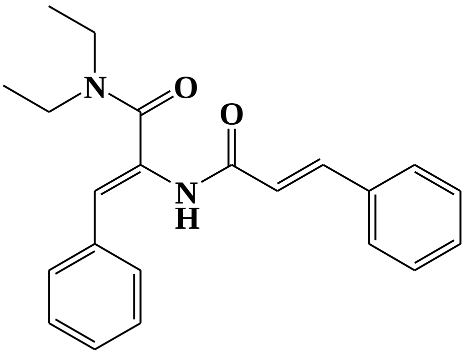


#### $^1$H NMR

This report was created by ACD/NMR Processor Academic Edition. For more information go to www.acdlabs.com/nmrproc/

**Cin-1535A**

4/7/2015 05:00:33 م

| | | | | | | | |
|---|---|---|---|---|---|---|---|
| *Acquisition Time (sec)* | 2.6564 | *Comment* | Dr.A.Mansour DMSO RAKAN 1535A | | | *Date* | 25 Feb 2015 14:20:32 |
| *Date Stamp* | 25 Feb 2015 14:20:32 | | | *File Name* | E:\Google Drive\Projects\CIN-Extension\Spectra\Rakan\RAKAN 1535A\1\fid | | |
| *Frequency (MHz)* | 600.13 | *Nucleus* | 1H | *Number of Transients* | 16 | *Origin* | spect |
| *Original Points Count* | 32768 | *Owner* | nmr | *Points Count* | 32768 | *Pulse Sequence* | zg30 |
| *Receiver Gain* | 37.66 | *SW(cyclical) (Hz)* | 12335.53 | *Solvent* | DMSO-d6 | *Spectrum Offset (Hz)* | 3706.0493 |
| *Spectrum Type* | STANDARD | *Sweep Width (Hz)* | 12335.15 | *Temperature (degree C)* | 24.993 | | |

$^1$H NMR (600 MHz, DMSO-$d_6$) δ ppm 1.14 (br. s., 6 H) 3.26 - 3.43 (m, 3 H) 6.10 (br. s., 1 H) 6.89 - 6.98 (m, 1 H) 7.27 - 7.35 (m, 1 H) 7.36 - 7.49 (m, 5 H) 7.52 - 7.59 (m, 3 H) 7.60 (d, J=7.15 Hz, 2 H) 9.98 (br. s., 1 H)

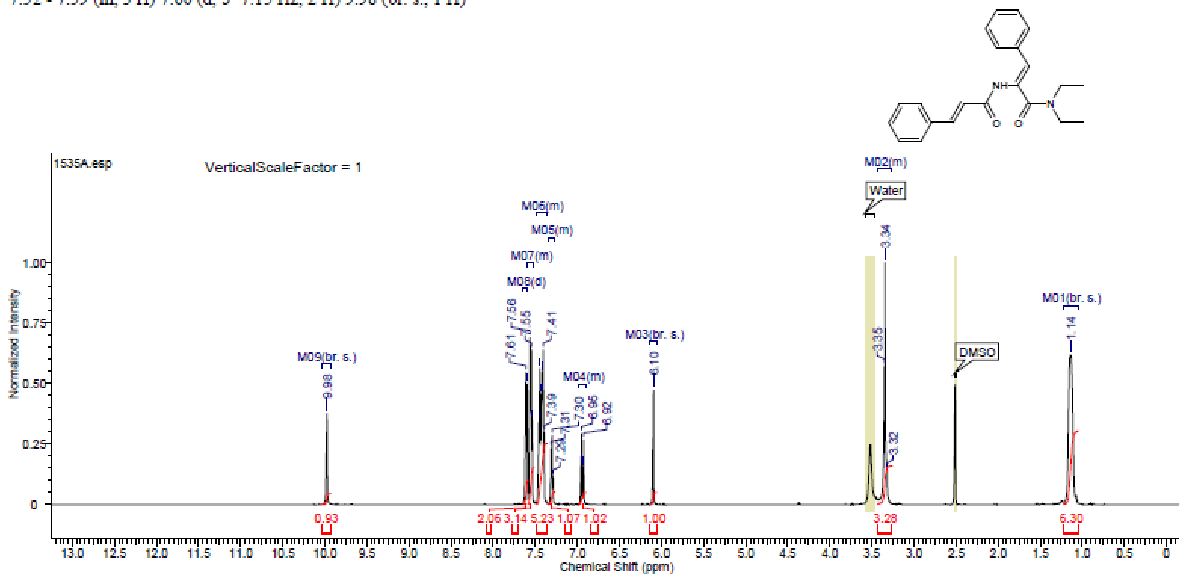

#### Spectra of (1536)

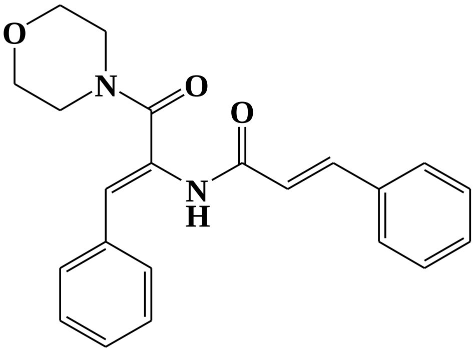

##### $^1$H NMR

This report was created by ACD/NMR Processor Academic Edition. For more information go to www.acdlabs.com/nmrproc/

**Cin-1536A**

4/7/2015 05:06:29 م

| | | | | | | | |
|---|---|---|---|---|---|---|---|
| Acquisition Time (sec) | 2.6564 | Comment | Dr.A.Mansour DMSO RAKAN 1536A | | | Date | 25 Feb 2015 14:22:40 |
| Date Stamp | 25 Feb 2015 14:22:40 | | | File Name | E:\Google Drive\Projects\CIN-Extension\Spectra\Rakan\RAKAN 1536A\1\fid | | |
| Frequency (MHz) | 600.13 | Nucleus | 1H | Number of Transients | 16 | Origin | spect |
| Original Points Count | 32768 | Owner | nmr | Points Count | 32768 | Pulse Sequence | zg30 |
| Receiver Gain | 37.66 | SW(cyclical) (Hz) | 12335.53 | Solvent | DMSO-d6 | Spectrum Offset (Hz) | 3706.0500 |
| Spectrum Type | STANDARD | Sweep Width (Hz) | 12335.15 | Temperature (degree C) | 25.000 | | |

$^1$H NMR (600 MHz, DMSO-$d_6$) δ ppm 3.35 (br. s., 1 H) 3.53 (br. s., 3 H) 6.19 (s, 1 H) 6.93 (d, $J$=15.81 Hz, 1 H) 7.27 - 7.36 (m, 2 H) 7.40 - 7.48 (m, 6 H) 7.56 - 7.63 (m, 6 H) 10.07 (s, 1 H)

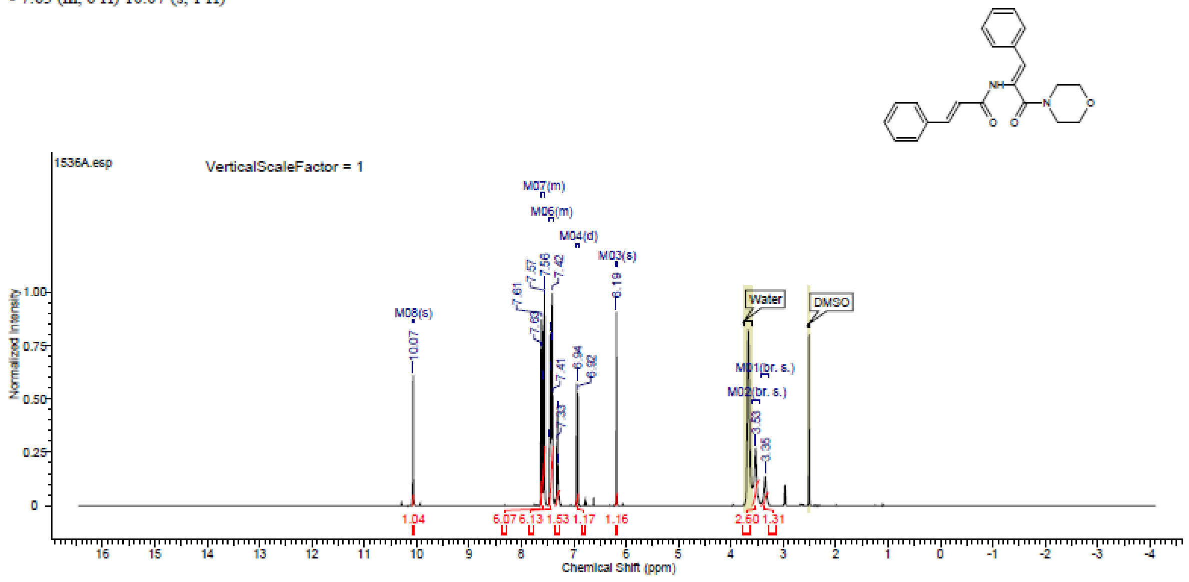

### Spectra of (1555)

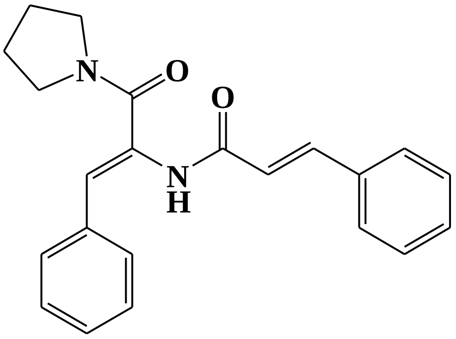

#### $^1$H NMR

This report was created by ACD/NMR Processor Academic Edition. For more information go to www.acdlabs.com/nmrproc/

##### Cin-1555A

4/7/2015 05:09:42

| | | | | | |
|---|---|---|---|---|---|
| *Acquisition Time (sec)* | 2.6564 | *Comment* | Dr.A.Mansour DMSO RAKAN 1555A | *Date* | 25 Feb 2015 14:58:56 |
| *Date Stamp* | 25 Feb 2015 14:58:56 | *File Name* | E:\Google Drive\Projects\CIN-Extension\Spectra\Rakan\RAKAN 1555A\1\fid | | |
| *Frequency (MHz)* | 600.13 | *Nucleus* | 1H | *Number of Transients* | 16 |
| *Origin* | spect | | | | |
| *Original Points Count* | 32768 | *Owner* | nmr | *Points Count* | 32768 |
| *Pulse Sequence* | zg30 | | | | |
| *Receiver Gain* | 87.48 | *SW(cyclical) (Hz)* | 12335.53 | *Solvent* | DMSO-d6 |
| *Spectrum Offset (Hz)* | 3706.0500 | | | | |
| *Spectrum Type* | STANDARD | *Sweep Width (Hz)* | 12335.15 | *Temperature (degree C)* | 24.992 |

$^1$H NMR (600 MHz, DMSO-$d_6$) δ ppm 1.83 - 1.87 (m, 18 H) 3.32 (br. s., 1 H) 3.37 (t, J=6.59 Hz, 9 H) 3.58 (t, J=6.02 Hz, 8 H) 6.32 (s, 4 H) 6.90 (s, 2 H) 6.92 (s, 2 H) 7.28 - 7.33 (m, 6 H) 7.40 - 7.47 (m, 24 H) 7.54 (s, 2 H) 7.56 - 7.58 (m, 11 H) 7.61 (d, J=6.78 Hz, 9 H) 10.00 (s, 3 H)

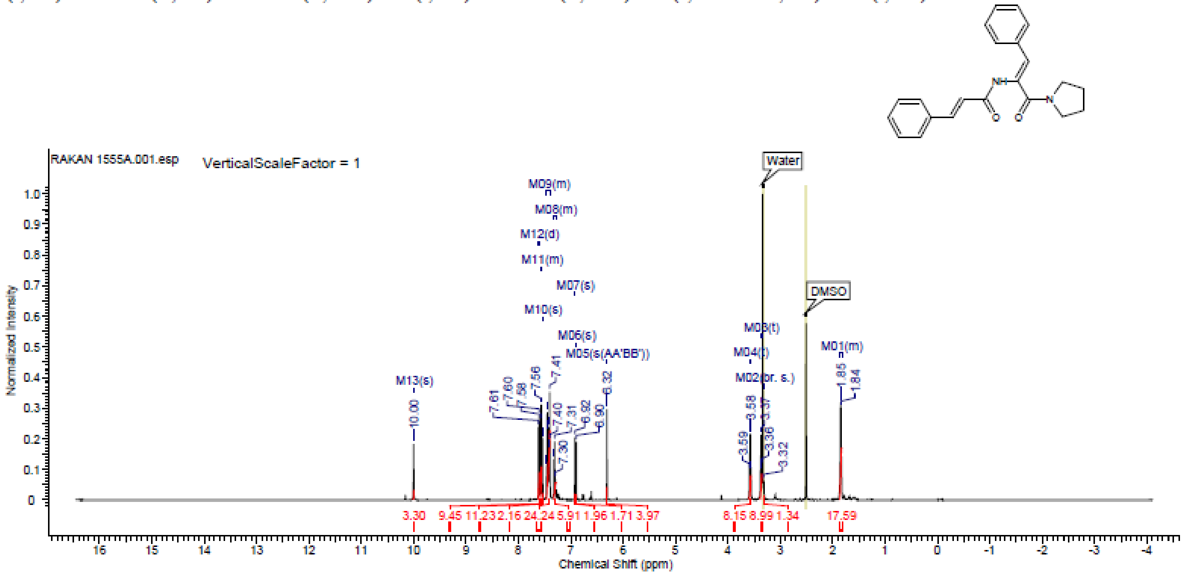

### Spectra of (1556)

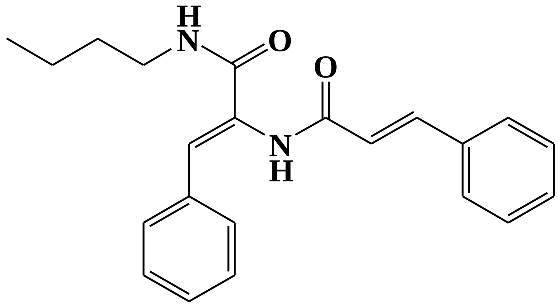

#### $^{1}$H NMR

This report was created by ACD/NMR Processor Academic Edition. For more information go to www.acdlabs.com/nmrproc/

##### Cin-1556A

4/7/2015 05:40:40

| | | | | | | | |
|---|---|---|---|---|---|---|---|
| *Acquisition Time (sec)* | 2.6564 | *Comment* | Dr.A.Mansour DMSO RAKAN 1556A | | | *Date* | 25 Feb 2015 14:33:20 |
| *Date Stamp* | 25 Feb 2015 14:33:20 | | | *File Name* | E:\Google Drive\Projects\CIN-Extension\Spectra\Rakan\RAKAN 1556A\1\fid | | |
| *Frequency (MHz)* | 600.13 | *Nucleus* | 1H | *Number of Transients* | 16 | *Origin* | spect |
| *Original Points Count* | 32768 | *Owner* | nmr | *Points Count* | 32768 | *Pulse Sequence* | zg30 |
| *Receiver Gain* | 43.27 | *SW(cyclical) (Hz)* | 12335.53 | *Solvent* | DMSO-d6 | *Spectrum Offset (Hz)* | 3706.0500 |
| *Spectrum Type* | STANDARD | *Sweep Width (Hz)* | 12335.15 | *Temperature (degree C)* | 24.996 | | |

$^{1}$H NMR (600 MHz, DMSO-$d_6$) δ ppm 0.90 (t, *J*=7.34 Hz, 3 H) 1.28 - 1.37 (m, 2 H) 1.43 - 1.51 (m, 2 H) 3.17 (q, *J*=6.78 Hz, 2 H) 6.88 (d, *J*=16.19 Hz, 1 H) 7.01 (s, 1 H) 7.31 (t, *J*=7.34 Hz, 1 H) 7.36 - 7.44 (m, 3 H) 7.44 - 7.48 (m, 2 H) 7.51 (d, *J*=15.81 Hz, 1 H) 7.56 (d, *J*=7.53 Hz, 2 H) 7.62 (d, *J*=7.15 Hz, 2 H) 8.10 (t, *J*=5.83 Hz, 1 H) 9.69 (s, 1 H)

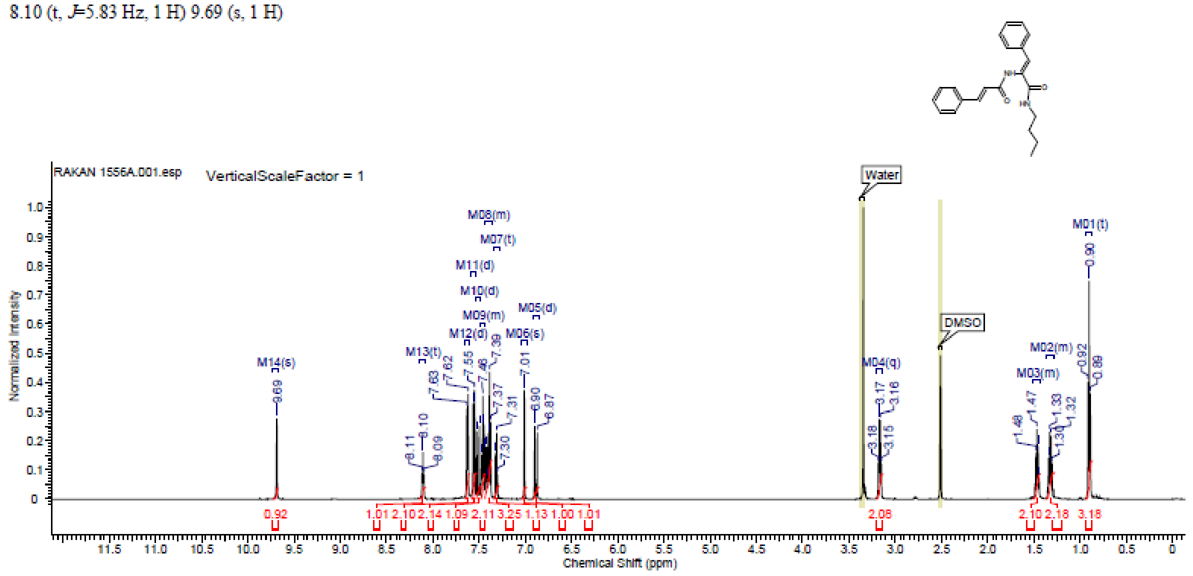

### Spectra of (1557)

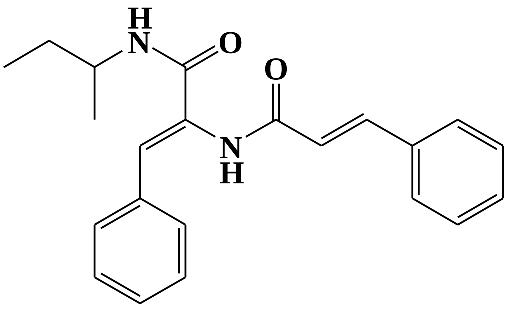

#### $^1$H NMR

This report was created by ACD/NMR Processor Academic Edition. For more information go to www.acdlabs.com/nmrproc/

**Cin-1557A**

4/7/2015 05:47:10 م

| | | | | | | | |
|---|---|---|---|---|---|---|---|
| *Acquisition Time (sec)* | 2.6564 | *Comment* | Dr.A.Mansour DMSO RAKAN 1557A | | | *Date* | 25 Feb 2015 15:03:12 |
| *Date Stamp* | 25 Feb 2015 15:03:12 | | | *File Name* | E:\Google Drive\Projects\CIN-Extension\Spectra\Rakan\RAKAN 1557A\1\fid | | |
| *Frequency (MHz)* | 600.13 | *Nucleus* | 1H | *Number of Transients* | 16 | *Origin* | spect |
| *Original Points Count* | 32768 | *Owner* | nmr | *Points Count* | 32768 | *Pulse Sequence* | zg30 |
| *Receiver Gain* | 83.44 | *SW(cyclical) (Hz)* | 12335.53 | *Solvent* | DMSO-d6 | *Spectrum Offset (Hz)* | 3706.0500 |
| *Spectrum Type* | STANDARD | *Sweep Width (Hz)* | 12335.15 | *Temperature (degree C)* | 24.996 | | |

$^1$H NMR (600 MHz, DMSO-$d_6$) δ ppm 0.84 - 0.93 (m, 3 H) 1.11 (d, *J*=6.78 Hz, 3 H) 1.39 - 1.59 (m, 2 H) 3.81 (dt, *J*=13.93, 7.34 Hz, 1 H) 6.86 - 6.94 (m, 2 H) 7.28 - 7.34 (m, 1 H) 7.39 (t, *J*=7.72 Hz, 2 H) 7.41 - 7.49 (m, 3 H) 7.52 (d, *J*=15.81 Hz, 1 H) 7.56 (d, *J*=7.91 Hz, 2 H) 7.62 (d, *J*=7.15 Hz, 2 H) 7.84 (d, *J*=8.66 Hz, 1 H) 9.66 (s, 1 H)

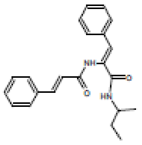

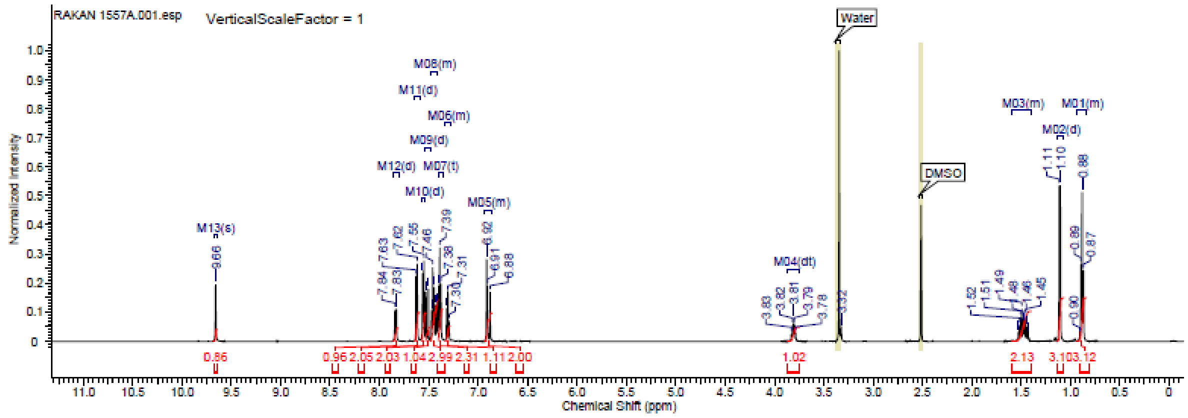

### Spectra of (1559)

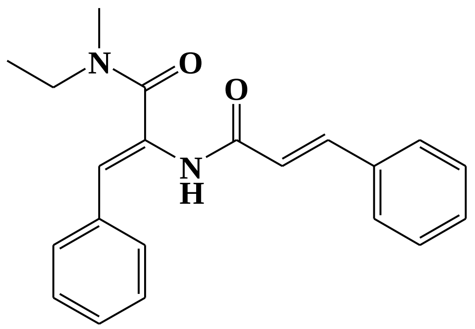


#### $^{1}$H NMR

This report was created by ACD/NMR Processor Academic Edition. For more information go to www.acdlabs.com/nmrproc/

**CIN-1559A-4m**

1/6/2017 05:28:08 PM

| | | | | | | | |
|---|---|---|---|---|---|---|---|
| *Acquisition Time (sec)* | 2.6564 | *Comment* | Dr.A.Mansour DMSO RAKAN 1559A | | | *Date* | 25 Feb 2015 14:50:24 |
| *Date Stamp* | 25 Feb 2015 14:50:24 | | | | | | |
| *File Name* | E:\Google Drive\Projects\CIN\Safo-Cinamamide\Spectra Bis-Cinnamamide\1H NMR\SecondBatch-Table4I-4t\ConfirmedCpdsNMR\RAKAN 1559A\1\fid | | | | | | |
| *Frequency (MHz)* | 600.13 | *Nucleus* | 1H | *Number of Transients* | 16 | *Origin* | spect |
| *Original Points Count* | 32768 | *Owner* | nmr | *Points Count* | 32768 | *Pulse Sequence* | zg30 |
| *Receiver Gain* | 30.24 | *SW(cyclical) (Hz)* | 12335.53 | *Solvent* | DMSO-d6 | *Spectrum Offset (Hz)* | 3706.0500 |
| *Spectrum Type* | STANDARD | *Sweep Width (Hz)* | 12335.15 | *Temperature (degree C)* | 24.997 | | |

$^{1}$H NMR (DMSO-$d_6$) δ ppm 10.01 (br. s., 1 H), 7.61 (d, $J$=7.15 Hz, 2 H), 7.52 - 7.59 (m, 3 H), 7.37 - 7.49 (m, 5 H), 7.27 - 7.34 (m, 1 H), 6.94 (d, $J$=15.81 Hz, 1 H), 6.09 - 6.20 (m, 1 H), 3.55 (br. s., 1 H), 3.08 (br. s., 1 H), 2.85 (br. s., 1 H), 1.17 (br. s., 1 H), 1.12 (br. s., 2 H)

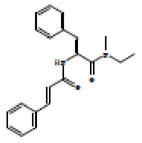

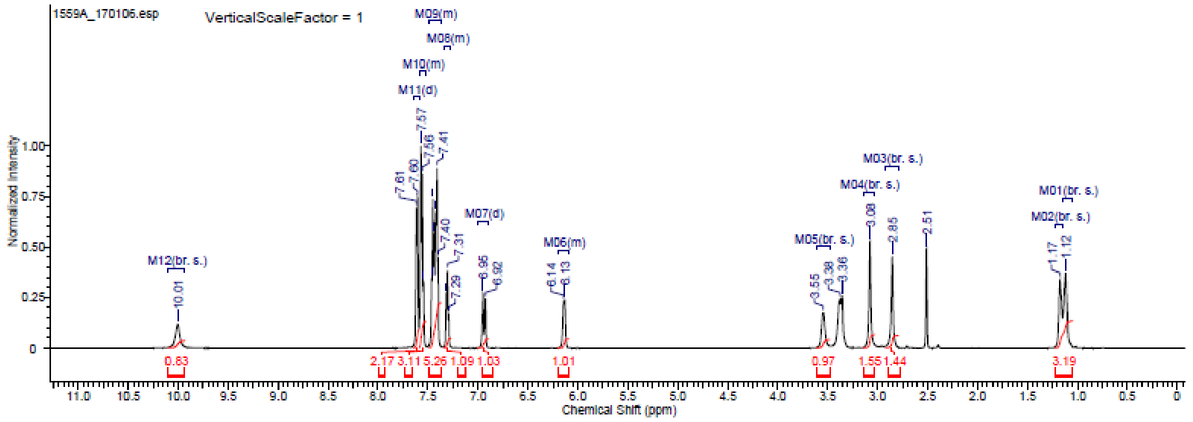

### Spectra of (15EE)[4]

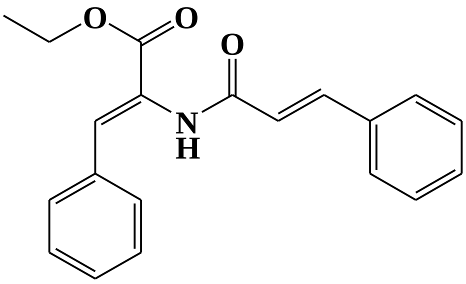

#### $^{1}$H NMR

[illegible]

**CIN-15EE, 5**

12/17/2016 07:09:34 PM

| | | | | | | | |
|---|---|---|---|---|---|---|---|
| *Acquisition Time (sec)* | 2.6564 | *Comment* | Dr A.Mansour Acetone Cin-1522B 27-3-2012 | | | *Date* | 27 Mar 2013 13:46:24 |
| *Date Stamp* | 27 Mar 2013 13:46:24 | | | | | | |
| *File Name* | E:\AAA_Research\AAA_Ongoing\CINN\Spectra\NMR\CIN15Series_NMR_20130707\CIN15series_1513to31_20130707\Cin-1522B\1\fid | | | | | | |
| *Frequency (MHz)* | 600.13 | *Nucleus* | 1H | *Number of Transients* | 16 | *Origin* | spect |
| *Original Points Count* | 32768 | *Owner* | nmr | *Points Count* | 32768 | *Pulse Sequence* | zg30 |
| *Receiver Gain* | 106.17 | *SW(cyclical) (Hz)* | 12335.53 | *Solvent* | Acetone | *Spectrum Offset (Hz)* | 3706.0493 |
| *Spectrum Type* | STANDARD | *Sweep Width (Hz)* | 12335.15 | *Temperature (degree C)* | 25.009 | | |

$^{1}$H NMR (Acetone) δ ppm 8.97 (br. s., 1 H), 7.62 - 7.69 (m, 5 H), 7.34 - 7.50 (m, 6 H), 7.30 (s, 1 H), 6.99 (d, *J*=15.81 Hz, 1 H), 4.27 (q, *J*=7.03 Hz, 2 H), 1.31 (t, *J*=7.15 Hz, 3 H)

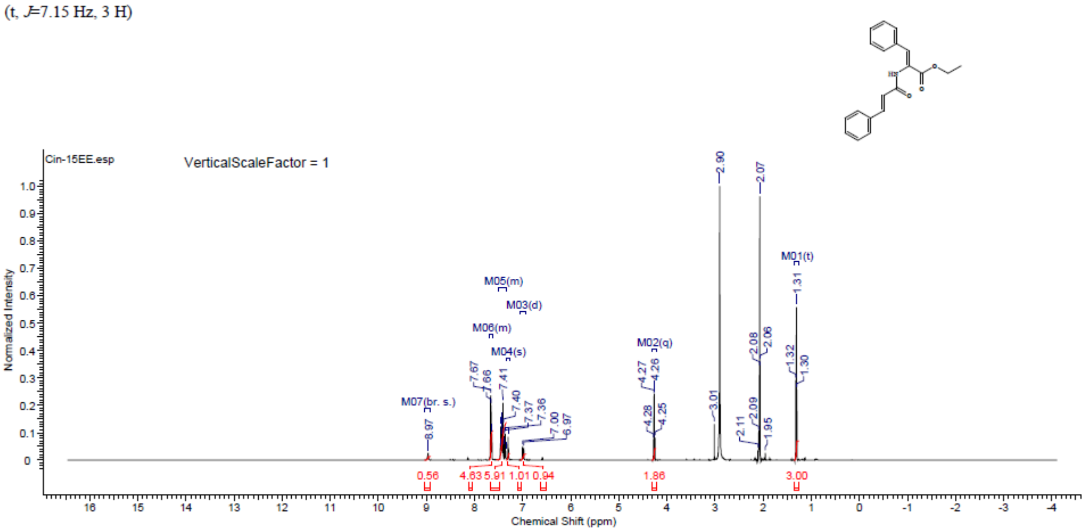


#### $^{13}$C NMR

This report was created by ACD/NMR Processor Academic Edition. For more information go to www.acdlabs.com/nmrproc/

**CIN-15EE**

11/20/2014 01:27:01 م

| | | | | | | | |
|---|---|---|---|---|---|---|---|
| *Acquisition Time (sec)* | 0.4719 | *Comment* | Dr.Mansour CDCl3 Cin15EE C13 | | | *Date* | 14 Nov 2014 07:24:32 |
| *Date Stamp* | 14 Nov 2014 07:24:32 | | | *File Name* | E:\AAA_Research\Spectra Reservoir\CIN_13C_20-11-14\Cin15EE C13\1\fid | | |
| *Frequency (MHz)* | 150.92 | *Nucleus* | 13C | *Number of Transients* | 4096 | *Origin* | spect |
| *Original Points Count* | 16384 | *Owner* | nmr | *Points Count* | 16384 | *Pulse Sequence* | zgpg30 |
| *Receiver Gain* | 173.48 | *SW(cyclical) (Hz)* | 34722.22 | *Solvent* | CHLOROFORM-d | | |
| *Spectrum Offset (Hz)* | 16599.3105 | *Spectrum Type* | STANDARD | *Sweep Width (Hz)* | 34720.10 | *Temperature (degree C)* | 25.001 |

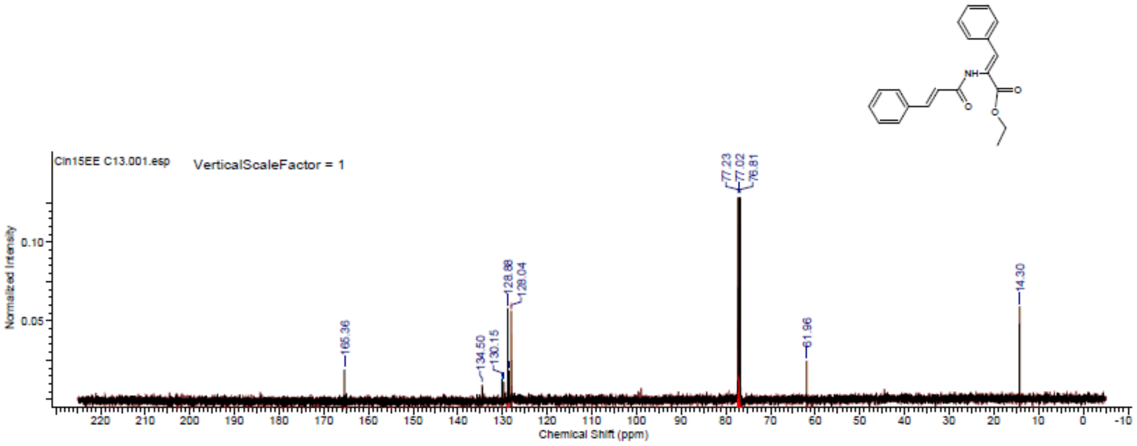

### Spectra of (1612)

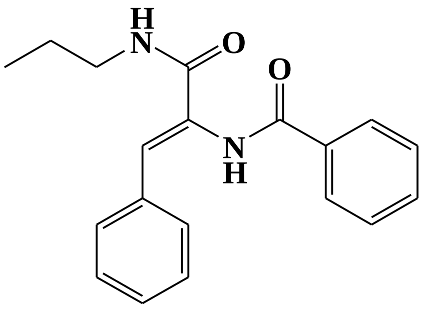


**LC/MS**

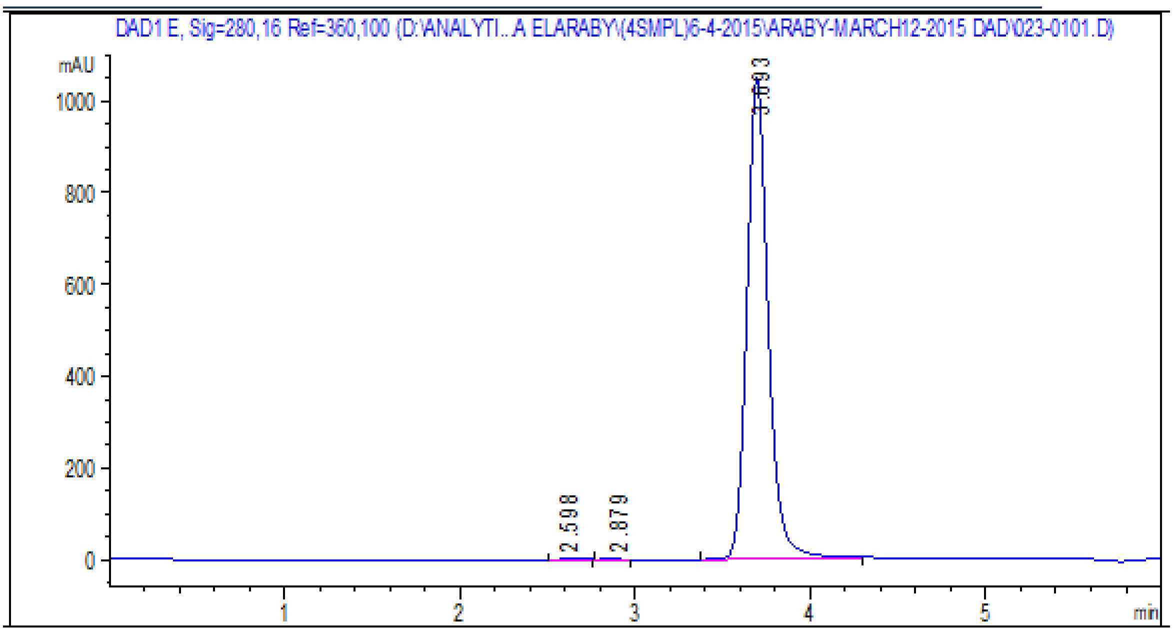


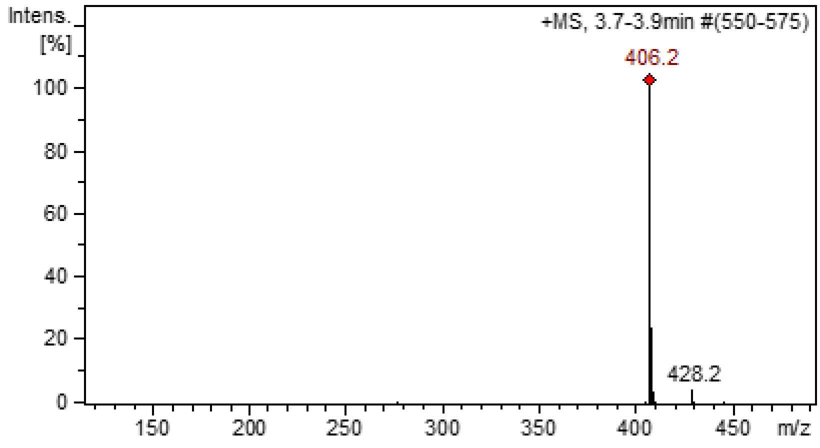

#### $^1$H NMR

This report was created by ACD/NMR Processor Academic Edition. For more information go to www.acdlabs.com/nmrproc/

### 1612

2016-10-06 5:34:48 PM

| Formula $C_{18}H_{20}N_2O_2$ | FW 308.3743 |
|---|---|

| | | | | | | | |
|---|---|---|---|---|---|---|---|
| Acquisition Time (sec) | 1.9266 | Comment | Dr.Mustafa El-Araby Sample : CIN-1612 DMSO | | | Date | 08 Apr 2015 10:15:12 |
| Date Stamp | 08 Apr 2015 10:15:12 | | | | | | |
| File Name | E:\AAA Research\Spectra Reservoir\April2015\Cin 1612 1712 1532\MUSTAFA CIN-1612 08-04-2015\20\fid | | | | | Frequency (MHz) | 850.15 |
| Nucleus | 1H | Number of Transients | 20 | Origin | spect | Original Points Count | 32768 |
| Owner | nmr | Points Count | 32768 | Pulse Sequence | zg30 | Receiver Gain | 10.55 |
| SW(cyclical) (Hz) | 17006.80 | Solvent | DMSO-d6 | Spectrum Offset (Hz) | 5250.0283 | Spectrum Type | STANDARD |
| Sweep Width (Hz) | 17006.28 | Temperature (degree C) | 25.001 | | | | |

$^1$H NMR (850 MHz, DMSO-$d_6$) δ ppm 7.82 - 7.92 (m, 2 H) 7.57 (d, $J$=8.30 Hz, 3 H) 7.50 (t, $J$=7.53 Hz, 2 H) 7.40 (d, $J$=8.30 Hz, 2 H) 7.22 - 7.32 (m, 3 H) 1.60 (s, 3 H) 0.86 (t, $J$=7.27 Hz, 2 H)

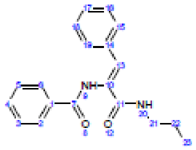

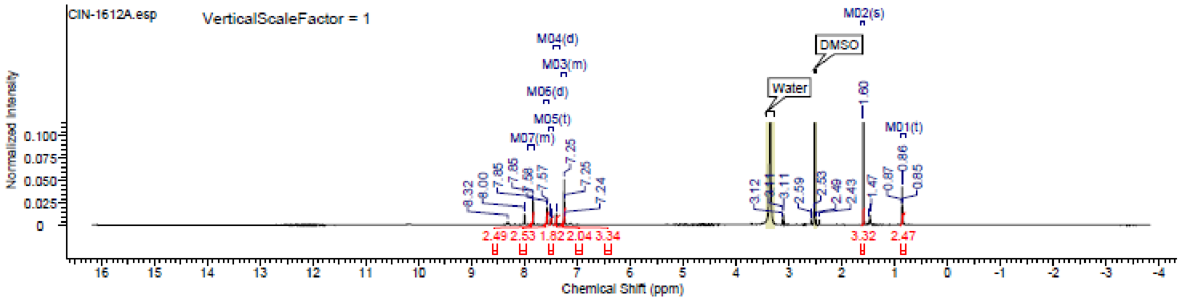

### Spectra of (1712)

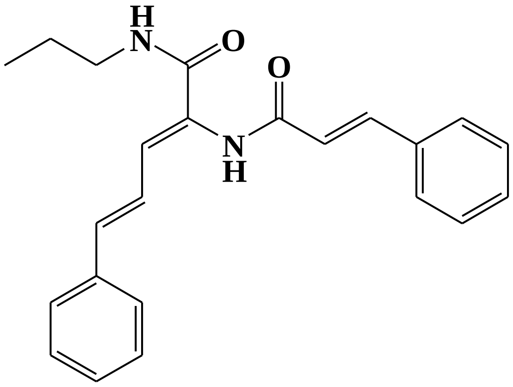

**LC/MS**

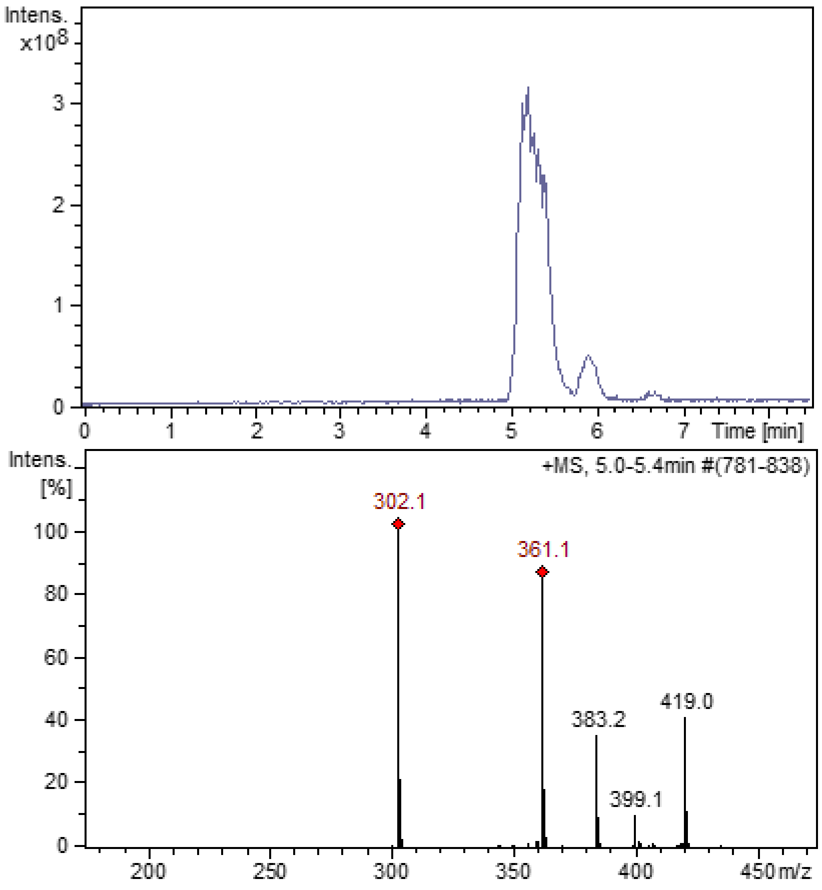

#### $^{1}$H NMR

This report was created by ACD/NMR Processor Academic Edition. For more information go to www.acdlabs.com/nmrproc/

**1712**

2018-10-06 5:35:14 PM

| Formula | $C_{23}H_{24}N_2O_2$ | FW | 360.4489 |
|---|---|---|---|

| Acquisition Time (sec) | 1.9268 | Comment | Dr.Mustafa El-Araby Sample : CUR-1712 DMSO | | | Date | 08 Apr 2015 10:19:28 |
|---|---|---|---|---|---|---|---|
| Date Stamp | 08 Apr 2015 10:19:28 | | | | | | |
| File Name | E:\AAA Research\Spectra Reservoir\April2015\Cin 1612 1712 1532\MUSTAFA CUR-1712 08-04-2015\30\fid | | | | | Frequency (MHz) | 850.15 |
| Nucleus | 1H | Number of Transients | 20 | Origin | spect | Original Points Count | 32768 |
| Owner | nmr | Points Count | 32768 | Pulse Sequence | zg30 | Receiver Gain | 9.04 |
| SW(cyclical) (Hz) | 17006.80 | Solvent | DMSO-d6 | Spectrum Offset (Hz) | 5250.0283 | Spectrum Type | STANDARD |
| Sweep Width (Hz) | 17006.28 | Temperature (degree C) | 24.999 | | | | |

$^{1}$H NMR (850 MHz, DMSO-$d_6$) δ ppm 9.69 (s, 1 H) 8.02 (t, $J$=5.71 Hz, 1 H) 7.64 (d, $J$=7.27 Hz, 2 H) 7.59 - 7.62 (m, 1 H) 7.50 - 7.55 (m, 3 H) 7.44 - 7.48 (m, 3 H) 7.40 - 7.44 (m, 2 H) 7.34 - 7.39 (m, 3 H) 7.28 - 7.31 (m, 1 H) 7.00 - 7.05 (m, 1 H) 6.87 - 6.92 (m, 2 H) 6.73 (d, $J$=10.90 Hz, 1 H) 3.17 - 3.21 (m, 1 H) 3.08 - 3.12 (m, 2 H) 1.53 - 1.60 (m, 1 H) 1.44 - 1.51 (m, 2 H) 0.94 (t, $J$=7.53 Hz, 1 H) 0.87 (t, $J$=7.27 Hz, 3 H)

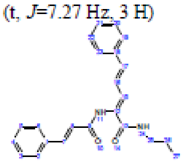

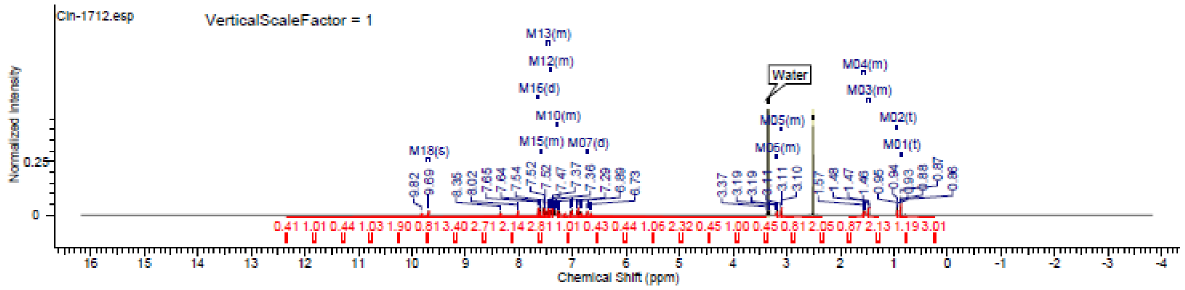

### Spectra of (1812)

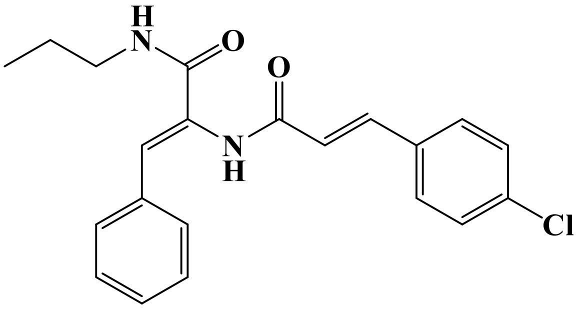


## LC/MS

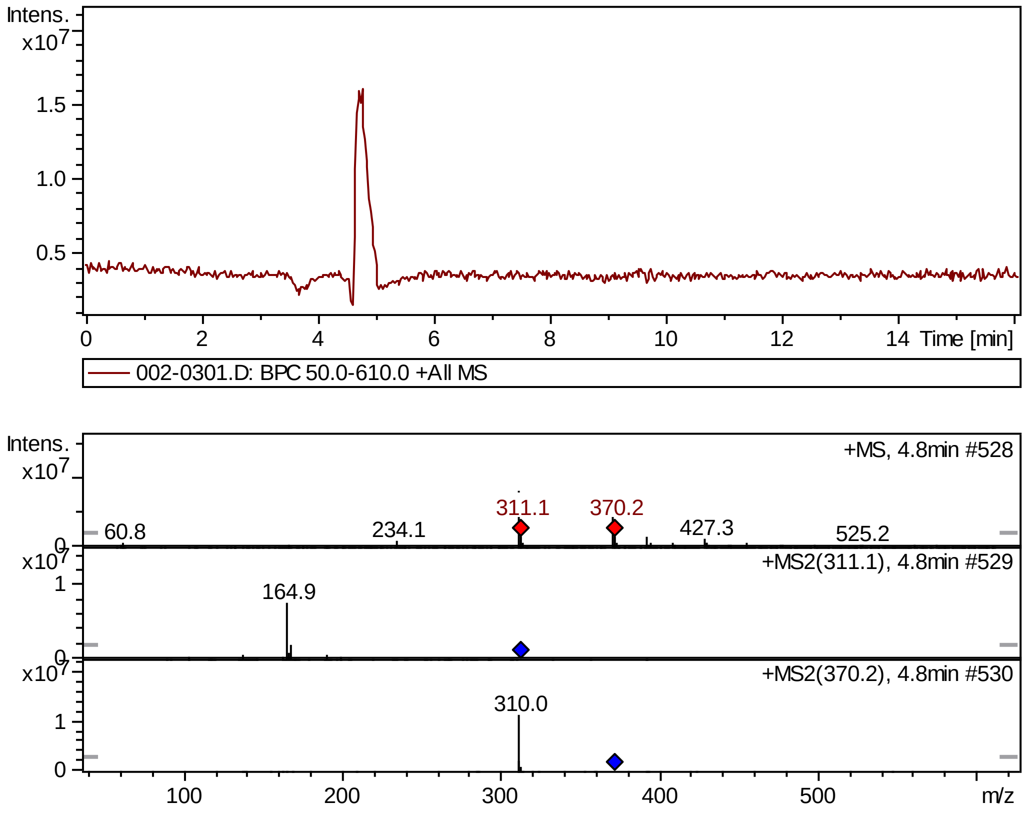


## FT-IR

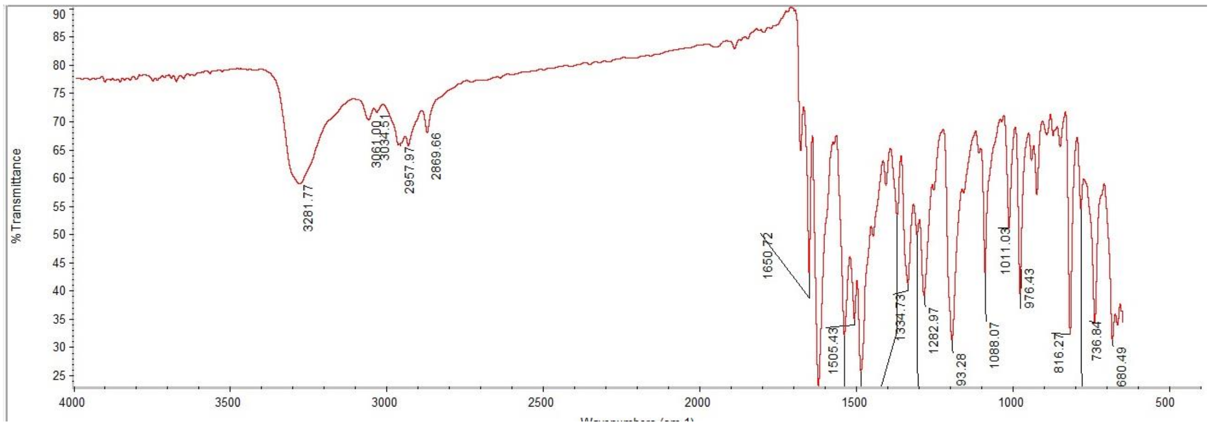

#### $^{1}$H NMR

This report was created by ACD/NMR Processor Academic Edition. For more information go to www.acdlabs.com/nmrproc/

##### RAD-312

05/11/2016 9:57:37 AM
Dr.Mustafa El-Araby Sample : RAD-312 DMSO

| | | | |
|---|---|---|---|
| *Formula* $C_{21}H_{21}ClN_2O_2$ | | *FW* | 368.8566 |

| | | | | | | | |
|---|---|---|---|---|---|---|---|
| *Acquisition Time (sec)* | 1.9268 | *Comment* | Dr.Mustafa El-Araby Sample : RAD-312 DMSO | | | *Date* | 08 Apr 2015 11:25:36 |
| *Date Stamp* | 08 Apr 2015 11:25:36 | | | | | | |
| *File Name* | E:\Mostafa Alaraby Project\Oxazolone NMR\NMR final oxazolone\MUSTAFA RAD-312 08-04-2015\180\fid | | | | | *Frequency (MHz)* | 850.15 |
| *Nucleus* | 1H | *Number of Transients* | 20 | *Origin* | spect | *Original Points Count* | 32768 |
| *Owner* | nmr | *Points Count* | 32768 | *Pulse Sequence* | zg30 | *Receiver Gain* | 10.55 |
| *SW(cyclical) (Hz)* | 17006.80 | *Solvent* | DMSO-d6 | *Spectrum Offset (Hz)* | 5250.0283 | *Spectrum Type* | STANDARD |
| *Sweep Width (Hz)* | 17006.28 | *Temperature (degree C)* | 25.001 | | | | |

$^{1}$H NMR (850 MHz, DMSO-$d_6$) δ 9.71 (br. s., 1H), 8.12 (t, $J$ = 5.71 Hz, 1H), 7.64 (d, $J$ = 8.30 Hz, 2H), 7.46 - 7.58 (m, 4H), 7.38 (t, $J$ = 7.79 Hz, 2H), 7.29 - 7.33 (m, 1H), 7.00 (s, 1H), 6.87 (d, $J$ = 16.09 Hz, 1H), 3.12 (q, $J$ = 6.75 Hz, 2H), 1.49 (sxt, $J$ = 7.27 Hz, 2H), 0.88 (t, $J$ = 7.27 Hz, 3H)

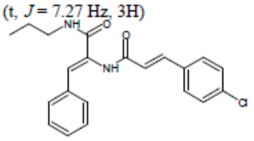


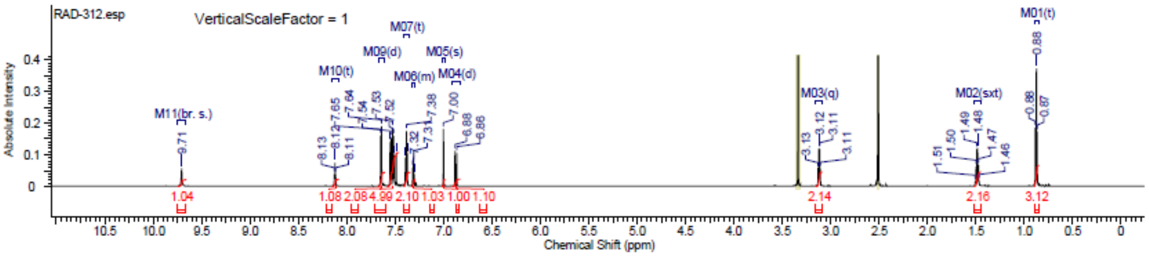


#### $^{13}$C NMR

This report was created by ACD/NMR Processor Academic Edition. For more information go to www.acdlabs.com/nmrproc/

##### RAD-312

20/09/2016 8:33:47 AM
Dr.Mostafa Sample ; RAD-312 DMSO

| | | | |
|---|---|---|---|
| *Formula* $C_{21}H_{21}ClN_2O_2$ | | *FW* | 368.8566 |

| | | | | | | | |
|---|---|---|---|---|---|---|---|
| *Acquisition Time (sec)* | 0.6423 | *Comment* | Dr.Mostafa Sample ; RAD-312 DMSO | | | *Date* | 20 Apr 2016 09:04:48 |
| *Date Stamp* | 20 Apr 2016 09:04:48 | | | | | | |
| *File Name* | E:\Projects\Mostafa Alaraby Project\Oxazolone\Oxazolone NMR\NMR final oxazolone\13 CNMR New 18-4-2016\MUSTAFA RAD-312 19-04-2016\80\fid | | | | | | |
| *Frequency (MHz)* | 213.77 | *Nucleus* | 13C | *Number of Transients* | 2396 | *Origin* | spect |
| *Original Points Count* | 32768 | *Owner* | nmr | *Points Count* | 32768 | *Pulse Sequence* | zgpg30 |
| *Receiver Gain* | 186.93 | *SW(cyclical) (Hz)* | 51020.41 | *Solvent* | DMSO-d6 | *Spectrum Offset (Hz)* | 21301.5977 |
| *Spectrum Type* | STANDARD | *Sweep Width (Hz)* | 51018.85 | *Temperature (degree C)* | 25.000 | | |

$^{13}$C NMR (214 MHz, DMSO-$d_6$) δ 165.1, 164.5, 138.7, 134.3, 133.8, 130.4, 129.5, 129.4, 129.2, 128.7, 128.6, 127.0, 122.4, 41.1, 39.8, 39.7, 39.6, 39.4, 39.3, 39.2, 22.4, 11.5

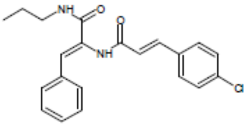


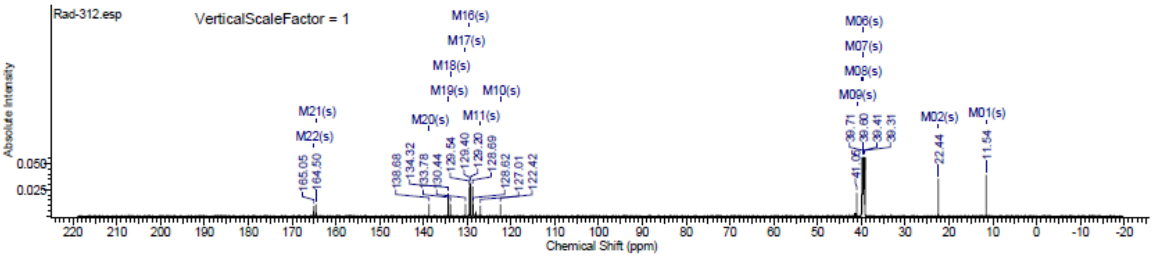

#### Spectra of (1912)

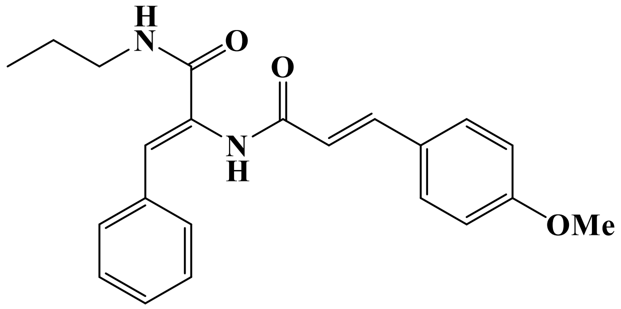


### LC/MS

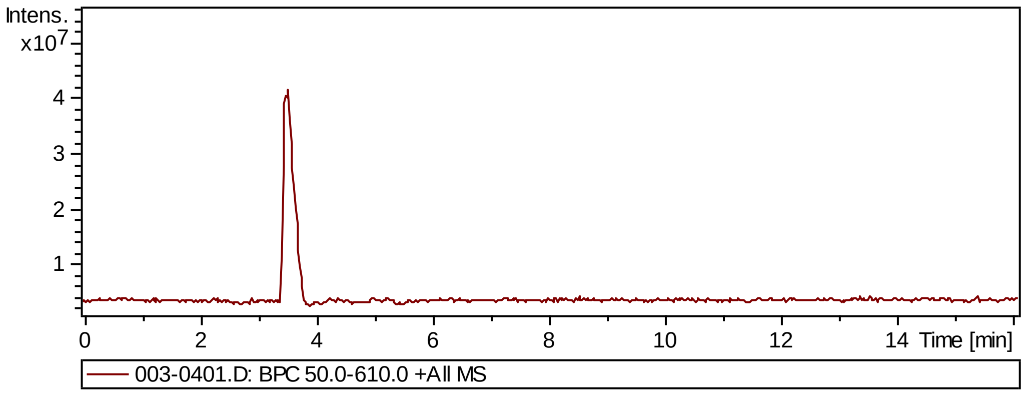


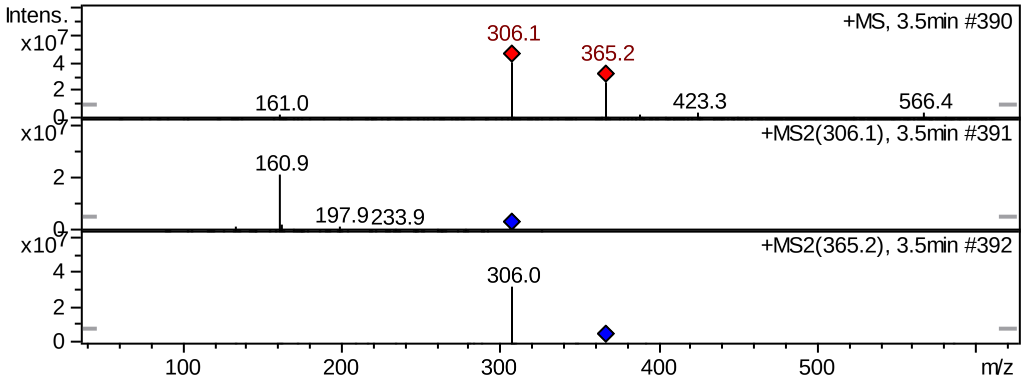


### FT-IR

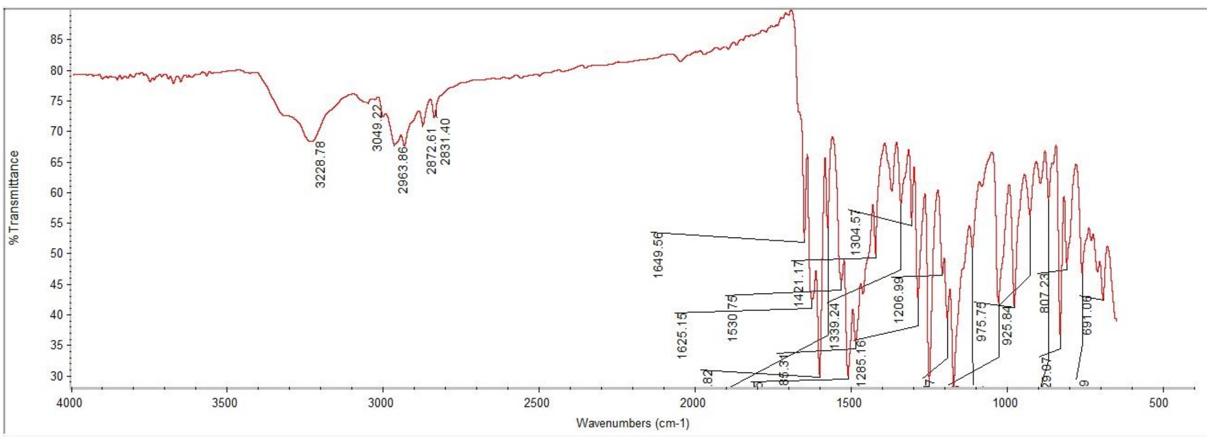

### Supplementary Information

#### $^{1}$H NMR

This report was created by ACD/NMR Processor Academic Edition. For more information go to www.acdlabs.com/nmrproc/

**RAD-313**

05/11/2016 10:04:00 AM
Dr.Mustafa Sample : RAD-313 DMSO PROTON DMSO {D:\Magdy} nmr 32

| Formula | $C_{20}H_{24}N_2O_3$ | FW | 364.4376 |
|---|---|---|---|

| | | | | | | | |
|---|---|---|---|---|---|---|---|
| Acquisition Time (sec) | 2.6564 | Comment | Dr.Mustafa Sample : RAD-313 DMSO PROTON DMSO {D:\Magdy} nmr 32 | | | | |
| Date | 18 Jun 2015 16:19:44 | | | Date Stamp | 18 Jun 2015 16:19:44 | | |
| File Name | E:\Mostafa Alaraby Project\Oxazolone NMR\NMR final oxazolone\MUSTAFA RAD-313 18-06-2015\10\fid | | | | | Frequency (MHz) | 600.15 |
| Nucleus | 1H | Number of Transients | 32 | Origin | spect | Original Points Count | 32768 |
| Owner | nmr | Points Count | 32768 | Pulse Sequence | zg30 | Receiver Gain | 144.00 |
| SW(cyclical) (Hz) | 12335.53 | Solvent | DMSO-d6 | Spectrum Offset (Hz) | 3706.1750 | Spectrum Type | STANDARD |
| Sweep Width (Hz) | 12335.15 | Temperature (degree C) | 25.000 | | | | |

$^{1}$H NMR (600 MHz, DMSO-$d_6$) δ 9.62 (s, 1H), 8.12 (t, $J$ = 5.65 Hz, 1H), 7.51 - 7.62 (m, 4H), 7.45 (d, $J$ = 15.43 Hz, 1H), 7.38 (t, $J$ = 7.72 Hz, 2H), 7.28 - 7.33 (m, 1H), 7.02 (d, $J$ = 8.66 Hz, 2H), 6.97 (s, 1H), 6.72 (d, $J$ = 15.81 Hz, 1H), 3.81 (s, 3H), 3.11 (q, $J$ = 6.40 Hz, 2H), 1.48 (sxt, $J$ = 7.30 Hz, 2H), 0.87 (t, $J$ = 7.34 Hz, 3H)

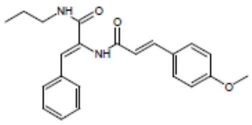


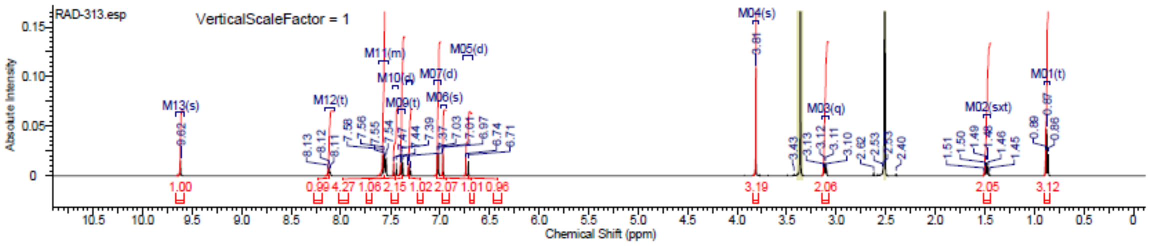


#### $^{13}$C NMR

This report was created by ACD/NMR Processor Academic Edition. For more information go to www.acdlabs.com/nmrproc/

**RAD-313**

20/09/2016 8:34:14 AM
Dr.Mostafa Sample ; RAD-313 DMSO

| Formula | $C_{20}H_{24}N_2O_3$ | FW | 364.4375 |
|---|---|---|---|

| | | | | | | | |
|---|---|---|---|---|---|---|---|
| Acquisition Time (sec) | 0.6423 | Comment | Dr.Mostafa Sample ; RAD-313 DMSO | | | Date | 19 Apr 2016 20:25:20 |
| Date Stamp | 19 Apr 2016 20:25:20 | | | | | | |
| File Name | E:\Projects\Mostafa Alaraby Project\Oxazolone\Oxazolone NMR\NMR final oxazolone\13 CNMR New 18-4-2016\MUSTAFA RAD-313 19-04-2016\30\fid | | | | | | |
| Frequency (MHz) | 213.77 | Nucleus | 13C | Number of Transients | 3500 | Origin | spect |
| Original Points Count | 32768 | Owner | nmr | Points Count | 32768 | Pulse Sequence | zgpg30 |
| Receiver Gain | 186.93 | SW(cyclical) (Hz) | 51020.41 | Solvent | DMSO-d6 | Spectrum Offset (Hz) | 21289.1406 |
| Spectrum Type | STANDARD | Sweep Width (Hz) | 51018.85 | Temperature (degree C) | 25.000 | | |

$^{13}$C NMR (214 MHz, DMSO-$d_6$) δ 165.1, 164.9, 160.6, 139.7, 134.4, 130.7, 129.4, 129.3, 128.6, 128.5, 127.3, 126.6, 119.0, 114.5, 55.4, 41.0, 39.8, 39.7, 39.6, 39.4, 39.3, 39.2, 22.4, 11.5

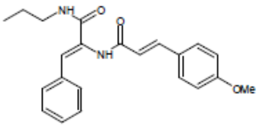


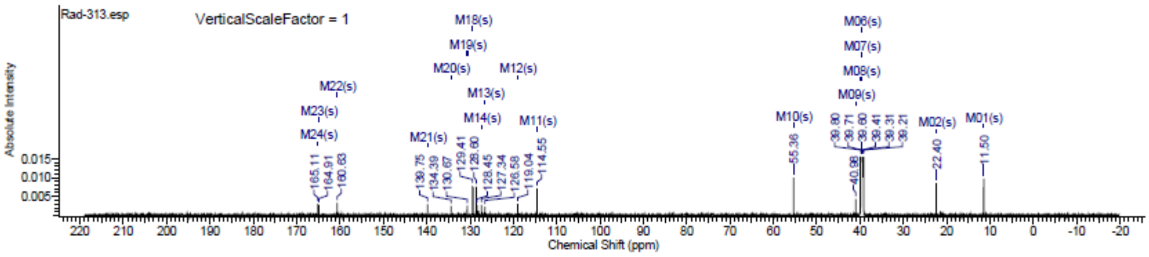

### Spectra of (2012)

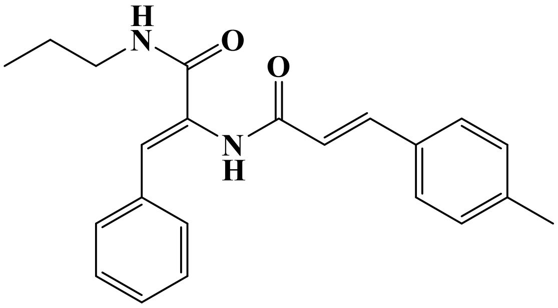

## LC/MS

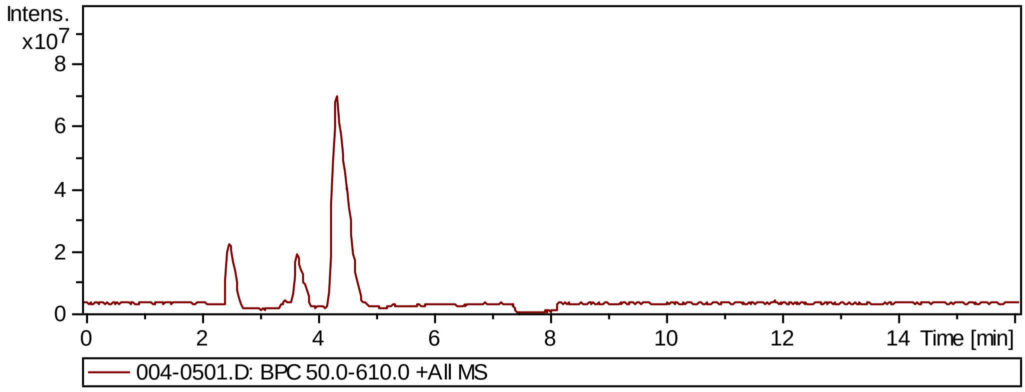


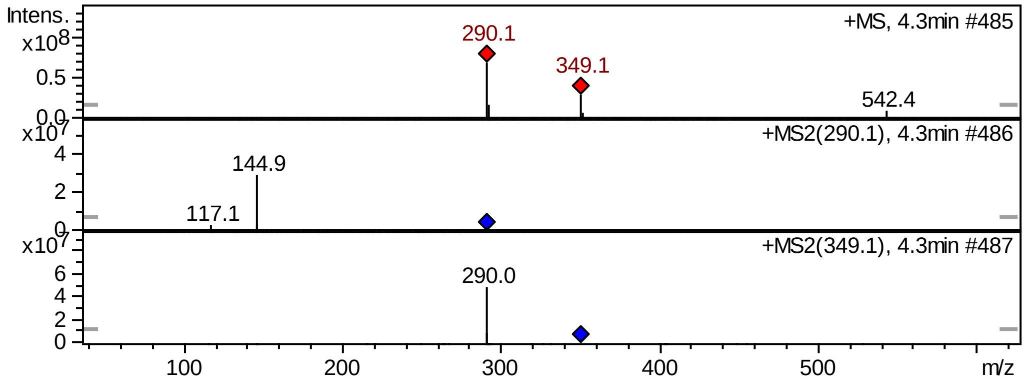

### Supplementary Information

#### [1]H NMR

This report was created by ACD/NMR Processor Academic Edition. For more information go to www.acdlabs.com/nmrproc/

**RAD-314**

02/10/2016 6:30:24 AM
Dr.Mustafa El-Araby Sample : RAD-314 DMSO

| | |
|---|---|
| Formula $C_{22}H_{24}N_2O_2$ | FW 348.4382 |

| | | | | | | | |
|---|---|---|---|---|---|---|---|
| Acquisition Time (sec) | 1.9268 | Comment | Dr.Mustafa El-Araby Sample : RAD-314 DMSO | | | Date | 08 Apr 2015 10:23:44 |
| Date Stamp | 08 Apr 2015 10:23:44 | | | | | | |
| File Name | E:\Mostafa Alaraby Project\Oxazolone NMR\NMR final oxazolone\MUSTAFA RAD-314 08-04-2015\40\fid | | | | | Frequency (MHz) | 850.15 |
| Nucleus | 1H | Number of Transients | 20 | Origin | spect | Original Points Count | 32768 |
| Owner | nmr | Points Count | 32768 | Pulse Sequence | zg30 | Receiver Gain | 9.04 |
| SW(cyclical) (Hz) | 17006.80 | Solvent | DMSO-d6 | Spectrum Offset (Hz) | 5250.0283 | Spectrum Type | STANDARD |
| Sweep Width (Hz) | 17006.28 | Temperature (degree C) | 25.000 | | | | |

[1]H NMR (850 MHz, DMSO-$d_6$) δ 9.65 (br. s., 1H), 8.11 (t, $J$ = 5.45 Hz, 1H), 7.49 - 7.57 (m, 3H), 7.46 (d, $J$ = 15.57 Hz, 1H), 7.34 - 7.43 (m, 3H), 7.28 - 7.34 (m, 2H), 7.26 (d, $J$ = 7.78 Hz, 1H), 6.95 - 7.03 (m, 1H), 6.82 (d, $J$ = 15.57 Hz, 1H), 3.11 (td, $J$ = 6.68, 13.10 Hz, 2H), 1.48 (qd, $J$ = 7.07, 14.08 Hz, 2H), 0.84 - 0.90 (m, 3H)

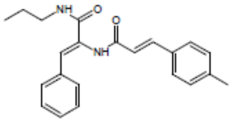

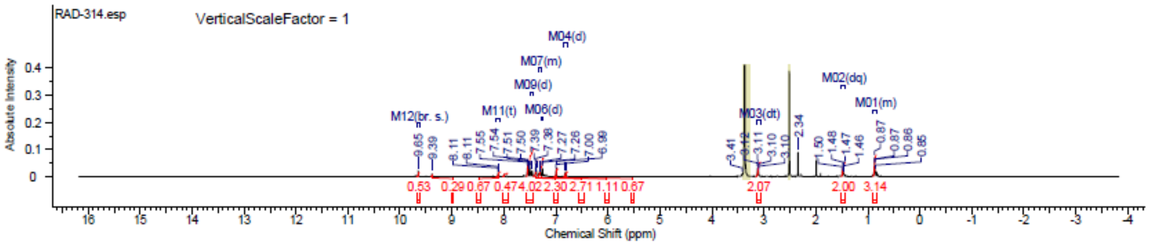


#### [13]C NMR

This report was created by ACD/NMR Processor Academic Edition. For more information go to www.acdlabs.com/nmrproc/

**RAD-314**

20/09/2016 8:34:47 AM
Dr.Mostafa Sample ; RAD-314 DMSO

| | |
|---|---|
| Formula $C_{22}H_{24}N_2O_2$ | FW 348.4382 |

| | | | | | | | |
|---|---|---|---|---|---|---|---|
| Acquisition Time (sec) | 0.6423 | Comment | Dr.Mostafa Sample ; RAD-314 DMSO | | | Date | 19 Apr 2016 23:07:28 |
| Date Stamp | 19 Apr 2016 23:07:28 | | | | | | |
| File Name | E:\Projects\Mostafa Alaraby Project\Oxazolone\Oxazolone NMR\NMR final oxazolone\13 CNMR New 18-4-2016\MUSTAFA RAD-314 19-04-2016\40\fid | | | | | | |
| Frequency (MHz) | 213.77 | Nucleus | 13C | Number of Transients | 3500 | Origin | spect |
| Original Points Count | 32768 | Owner | nmr | Points Count | 32768 | Pulse Sequence | zgpg30 |
| Receiver Gain | 186.93 | SW(cyclical) (Hz) | 51020.41 | Solvent | DMSO-d6 | Spectrum Offset (Hz) | 21309.3809 |
| Spectrum Type | STANDARD | Sweep Width (Hz) | 51018.85 | Temperature (degree C) | 25.006 | | |

[13]C NMR (214 MHz, DMSO-$d_6$) δ 165.2, 164.9, 140.1, 139.8, 134.4, 132.1, 130.6, 129.8, 129.4, 128.7, 127.9, 127.5, 126.9, 120.6, 41.1, 39.8, 39.7, 39.6, 39.4, 39.3, 39.2, 22.5, 21.1, 11.6

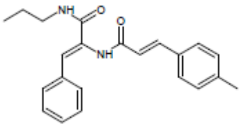

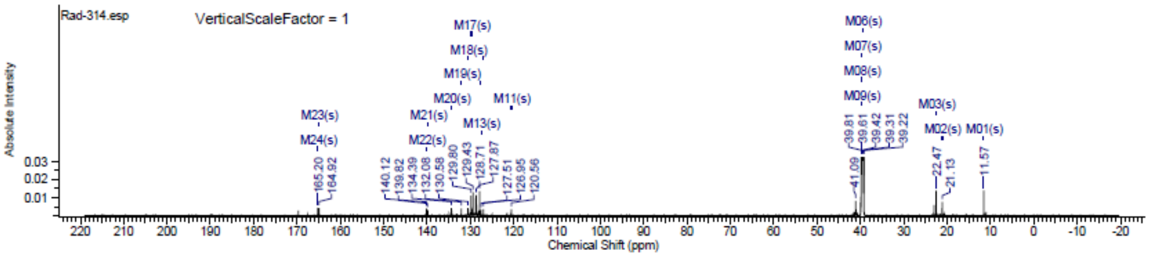

### Spectra of (2112)

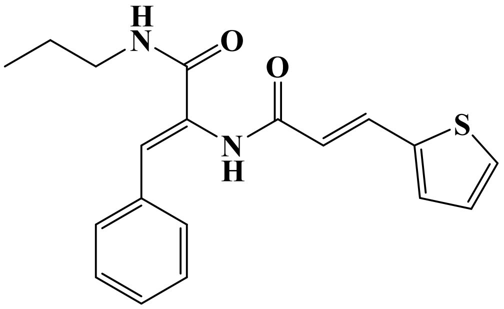

## LC/MS

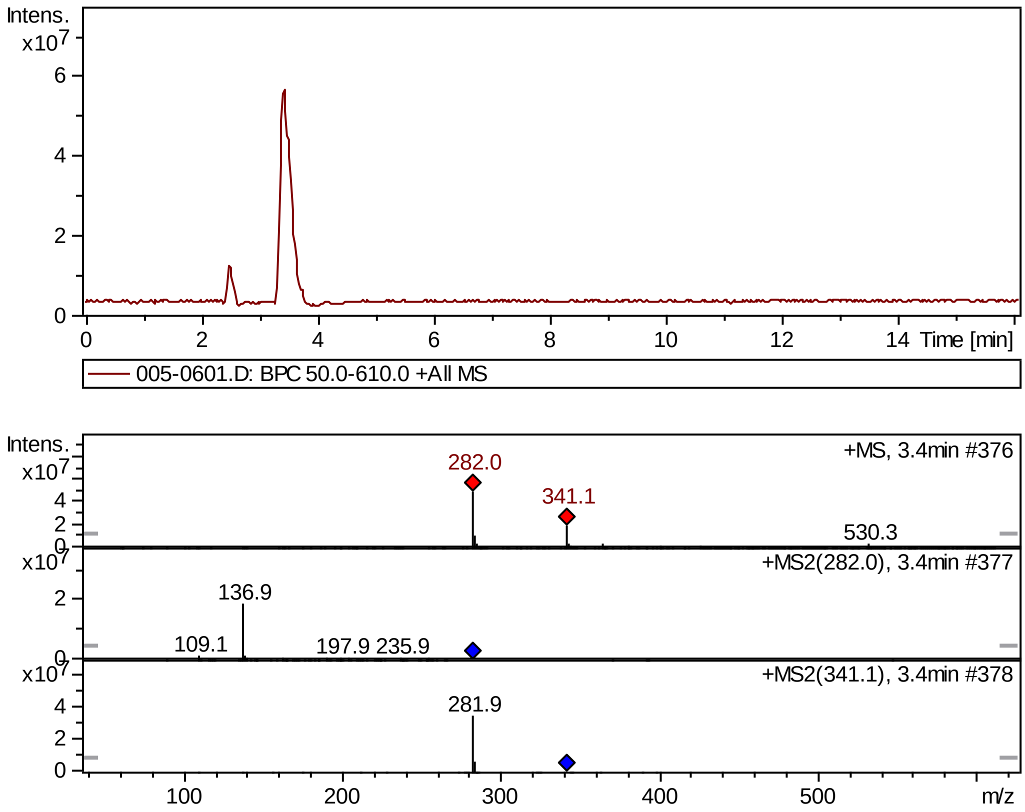


## FT-IR

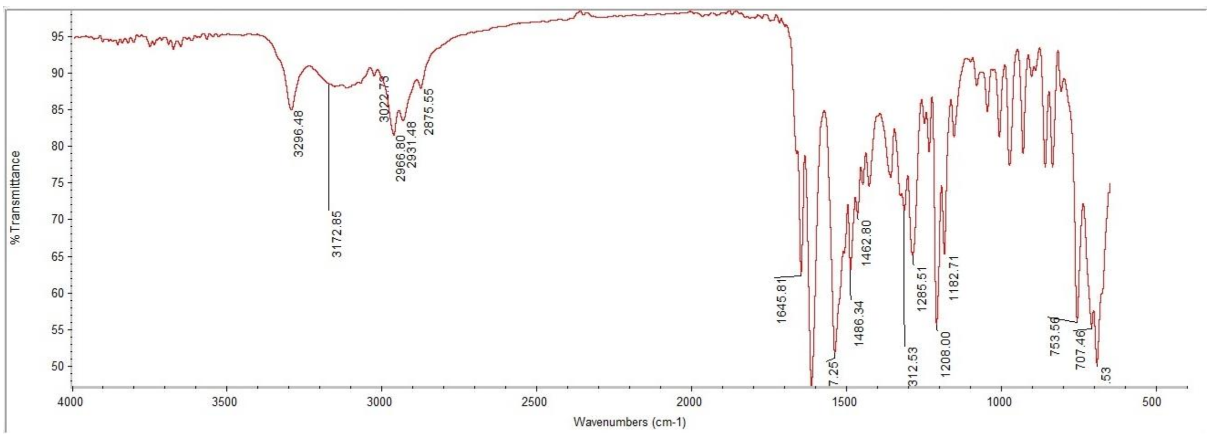

#### $^1$H NMR

This report was created by ACD/NMR Processor Academic Edition. For more information go to www.acdlabs.com/nmrproc/

**RAD-315**

08/11/2016 7:51:58 AM
Dr.Mustafa El-Araby Sample : RAD-315 DMSO

| Formula $C_{19}H_{20}N_2O_2S$ | FW 340.4393 |
|---|---|

| | | | | | | | |
|---|---|---|---|---|---|---|---|
| *Acquisition Time (sec)* | 1.9268 | *Comment* | Dr.Mustafa El-Araby Sample : RAD-315 DMSO | | | *Date* | 08 Apr 2015 10:53:36 |
| *Date Stamp* | 08 Apr 2015 10:53:36 | | | | | | |
| *File Name* | E:\Mostafa Alaraby Project\Oxazolone NMR\NMR final oxazolone\MUSTAFA RAD-315 08-04-2015\110\fid | | | | | *Frequency (MHz)* | 850.15 |
| *Nucleus* | 1H | *Number of Transients* | 20 | *Origin* | spect | *Original Points Count* | 32768 |
| *Owner* | nmr | *Points Count* | 32768 | *Pulse Sequence* | zg30 | *Receiver Gain* | 10.55 |
| *SW(cyclical) (Hz)* | 17006.80 | *Solvent* | DMSO-d6 | *Spectrum Offset (Hz)* | 5250.0283 | *Spectrum Type* | STANDARD |
| *Sweep Width (Hz)* | 17006.28 | *Temperature (degree C)* | 25.001 | | | | |

$^1$H NMR (850 MHz, DMSO-$d_6$) δ 9.65 (s, 1H), 8.10 (t, $J$ = 5.71 Hz, 1H), 7.61 - 7.68 (m, 2H), 7.53 (d, $J$ = 7.79 Hz, 2H), 7.43 (d, $J$ = 3.11 Hz, 1H), 7.39 (t, $J$ = 7.79 Hz, 2H), 7.28 - 7.33 (m, 1H), 7.14 (dd, $J$ = 3.37, 4.93 Hz, 1H), 6.99 (s, 1H), 6.62 (d, $J$ = 15.57 Hz, 1H), 3.11 (q, $J$ = 6.75 Hz, 2H), 1.48 (sxt, $J$ = 7.27 Hz, 2H), 0.87 (t, $J$ = 7.53 Hz, 3H)

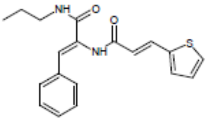


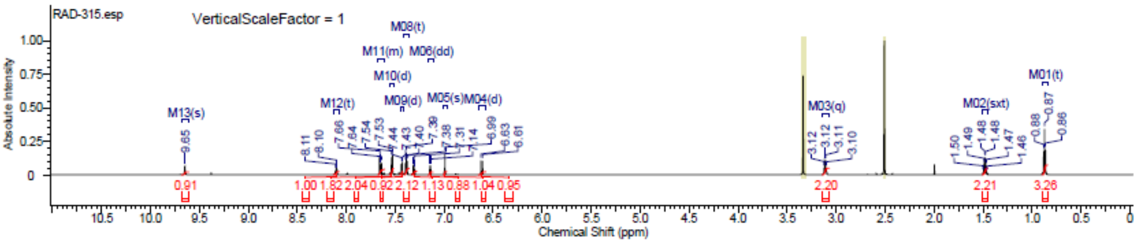


#### $^{13}$C NMR

This report was created by ACD/NMR Processor Academic Edition. For more information go to www.acdlabs.com/nmrproc/

**RAD-315**

20/09/2016 8:35:12 AM
Dr.Mostafa Sample ; RAD-315 DMSO

| Formula $C_{19}H_{20}N_2O_2S$ | FW 340.4393 |
|---|---|

| | | | | | | | |
|---|---|---|---|---|---|---|---|
| *Acquisition Time (sec)* | 0.6423 | *Comment* | Dr.Mostafa Sample ; RAD-315 DMSO | | | *Date* | 20 Apr 2016 01:49:36 |
| *Date Stamp* | 20 Apr 2016 01:49:36 | | | | | | |
| *File Name* | E:\Projects\Mostafa Alaraby Project\Oxazolone\Oxazolone NMR\NMR final oxazolone\13 CNMR New 18-4-2016\MUSTAFA RAD-315 19-04-2016\50\fid | | | | | | |
| *Frequency (MHz)* | 213.77 | *Nucleus* | 13C | *Number of Transients* | 3500 | *Origin* | spect |
| *Original Points Count* | 32768 | *Owner* | nmr | *Points Count* | 32768 | *Pulse Sequence* | zgpg30 |
| *Receiver Gain* | 186.93 | *SW(cyclical) (Hz)* | 51020.41 | *Solvent* | DMSO-d6 | *Spectrum Offset (Hz)* | 21293.8125 |
| *Spectrum Type* | STANDARD | *Sweep Width (Hz)* | 51018.85 | *Temperature (degree C)* | 25.000 | | |

$^{13}$C NMR (214 MHz, DMSO-$d_6$) δ 165.0, 164.4, 139.8, 134.3, 133.0, 131.3, 130.5, 129.3, 128.6, 128.6, 128.5, 128.5, 126.9, 120.3, 41.0, 39.8, 39.7, 39.6, 39.4, 39.3, 39.2, 22.4, 11.5

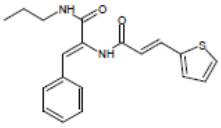


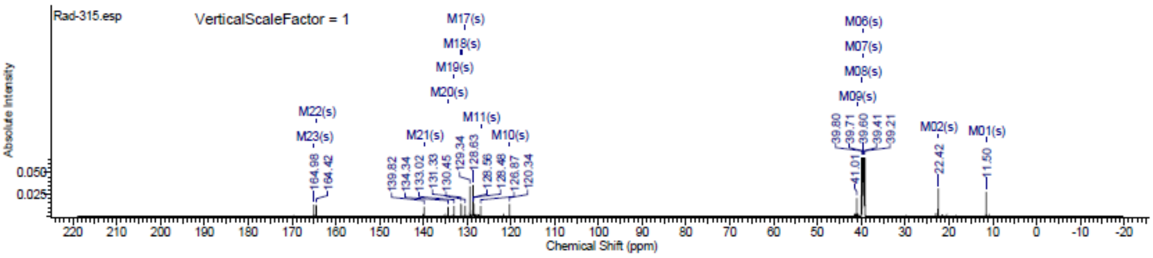

### Spectra of (2212)

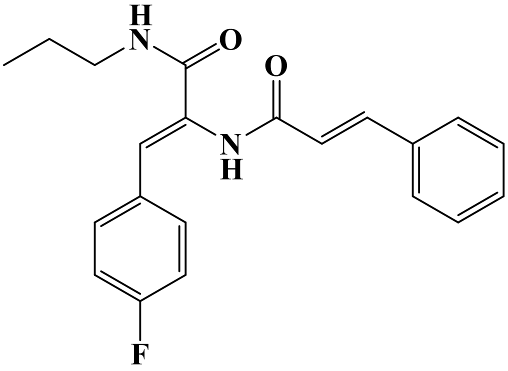

## LC/MS

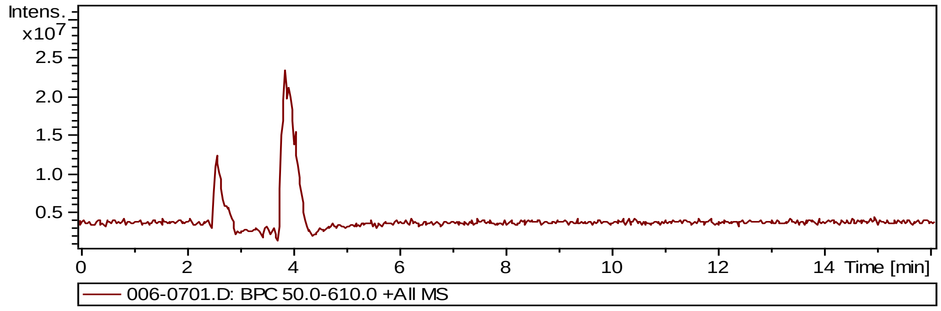


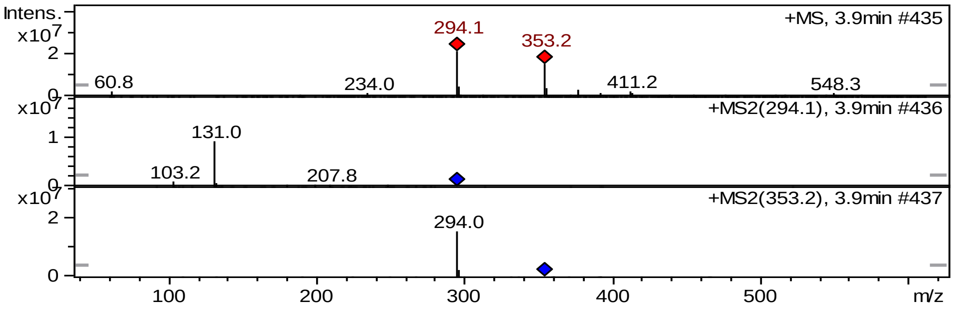


#### $^1$H NMR

**2212**

10/4/2018 11:13:25 PM
Dr.Mostafa Sample : RAD-321 DMSO

| | | | | | | | |
|---|---|---|---|---|---|---|---|
| Acquisition Time (sec) | 1.9268 | Comment | Dr.Mostafa Sample : RAD-321 DMSO | | | Date | 14 Feb 2017 12:29:36 |
| Date Stamp | 14 Feb 2017 12:29:36 | | | | | | |
| File Name | E:\GoogleDrive 2018\Shuttle\MOS-Final-Sep2018\Raw Data for Reviewers\MOSTAFA RAD-321 14-02-2017\10\fid | | | | | | |
| Frequency (MHz) | 850.15 | Nucleus | 1H | Number of Transients | 64 | Origin | spect |
| Original Points Count | 32768 | Owner | nmr | Points Count | 32768 | Pulse Sequence | zg30 |
| Receiver Gain | 9.04 | SW(cyclical) (Hz) | 17006.80 | Solvent | DMSO-d6 | Spectrum Offset (Hz) | 5250.0283 |
| Spectrum Type | STANDARD | Sweep Width (Hz) | 17006.28 | Temperature (degree C) | 25.000 | | |

$^1$H NMR (DMSO-$d_6$) δ ppm 9.68 (s, 1 H), 8.12 (s, 1 H), 7.56 - 7.67 (m, 4 H), 7.50 (d, $J$=16.09 Hz, 1 H), 7.44 - 7.47 (m, 2 H), 7.42 (d, $J$=7.78 Hz, 1 H), 7.23 (t, $J$=8.82 Hz, 2 H), 7.01 (s, 1 H), 6.86 (d, $J$=16.09 Hz, 1 H), 3.12 (q, $J$=6.75 Hz, 2 H), 1.45 - 1.52 (m, 2 H), 0.87 (t, $J$=7.53 Hz, 3 H)

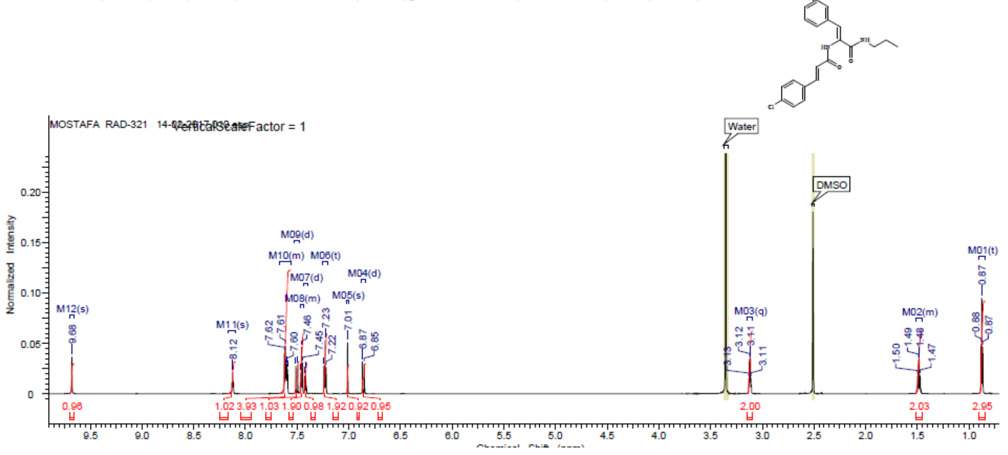

### Spectra of (2312)

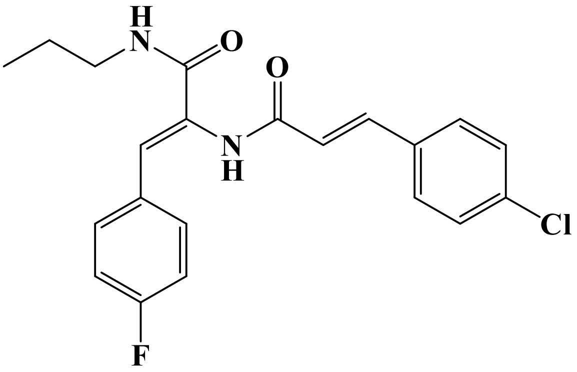

## LC/MS

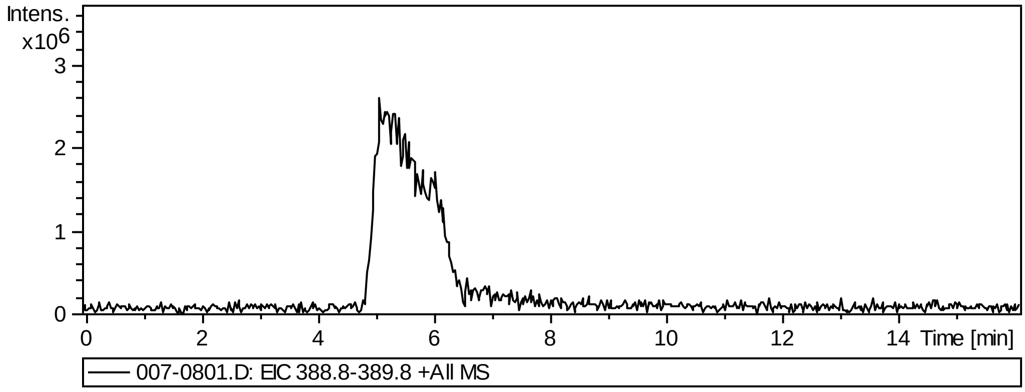


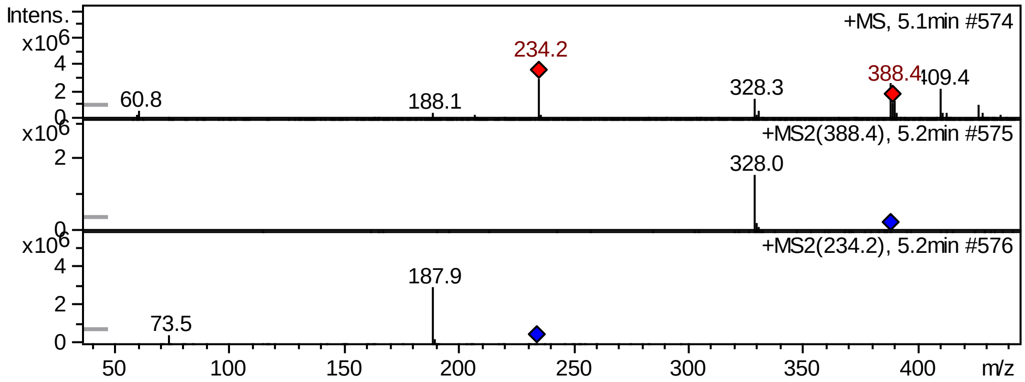


## FT-IR

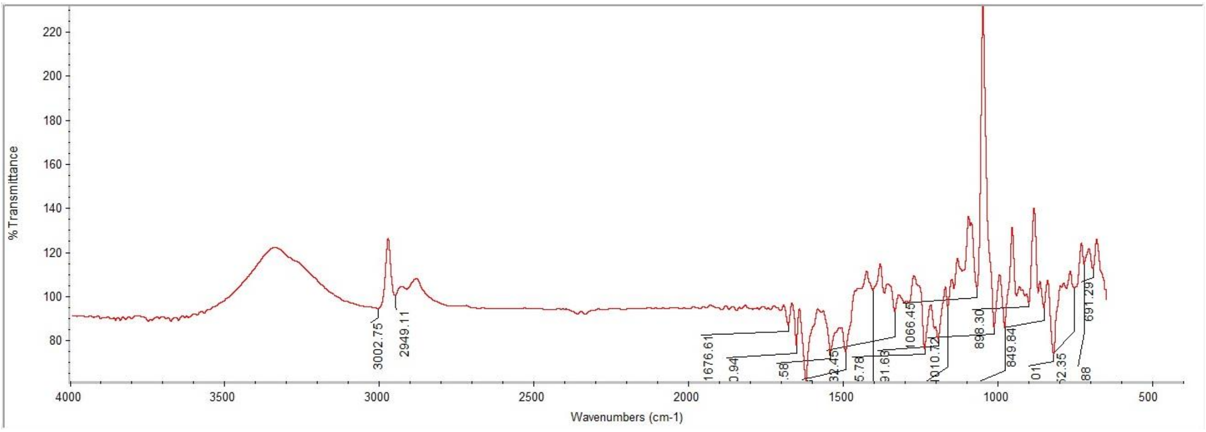

#### [1]H NMR

This report was created by ACD/NMR Processor Academic Edition. For more information go to www.acdlabs.com/nmrproc/

**RAD-322**

08/11/2016 7:58:08 AM
Dr.Mustafa El-Araby Sample : RAD-322 DMSO

| Formula | $C_{21}H_{20}ClFN_2O_2$ | FW | 386.8471 |
|---|---|---|---|

| | | | | | | | |
|---|---|---|---|---|---|---|---|
| Acquisition Time (sec) | 1.9268 | Comment | Dr.Mustafa El-Araby Sample : RAD-322 DMSO | | | Date | 08 Apr 2015 10:45:04 |
| Date Stamp | 08 Apr 2015 10:45:04 | | | | | | |
| File Name | E:\Mostafa Alaraby Project\Oxazolone NMR\NMR final oxazolone\MUSTAFA RAD-322 08-04-2015\90\fid | | | | | Frequency (MHz) | 850.15 |
| Nucleus | 1H | Number of Transients | 20 | Origin | spect | Original Points Count | 32768 |
| Owner | nmr | Points Count | 32768 | Pulse Sequence | zg30 | Receiver Gain | 10.55 |
| SW(cyclical) (Hz) | 17006.80 | Solvent | DMSO-d6 | Spectrum Offset (Hz) | 5250.0283 | Spectrum Type | STANDARD |
| Sweep Width (Hz) | 17006.28 | Temperature (degree C) | 25.000 | | | | |

$^1$H NMR (850 MHz, DMSO-$d_6$) δ 9.69 (s, 1H), 8.12 (t, $J$ = 5.71 Hz, 1H), 7.64 (d, $J$ = 8.30 Hz, 2H), 7.60 (dd, $J$ = 5.71, 8.82 Hz, 2H), 7.46 - 7.55 (m, 3H), 7.21 - 7.25 (m, 2H), 7.01 (s, 1H), 6.85 (d, $J$ = 16.09 Hz, 1H), 3.11 (q, $J$ = 6.40 Hz, 2H), 1.48 (sxt, $J$ = 7.27 Hz, 2H), 0.87 (t, $J$ = 7.27 Hz, 3H)

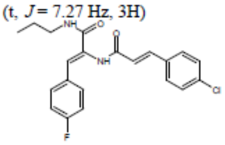


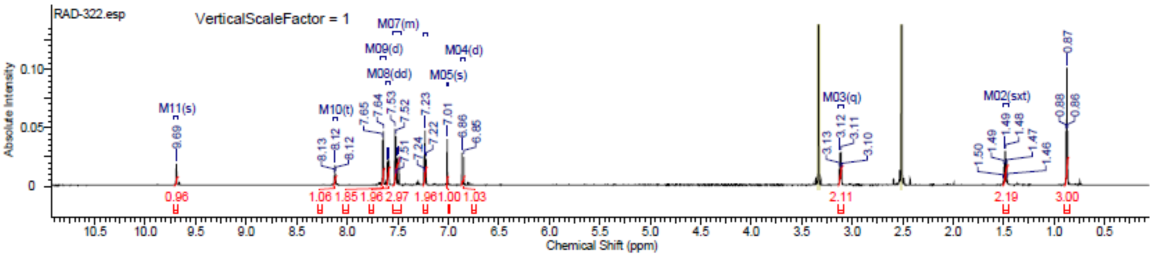


#### [13]C NMR

This report was created by ACD/NMR Processor Academic Edition. For more information go to www.acdlabs.com/nmrproc/

**RAD-322**

20/09/2016 8:35:42 AM
Dr.Mostafa Sample : RAD-322 DMSO

| Formula | $C_{21}H_{20}ClFN_2O_2$ | FW | 386.8471 |
|---|---|---|---|

| | | | | | | | |
|---|---|---|---|---|---|---|---|
| Acquisition Time (sec) | 0.6423 | Comment | Dr.Mostafa Sample : RAD-322 DMSO | | | Date | 19 Apr 2016 12:53:04 |
| Date Stamp | 19 Apr 2016 12:53:04 | | | | | | |
| File Name | E:\Projects\Mostafa Alaraby Project\Oxazolone\Oxazolone NMR\NMR final oxazolone\13 CNMR New 18-4-2016\MUSTAFA RAD-322 18-04-2016\150\fid | | | | | | |
| Frequency (MHz) | 213.77 | Nucleus | 13C | Number of Transients | 2384 | Origin | spect |
| Original Points Count | 32768 | Owner | nmr | Points Count | 32768 | Pulse Sequence | zgpg30 |
| Receiver Gain | 186.93 | SW(cyclical) (Hz) | 51020.41 | Solvent | DMSO-d6 | Spectrum Offset (Hz) | 21292.2539 |
| Spectrum Type | STANDARD | Sweep Width (Hz) | 51018.85 | Temperature (degree C) | 25.000 | | |

$^{13}$C NMR (214 MHz, DMSO-$d_6$) δ 164.9, 164.4, 138.7, 134.3, 133.7, 131.5, 130.9, 130.1, 129.5, 129.2, 125.9, 122.4, 115.6, 115.5, 41.0, 39.8, 39.7, 39.6, 39.4, 39.3, 39.2, 22.4, 11.5

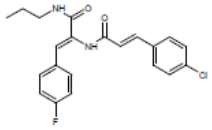


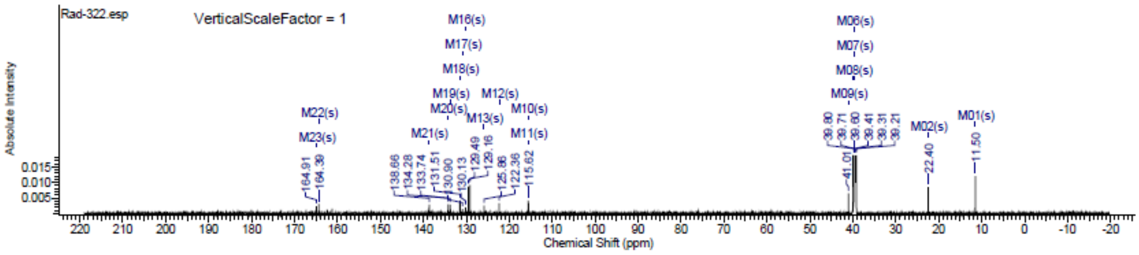

### Spectra of (2412)

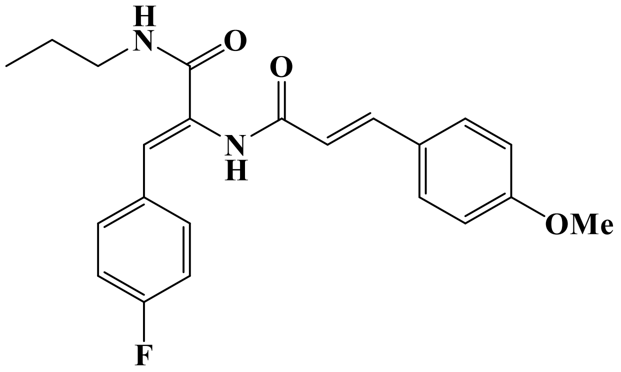


## LC/MS

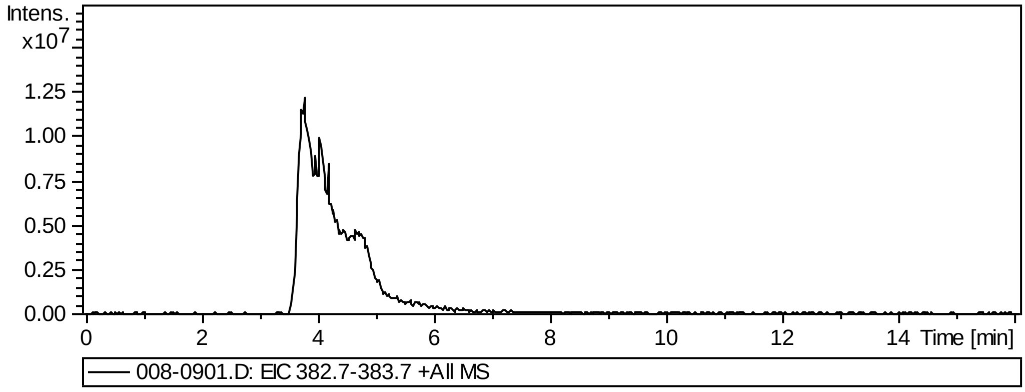


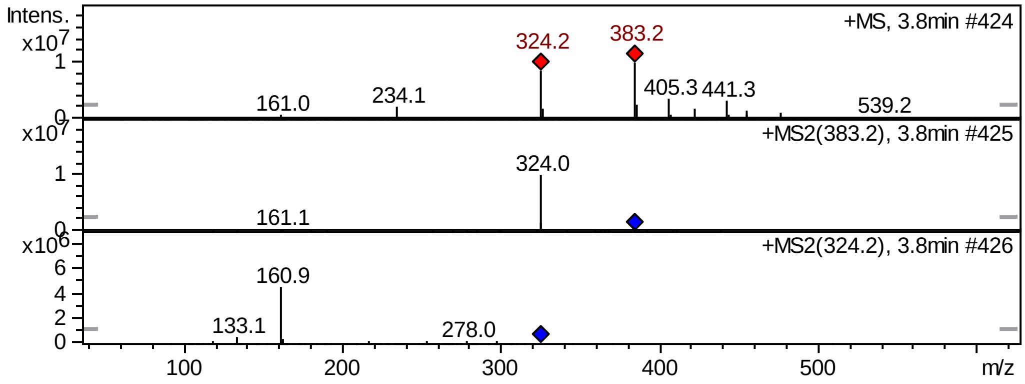


## FT-IR

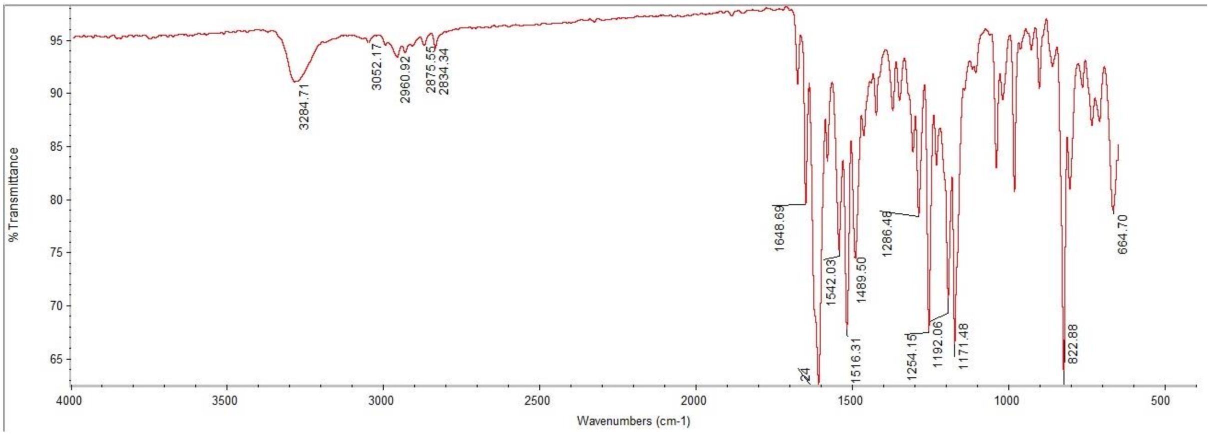

#### $^{1}$H NMR

This report was created by ACD/NMR Processor Academic Edition. For more information go to www.acdlabs.com/nmrproc/

**RAD-323**

08/11/2016 8:02:57 AM
Dr.Mustafa Sample : RAD-323 DMSO PROTON DMSO {D:\Magdy} nmr 26

| Formula $C_{22}H_{23}FN_2O_3$ | FW 382.4280 |
|---|---|

| | | | | | | | |
|---|---|---|---|---|---|---|---|
| *Acquisition Time (sec)* | 2.6564 | *Comment* | Dr.Mustafa Sample : RAD-323 DMSO PROTON DMSO {D:\Magdy} nmr 26 | | | | |
| *Date* | 18 Jun 2015 15:47:44 | | | *Date Stamp* | 18 Jun 2015 15:47:44 | | |
| *File Name* | E:\Mostafa Alaraby Project\Oxazolone NMR\NMR final oxazolone\MUSTAFA RAD-323 18-06-2015\10\fid | | | | | *Frequency (MHz)* | 600.15 |
| *Nucleus* | 1H | *Number of Transients* | 32 | *Origin* | spect | *Original Points Count* | 32768 |
| *Owner* | nmr | *Points Count* | 32768 | *Pulse Sequence* | zg30 | *Receiver Gain* | 128.00 |
| *SW(cyclical) (Hz)* | 12335.53 | *Solvent* | DMSO-d6 | *Spectrum Offset (Hz)* | 3706.1750 | *Spectrum Type* | STANDARD |
| *Sweep Width (Hz)* | 12335.15 | *Temperature (degree C)* | 25.000 | | | | |

$^{1}$H NMR (600 MHz, DMSO-$d_6$) δ 9.61 (s, 1H), 8.12 (t, $J$ = 5.65 Hz, 1H), 7.56 - 7.61 (m, 4H), 7.45 (d, $J$ = 15.81 Hz, 1H), 7.20 - 7.26 (m, 2H), 7.02 (d, $J$ = 8.66 Hz, 2H), 6.98 (s, 1H), 6.71 (d, $J$ = 15.81 Hz, 1H), 3.81 (s, 3H), 3.11 (q, $J$ = 6.65 Hz, 2H), 1.48 (sxt, $J$ = 7.30 Hz, 2H), 0.87 (t, $J$ = 7.53 Hz, 3H)

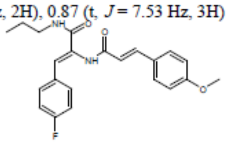


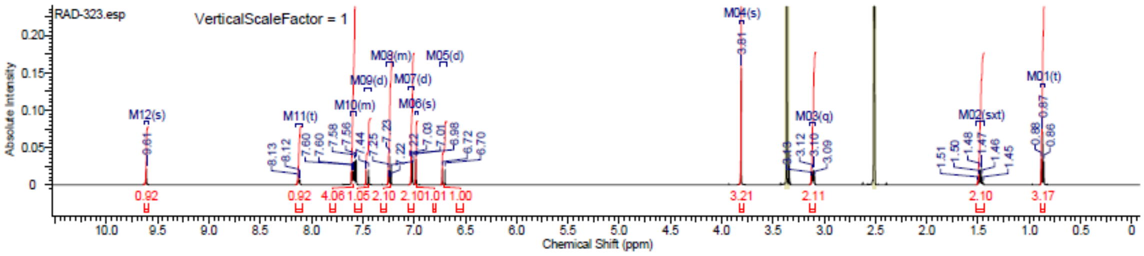


#### $^{13}$C NMR

This report was created by ACD/NMR Processor Academic Edition. For more information go to www.acdlabs.com/nmrproc/

**RAD-323**

20/09/2016 8:36:09 AM
Dr.Mostafa Sample : RAD-323 DMSO

| Formula $C_{22}H_{23}FN_2O_3$ | FW 382.4280 |
|---|---|

| | | | | | | | |
|---|---|---|---|---|---|---|---|
| *Acquisition Time (sec)* | 0.6423 | *Comment* | Dr.Mostafa Sample : RAD-323 DMSO | | | *Date* | 19 Apr 2016 10:15:12 |
| *Date Stamp* | 19 Apr 2016 10:15:12 | | | | | | |
| *File Name* | E:\Projects\Mostafa Alaraby Project\Oxazolone\Oxazolone NMR\NMR final oxazolone\13 CNMR New 18-4-2016\MUSTAFA RAD-323 18-04-2016\110\fid | | | | | | |
| *Frequency (MHz)* | 213.77 | *Nucleus* | 13C | *Number of Transients* | 604 | *Origin* | spect |
| *Original Points Count* | 32768 | *Owner* | nmr | *Points Count* | 32768 | *Pulse Sequence* | zgpg30 |
| *Receiver Gain* | 186.93 | *SW(cyclical) (Hz)* | 51020.41 | *Solvent* | DMSO-d6 | *Spectrum Offset (Hz)* | 21295.3691 |
| *Spectrum Type* | STANDARD | *Sweep Width (Hz)* | 51018.85 | *Temperature (degree C)* | 24.999 | | |

$^{13}$C NMR (214 MHz, DMSO-$d_6$) δ 165.1, 164.9, 160.7, 139.9, 131.5, 131.0, 130.4, 129.5, 127.3, 125.6, 119.0, 115.6, 115.5, 114.6, 55.4, 41.0, 39.8, 39.7, 39.6, 39.4, 39.3, 39.2, 22.4, 11.5

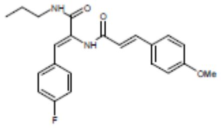


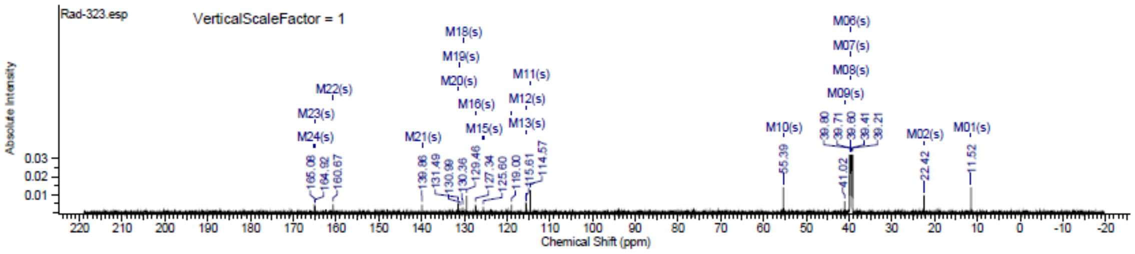

### Spectra of (2512)

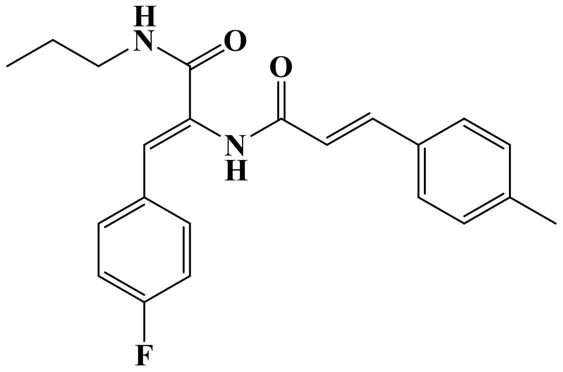


## LC/MS

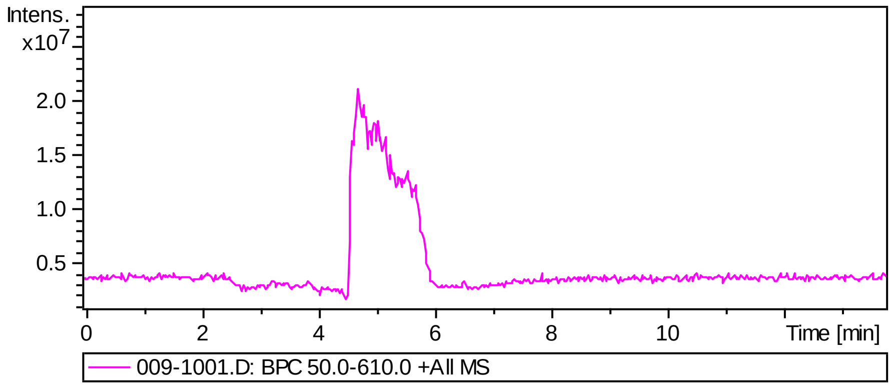


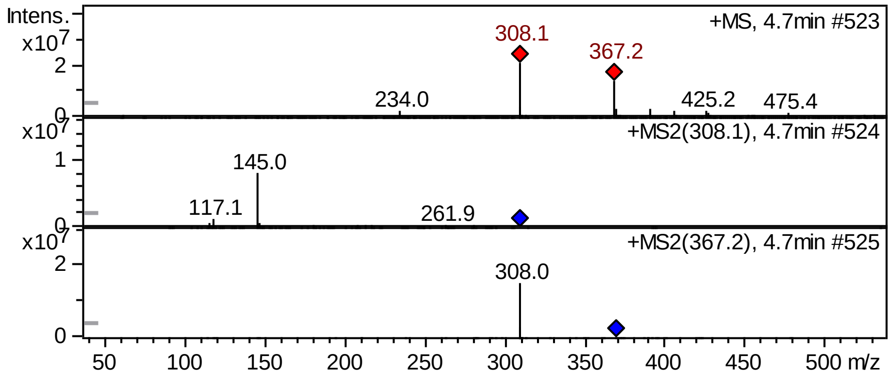


## FT-IR

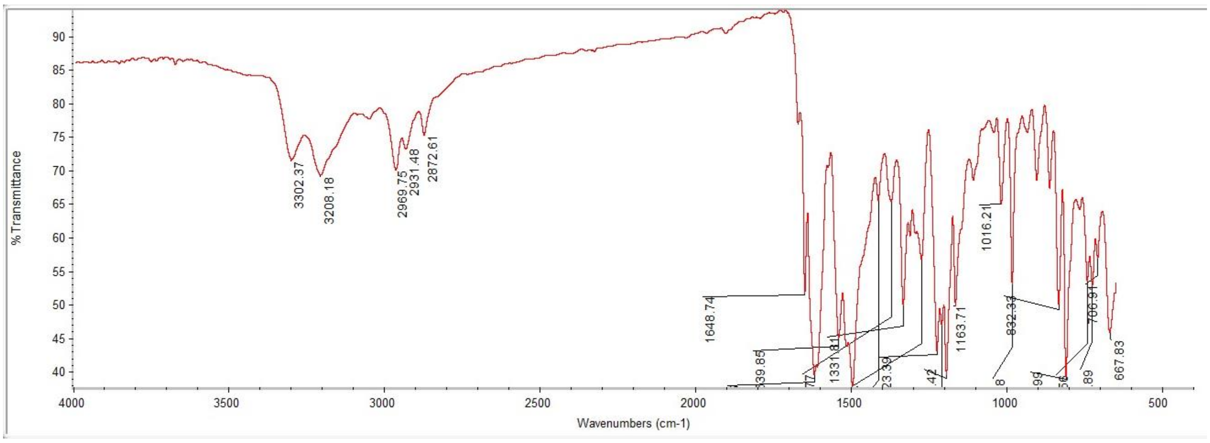

#### $^{1}$H NMR

This report was created by ACD/NMR Processor Academic Edition. For more information go to www.acdlabs.com/nmrproc/

##### RAD-324

08/11/2016 8:11:51 AM
Dr.Mustafa El-Araby Sample : RAD-324 DMSO

| Formula $C_{22}H_{23}FN_2O_2$ | FW 366.4286 |
|---|---|

| Acquisition Time (sec) | 1.9268 | Comment | Dr.Mustafa El-Araby Sample : RAD-324 DMSO | | | Date | 08 Apr 2015 11:21:20 |
|---|---|---|---|---|---|---|---|
| Date Stamp | 08 Apr 2015 11:21:20 | | | | | | |
| File Name | E:\Mostafa Alaraby Project\Oxazolone NMR\NMR final oxazolone\MUSTAFA RAD-324 08-04-2015\170\fid | | | | | Frequency (MHz) | 850.15 |
| Nucleus | 1H | Number of Transients | 20 | Origin | spect | Original Points Count | 32768 |
| Owner | nmr | Points Count | 32768 | Pulse Sequence | zg30 | Receiver Gain | 9.04 |
| SW(cyclical) (Hz) | 17006.80 | Solvent | DMSO-d6 | Spectrum Offset (Hz) | 5250.0283 | Spectrum Type | STANDARD |
| Sweep Width (Hz) | 17006.28 | Temperature (degree C) | 25.001 | | | | |

$^{1}$H NMR (850 MHz, DMSO-$d_6$) δ 9.64 (s, 1H), 8.11 (t, $J$ = 5.71 Hz, 1H), 7.60 (dd, $J$ = 5.71, 8.82 Hz, 2H), 7.48 - 7.53 (m, $J$ = 7.78 Hz, 2H), 7.46 (d, $J$ = 16.09 Hz, 1H), 7.25 - 7.28 (m, $J$ = 8.30 Hz, 2H), 7.20 - 7.25 (m, 2H), 7.00 (s, 1H), 6.81 (d, $J$ = 15.57 Hz, 1H), 3.12 (q, $J$ = 6.75 Hz, 2H), 2.35 (s, 3H), 1.48 (sxt, $J$ = 7.27 Hz, 2H), 0.87 (t, $J$ = 7.27 Hz, 3H)

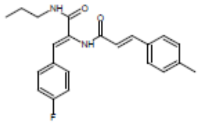


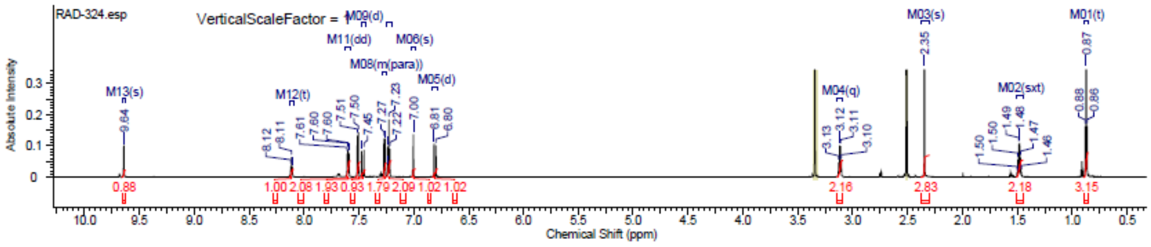


#### $^{13}$C NMR

This report was created by ACD/NMR Processor Academic Edition. For more information go to www.acdlabs.com/nmrproc/

##### RAD-324

20/09/2016 8:36:44 AM
Dr.Mostafa Sample : RAD-324 DMSO

| Formula $C_{22}H_{23}FN_2O_2$ | FW 366.4286 |
|---|---|

| Acquisition Time (sec) | 0.6423 | Comment | Dr.Mostafa Sample : RAD-324 DMSO | | | Date | 18 Apr 2016 12:12:32 |
|---|---|---|---|---|---|---|---|
| Date Stamp | 18 Apr 2016 12:12:32 | | | | | | |
| File Name | E:\Projects\Mostafa Alaraby Project\Oxazolone\Oxazolone NMR\NMR final oxazolone\13 CNMR New 18-4-2016\MUSTAFA RAD-324 18-04-2016\10\fid | | | | | | |
| Frequency (MHz) | 213.77 | Nucleus | 13C | Number of Transients | 1464 | Origin | spect |
| Original Points Count | 32768 | Owner | nmr | Points Count | 32768 | Pulse Sequence | zgpg30 |
| Receiver Gain | 186.93 | SW(cyclical) (Hz) | 51020.41 | Solvent | DMSO-d6 | Spectrum Offset (Hz) | 21293.8125 |
| Spectrum Type | STANDARD | Sweep Width (Hz) | 51018.85 | Temperature (degree C) | 25.002 | | |

$^{13}$C NMR (214 MHz, DMSO-$d_6$) δ 165.0, 164.8, 140.0, 139.7, 132.0, 131.5, 131.0, 130.3, 129.7, 127.8, 125.7, 120.5, 115.6, 115.5, 41.0, 39.8, 39.7, 39.6, 39.4, 39.3, 39.2, 22.4, 21.1, 11.5

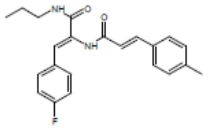


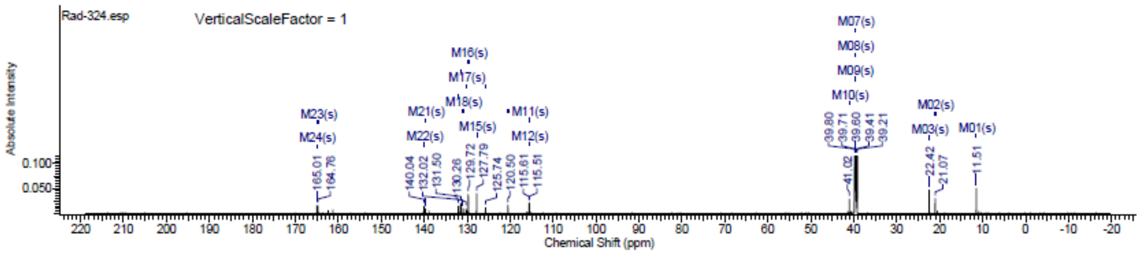

#### Spectra of (2612)

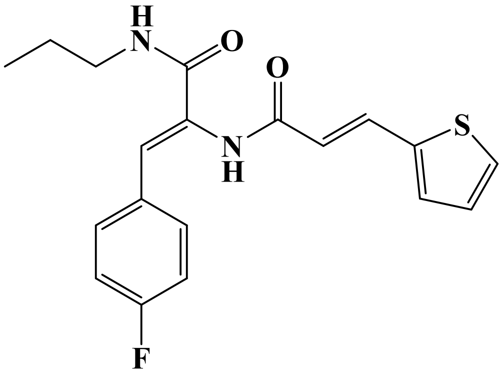


### LC/MS

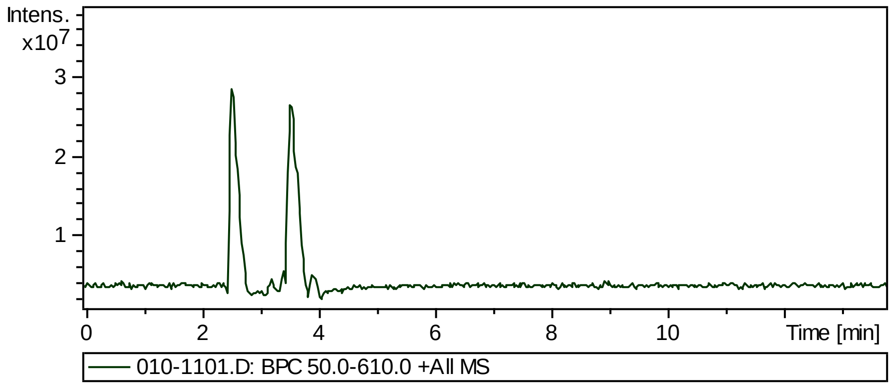


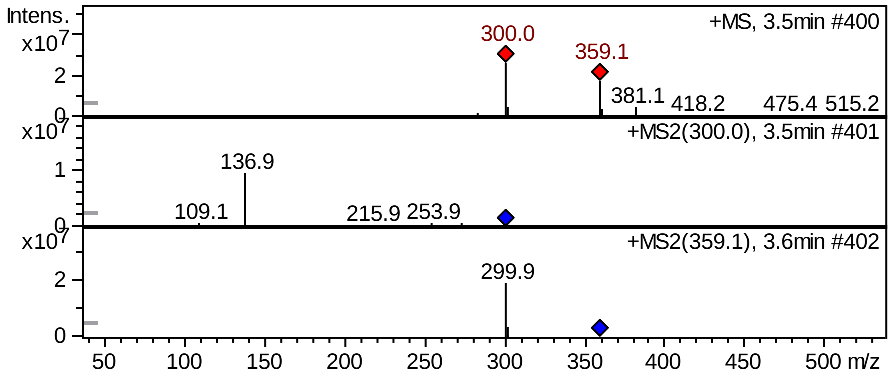


### FT-IR

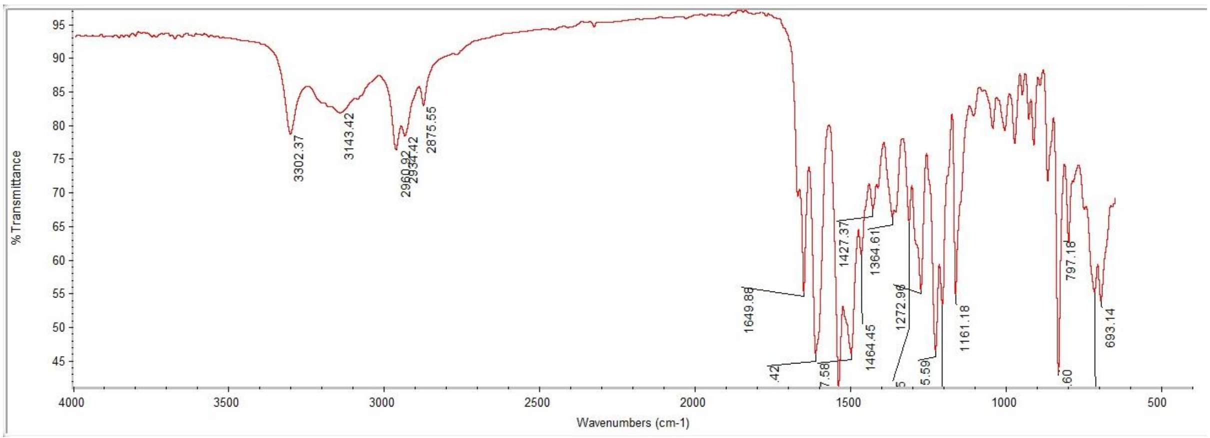

#### $^{1}$H NMR

This report was created by ACD/NMR Processor Academic Edition. For more information go to www.acdlabs.com/nmrproc/

**RAD-325**

18/02/2017 10:53:56 AM
Dr.Mostafa Sample : RAD-325 DMSO

| Formula $C_{19}H_{19}FN_2O_2S$ | FW 358.4298 |
|---|---|

| | | | | | | | |
|---|---|---|---|---|---|---|---|
| Acquisition Time (sec) | 1.9268 | Comment | Dr.Mostafa Sample : RAD-325 DMSO | | | Date | 14 Feb 2017 12:38:08 |
| Date Stamp | 14 Feb 2017 12:38:08 | | | | | | |
| File Name | E:\Projects\Mostafa Alaraby Project\Oxazolone\Rad 355 new amines\MOSTAFA RAD-325 14-02-2017\20\fid | | | | | Frequency (MHz) | 850.15 |
| Nucleus | 1H | Number of Transients | 64 | Origin | spect | Original Points Count | 32768 |
| Owner | nmr | Points Count | 32768 | Pulse Sequence | zg30 | Receiver Gain | 10.55 |
| SW(cyclical) (Hz) | 17006.80 | Solvent | DMSO-d6 | Spectrum Offset (Hz) | 5250.0283 | Spectrum Type | STANDARD |
| Sweep Width (Hz) | 17006.28 | Temperature (degree C) | 25.001 | | | | |

$^{1}$H NMR (850 MHz, DMSO-$d_6$) δ 9.63 (s, 1H), 8.11 (t, $J$ = 5.71 Hz, 1H), 7.61 - 7.68 (m, 2H), 7.58 (dd, $J$ = 5.97, 8.04 Hz, 2H), 7.43 (d, $J$ = 3.11 Hz, 1H), 7.23 (t, $J$ = 8.56 Hz, 2H), 7.11 - 7.16 (m, 1H), 7.00 (s, 1H), 6.60 (d, $J$ = 15.57 Hz, 1H), 3.11 (q, $J$ = 6.40 Hz, 2H), 1.48 (sxt, $J$ = 7.27 Hz, 2H), 0.87 (t, $J$ = 7.53 Hz, 3H)

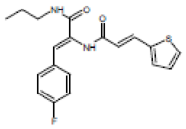


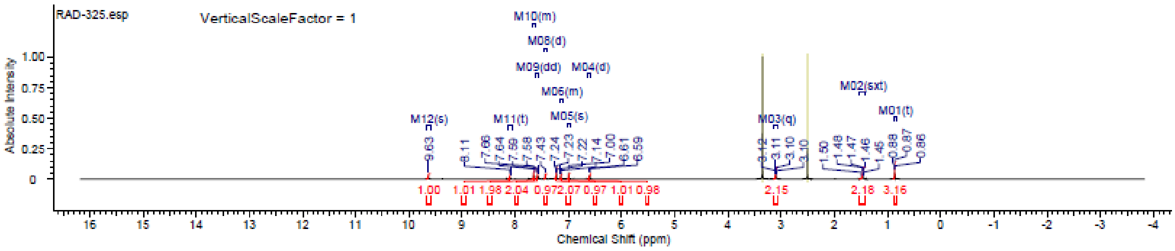


#### $^{13}$C NMR

This report was created by ACD/NMR Processor Academic Edition. For more information go to www.acdlabs.com/nmrproc/

**RAD-325**

25/09/2016 2:04:21 PM
Dr.Mostafa Sample : RAD-325 DMSO

| Formula $C_{19}H_{19}FN_2O_2S$ | FW 358.4298 |
|---|---|

| | | | | | | | |
|---|---|---|---|---|---|---|---|
| Acquisition Time (sec) | 0.6423 | Comment | Dr.Mostafa Sample : RAD-325 DMSO | | | Date | 19 Apr 2016 09:47:28 |
| Date Stamp | 19 Apr 2016 09:47:28 | | | | | | |
| File Name | E:\Projects\Mostafa Alaraby Project\Oxazolone\Oxazolone NMR\NMR final oxazolone\13 CNMR New 18-4-2016\MUSTAFA RAD-325 18-04-2016\100\fid | | | | | | |
| Frequency (MHz) | 213.77 | Nucleus | 13C | Number of Transients | 1930 | Origin | spect |
| Original Points Count | 32768 | Owner | nmr | Points Count | 32768 | Pulse Sequence | zgpg30 |
| Receiver Gain | 186.93 | SW(cyclical) (Hz) | 51020.41 | Solvent | DMSO-d6 | Spectrum Offset (Hz) | 21298.4824 |
| Spectrum Type | STANDARD | Sweep Width (Hz) | 51018.85 | Temperature (degree C) | 25.000 | | |

$^{13}$C NMR (214 MHz, DMSO-$d_6$) δ 165.0, 164.4, 139.8, 133.1, 131.6, 131.5, 131.4, 130.9, 130.2, 128.6, 128.5, 126.3, 125.9, 120.3, 41.0, 39.8, 39.7, 39.6, 39.4, 39.3, 39.2, 22.4, 11.5

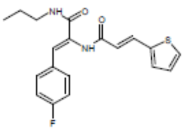


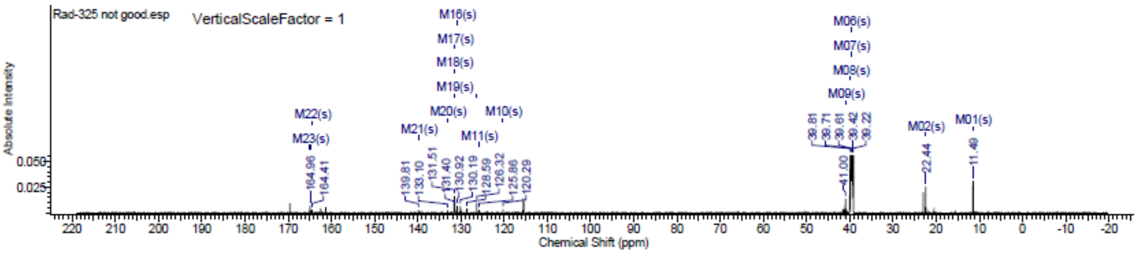

### Spectra of (2712)

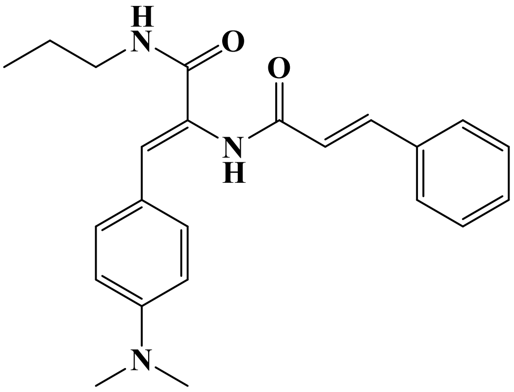


## LC/MS

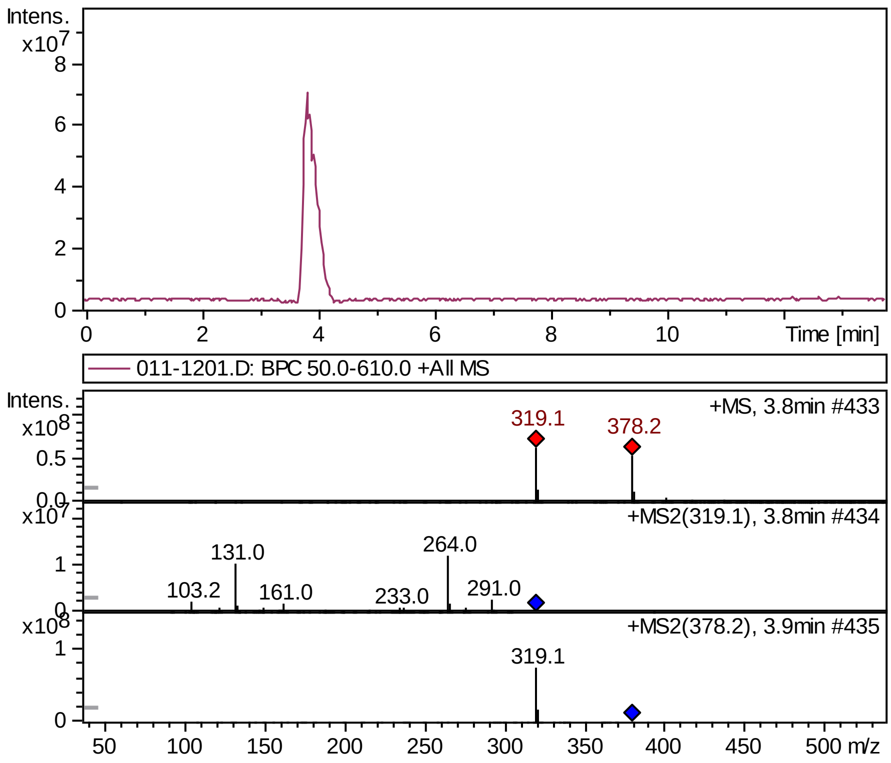


## FT-IR

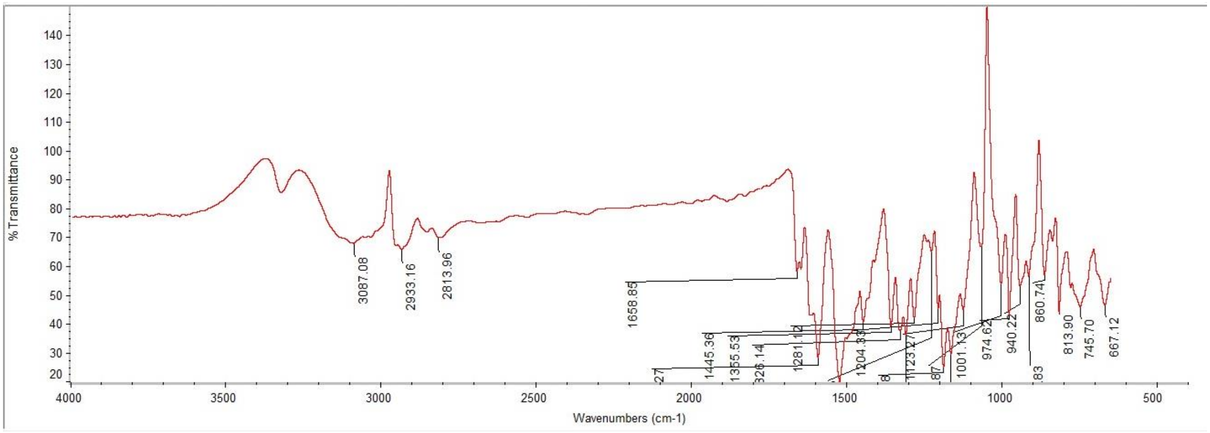

### Supplementary Information

#### 1H NMR

This report was created by ACD/NMR Processor Academic Edition. For more information go to www.acdlabs.com/nmrproc/

**RAD-331**

08/11/2016 8:20:31 AM
Dr.Mustafa Sample : RAD-331 DMSO PROTON DMSO {D:\Magdy} nmr 34

| | | | |
|---|---|---|---|
| *Formula* $C_{22}H_{27}N_3O_2$ | | *FW* | 377.4794 |

| | | | | | | | |
|---|---|---|---|---|---|---|---|
| *Acquisition Time (sec)* | 2.6564 | *Comment* | Dr.Mustafa Sample : RAD-331 DMSO PROTON DMSO {D:\Magdy} nmr 34 | | | | |
| *Date* | 18 Jun 2015 16:30:24 | | | *Date Stamp* | 18 Jun 2015 16:30:24 | | |
| *File Name* | E:\Mostafa Alaraby Project\Oxazolone NMR\NMR final oxazolone\MUSTAFA RAD-331 18-06-2015\10\fid | | | | | *Frequency (MHz)* | 600.15 |
| *Nucleus* | 1H | *Number of Transients* | 32 | *Origin* | spect | *Original Points Count* | 32768 |
| *Owner* | nmr | *Points Count* | 32768 | *Pulse Sequence* | zg30 | *Receiver Gain* | 128.00 |
| *SW(cyclical) (Hz)* | 12335.53 | *Solvent* | DMSO-d6 | *Spectrum Offset (Hz)* | 3706.1750 | *Spectrum Type* | STANDARD |
| *Sweep Width (Hz)* | 12335.15 | *Temperature (degree C)* | 25.000 | | | | |

$^1$H NMR (600 MHz, DMSO-$d_6$) δ 9.51 (s, 1H), 7.91 (t, $J$ = 5.84 Hz, 1H), 7.63 (d, $J$ = 7.15 Hz, 2H), 7.45 - 7.52 (m, 3H), 7.40 - 7.44 (m, 3H), 7.04 (s, 1H), 6.90 (d, $J$ = 15.81 Hz, 1H), 6.70 (d, $J$ = 9.03 Hz, 2H), 3.07 - 3.14 (m, 2H), 2.92 (s, 6H), 1.46 (sxt, $J$ = 7.23 Hz, 2H), 0.86 (t, $J$ = 7.34 Hz, 3H)

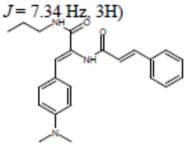


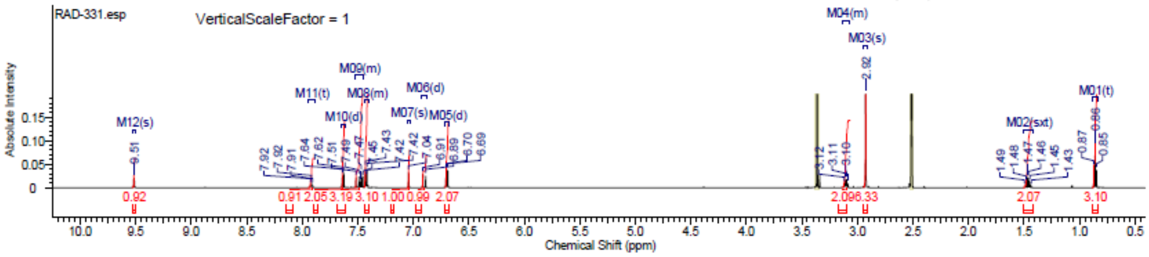


#### 13C NMR

This report was created by ACD/NMR Processor Academic Edition. For more information go to www.acdlabs.com/nmrproc/

**RAD-331**

25/09/2016 2:12:54 PM
Dr.Mustafa Sample ; RAD-331 DMSO

| | | | |
|---|---|---|---|
| *Formula* $C_{22}H_{27}N_3O_2$ | | *FW* | 377.4794 |

| | | | | | | | |
|---|---|---|---|---|---|---|---|
| *Acquisition Time (sec)* | 0.6423 | *Comment* | Dr.Mustafa Sample ; RAD-331 DMSO | | | *Date* | 20 Apr 2016 10:49:20 |
| *Date Stamp* | 20 Apr 2016 10:49:20 | | | | | | |
| *File Name* | E:\Projects\Mostafa Alaraby Project\Oxazolone\Oxazolone NMR\NMR final oxazolone\13 CNMR New 18-4-2016\MUSTAFA RAD-331 19-04-2016\100\fid | | | | | | |
| *Frequency (MHz)* | 213.77 | *Nucleus* | 13C | *Number of Transients* | 3007 | *Origin* | spect |
| *Original Points Count* | 32768 | *Owner* | nmr | *Points Count* | 32768 | *Pulse Sequence* | zgpg30 |
| *Receiver Gain* | 186.93 | *SW(cyclical) (Hz)* | 51020.41 | *Solvent* | DMSO-d6 | *Spectrum Offset (Hz)* | 21320.2813 |
| *Spectrum Type* | STANDARD | *Sweep Width (Hz)* | 51018.85 | *Temperature (degree C)* | 24.999 | | |

$^{13}$C NMR (214 MHz, DMSO-$d_6$) δ 169.6, 168.0, 142.5, 141.1, 135.8, 134.7, 131.2, 129.3, 129.0, 128.7, 128.1, 127.5, 126.3, 111.0, 41.6, 40.7, 39.8, 39.7, 39.6, 39.4, 39.3, 39.2, 22.3, 11.4

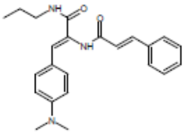


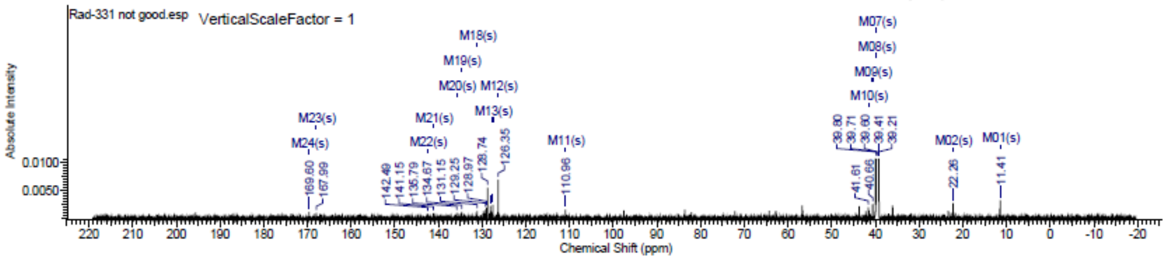

#### Spectra of (2812)

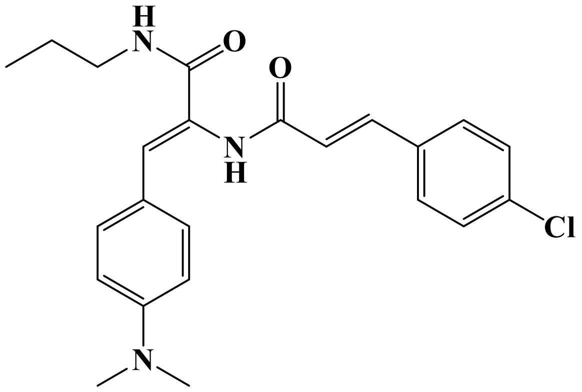

### LC/MS

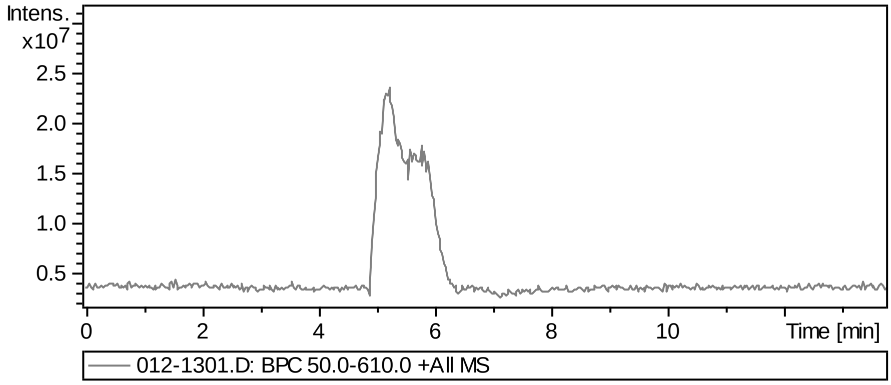


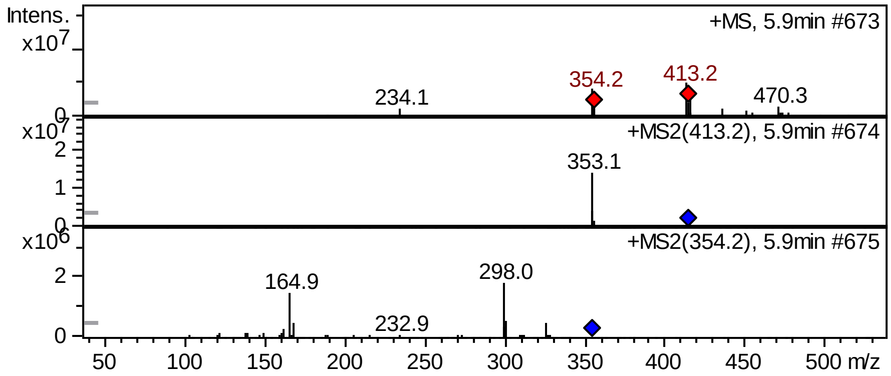


### FT-IR

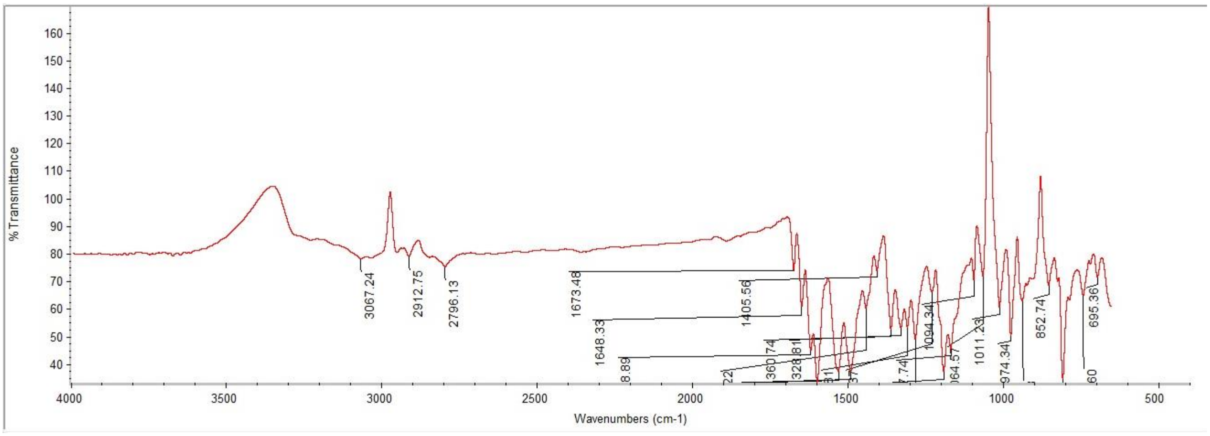

### Supplementary Information

#### $^{1}$H NMR

This report was created by ACD/NMR Processor Academic Edition. For more information go to www.acdlabs.com/nmrproc/

**RAD-332**

08/11/2016 8:27:20 AM
Dr.Mustafa Sample : RAD-332 DMSO PROTON DMSO {D:\Magdy} nmr 15

| Formula | $C_{23}H_{26}ClN_3O_2$ | FW | 411.9244 |
|---|---|---|---|

| Acquisition Time (sec) | 2.6564 | Comment | Dr.Mustafa Sample : RAD-332 DMSO PROTON DMSO {D:\Magdy} nmr 15 | | | | |
|---|---|---|---|---|---|---|---|
| Date | 18 Jun 2015 14:52:16 | | | Date Stamp | 18 Jun 2015 14:52:16 | | |
| File Name | E:\Mostafa Alaraby Project\Oxazolone NMR\NMR final oxazolone\MUSTAFA RAD-332 18-06-2015\10\fid | | | | | Frequency (MHz) | 600.15 |
| Nucleus | 1H | Number of Transients | 32 | Origin | spect | Original Points Count | 32768 |
| Owner | nmr | Points Count | 32768 | Pulse Sequence | zg30 | Receiver Gain | 128.00 |
| SW(cyclical) (Hz) | 12335.53 | Solvent | DMSO-d6 | Spectrum Offset (Hz) | 3706.1750 | Spectrum Type | STANDARD |
| Sweep Width (Hz) | 12335.15 | Temperature (degree C) | 25.000 | | | | |

$^{1}$H NMR (600 MHz, DMSO-$d_6$) δ 9.53 (s, 1H), 7.92 (t, $J$ = 6.02 Hz, 1H), 7.63 - 7.68 (m, $J$ = 8.66 Hz, 2H), 7.52 - 7.55 (m, $J$ = 8.28 Hz, 2H), 7.50 (d, $J$ = 15.81 Hz, 1H), 7.39 - 7.44 (m, $J$ = 9.03 Hz, 2H), 7.04 (s, 1H), 6.90 (d, $J$ = 15.81 Hz, 1H), 6.67 - 6.72 (m, $J$ = 9.03 Hz, 2H), 3.10 (q, $J$ = 6.40 Hz, 2H), 2.93 (s, 6H), 1.46 (sxt, $J$ = 7.23 Hz, 2H), 0.86 (t, $J$ = 7.34 Hz, 3H)

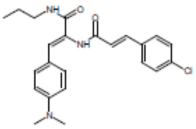


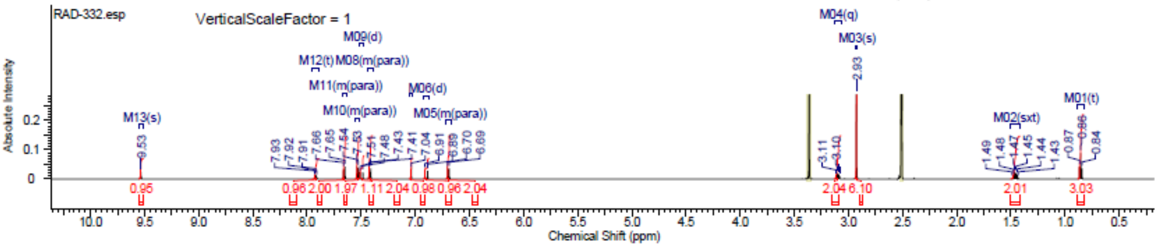


#### $^{13}$C NMR

This report was created by ACD/NMR Processor Academic Edition. For more information go to www.acdlabs.com/nmrproc/

**RAD-332**

25/09/2016 3:09:37 PM
Dr.Mostafa Sample ; RAD-332 DMSO

| Formula | $C_{23}H_{26}ClN_3O_2$ | FW | 411.9244 |
|---|---|---|---|

| Acquisition Time (sec) | 0.6423 | Comment | Dr.Mostafa Sample ; RAD-332 DMSO | | | Date | 20 Apr 2016 07:13:52 |
|---|---|---|---|---|---|---|---|
| Date Stamp | 20 Apr 2016 07:13:52 | | | | | | |
| File Name | E:\Projects\Mostafa Alaraby Project\Oxazolone\Oxazolone NMR\NMR final oxazolone\13 CNMR New 18-4-2016\MUSTAFA RAD-332 19-04-2016\70\fid | | | | | | |
| Frequency (MHz) | 213.77 | Nucleus | 13C | Number of Transients | 3500 | Origin | spect |
| Original Points Count | 32768 | Owner | nmr | Points Count | 32768 | Pulse Sequence | zgpg30 |
| Receiver Gain | 186.93 | SW(cyclical) (Hz) | 51020.41 | Solvent | DMSO-d6 | Spectrum Offset (Hz) | 21286.0273 |
| Spectrum Type | STANDARD | Sweep Width (Hz) | 51018.85 | Temperature (degree C) | 25.001 | | |

$^{13}$C NMR (214 MHz, DMSO-$d_6$) δ 166.5, 163.1, 140.1, 137.9, 134.6, 133.4, 131.2, 129.4, 129.3, 128.8, 128.6, 128.2, 127.8, 127.5, 41.3, 40.5, 39.8, 39.7, 39.6, 39.4, 39.3, 39.2, 22.4, 11.4

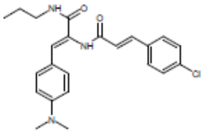


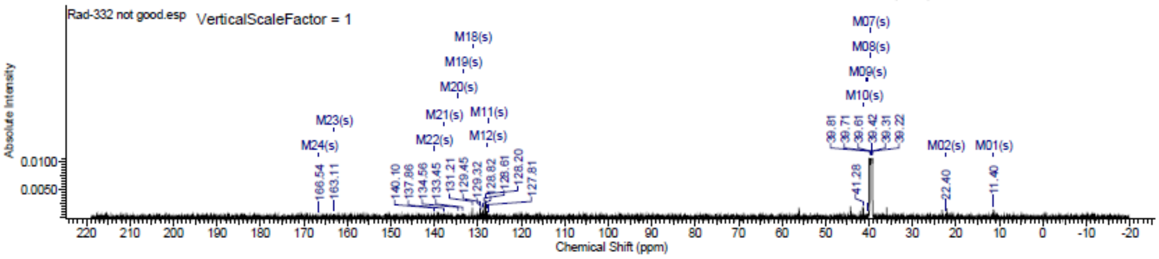

### Spectra of (2912)

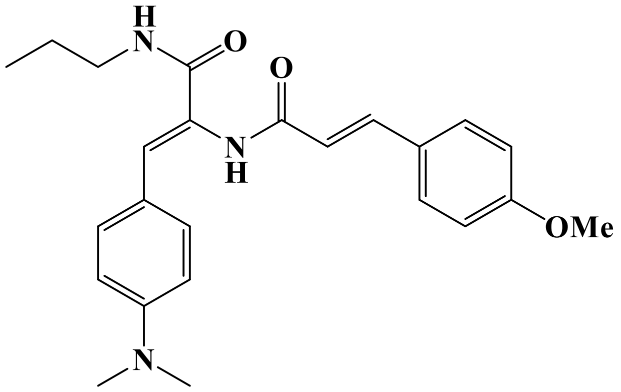


## LC/MS

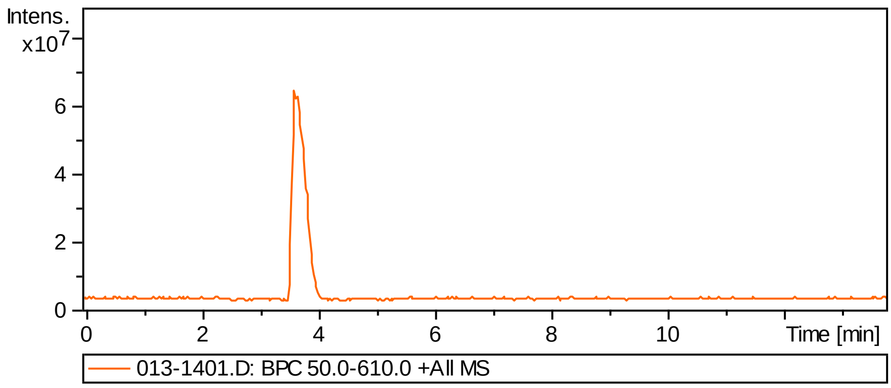


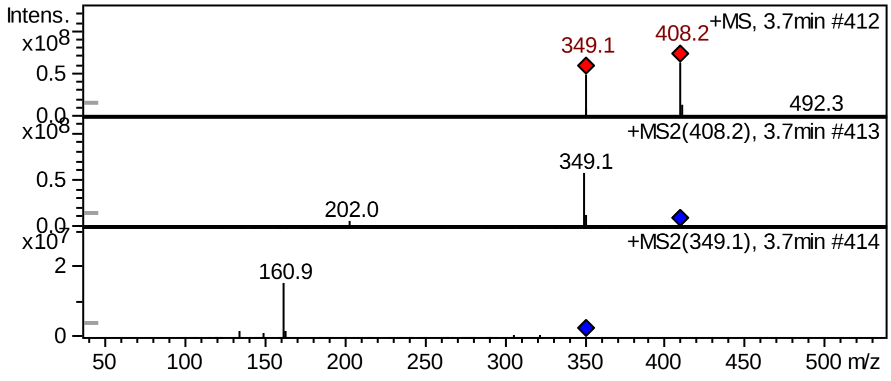


## FT-IR

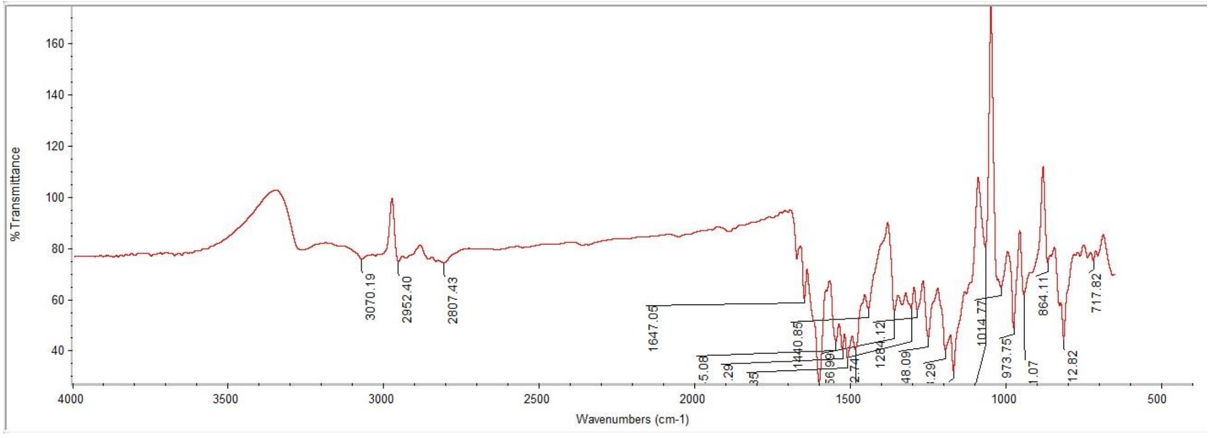

#### $^{1}$H NMR

This report was created by ACD/NMR Processor Academic Edition. For more information go to www.acdlabs.com/nmrproc/

**RAD-333**

08/11/2016 8:30:41 AM
Dr.Mustafa Sample : RAD-333 DMSO PROTON DMSO {D:\Magdy} nmr 18

| | | | |
|---|---|---|---|
| *Formula* $C_{24}H_{[illegible]}N_3O_3$ | | *FW* | 407.5054 |

| | | | | | | | |
|---|---|---|---|---|---|---|---|
| *Acquisition Time (sec)* | 2.6564 | *Comment* | Dr.Mustafa Sample : RAD-333 DMSO PROTON DMSO {D:\Magdy} nmr 18 | | | | |
| *Date* | 18 Jun 2015 15:07:12 | | | *Date Stamp* | 18 Jun 2015 15:07:12 | | |
| *File Name* | E:\Mostafa Alaraby Project\Oxazolone NMR\NMR final oxazolone\MUSTAFA RAD-333 18-06-2015\10\fid | | | | | *Frequency (MHz)* | 600.15 |
| *Nucleus* | 1H | *Number of Transients* | 32 | *Origin* | spect | *Original Points Count* | 32768 |
| *Owner* | nmr | *Points Count* | 32768 | *Pulse Sequence* | zg30 | *Receiver Gain* | 128.00 |
| *SW(cyclical) (Hz)* | 12335.53 | *Solvent* | DMSO-d6 | *Spectrum Offset (Hz)* | 3706.1750 | *Spectrum Type* | STANDARD |
| *Sweep Width (Hz)* | 12335.15 | *Temperature (degree C)* | 25.000 | | | | |

$^{1}$H NMR (600 MHz, DMSO-$d_6$) δ 9.41 (s, 1H), 7.88 (t, *J* = 5.83 Hz, 1H), 7.58 (d, *J* = 9.04 Hz, 2H), 7.40 - 7.47 (m, 3H), 7.01 - 7.04 (m, 3H), 6.75 (d, *J* = 15.81 Hz, 1H), 6.69 (d, *J* = 9.03 Hz, 2H), 3.81 (s, 3H), 3.10 (q, *J* = 6.40 Hz, 2H), 2.92 (s, 6H), 1.46 (sxt, *J* = 7.30 Hz, 2H), 0.86 (t, *J* = 7.34 Hz, 3H)

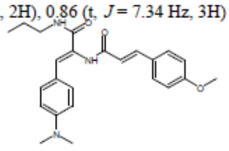

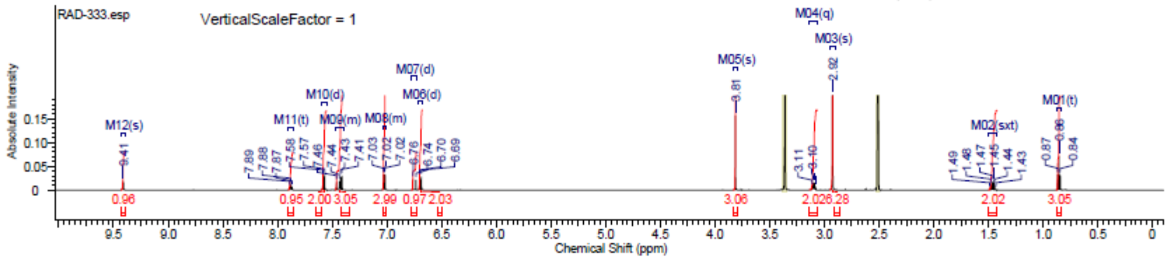


#### $^{13}$C NMR

This report was created by ACD/NMR Processor Academic Edition. For more information go to www.acdlabs.com/nmrproc/

**RAD-333**

25/09/2016 3:18:33 PM
Dr.Moustafa Sample : RAD-333 DMSO

| | | | |
|---|---|---|---|
| *Formula* $C_{24}H_{[illegible]}N_3O_3$ | | *FW* | 407.5053 |

| | | | | | | | |
|---|---|---|---|---|---|---|---|
| *Acquisition Time (sec)* | 0.6423 | *Comment* | Dr.Moustafa Sample : RAD-333 DMSO | | | *Date* | 22 Apr 2016 05:48:32 |
| *Date Stamp* | 22 Apr 2016 05:48:32 | | | | | | |
| *File Name* | E:\Projects\Mostafa Alaraby Project\Oxazolone\Oxazolone NMR\NMR final oxazolone\13 CNMR New 18-4-2016\MUSTAFA RAD-333 21-04-2016\20\fid | | | | | | |
| *Frequency (MHz)* | 213.77 | *Nucleus* | 13C | *Number of Transients* | 12288 | *Origin* | spect |
| *Original Points Count* | 32768 | *Owner* | nmr | *Points Count* | 32768 | *Pulse Sequence* | zgpg30 |
| *Receiver Gain* | 186.93 | *SW(cyclical) (Hz)* | 51020.41 | *Solvent* | DMSO-d6 | *Spectrum Offset (Hz)* | 21286.0273 |
| *Spectrum Type* | STANDARD | *Sweep Width (Hz)* | 51018.85 | *Temperature (degree C)* | 24.999 | | |

$^{13}$C NMR (214 MHz, DMSO-$d_6$) δ 164.3, 163.1, 158.6, 142.3, 134.5, 133.1, 131.9, 131.4, 129.4, 128.6, 119.0, 114.6, 113.9, 113.2, 55.3, 41.5, 40.5, 39.8, 39.7, 39.6, 39.4, 39.3, 39.2, 22.1, 11.3

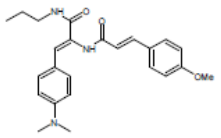

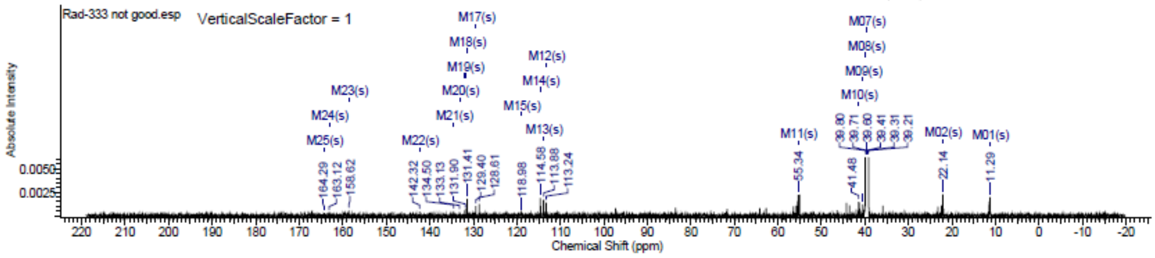

### Spectra of (3012)

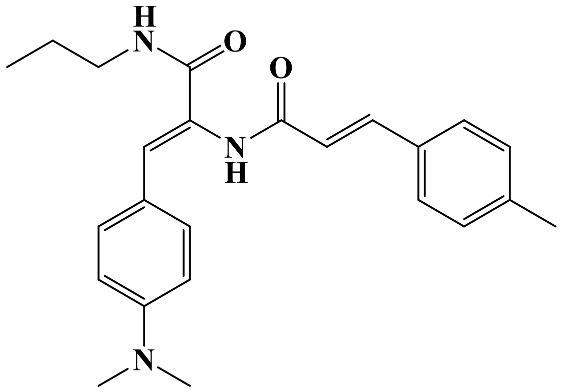

## LC/MS

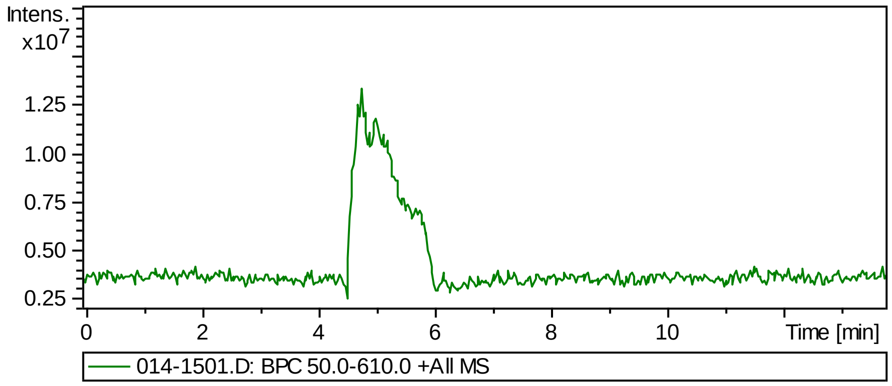


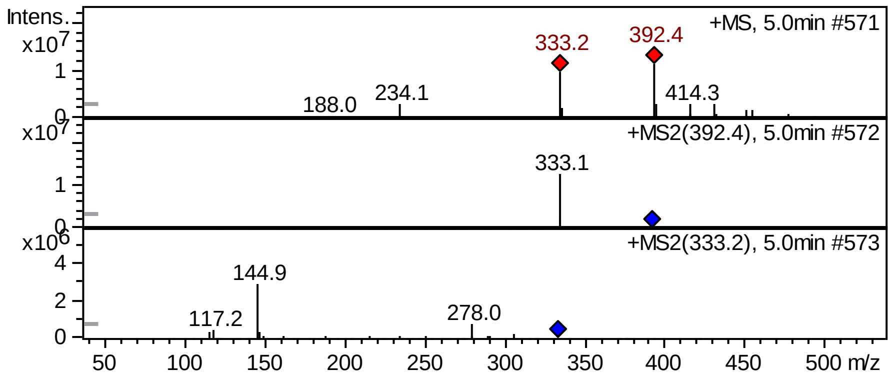


## FT-IR

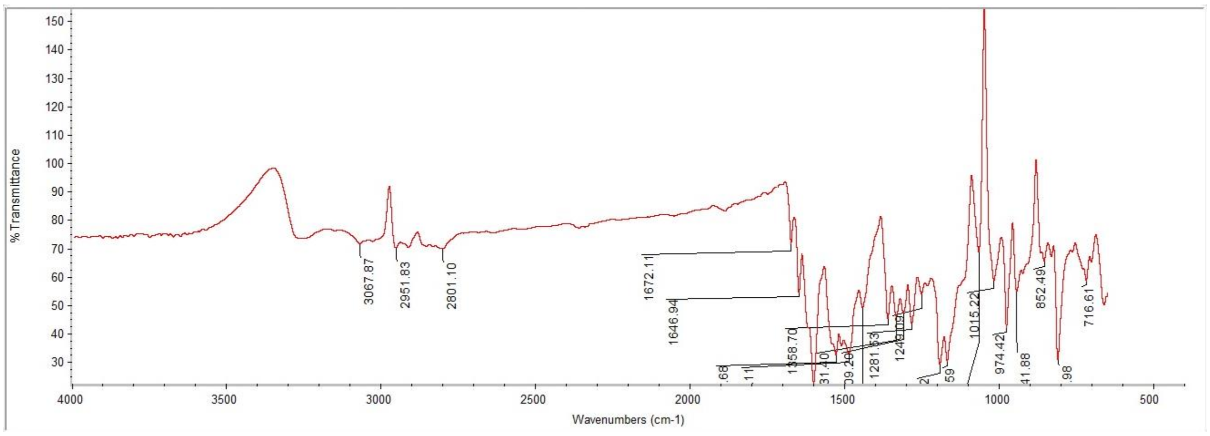

#### $^1$H NMR

This report was created by ACD/NMR Processor Academic Edition. For more information go to www.acdlabs.com/nmrproc/

##### RAD-334

18/02/2017 10:53:27 AM
Dr.Mostafa Sample : RAD-334 DMSO

| Formula $C_{24}H_{29}N_3O_2$ | FW 391.5060 |
|---|---|

| | | | | | | | |
|---|---|---|---|---|---|---|---|
| Acquisition Time (sec) | 1.9268 | Comment | Dr.Mostafa Sample : RAD-334 DMSO | | | Date | 14 Feb 2017 12:44:32 |
| Date Stamp | 14 Feb 2017 12:44:32 | | | | | | |
| File Name | E:\Projects\Mostafa Alaraby Project\Oxazolone\Rad 355 new amines\MOSTAFA RAD-334 14-02-2017\30\fid | | | | | Frequency (MHz) | 850.15 |
| Nucleus | 1H | Number of Transients | 64 | Origin | spect | Original Points Count | 32768 |
| Owner | nmr | Points Count | 32768 | Pulse Sequence | zg30 | Receiver Gain | 10.55 |
| SW(cyclical) (Hz) | 17006.80 | Solvent | DMSO-d6 | Spectrum Offset (Hz) | 5250.0283 | Spectrum Type | STANDARD |
| Sweep Width (Hz) | 17006.28 | Temperature (degree C) | 25.000 | | | | |

$^1$H NMR (850 MHz, DMSO-$d_6$) δ 9.51 (br. s., 1H), 7.96 (t, $J$ = 5.97 Hz, 1H), 7.45 - 7.52 (m, 2H), 7.24 - 7.28 (m, $J$ = 7.78 Hz, 2H), 7.21 (s, 1H), 7.04 (s, 1H), 6.99 (d, $J$ = 8.30 Hz, 1H), 6.84 (d, $J$ = 15.57 Hz, 1H), 6.76 (d, $J$ = 8.30 Hz, 1H), 3.11 (q, $J$ = 6.57 Hz, 2H), 2.51 (br. s., 9H), 2.34 (s, 2H), 1.47 (sxt, $J$ = 7.16 Hz, 2H), 0.85 - 0.87 (m, 3H)

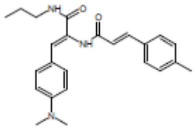

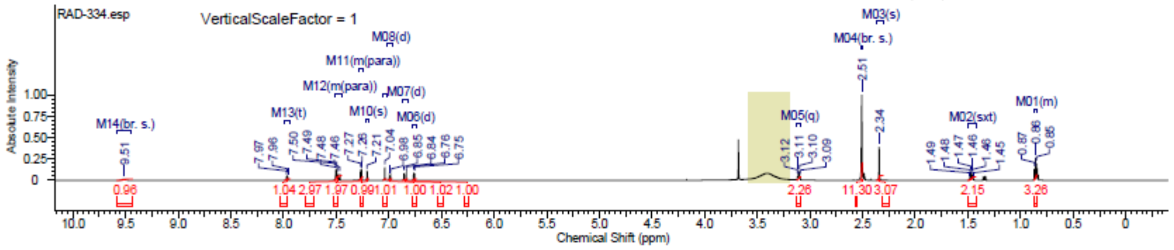

#### Spectra of (3112)

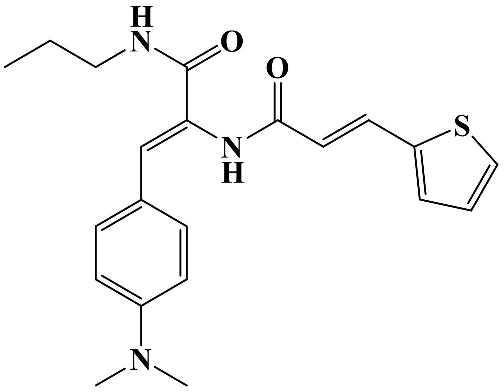

### LC/MS

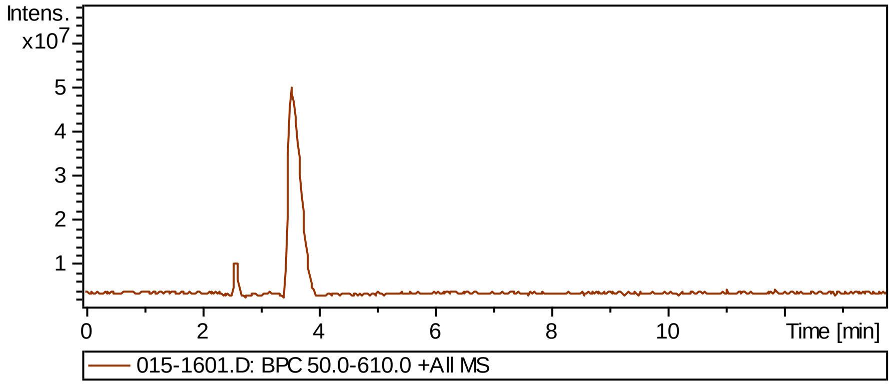


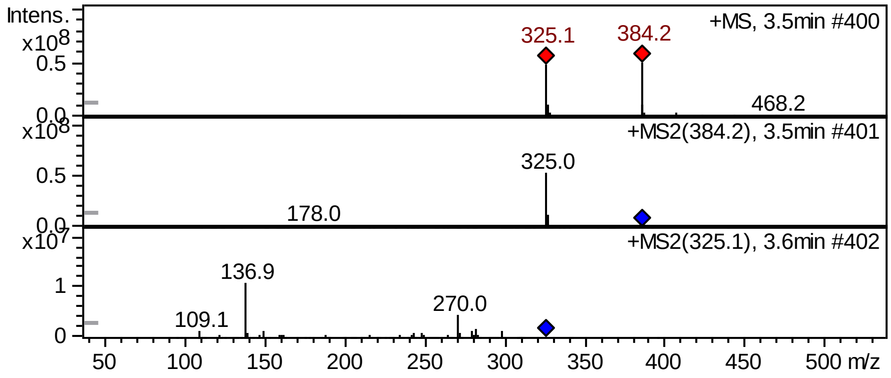


### FT-IR

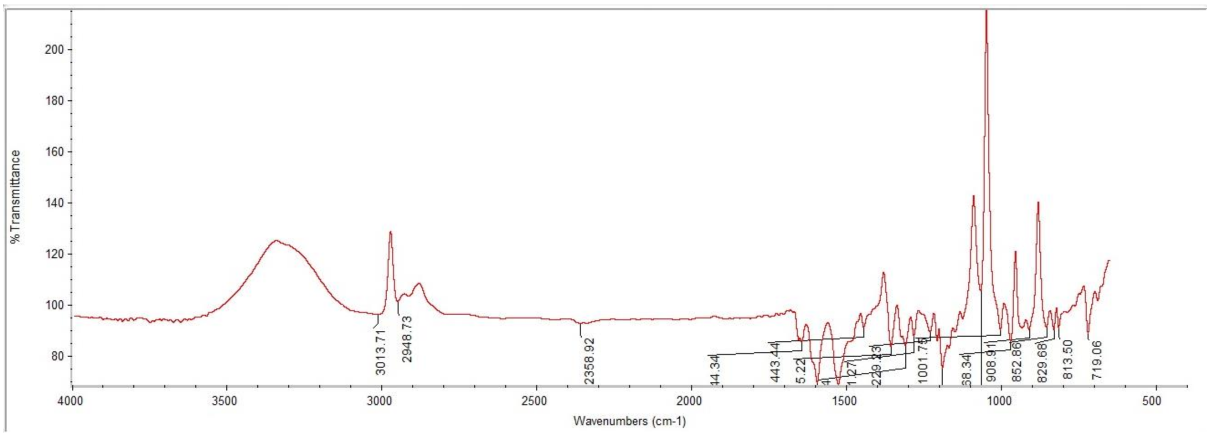

### [1]H NMR

This report was created by ACD/NMR Processor Academic Edition. For more information go to www.acdlabs.com/nmrproc/

#### RAD-335

08/11/2016 8:37:35 AM
Dr.Mustafa Sample : RAD-335 DMSO PROTON DMSO {D:\Magdy} nmr 16

| Formula | $C_{21}H_{25}N_3O_2S$ | FW | 383.5071 |
|---|---|---|---|

| Acquisition Time (sec) | 2.6564 | Comment | Dr.Mustafa Sample : RAD-335 DMSO PROTON DMSO {D:\Magdy} nmr 16 | | | | |
|---|---|---|---|---|---|---|---|
| Date | 18 Jun 2015 14:56:32 | | | Date Stamp | 18 Jun 2015 14:56:32 | | |
| File Name | E:\Mostafa Alaraby Project\Oxazolone NMR\NMR final oxazolone\MUSTAFA RAD-335 18-06-2015\10\fid | | | | | Frequency (MHz) | 600.15 |
| Nucleus | 1H | Number of Transients | 32 | Origin | spect | Original Points Count | 32768 |
| Owner | nmr | Points Count | 32768 | Pulse Sequence | zg30 | Receiver Gain | 161.00 |
| SW(cyclical) (Hz) | 12335.53 | Solvent | DMSO-d6 | Spectrum Offset (Hz) | 3706.1750 | Spectrum Type | STANDARD |
| Sweep Width (Hz) | 12335.15 | Temperature (degree C) | 25.000 | | | | |

[1]H NMR (600 MHz, DMSO-$d_6$) δ 9.47 (s, 1H), 7.90 (t, $J$ = 5.83 Hz, 1H), 7.61 - 7.68 (m, 2H), 7.44 (d, $J$ = 3.39 Hz, 1H), 7.38 - 7.42 (m, $J$ = 9.04 Hz, 2H), 7.15 (dd, $J$ = 3.58, 5.08 Hz, 1H), 7.03 (s, 1H), 6.68 - 6.72 (m, $J$ = 9.03 Hz, 2H), 6.64 (d, $J$ = 15.81 Hz, 1H), 3.09 (q, $J$ = 6.40 Hz, 2H), 2.93 (s, 5H), 1.46 (sxt, $J$ = 7.30 Hz, 2H), 0.85 (t, $J$ = 7.34 Hz, 3H)

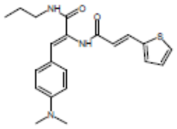


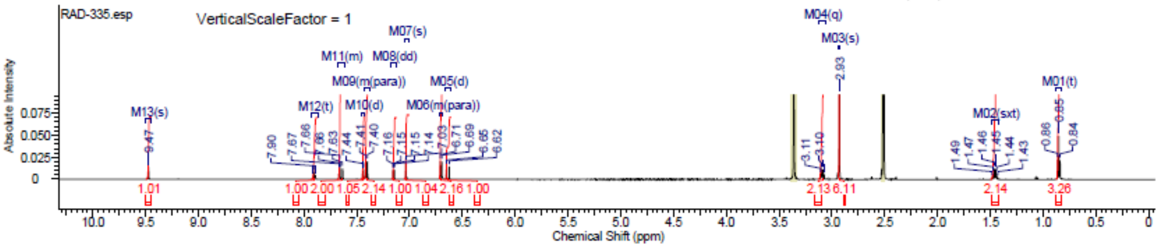

### Spectra of (3212)

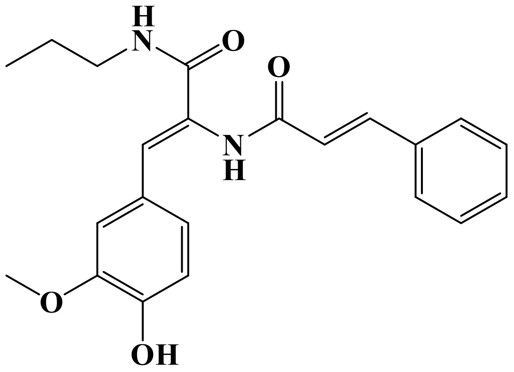

## LC/MS

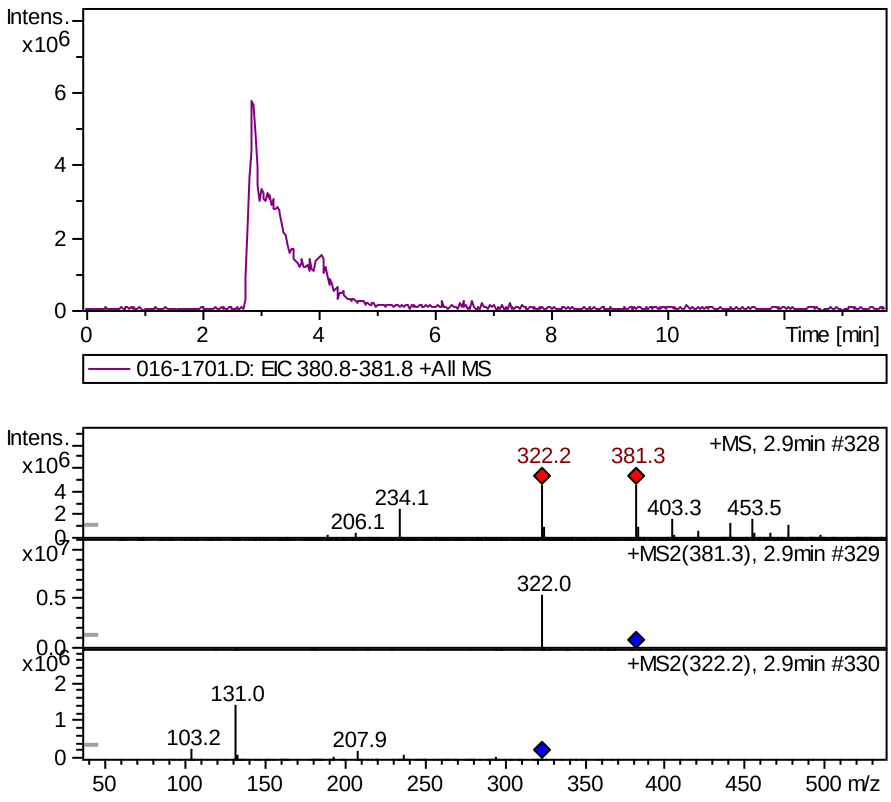


## FT-IR

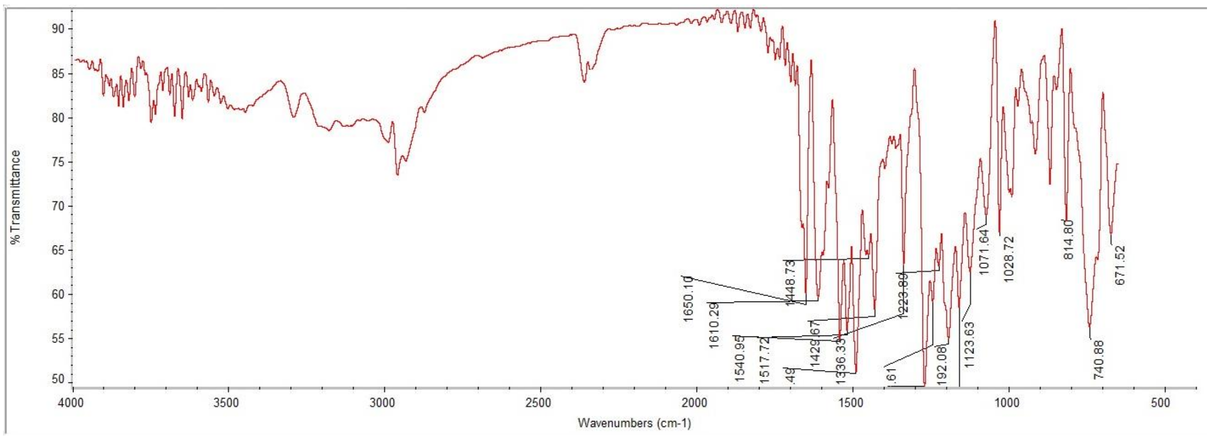

#### $^{1}$H NMR

This report was created by ACD/NMR Processor Academic Edition. For more information go to www.acdlabs.com/nmrproc/

**RAD-341**

08/11/2016 8:41:07 AM
Dr.Mustafa Sample : RAD-341 DMSO PROTON DMSO {D:\Magdy} nmr 23

| | |
|---|---|
| *Formula* $C_{20}H_{24}N_2O_4$ | *FW* 380.4370 |

| | | | | | | | |
|---|---|---|---|---|---|---|---|
| *Acquisition Time (sec)* | 2.6564 | *Comment* | Dr.Mustafa Sample : RAD-341 DMSO PROTON DMSO {D:\Magdy} nmr 23 | | | | |
| *Date* | 18 Jun 2015 15:32:48 | | | *Date Stamp* | 18 Jun 2015 15:32:48 | | |
| *File Name* | E:\Mostafa Alaraby Project\Oxazolone NMR\NMR final oxazolone\MUSTAFA RAD-341 18-06-2015\10\fid | | | | | *Frequency (MHz)* | 600.15 |
| *Nucleus* | 1H | *Number of Transients* | 32 | *Origin* | spect | *Original Points Count* | 32768 |
| *Owner* | nmr | *Points Count* | 32768 | *Pulse Sequence* | zg30 | *Receiver Gain* | 114.00 |
| *SW(cyclical) (Hz)* | 12335.53 | *Solvent* | DMSO-d6 | *Spectrum Offset (Hz)* | 3706.1750 | *Spectrum Type* | STANDARD |
| *Sweep Width (Hz)* | 12335.15 | *Temperature (degree C)* | 25.000 | | | | |

$^{1}$H NMR (600 MHz, DMSO-$d_6$) δ 8.02 (br. s., 1H), 7.61 (d, $J$ = 7.15 Hz, 2H), 7.39 - 7.55 (m, 4H), 7.21 (s, 1H), 7.04 (s, 1H), 6.91 (d, $J$ = 16.19 Hz, 1H), 6.77 (d, $J$ = 8.28 Hz, 1H), 3.69 (s, 3H), 3.11 (q, $J$ = 6.65 Hz, 2H), 1.47 (sxt, $J$ = 7.15 Hz, 2H), 0.86 (t, $J$ = 7.34 Hz, 3H)

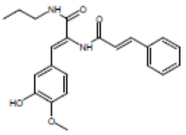


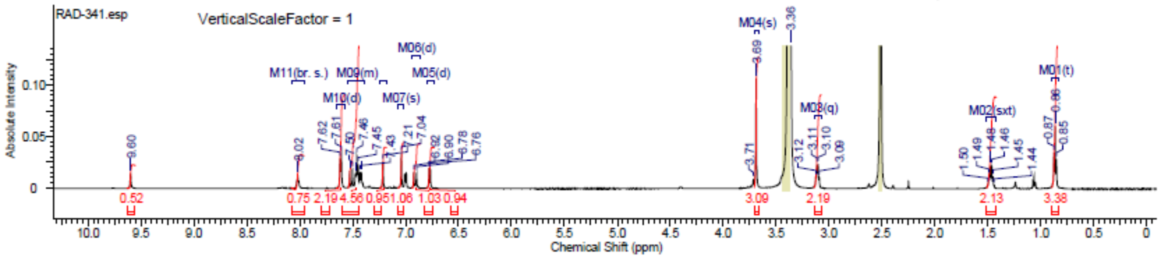


#### $^{13}$C NMR

This report was created by ACD/NMR Processor Academic Edition. For more information go to www.acdlabs.com/nmrproc/

**RAD-341**

20/09/2016 9:24:57 AM
Dr.Mustafa Sample : RAD-341 DMSO

| | |
|---|---|
| *Formula* $C_{20}H_{24}N_2O_4$ | *FW* 380.4370 |

| | | | | | | | |
|---|---|---|---|---|---|---|---|
| *Acquisition Time (sec)* | 0.6423 | *Comment* | Dr.Mustafa Sample : RAD-341 DMSO | | | *Date* | 19 Apr 2016 05:54:56 |
| *Date Stamp* | 19 Apr 2016 05:54:56 | | | | | | |
| *File Name* | E:\Projects\Mostafa Alaraby Project\Oxazolone\Oxazolone NMR\NMR final oxazolone\13 CNMR New 18-4-2016\MUSTAFA RAD-341 18-04-2016\80\fid | | | | | | |
| *Frequency (MHz)* | 213.77 | *Nucleus* | 13C | *Number of Transients* | 3072 | *Origin* | spect |
| *Original Points Count* | 32768 | *Owner* | nmr | *Points Count* | 32768 | *Pulse Sequence* | zgpg30 |
| *Receiver Gain* | 186.93 | *SW(cyclical) (Hz)* | 51020.41 | *Solvent* | DMSO-d6 | *Spectrum Offset (Hz)* | 21296.9258 |
| *Spectrum Type* | STANDARD | *Sweep Width (Hz)* | 51018.85 | *Temperature (degree C)* | 25.000 | | |

$^{13}$C NMR (214 MHz, DMSO-$d_6$) δ 165.2, 164.8, 147.5, 147.3, 139.8, 134.9, 129.9, 129.2, 128.6, 127.7, 127.2, 125.5, 123.8, 121.8, 115.5, 112.9, 55.4, 41.0, 39.8, 39.7, 39.6, 39.4, 39.3, 39.2, 22.5, 11.5

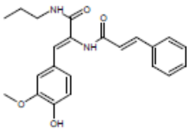


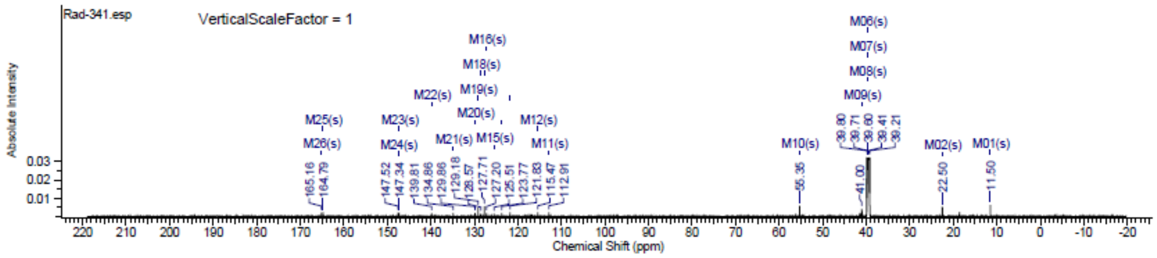

### Spectra of (3312)

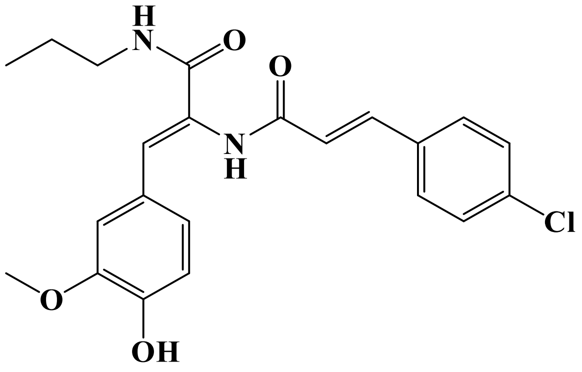


## LC/MS

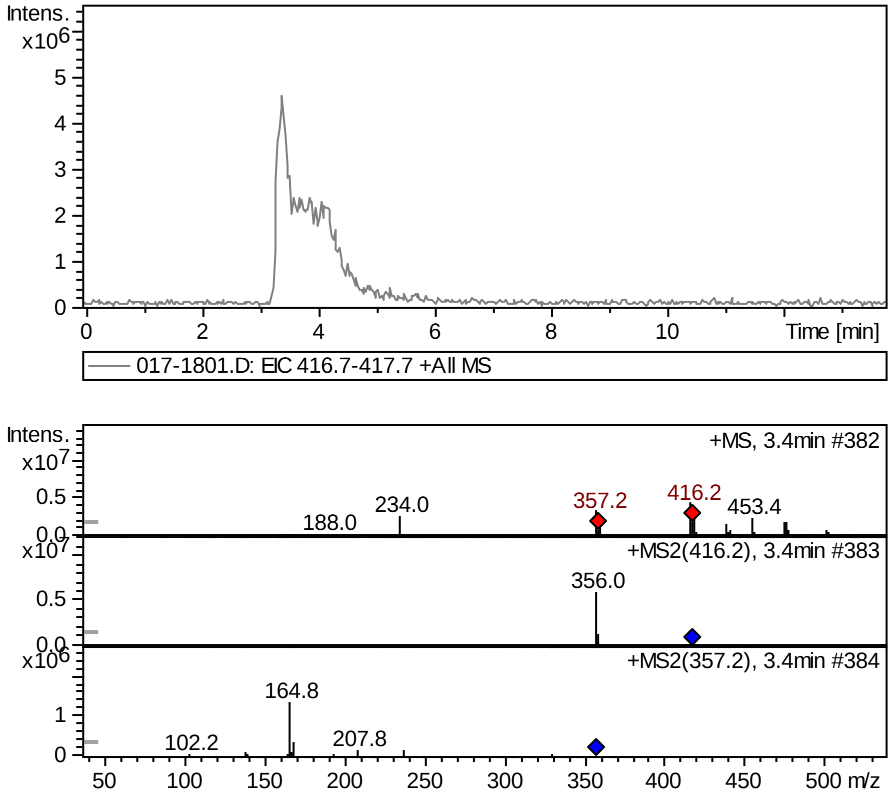


## FT-IR

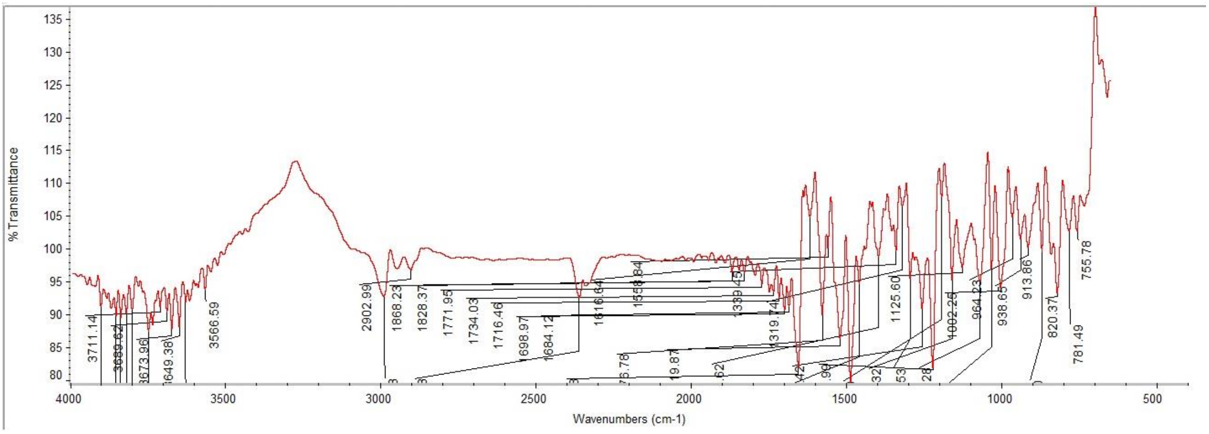

### $^1$H NMR

This report was created by ACD/NMR Processor Academic Edition. For more information go to www.acdlabs.com/nmrproc/

#### RAD-342

08/11/2016 8:51:48 AM
Dr.Mustafa Sample : RAD-342 DMSO PROTON DMSO {D:\Magdy} nmr 33

| Formula $C_{22}H_{23}ClN_2O_4$ | FW 414.8820 |
|---|---|

| | | | | | | | |
|---|---|---|---|---|---|---|---|
| Acquisition Time (sec) | 2.6564 | Comment | Dr.Mustafa Sample : RAD-342 DMSO PROTON DMSO {D:\Magdy} nmr 33 | | | | |
| Date | 18 Jun 2015 16:24:00 | | | Date Stamp | 18 Jun 2015 16:24:00 | | |
| File Name | E:\Mostafa Alaraby Project\Oxazolone NMR\NMR final oxazolone\MUSTAFA RAD-342 18-06-2015\10\fid | | | | | Frequency (MHz) | 600.15 |
| Nucleus | 1H | Number of Transients | 32 | Origin | spect | Original Points Count | 32768 |
| Owner | nmr | Points Count | 32768 | Pulse Sequence | zg30 | Receiver Gain | 144.00 |
| SW(cyclical) (Hz) | 12335.53 | Solvent | DMSO-d6 | Spectrum Offset (Hz) | 3706.1750 | Spectrum Type | STANDARD |
| Sweep Width (Hz) | 12335.15 | Temperature (degree C) | 25.000 | | | | |

$^1$H NMR (600 MHz, DMSO-$d_6$) δ 9.61 (br. s., 1H), 8.01 (t, $J$ = 5.83 Hz, 1H), 7.64 (d, $J$ = 8.28 Hz, 2H), 7.48 - 7.56 (m, 3H), 7.20 (d, $J$ = 2.26 Hz, 1H), 7.04 (s, 1H), 6.99 (dd, $J$ = 1.88, 8.28 Hz, 1H), 6.90 (d, $J$ = 16.19 Hz, 1H), 6.76 (d, $J$ = 8.28 Hz, 1H), 3.68 (s, 3H), 3.08 - 3.13 (m, 2H), 1.47 (sxt, $J$ = 7.23 Hz, 2H), 0.85 - 0.88 (m, 3H)

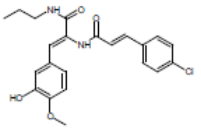


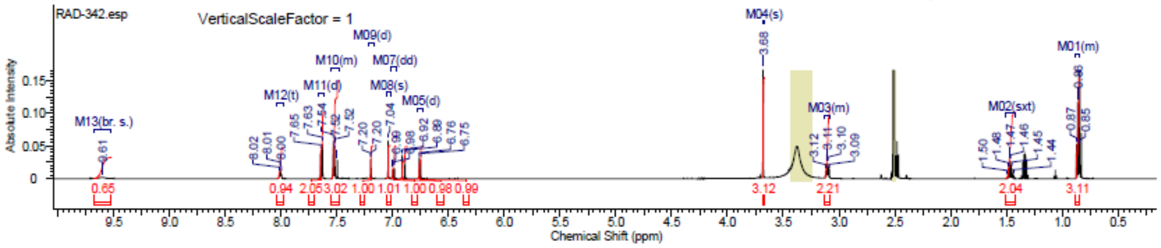

#### Spectra of (3412)

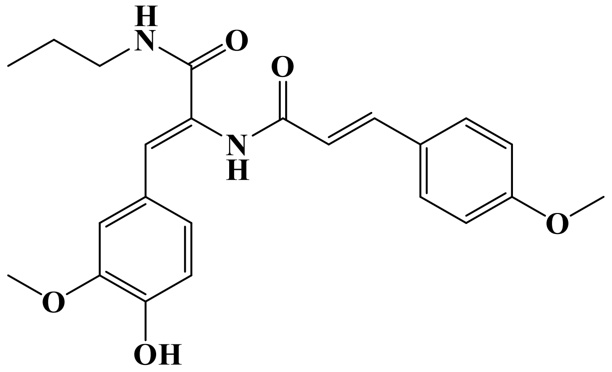

### LC/MS

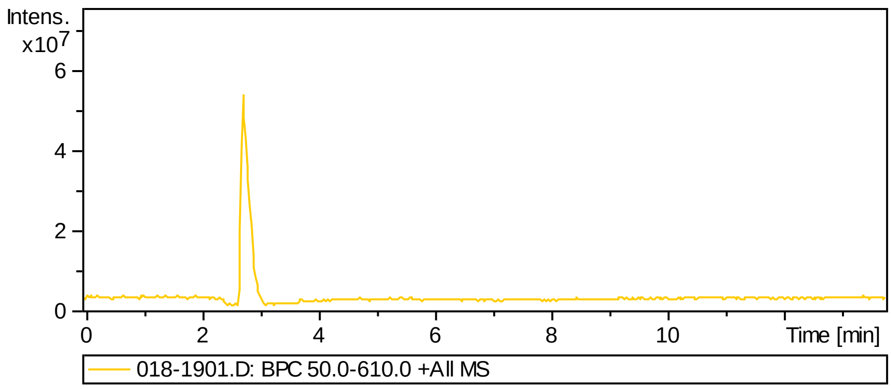


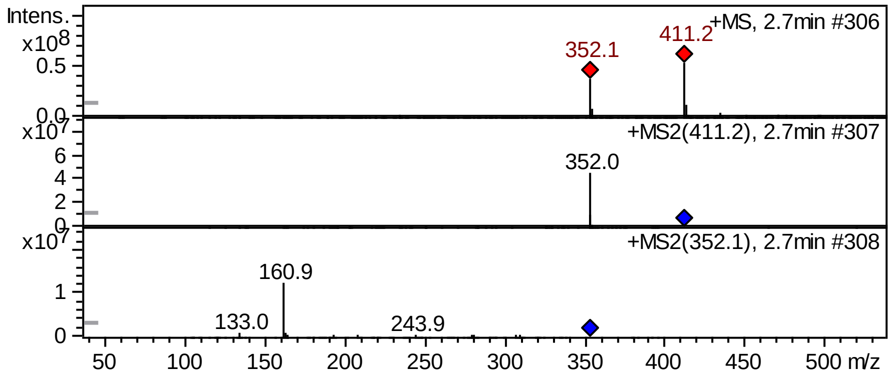


### FT-IR

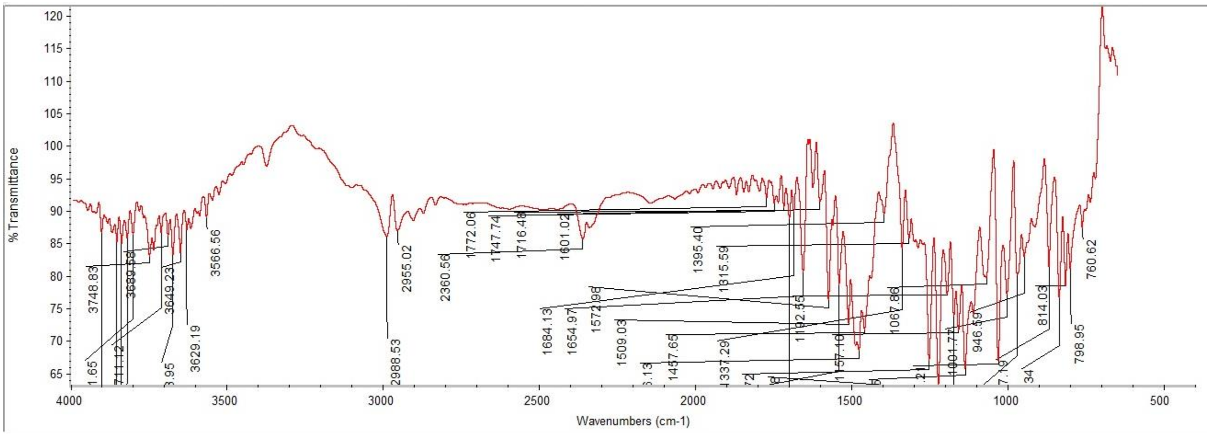

#### $^{1}$H NMR

This report was created by ACD/NMR Processor Academic Edition. For more information go to www.acdlabs.com/nmrproc/

##### RAD-343

11/10/2016 8:44:34 AM
Dr.Mustafa Sample : RAD-343 DMSO PROTON DMSO {D:\Magdy} nmr 19

| Formula $C_{23}H_{26}N_2O_5$ | FW 410.4629 |
|---|---|

| | | | | | | | |
|---|---|---|---|---|---|---|---|
| Acquisition Time (sec) | 2.6564 | Comment | Dr.Mustafa Sample : RAD-343 DMSO PROTON DMSO {D:\Magdy} nmr 19 | | | | |
| Date | 18 Jun 2015 15:11:28 | | | Date Stamp | 18 Jun 2015 15:11:28 | | |
| File Name | E:\Mostafa Alaraby Project\Oxazolone NMR\NMR final oxazolone\MUSTAFA RAD-343 18-06-2015\10\fid | | | | | Frequency (MHz) | 600.15 |
| Nucleus | 1H | Number of Transients | 32 | Origin | spect | Original Points Count | 32768 |
| Owner | nmr | Points Count | 32768 | Pulse Sequence | zg30 | Receiver Gain | 128.00 |
| SW(cyclical) (Hz) | 12335.53 | Solvent | DMSO-d6 | Spectrum Offset (Hz) | 3706.1750 | Spectrum Type | STANDARD |
| Sweep Width (Hz) | 12335.15 | Temperature (degree C) | 25.000 | | | | |

$^{1}$H NMR (600 MHz, DMSO-$d_6$) δ 9.49 (br. s., 1H), 7.97 (t, *J* = 6.02 Hz, 1H), 7.53 - 7.59 (m, 2H), 7.46 (d, *J* = 15.81 Hz, 1H), 7.21 (d, *J* = 1.88 Hz, 1H), 7.00 - 7.03 (m, 3H), 6.99 (dd, *J* = 1.88, 8.28 Hz, 1H), 6.77 (d, *J* = 6.78 Hz, 1H), 6.75 (s, 1H), 3.81 (s, 3H), 3.68 (s, 3H), 3.08 - 3.13 (m, 2H), 1.47 (sxt, *J* = 7.30 Hz, 2H), 0.85 - 0.88 (m, 3H)

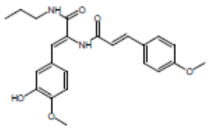


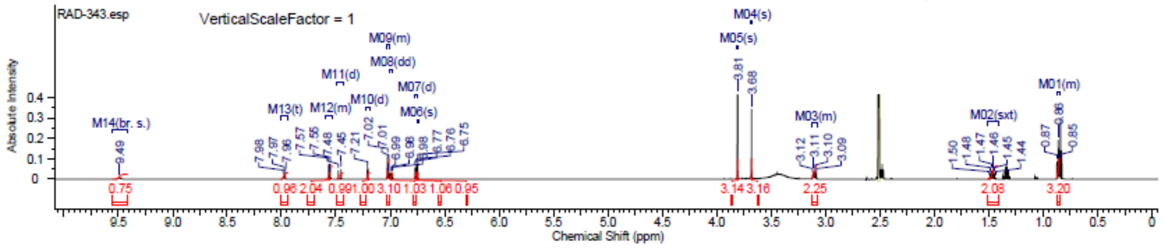

### Spectra of (3512)

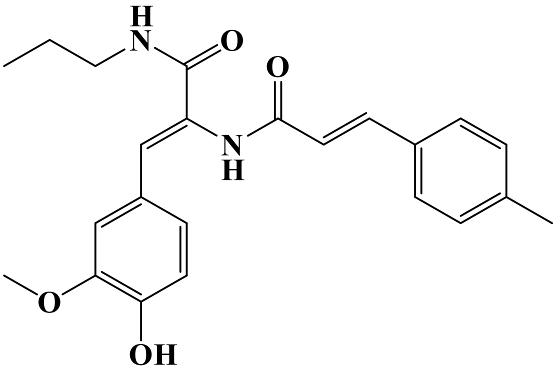


## LC/MS

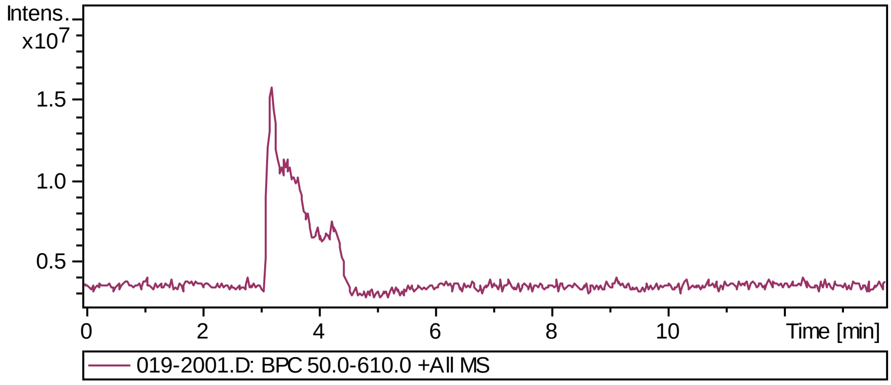


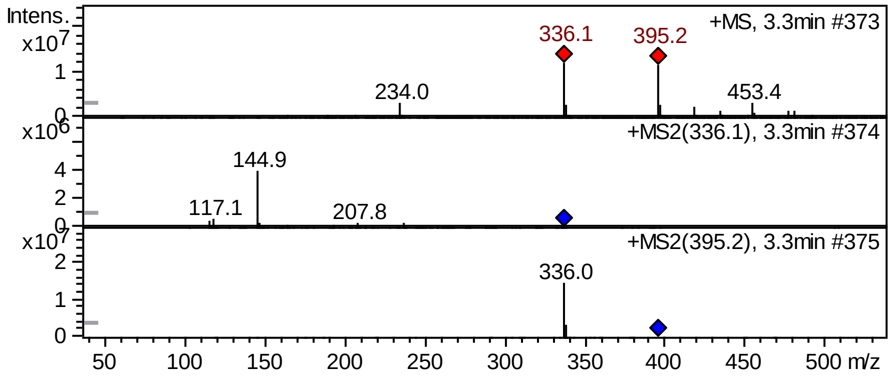


## FT-IR

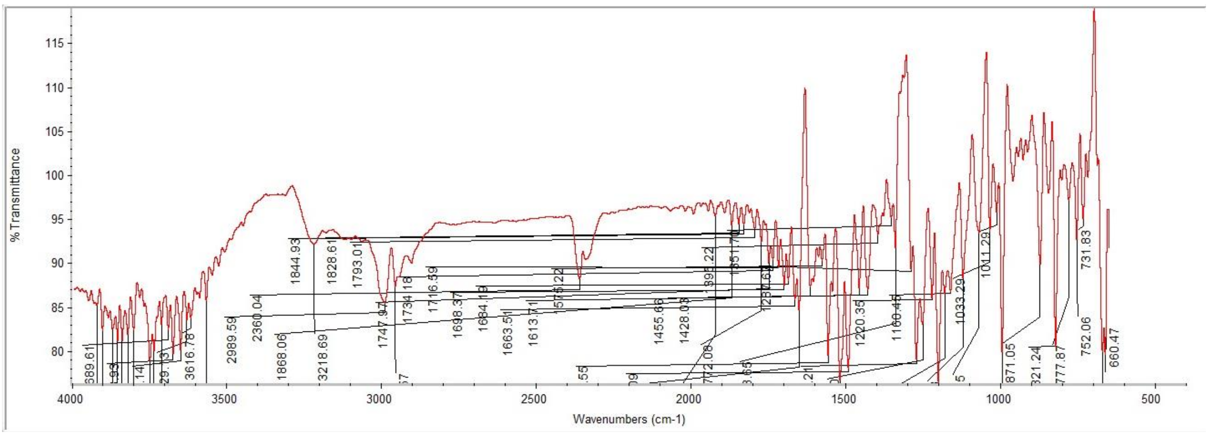

### Supplementary Information

#### [1]H NMR

This report was created by ACD/NMR Processor Academic Edition. For more information go to www.acdlabs.com/nmrproc/

**RAD-344**

08/11/2016 9:06:10 AM
Dr.Mustafa Sample : RAD-344 DMSO PROTON DMSO {D:\Magdy} nmr 10

| Formula $C_{23}H_{26}N_2O_4$ | FW 394.4635 |
|---|---|

| | | | | | | | |
|---|---|---|---|---|---|---|---|
| *Acquisition Time (sec)* | 2.6564 | *Comment* | Dr.Mustafa Sample : RAD-344 DMSO PROTON DMSO {D:\Magdy} nmr 10 | | | | |
| *Date* | 18 Jun 2015 14:24:32 | | | *Date Stamp* | 18 Jun 2015 14:24:32 | | |
| *File Name* | E:\Mostafa Alaraby Project\Oxazolone NMR\NMR final oxazolone\MUSTAFA RAD-344 18-06-2015\10\fid | | | | | *Frequency (MHz)* | 600.15 |
| *Nucleus* | 1H | *Number of Transients* | 32 | *Origin* | spect | *Original Points Count* | 32768 |
| *Owner* | nmr | *Points Count* | 32768 | *Pulse Sequence* | zg30 | *Receiver Gain* | 144.00 |
| *SW(cyclical) (Hz)* | 12335.53 | *Solvent* | DMSO-d6 | *Spectrum Offset (Hz)* | 3706.1750 | *Spectrum Type* | STANDARD |
| *Sweep Width (Hz)* | 12335.15 | *Temperature (degree C)* | 25.000 | | | | |

[1]H NMR (600 MHz, DMSO-$d_6$) δ 9.54 (s, 1H), 9.39 (br. s., 1H), 8.00 (t, $J$ = 5.65 Hz, 1H), 7.44 - 7.53 (m, 3H), 7.27 (d, $J$ = 7.91 Hz, 2H), 7.21 (s, 1H), 7.03 (s, 1H), 6.99 (dd, $J$ = 1.88, 8.28 Hz, 1H), 6.85 (d, $J$ = 15.81 Hz, 1H), 6.76 (d, $J$ = 8.28 Hz, 1H), 3.68 (s, 3H), 3.10 (q, $J$ = 6.65 Hz, 2H), 2.35 (s, 3H), 1.47 (sxt, $J$ = 7.23 Hz, 2H), 0.86 (t, $J$ = 7.34 Hz, 3H)

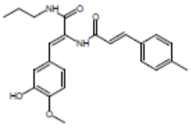


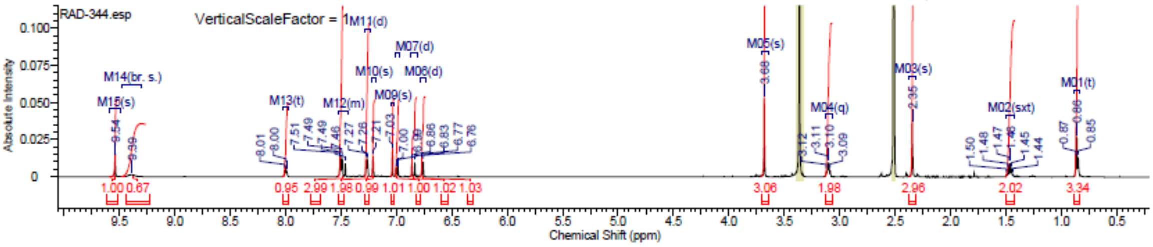


#### [13]C NMR

This report was created by ACD/NMR Processor Academic Edition. For more information go to www.acdlabs.com/nmrproc/

**RAD-344**

20/09/2016 9:31:55 AM
Dr.Mostafa Sample ; RAD-344 DMSO

| Formula $C_{23}H_{26}N_2O_4$ | FW 394.4635 |
|---|---|

| | | | | | | | |
|---|---|---|---|---|---|---|---|
| *Acquisition Time (sec)* | 0.6423 | *Comment* | Dr.Mostafa Sample ; RAD-344 DMSO | | | *Date* | 20 Apr 2016 20:06:08 |
| *Date Stamp* | 20 Apr 2016 20:06:08 | | | | | | |
| *File Name* | E:\Projects\Mostafa Alaraby Project\Oxazolone\Oxazolone NMR\NMR final oxazolone\13 CNMR New 18-4-2016\MUSTAFA RAD-344 19-04-2016\140\fid | | | | | | |
| *Frequency (MHz)* | 213.77 | *Nucleus* | 13C | *Number of Transients* | 3500 | *Origin* | spect |
| *Original Points Count* | 32768 | *Owner* | nmr | *Points Count* | 32768 | *Pulse Sequence* | zgpg30 |
| *Receiver Gain* | 186.93 | *SW(cyclical) (Hz)* | 51020.41 | *Solvent* | DMSO-d6 | *Spectrum Offset (Hz)* | 21290.6973 |
| *Spectrum Type* | STANDARD | *Sweep Width (Hz)* | 51018.85 | *Temperature (degree C)* | 25.000 | | |

[13]C NMR (214 MHz, DMSO-$d_6$) δ 165.1, 164.9, 147.5, 147.3, 139.7, 139.6, 132.1, 129.7, 128.5, 127.7, 127.3, 125.5, 123.7, 120.8, 115.4, 112.9, 55.3, 41.0, 39.8, 39.7, 39.6, 39.4, 39.3, 39.2, 22.5, 11.5

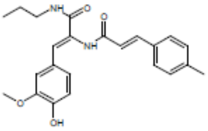


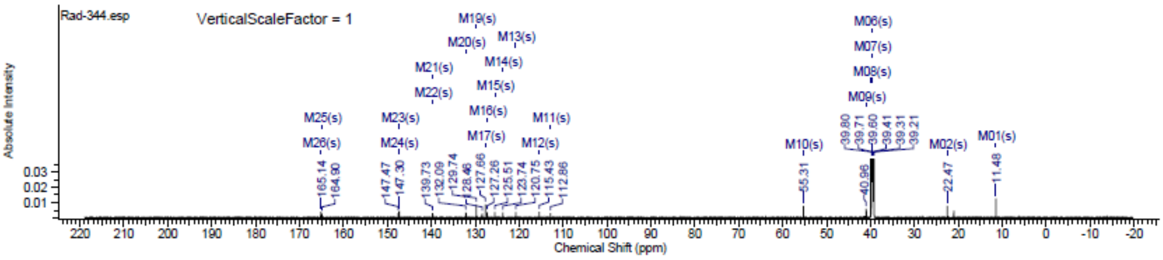

#### Spectra of (3612)

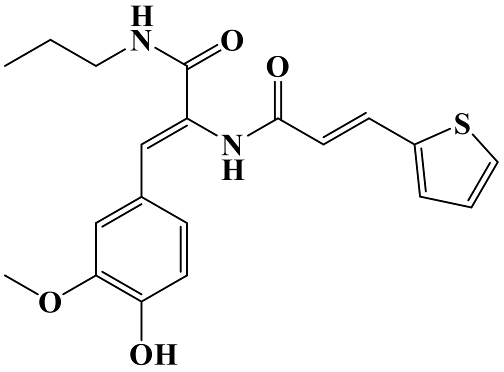


### LC/MS

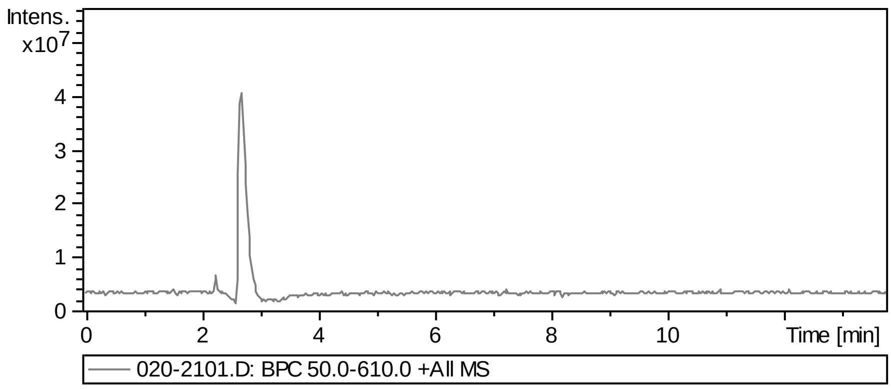


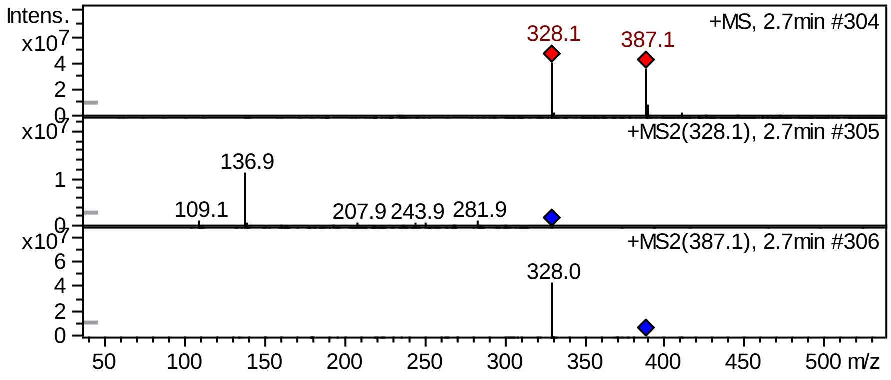


### FT-IR

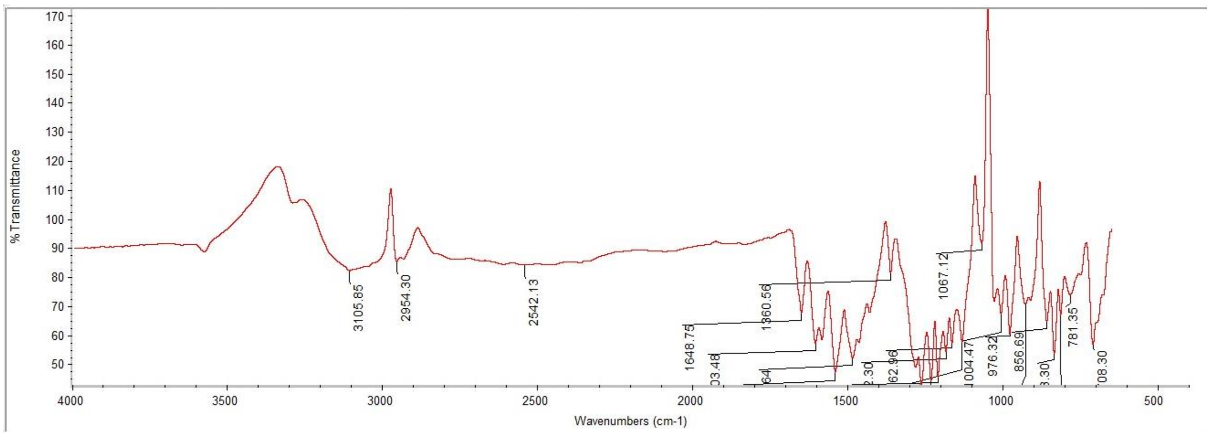

#### $^{1}$H NMR

This report was created by ACD/NMR Processor Academic Edition. For more information go to www.acdlabs.com/nmrproc/

##### RAD-345

11/10/2016 8:53:39 AM
Dr.Mustafa Sample : RAD-345 DMSO PROTON DMSO {D:\Magdy} nmr 14

| Formula | $C_{20}H_{22}N_2O_4S$ | FW | 386.4647 |
|---|---|---|---|

| | | | | | | | |
|---|---|---|---|---|---|---|---|
| Acquisition Time (sec) | 2.6564 | Comment | Dr.Mustafa Sample : RAD-345 DMSO PROTON DMSO {D:\Magdy} nmr 14 | | | | |
| Date | 18 Jun 2015 14:45:52 | | | Date Stamp | 18 Jun 2015 14:45:52 | | |
| File Name | E:\Mostafa Alaraby Project\Oxazolone NMR\NMR final oxazolone\MUSTAFA RAD-345 18-06-2015\10\fid | | | | | Frequency (MHz) | 600.15 |
| Nucleus | 1H | Number of Transients | 32 | Origin | spect | Original Points Count | 32768 |
| Owner | nmr | Points Count | 32768 | Pulse Sequence | zg30 | Receiver Gain | 144.00 |
| SW(cyclical) (Hz) | 12335.53 | Solvent | DMSO-d6 | Spectrum Offset (Hz) | 3706.1750 | Spectrum Type | STANDARD |
| Sweep Width (Hz) | 12335.15 | Temperature (degree C) | 25.000 | | | | |

$^{1}$H NMR (600 MHz, DMSO-$d_6$) δ 9.54 (br. s., 1H), 7.99 (t, $J$ = 5.83 Hz, 1H), 7.63 - 7.68 (m, 2H), 7.43 (d, $J$ = 3.39 Hz, 1H), 7.19 (d, $J$ = 1.88 Hz, 1H), 7.13 - 7.16 (m, 1H), 7.04 (s, 1H), 6.98 (dd, $J$ = 1.88, 8.28 Hz, 1H), 6.76 (d, $J$ = 7.91 Hz, 1H), 6.66 (d, $J$ = 15.81 Hz, 1H), 3.69 (s, 3H), 3.08 - 3.12 (m, 2H), 1.46 (sxt, $J$ = 7.23 Hz, 2H), 0.84 - 0.87 (m, 3H)

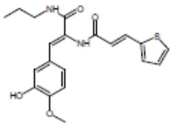


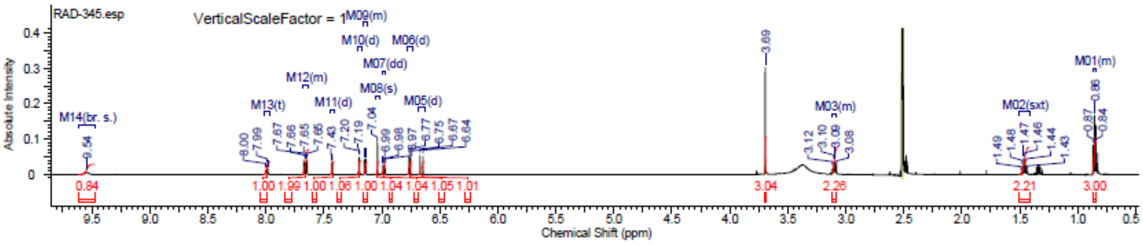

### Spectra of (3712)

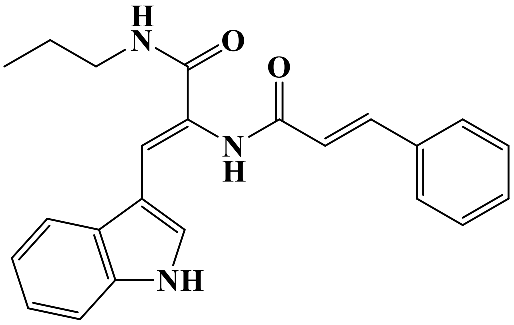


## LC/MS

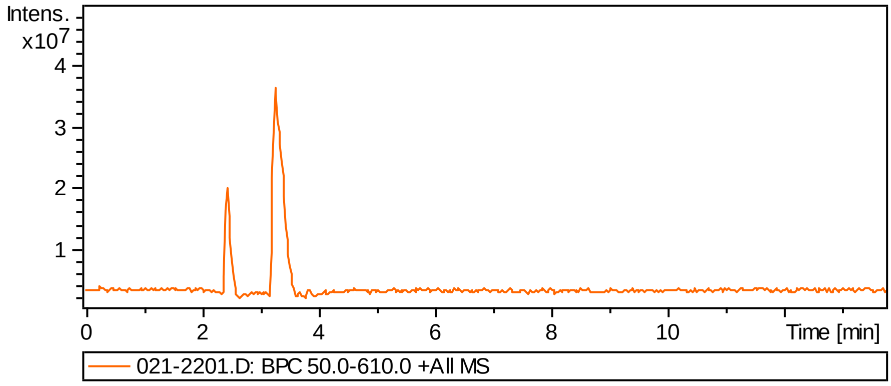


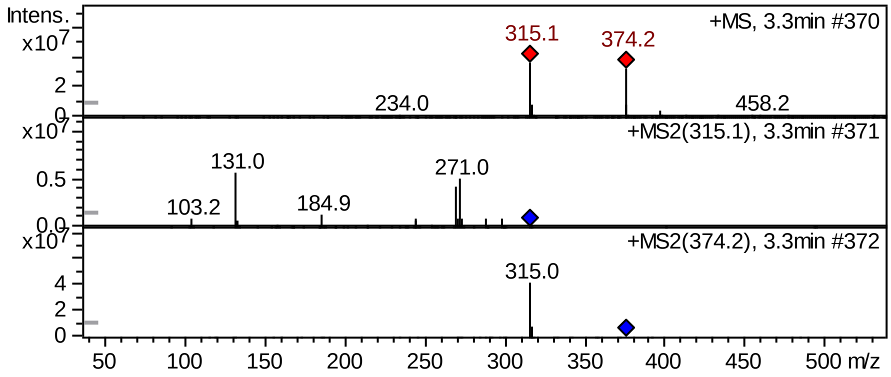


## FT-IR

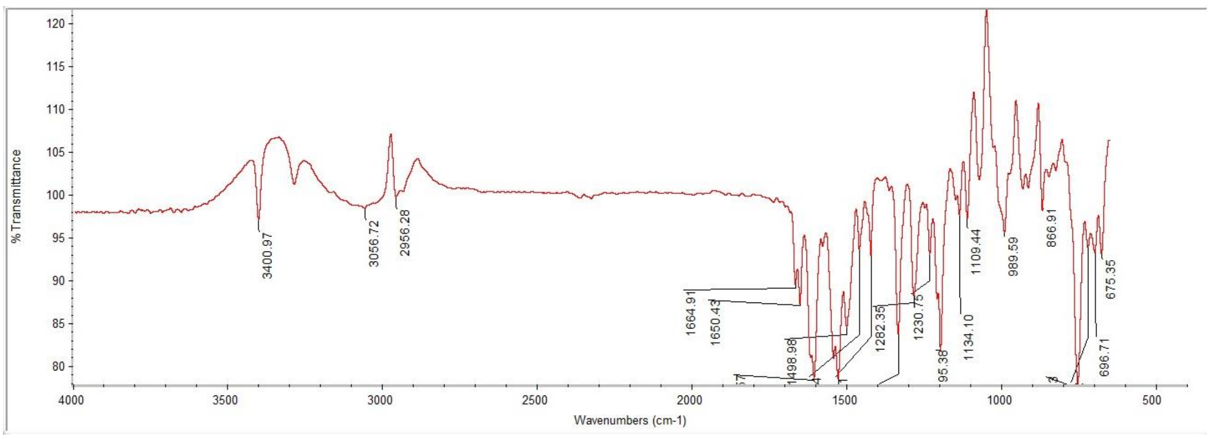

#### $^1$H NMR

This report was created by ACD/NMR Processor Academic Edition. For more information go to www.acdlabs.com/nmrproc/

##### RAD-351

08/11/2016 11:46:25 AM
Dr.Mustafa Sample : RAD-351 DMSO PROTON DMSO {D:\Magdy} nmr 30

| Formula $C_{23}H_{23}N_3O_2$ | FW 373.4476 |
|---|---|

| | | | | | | | |
|---|---|---|---|---|---|---|---|
| Acquisition Time (sec) | 2.6564 | Comment | Dr.Mustafa Sample : RAD-351 DMSO PROTON DMSO {D:\Magdy} nmr 30 | | | | |
| Date | 18 Jun 2015 16:09:04 | | | Date Stamp | 18 Jun 2015 16:09:04 | | |
| File Name | E:\Mostafa Alaraby Project\Oxazolone NMR\NMR final oxazolone\MUSTAFA RAD-351 18-06-2015\10\fid | | | | | Frequency (MHz) | 600.15 |
| Nucleus | 1H | Number of Transients | 32 | Origin | spect | Original Points Count | 32768 |
| Owner | nmr | Points Count | 32768 | Pulse Sequence | zg30 | Receiver Gain | 144.00 |
| SW(cyclical) (Hz) | 12335.53 | Solvent | DMSO-d6 | Spectrum Offset (Hz) | 3706.1750 | Spectrum Type | STANDARD |
| Sweep Width (Hz) | 12335.15 | Temperature (degree C) | 25.000 | | | | |

$^1$H NMR (600 MHz, DMSO-$d_6$) δ 11.56 (br. s., 1H), 9.46 (s, 1H), 7.98 (t, $J$ = 5.83 Hz, 1H), 7.74 (d, $J$ = 7.91 Hz, 1H), 7.63 - 7.67 (m, 3H), 7.42 - 7.55 (m, 5H), 7.17 (t, $J$ = 7.53 Hz, 1H), 7.12 (t, $J$ = 7.53 Hz, 1H), 6.98 (d, $J$ = 15.81 Hz, 1H), 3.14 (q, $J$ = 6.53 Hz, 2H), 1.50 (sxt, $J$ = 7.23 Hz, 2H), 0.88 (t, $J$ = 7.34 Hz, 3H)

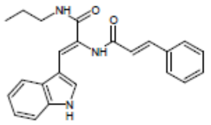


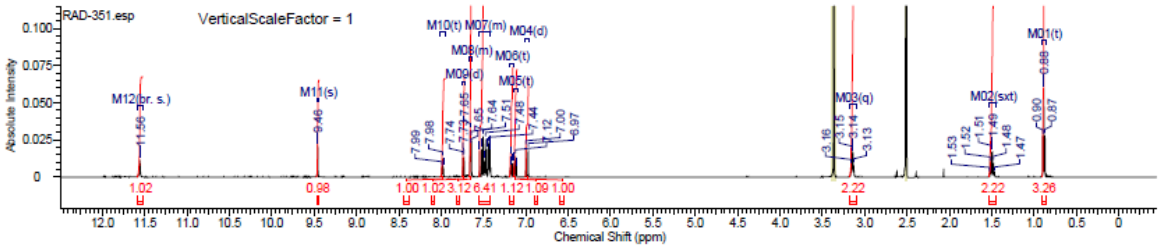


#### $^{13}$C NMR

This report was created by ACD/NMR Processor Academic Edition. For more information go to www.acdlabs.com/nmrproc/

##### RAD-351

25/09/2016 3:26:58 PM
Dr.Moustafa Sample : RAD-351 DMSO

| Formula $C_{23}H_{23}N_3O_2$ | FW 373.4476 |
|---|---|

| | | | | | | | |
|---|---|---|---|---|---|---|---|
| Acquisition Time (sec) | 0.6423 | Comment | Dr.Moustafa Sample : RAD-351 DMSO | | | Date | 22 Apr 2016 15:05:20 |
| Date Stamp | 22 Apr 2016 15:05:20 | | | | | | |
| File Name | E:\Projects\Mostafa Alaraby Project\Oxazolone\Oxazolone NMR\NMR final oxazolone\13 CNMR New 18-4-2016\MUSTAFA RAD-351 21-04-2016\30\fid | | | | | | |
| Frequency (MHz) | 213.77 | Nucleus | 13C | Number of Transients | 12288 | Origin | spect |
| Original Points Count | 32768 | Owner | nmr | Points Count | 32768 | Pulse Sequence | zgpg30 |
| Receiver Gain | 186.93 | SW(cyclical) (Hz) | 51020.41 | Solvent | DMSO-d6 | Spectrum Offset (Hz) | 21293.8125 |
| Spectrum Type | STANDARD | Sweep Width (Hz) | 51018.85 | Temperature (degree C) | 25.000 | | |

$^{13}$C NMR (214 MHz, DMSO-$d_6$) δ 164.5, 164.2, 141.5, 135.5, 134.8, 129.6, 129.3, 129.1, 129.0, 128.7, 127.9, 124.9, 124.6, 122.1, 120.0, 118.2, 111.9, 110.0, 41.0, 39.8, 39.7, 39.6, 39.4, 39.3, 39.2, 22.3, 11.5

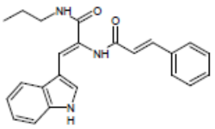


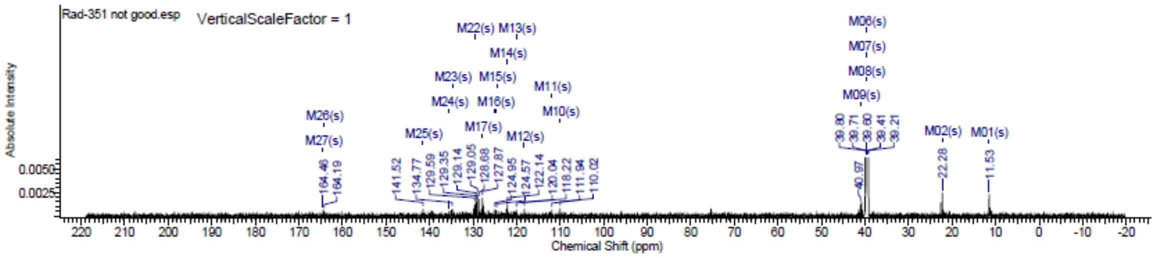

#### Spectra of (3812)

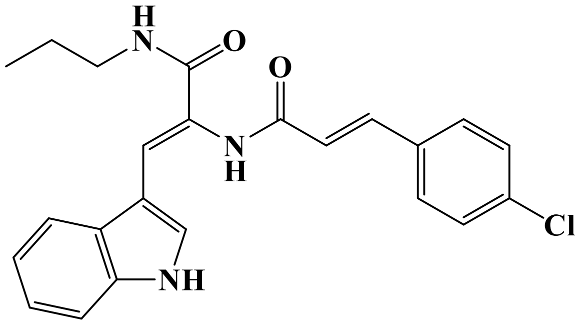

### LC/MS

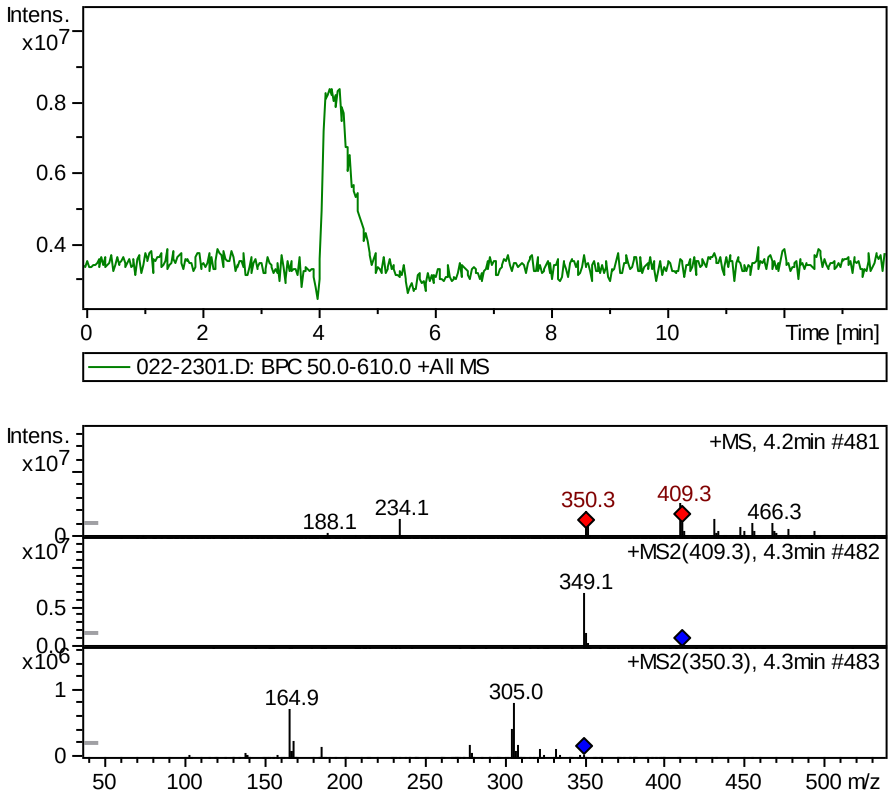


### FT-IR

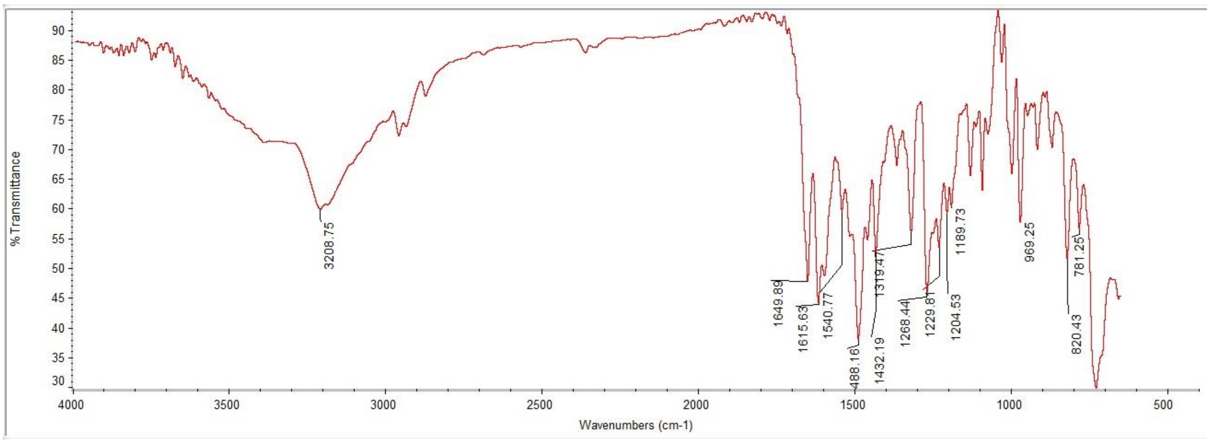

#### [1]H NMR

This report was created by ACD/NMR Processor Academic Edition. For more information go to www.acdlabs.com/nmrproc/

##### RAD-352

08/11/2016 11:51:59 AM
Dr.Mustafa Sample : RAD-352 DMSO PROTON DMSO {D:\Magdy} nmr 35

| Formula | $C_{23}H_{22}ClN_3O_2$ | FW | 407.8927 |
|---|---|---|---|

| Acquisition Time (sec) | 2.6564 | Comment | Dr.Mustafa Sample : RAD-352 DMSO PROTON DMSO {D:\Magdy} nmr 35 | | | | |
|---|---|---|---|---|---|---|---|
| Date | 18 Jun 2015 16:34:40 | | | Date Stamp | 18 Jun 2015 16:34:40 | | |
| File Name | E:\Mostafa Alaraby Project\Oxazolone NMR\NMR final oxazolone\MUSTAFA RAD-352 18-06-2015\10\fid | | | | | Frequency (MHz) | 600.15 |
| Nucleus | 1H | Number of Transients | 32 | Origin | spect | Original Points Count | 32768 |
| Owner | nmr | Points Count | 32768 | Pulse Sequence | zg30 | Receiver Gain | 144.00 |
| SW(cyclical) (Hz) | 12335.53 | Solvent | DMSO-d6 | Spectrum Offset (Hz) | 3706.1750 | Spectrum Type | STANDARD |
| Sweep Width (Hz) | 12335.15 | Temperature (degree C) | 25.000 | | | | |

[1]H NMR (600 MHz, DMSO-$d_6$) δ 11.56 (br. s., 1H), 9.47 (s, 1H), 7.98 (t, $J$ = 5.83 Hz, 1H), 7.73 (d, $J$ = 7.91 Hz, 1H), 7.68 (d, $J$ = 8.66 Hz, 2H), 7.65 (d, $J$ = 2.63 Hz, 1H), 7.50 - 7.56 (m, 3H), 7.43 (d, $J$ = 7.91 Hz, 1H), 7.17 (t, $J$ = 7.15 Hz, 1H), 7.12 (t, $J$ = 7.34 Hz, 1H), 6.98 (d, $J$ = 15.81 Hz, 1H), 3.14 (q, $J$ = 6.53 Hz, 2H), 1.50 (sxt, $J$ = 7.30 Hz, 2H), 0.88 (t, $J$ = 7.53 Hz, 3H)

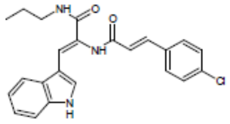


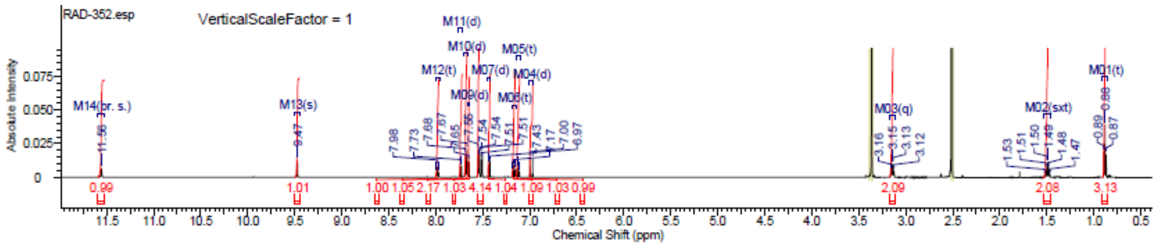


#### [13]C NMR

This report was created by ACD/NMR Processor Academic Edition. For more information go to www.acdlabs.com/nmrproc/

##### RAD-352

25/09/2016 3:33:14 PM
Dr.Moustafa Sample : RAD-352 DMSO

| Formula | $C_{23}H_{22}ClN_3O_2$ | FW | 407.8927 |
|---|---|---|---|

| Acquisition Time (sec) | 0.6423 | Comment | Dr.Moustafa Sample : RAD-352 DMSO | | | Date | 21 Apr 2016 11:25:36 |
|---|---|---|---|---|---|---|---|
| Date Stamp | 21 Apr 2016 11:25:36 | | | | | | |
| File Name | E:\Projects\Mostafa Alaraby Project\Oxazolone\Oxazolone NMR\NMR final oxazolone\13 CNMR New 18-4-2016\MUSTAFA RAD-352 21-04-2016\10\fid | | | | | | |
| Frequency (MHz) | 213.77 | Nucleus | 13C | Number of Transients | 12288 | Origin | spect |
| Original Points Count | 32768 | Owner | nmr | Points Count | 32768 | Pulse Sequence | zgpg30 |
| Receiver Gain | 186.93 | SW(cyclical) (Hz) | 51020.41 | Solvent | DMSO-d6 | Spectrum Offset (Hz) | 21516.4648 |
| Spectrum Type | STANDARD | Sweep Width (Hz) | 51018.85 | Temperature (degree C) | 24.999 | | |

[13]C NMR (214 MHz, DMSO-$d_6$) δ 165.7, 165.3, 140.5, 138.1, 135.7, 134.2, 132.3, 132.2, 130.7, 130.6, 130.5, 130.1, 129.8, 125.1, 123.9, 122.7, 113.7, 111.5, 110.1, 41.6, 39.8, 39.7, 39.6, 39.4, 39.3, 39.2, 23.0, 11.6

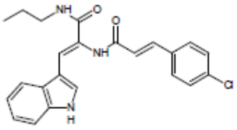


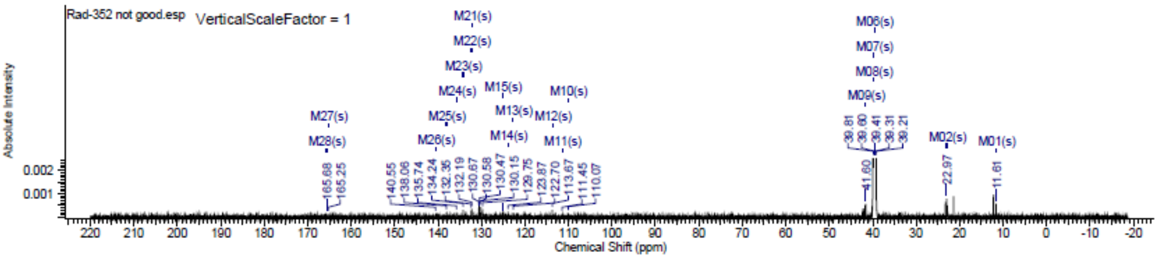

### Spectra of (3912)

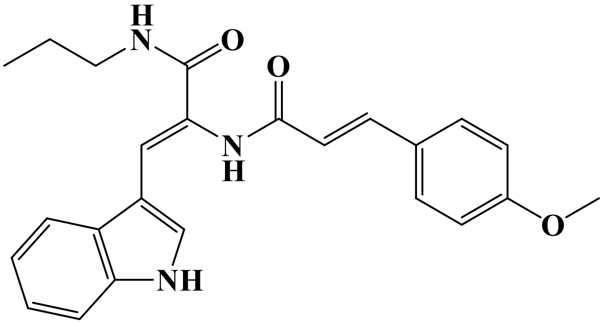


## LC/MS

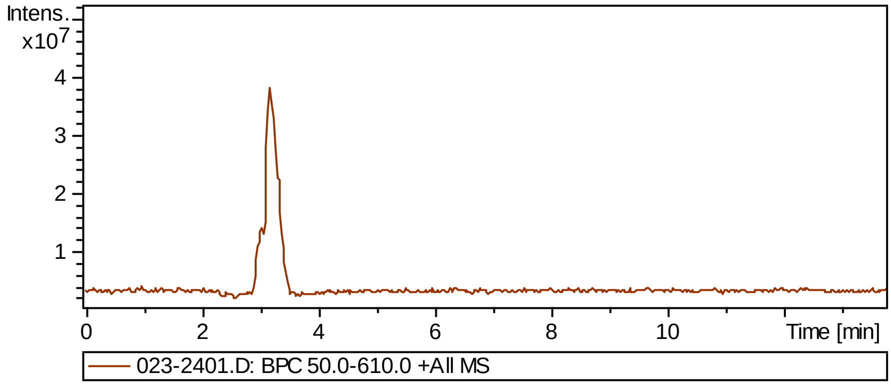


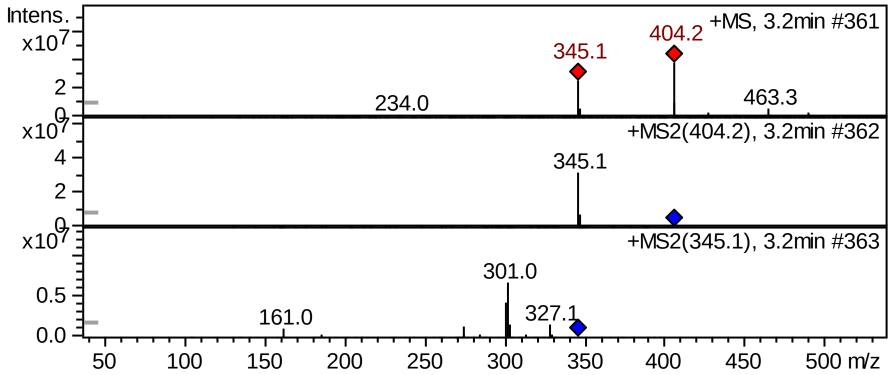


## FT-IR

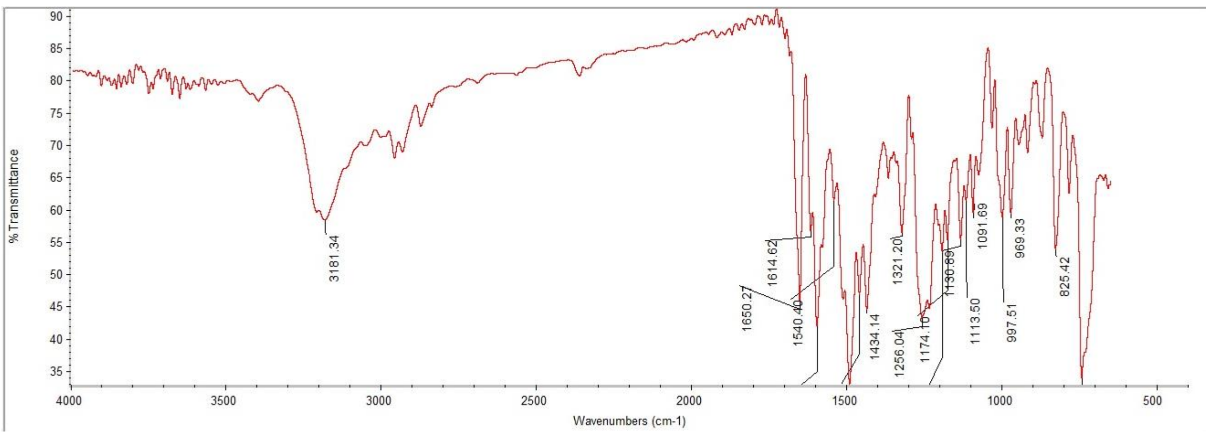

### Supplementary Information

#### $^{1}$H NMR

This report was created by ACD/NMR Processor Academic Edition. For more information go to www.acdlabs.com/nmrproc/

##### RAD-353

09/11/2016 7:58:16 AM
Dr.Mustafa Sample : RAD-353 DMSO PROTON DMSO {D:\Magdy} nmr 28

| Formula $C_{24}H_{25}N_3O_3$ | FW 403.4736 |
|---|---|

| | | | | | | | |
|---|---|---|---|---|---|---|---|
| *Acquisition Time (sec)* | 2.6564 | *Comment* | Dr.Mustafa Sample : RAD-353 DMSO PROTON DMSO {D:\Magdy} nmr 28 | | | | |
| *Date* | 18 Jun 2015 15:58:24 | | | *Date Stamp* | 18 Jun 2015 15:58:24 | | |
| *File Name* | E:\Mostafa Alaraby Project\Oxazolone NMR\NMR final oxazolone\MUSTAFA RAD-353 18-06-2015\10\fid | | | | | *Frequency (MHz)* | 600.15 |
| *Nucleus* | 1H | *Number of Transients* | 32 | *Origin* | spect | *Original Points Count* | 32768 |
| *Owner* | nmr | *Points Count* | 32768 | *Pulse Sequence* | zg30 | *Receiver Gain* | 144.00 |
| *SW(cyclical) (Hz)* | 12335.53 | *Solvent* | DMSO-d6 | *Spectrum Offset (Hz)* | 3706.1750 | *Spectrum Type* | STANDARD |
| *Sweep Width (Hz)* | 12335.15 | *Temperature (degree C)* | 25.000 | | | | |

$^{1}$H NMR (600 MHz, DMSO-$d_6$) δ 11.55 (br. s., 1H), 9.36 (s, 1H), 7.95 (t, *J* = 5.83 Hz, 1H), 7.73 (d, *J* = 7.91 Hz, 1H), 7.64 (d, *J* = 2.26 Hz, 1H), 7.60 (d, *J* = 8.66 Hz, 2H), 7.46 - 7.50 (m, 2H), 7.43 (d, *J* = 7.91 Hz, 1H), 7.17 (t, *J* = 7.53 Hz, 1H), 7.12 (t, *J* = 7.53 Hz, 1H), 7.03 (d, *J* = 8.66 Hz, 2H), 6.84 (d, *J* = 15.81 Hz, 1H), 3.82 (s, 3H), 3.14 (q, *J* = 6.27 Hz, 2H), 1.49 (sxt, *J* = 7.23 Hz, 2H), 0.88 (t, *J* = 7.34 Hz, 3H)

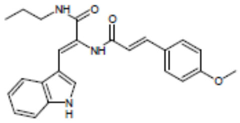


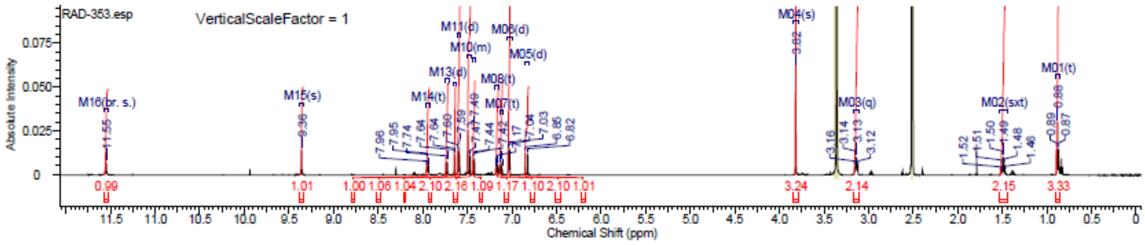


#### $^{13}$C NMR

This report was created by ACD/NMR Processor Academic Edition. For more information go to www.acdlabs.com/nmrproc/

##### RAD-353

25/09/2016 3:40:24 PM
Dr.Moustafa Sample : RAD-353 DMSO

| Formula $C_{24}H_{25}N_3O_3$ | FW 403.4736 |
|---|---|

| | | | | | | | |
|---|---|---|---|---|---|---|---|
| *Acquisition Time (sec)* | 0.6423 | *Comment* | Dr.Moustafa Sample : RAD-353 DMSO | | | *Date* | 23 Apr 2016 09:43:12 |
| *Date Stamp* | 23 Apr 2016 09:43:12 | | | | | | |
| *File Name* | E:\Projects\Mostafa Alaraby Project\Oxazolone\Oxazolone NMR\NMR final oxazolone\13 CNMR New 18-4-2016\MUSTAFA RAD-353 21-04-2016\50\fid | | | | | | |
| *Frequency (MHz)* | 213.77 | *Nucleus* | 13C | *Number of Transients* | 12288 | *Origin* | spect |
| *Original Points Count* | 32768 | *Owner* | nmr | *Points Count* | 32768 | *Pulse Sequence* | zgpg30 |
| *Receiver Gain* | 186.93 | *SW(cyclical) (Hz)* | 51020.41 | *Solvent* | DMSO-d6 | *Spectrum Offset (Hz)* | 21287.5840 |
| *Spectrum Type* | STANDARD | *Sweep Width (Hz)* | 51018.85 | *Temperature (degree C)* | 25.002 | | |

$^{13}$C NMR (214 MHz, DMSO-$d_6$) δ 166.0, 164.3, 160.6, 141.5, 138.5, 137.1, 132.3, 131.9, 131.4, 129.2, 123.5, 122.2, 121.0, 120.6, 114.6, 113.9, 112.5, 112.3, 55.8, 41.5, 39.8, 39.7, 39.6, 39.4, 39.3, 39.2, 22.6, 11.5

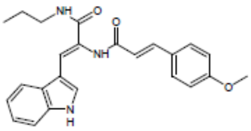


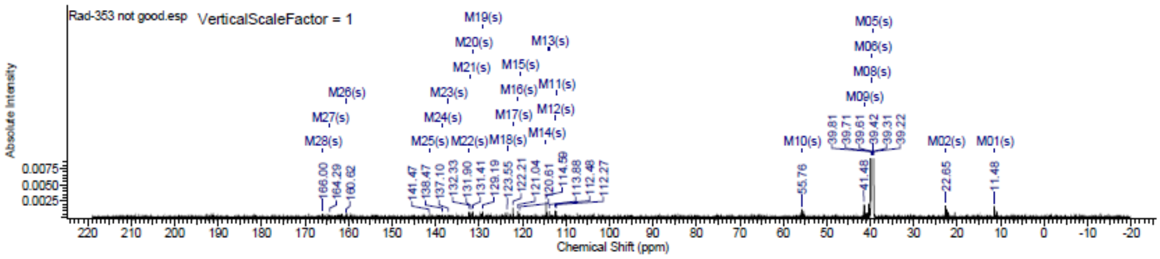

### Spectra of (4012)

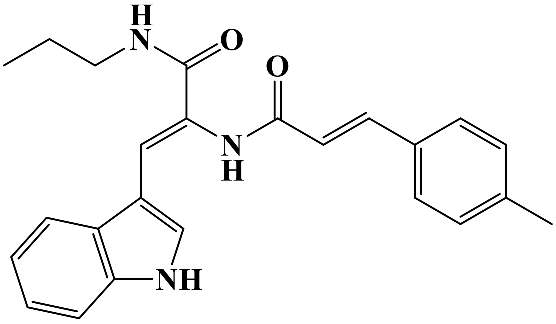


## LC/MS

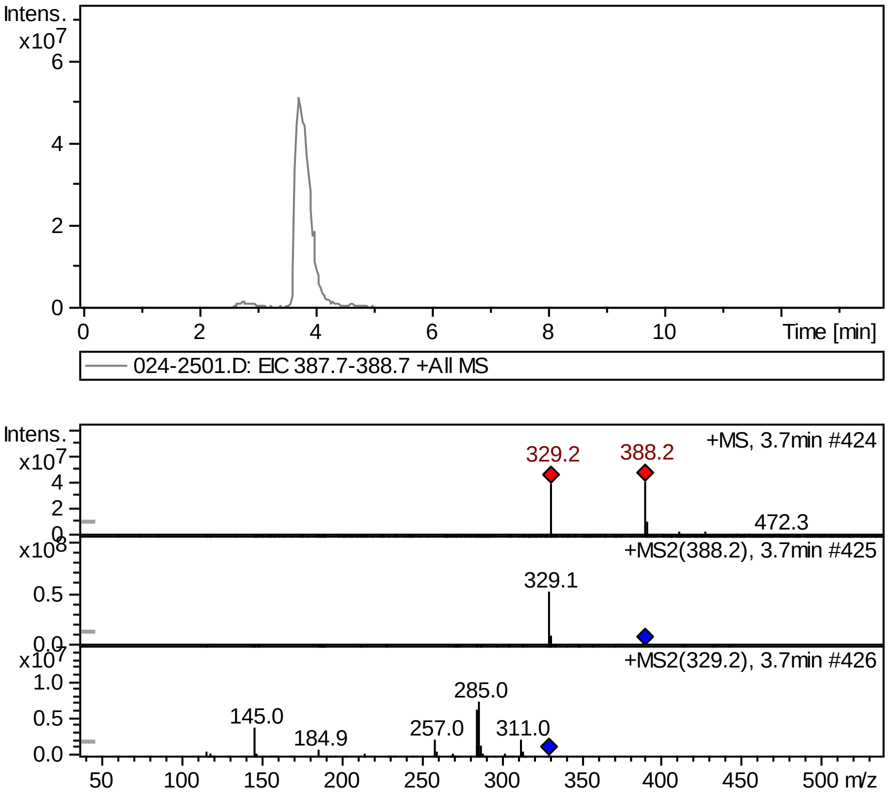


## FT-IR

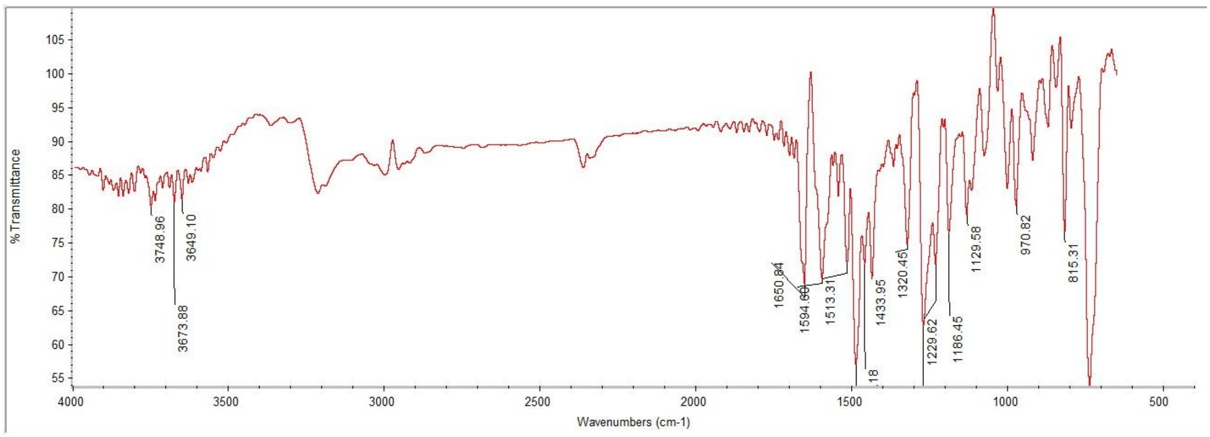

#### $^{1}$H NMR

This report was created by ACD/NMR Processor Academic Edition. For more information go to www.acdlabs.com/nmrproc/

**RAD-354**

09/11/2016 8:10:30 AM
Dr.Mustafa Sample : RAD-354 DMSO PROTON DMSO {D:\Magdy} nmr 29

| Formula $C_{24}H_{25}N_3O_2$ | FW 387.4742 |
|---|---|

| | | | | | | | |
|---|---|---|---|---|---|---|---|
| *Acquisition Time (sec)* | 2.6564 | *Comment* | Dr.Mustafa Sample : RAD-354 DMSO PROTON DMSO {D:\Magdy} nmr 29 | | | | |
| *Date* | 18 Jun 2015 16:04:48 | | | *Date Stamp* | 18 Jun 2015 16:04:48 | | |
| *File Name* | E:\Mostafa Alaraby Project\Oxazolone NMR\NMR final oxazolone\MUSTAFA RAD-354 18-06-2015\10\fid | | | | | *Frequency (MHz)* | 600.15 |
| *Nucleus* | 1H | *Number of Transients* | 32 | *Origin* | spect | *Original Points Count* | 32768 |
| *Owner* | nmr | *Points Count* | 32768 | *Pulse Sequence* | zg30 | *Receiver Gain* | 114.00 |
| *SW(cyclical) (Hz)* | 12335.53 | *Solvent* | DMSO-d6 | *Spectrum Offset (Hz)* | 3706.1750 | *Spectrum Type* | STANDARD |
| *Sweep Width (Hz)* | 12335.15 | *Temperature (degree C)* | 25.000 | | | | |

$^{1}$H NMR (600 MHz, DMSO-$d_6$) δ 11.56 (br. s., 1H), 9.42 (s, 1H), 7.97 (t, $J$ = 5.83 Hz, 1H), 7.73 (d, $J$ = 7.91 Hz, 1H), 7.65 (s, 1H), 7.52 - 7.56 (m, $J$ = 7.91 Hz, 2H), 7.47 - 7.52 (m, 2H), 7.43 (d, $J$ = 7.91 Hz, 1H), 7.26 - 7.30 (m, $J$ = 7.91 Hz, 2H), 7.15 - 7.18 (m, 1H), 7.11 - 7.14 (m, 1H), 6.93 (d, $J$ = 15.81 Hz, 1H), 3.14 (q, $J$ = 6.40 Hz, 2H), 2.36 (s, 3H), 1.50 (sxt, $J$ = 7.30 Hz, 2H), 0.88 (t, $J$ = 7.53 Hz, 3H)

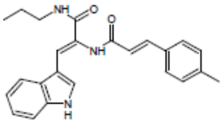


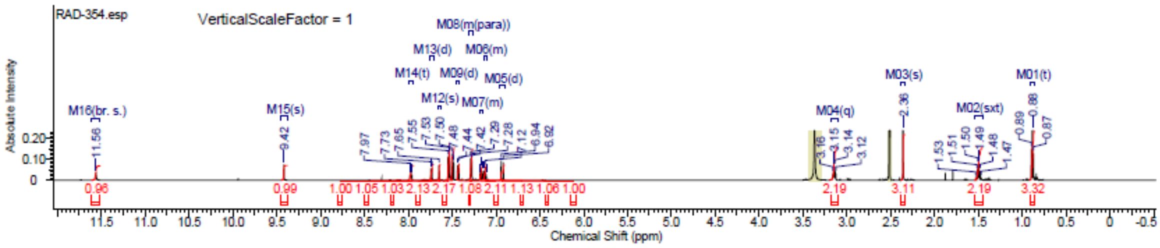


#### $^{13}$C NMR

This report was created by ACD/NMR Processor Academic Edition. For more information go to www.acdlabs.com/nmrproc/

**RAD-354**

25/09/2016 3:51:57 PM
Dr.Mostafa Sample ; RAD-354 DMSO

| Formula $C_{24}H_{25}N_3O_2$ | FW 387.4742 |
|---|---|

| | | | | | | | |
|---|---|---|---|---|---|---|---|
| *Acquisition Time (sec)* | 0.6423 | *Comment* | Dr.Mostafa Sample ; RAD-354 DMSO | | | *Date* | 19 Apr 2016 17:45:20 |
| *Date Stamp* | 19 Apr 2016 17:45:20 | | | | | | |
| *File Name* | E:\Projects\Mostafa Alaraby Project\Oxazolone\Oxazolone NMR\NMR final oxazolone\13 CNMR New 18-4-2016\MUSTAFA RAD-354 19-04-2016\20\fid | | | | | | |
| *Frequency (MHz)* | 213.77 | *Nucleus* | 13C | *Number of Transients* | 3500 | *Origin* | spect |
| *Original Points Count* | 32768 | *Owner* | nmr | *Points Count* | 32768 | *Pulse Sequence* | zgpg30 |
| *Receiver Gain* | 186.93 | *SW(cyclical) (Hz)* | 51020.41 | *Solvent* | DMSO-d6 | *Spectrum Offset (Hz)* | 21289.1406 |
| *Spectrum Type* | STANDARD | *Sweep Width (Hz)* | 51018.85 | *Temperature (degree C)* | 25.000 | | |

$^{13}$C NMR (214 MHz, DMSO-$d_6$) δ 165.1, 164.6, 141.5, 139.6, 135.5, 129.8, 129.6, 129.2, 127.9, 127.7, 127.6, 122.2, 121.3, 121.1, 118.2, 112.5, 111.9, 110.0, 40.9, 39.8, 39.7, 39.6, 39.4, 39.3, 39.2, 22.6, 21.0, 11.5

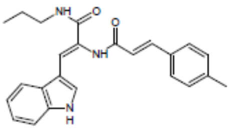


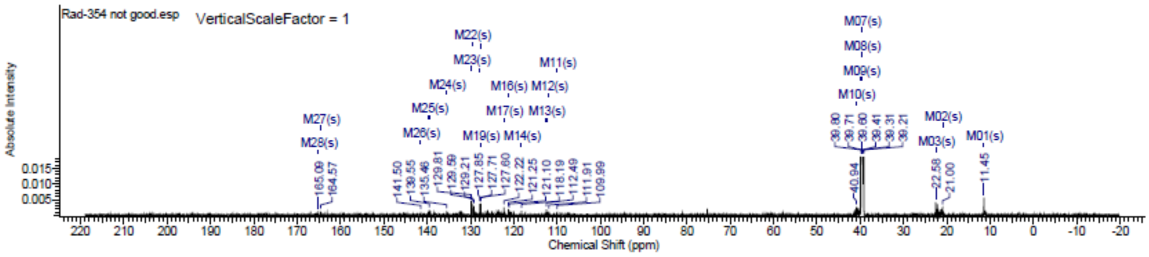

### Spectra of (4112)

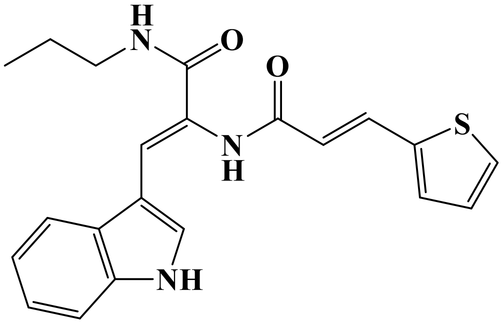


## LC/MS

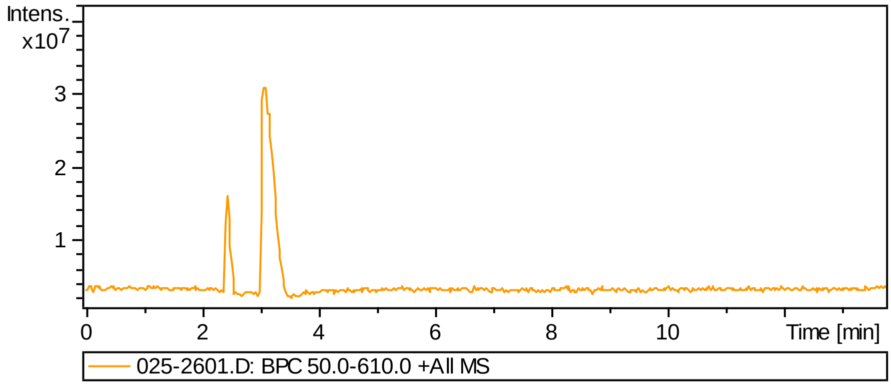


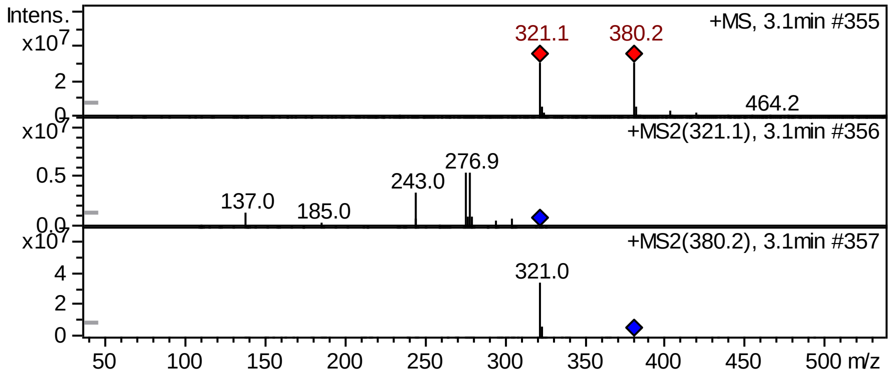


## FT-IR

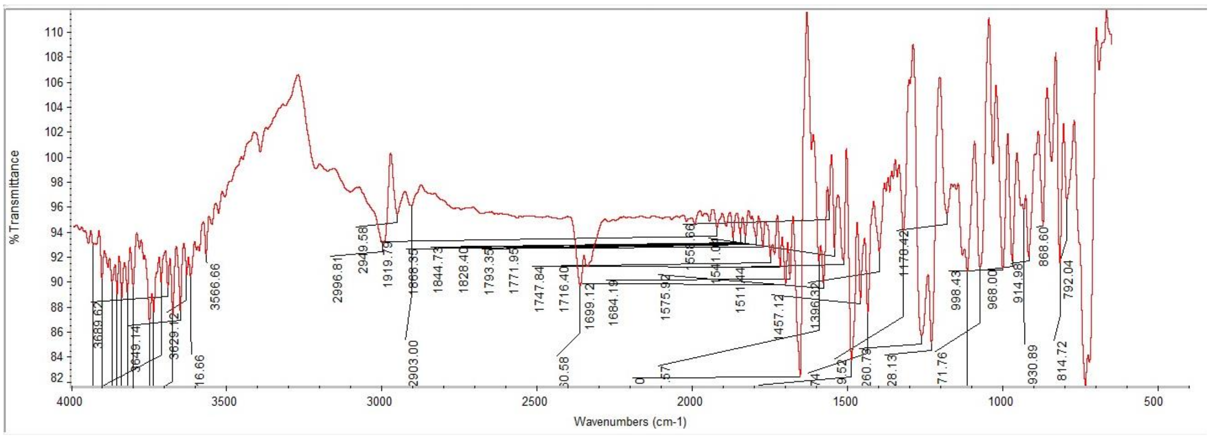

#### $^{1}H$ NMR

This report was created by ACD/NMR Processor Academic Edition. For more information go to www.acdlabs.com/nmrproc/

##### RAD-355

09/11/2016 8:15:32 AM
Dr.Mustafa Sample : RAD-355 DMSO PROTON DMSO {D:\Magdy} nmr 27

| Formula $C_{21}H_{21}N_3O_2S$ | FW 379.4753 |
|---|---|

| | | | | | | | |
|---|---|---|---|---|---|---|---|
| Acquisition Time (sec) | 2.6564 | Comment | Dr.Mustafa Sample : RAD-355 DMSO PROTON DMSO {D:\Magdy} nmr 27 | | | | |
| Date | 18 Jun 2015 15:54:08 | | | Date Stamp | 18 Jun 2015 15:54:08 | | |
| File Name | E:\Mostafa Alaraby Project\Oxazolone NMR\NMR final oxazolone\MUSTAFA RAD-355 18-06-2015\10\fid | | | | | Frequency (MHz) | 600.15 |
| Nucleus | 1H | Number of Transients | 32 | Origin | spect | Original Points Count | 32768 |
| Owner | nmr | Points Count | 32768 | Pulse Sequence | zg30 | Receiver Gain | 161.00 |
| SW(cyclical) (Hz) | 12335.53 | Solvent | DMSO-d6 | Spectrum Offset (Hz) | 3706.1750 | Spectrum Type | STANDARD |
| Sweep Width (Hz) | 12335.15 | Temperature (degree C) | 25.000 | | | | |

$^{1}H$ NMR (600 MHz, DMSO-$d_6$) δ 11.56 (br. s., 1H), 9.41 (s, 1H), 7.96 (t, $J$ = 5.83 Hz, 1H), 7.72 (d, $J$ = 7.91 Hz, 1H), 7.65 - 7.70 (m, 2H), 7.64 (d, $J$ = 1.88 Hz, 1H), 7.50 (s, 1H), 7.42 - 7.46 (m, 2H), 7.14 - 7.19 (m, 2H), 7.10 - 7.14 (m, 1H), 6.73 (d, $J$ = 15.43 Hz, 1H), 3.13 (q, $J$ = 6.40 Hz, 2H), 1.49 (sxt, $J$ = 7.23 Hz, 2H), 0.88 (t, $J$ = 7.53 Hz, 3H)

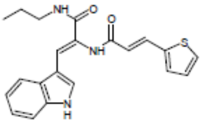


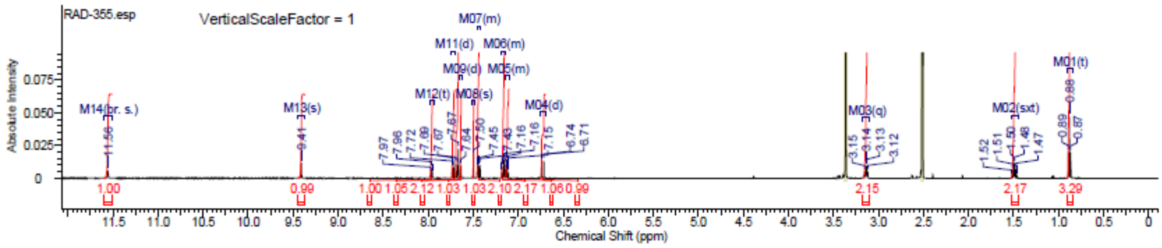


#### $^{13}C$ NMR

This report was created by ACD/NMR Processor Academic Edition. For more information go to www.acdlabs.com/nmrproc/

##### RAD-355

25/09/2016 3:57:30 PM
Dr.Moustafa Sample : RAD-355 DMSO

| Formula $C_{21}H_{21}N_3O_2S$ | FW 379.4753 |
|---|---|

| | | | | | | | |
|---|---|---|---|---|---|---|---|
| Acquisition Time (sec) | 0.6423 | Comment | Dr.Moustafa Sample : RAD-355 DMSO | | | Date | 23 Apr 2016 00:24:16 |
| Date Stamp | 23 Apr 2016 00:24:16 | | | | | | |
| File Name | E:\Projects\Mostafa Alaraby Project\Oxazolone\Oxazolone NMR\NMR final oxazolone\13 CNMR New 18-4-2016\MUSTAFA RAD-355 21-04-2016\40\fid | | | | | | |
| Frequency (MHz) | 213.77 | Nucleus | 13C | Number of Transients | 12288 | Origin | spect |
| Original Points Count | 32768 | Owner | nmr | Points Count | 32768 | Pulse Sequence | zgpg30 |
| Receiver Gain | 186.93 | SW(cyclical) (Hz) | 51020.41 | Solvent | DMSO-d6 | Spectrum Offset (Hz) | 21292.2539 |
| Spectrum Type | STANDARD | Sweep Width (Hz) | 51018.85 | Temperature (degree C) | 25.000 | | |

$^{13}C$ NMR (214 MHz, DMSO-$d_6$) δ 163.8, 163.2, 139.7, 138.1, 136.5, 136.2, 133.4, 133.3, 132.4, 129.1, 128.3, 123.6, 122.9, 122.2, 121.1, 120.6, 112.3, 107.4, 41.4, 39.8, 39.7, 39.6, 39.4, 39.3, 39.2, 22.3, 11.3

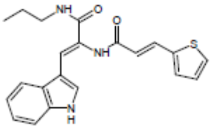


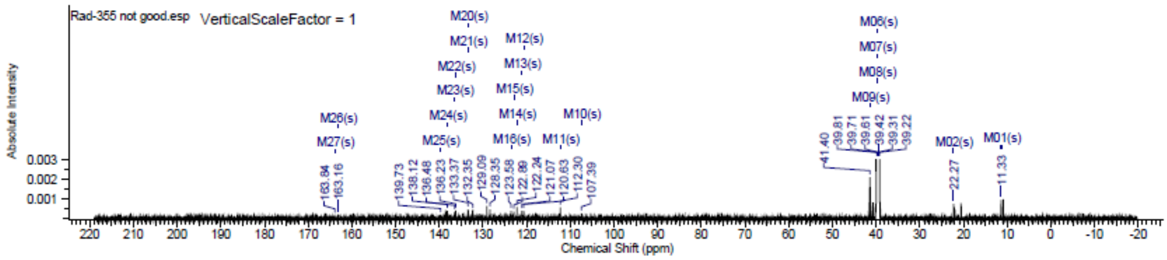

### Spectra of (4212)

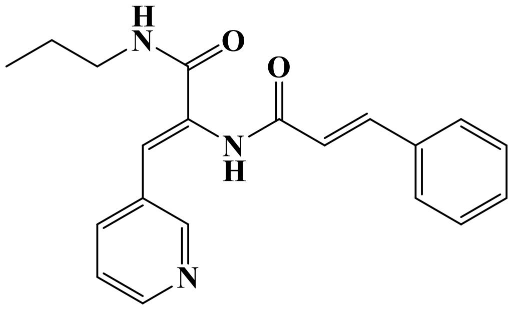

## LC/MS

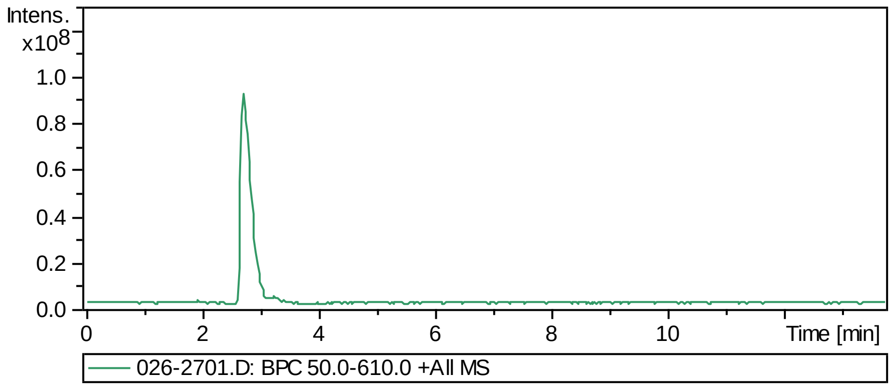


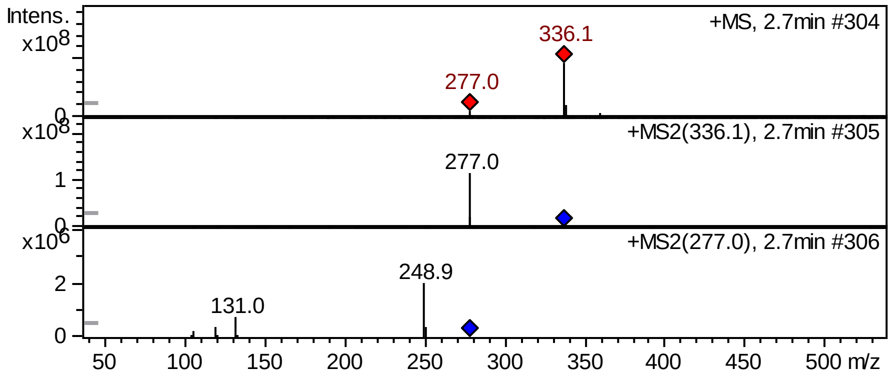


## FT-IR

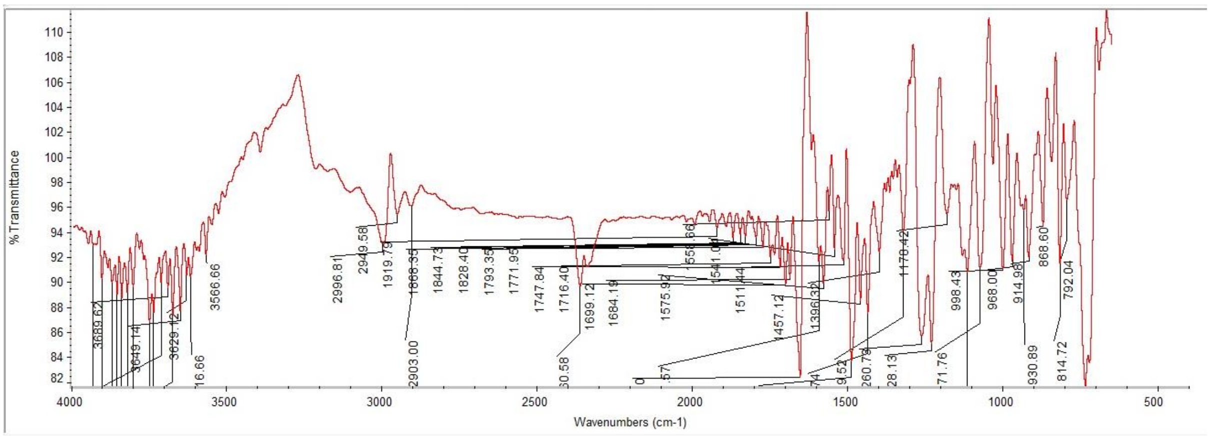

#### $^{1}$H NMR

This report was created by ACD/NMR Processor Academic Edition. For more information go to www.acdlabs.com/nmrproc/

##### RAD-361

09/11/2016 8:21:08 AM
Dr.Mustafa Sample : RAD-361 DMSO PROTON DMSO {D:\Magdy} nmr 31

| Formula $C_{20}H_{21}N_3O_2$ | FW 335.3996 |
|---|---|

| | | | | | | | |
|---|---|---|---|---|---|---|---|
| Acquisition Time (sec) | 2.6564 | Comment | Dr.Mustafa Sample : RAD-361 DMSO PROTON DMSO {D:\Magdy} nmr 31 | | | | |
| Date | 18 Jun 2015 16:15:28 | | | Date Stamp | 18 Jun 2015 16:15:28 | | |
| File Name | E:\Mostafa Alaraby Project\Oxazolone NMR\NMR final oxazolone\MUSTAFA RAD-361 18-06-2015\10\fid | | | | | Frequency (MHz) | 600.15 |
| Nucleus | 1H | Number of Transients | 32 | Origin | spect | Original Points Count | 32768 |
| Owner | nmr | Points Count | 32768 | Pulse Sequence | zg30 | Receiver Gain | 144.00 |
| SW(cyclical) (Hz) | 12335.53 | Solvent | DMSO-d6 | Spectrum Offset (Hz) | 3706.1750 | Spectrum Type | STANDARD |
| Sweep Width (Hz) | 12335.15 | Temperature (degree C) | 25.000 | | | | |

$^{1}$H NMR (600 MHz, DMSO-$d_6$) δ 9.82 (s, 1H), 8.70 (d, $J$ = 2.26 Hz, 1H), 8.47 (dd, $J$ = 1.69, 4.71 Hz, 1H), 8.25 (t, $J$ = 5.65 Hz, 1H), 7.92 (td, $J$ = 1.74, 8.19 Hz, 1H), 7.60 - 7.66 (m, 2H), 7.51 (d, $J$ = 15.81 Hz, 1H), 7.39 - 7.49 (m, 4H), 7.00 (s, 1H), 6.86 (d, $J$ = 16.19 Hz, 1H), 3.09 - 3.17 (m, 2H), 1.49 (sxt, $J$ = 7.30 Hz, 2H), 0.88 (t, $J$ = 7.34 Hz, 3H)

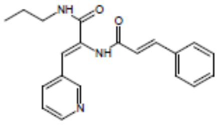


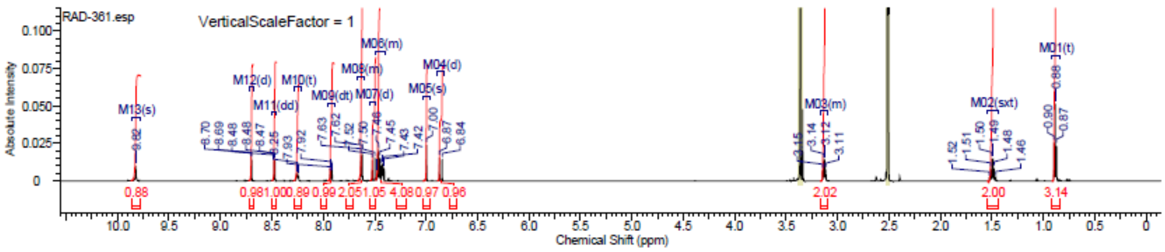


#### $^{13}$C NMR

This report was created by ACD/NMR Processor Academic Edition. For more information go to www.acdlabs.com/nmrproc/

##### RAD-361

20/09/2016 9:41:55 AM
Dr.Mostafa Sample : RAD-361 DMSO

| Formula $C_{20}H_{21}N_3O_2$ | FW 335.3996 |
|---|---|

| | | | | | | | |
|---|---|---|---|---|---|---|---|
| Acquisition Time (sec) | 0.6423 | Comment | Dr.Mostafa Sample : RAD-361 DMSO | | | Date | 19 Apr 2016 11:08:32 |
| Date Stamp | 19 Apr 2016 11:08:32 | | | | | | |
| File Name | E:\Projects\Mostafa Alaraby Project\Oxazolone\Oxazolone NMR\NMR final oxazolone\13 CNMR New 18-4-2016\MUSTAFA RAD-361 18-04-2016\130\fid | | | | | | |
| Frequency (MHz) | 213.77 | Nucleus | 13C | Number of Transients | 1320 | Origin | spect |
| Original Points Count | 32768 | Owner | nmr | Points Count | 32768 | Pulse Sequence | zgpg30 |
| Receiver Gain | 186.93 | SW(cyclical) (Hz) | 51020.41 | Solvent | DMSO-d6 | Spectrum Offset (Hz) | 21289.1406 |
| Spectrum Type | STANDARD | Sweep Width (Hz) | 51018.85 | Temperature (degree C) | 25.002 | | |

$^{13}$C NMR (214 MHz, DMSO-$d_6$) δ 164.7, 164.5, 150.1, 148.9, 140.3, 135.9, 134.7, 132.2, 130.5, 129.9, 129.1, 127.8, 123.7, 123.2, 121.4, 41.0, 39.8, 39.7, 39.6, 39.4, 39.3, 39.2, 22.4, 11.5

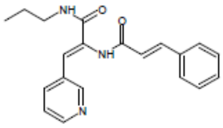


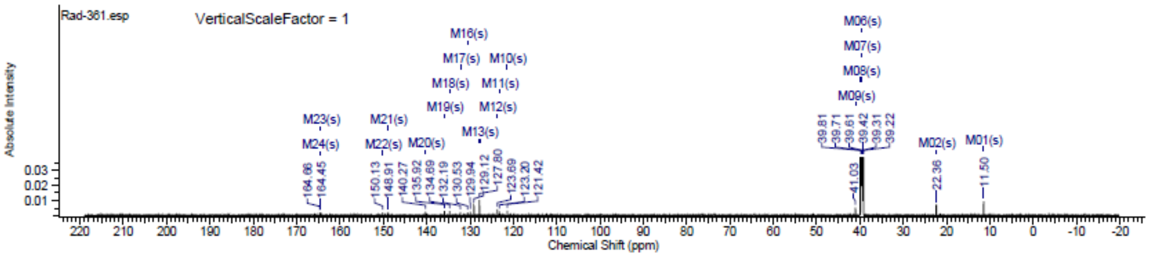

### Spectra of (4312)

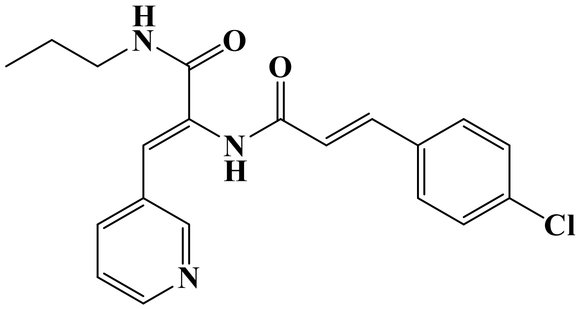

## FT-IR

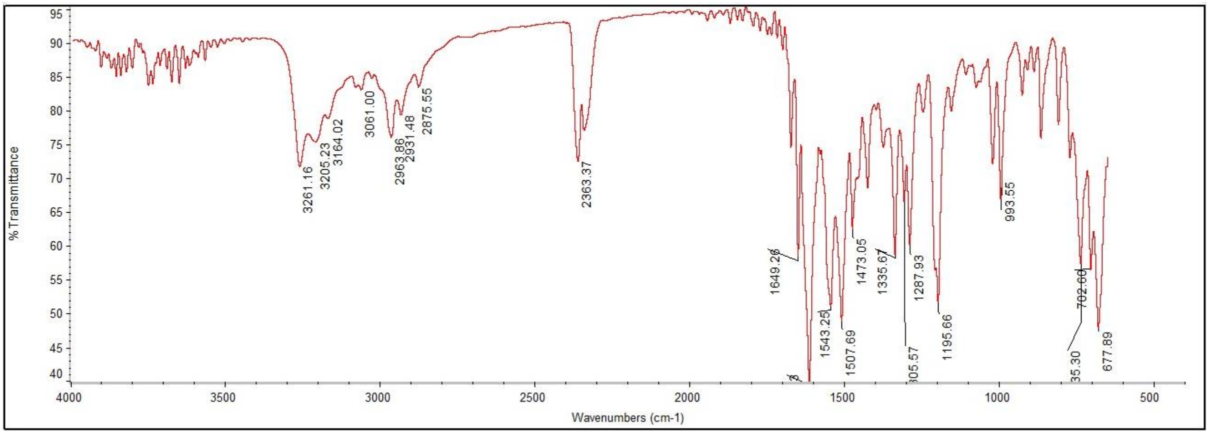


#### $^1$H NMR

This report was created by ACD/NMR Processor Academic Edition. For more information go to www.acdlabs.com/nmrproc/

**RAD-362**

09/11/2016 8:30:38 AM
Dr.Mustafa Sample : RAD-362 DMSO PROTON DMSO {D:\Magdy} nmr 12

| Formula $C_{20}H_{20}ClN_3O_2$ | FW 369.8447 |
|---|---|

| Acquisition Time (sec) | 2.6564 | Comment | Dr.Mustafa Sample : RAD-362 DMSO PROTON DMSO {D:\Magdy} nmr 12 | | | | |
|---|---|---|---|---|---|---|---|
| Date | 18 Jun 2015 14:35:12 | | | Date Stamp | 18 Jun 2015 14:35:12 | | |
| File Name | E:\Mostafa Alaraby Project\Oxazolone NMR\NMR final oxazolone\MUSTAFA RAD-362 18-06-2015\10\fid | | | | | Frequency (MHz) | 600.15 |
| Nucleus | 1H | Number of Transients | 32 | Origin | spect | Original Points Count | 32768 |
| Owner | nmr | Points Count | 32768 | Pulse Sequence | zg30 | Receiver Gain | 181.00 |
| SW(cyclical) (Hz) | 12335.53 | Solvent | DMSO-d6 | Spectrum Offset (Hz) | 3706.1750 | Spectrum Type | STANDARD |
| Sweep Width (Hz) | 12335.15 | Temperature (degree C) | 25.000 | | | | |

$^1$H NMR (600 MHz, DMSO-$d_6$) δ 9.83 (br. s., 1H), 8.70 (d, $J$ = 2.26 Hz, 1H), 8.47 (dd, $J$ = 1.69, 4.71 Hz, 1H), 8.25 (t, $J$ = 5.46 Hz, 1H), 7.92 (d, $J$ = 7.91 Hz, 1H), 7.63 (d, $J$ = 7.15 Hz, 2H), 7.51 (d, $J$ = 15.81 Hz, 1H), 7.44 - 7.48 (m, 2H), 7.40 - 7.44 (m, 2H), 7.00 (s, 1H), 6.86 (d, $J$ = 15.81 Hz, 1H), 3.13 (q, $J$ = 6.53 Hz, 2H), 1.46 - 1.53 (m, 1H), 0.88 (t, $J$ = 7.34 Hz, 3H)

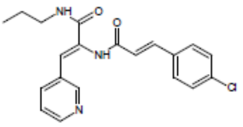


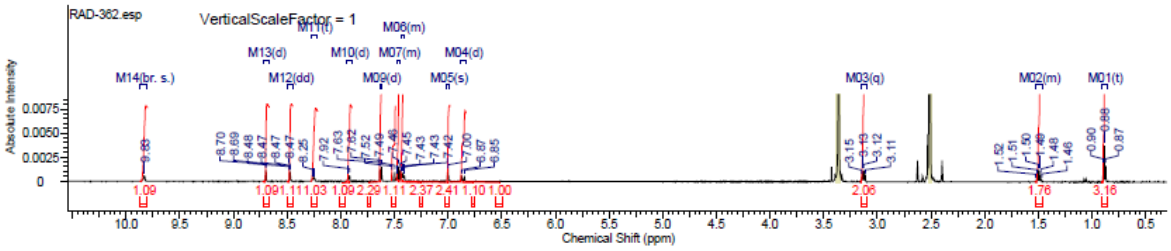

### Spectra of (4412)

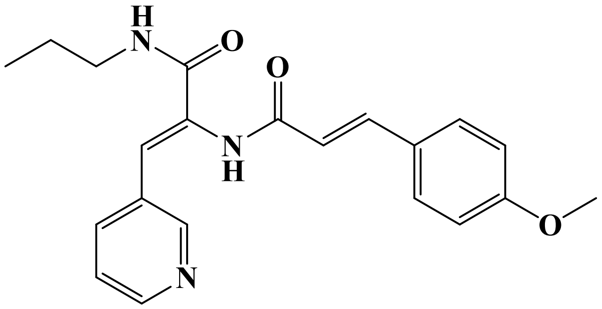

## LC/MS

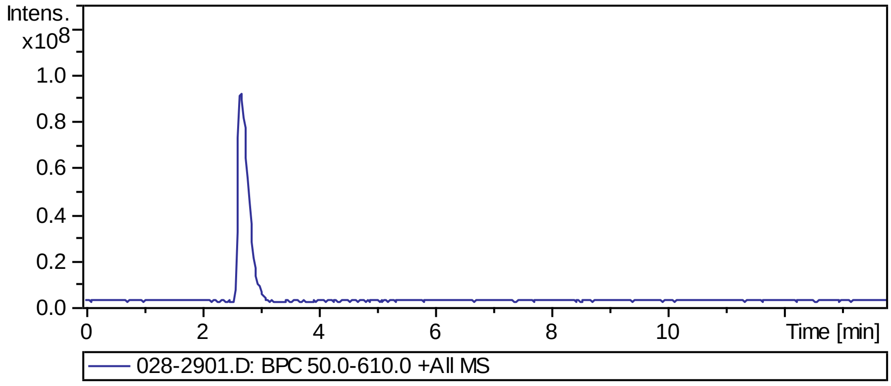


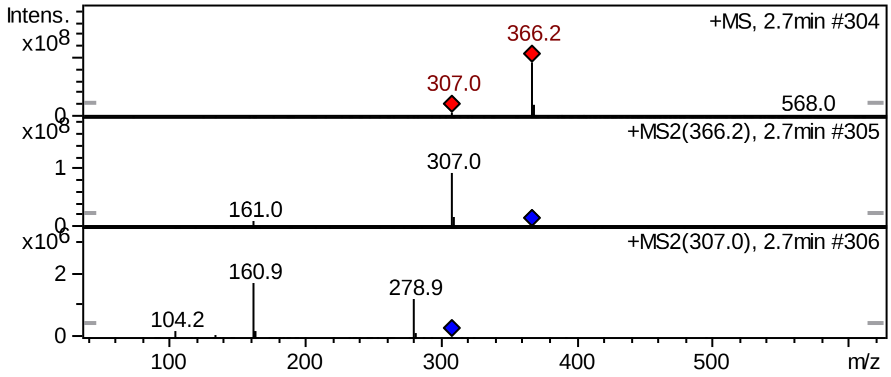


## FT-IR

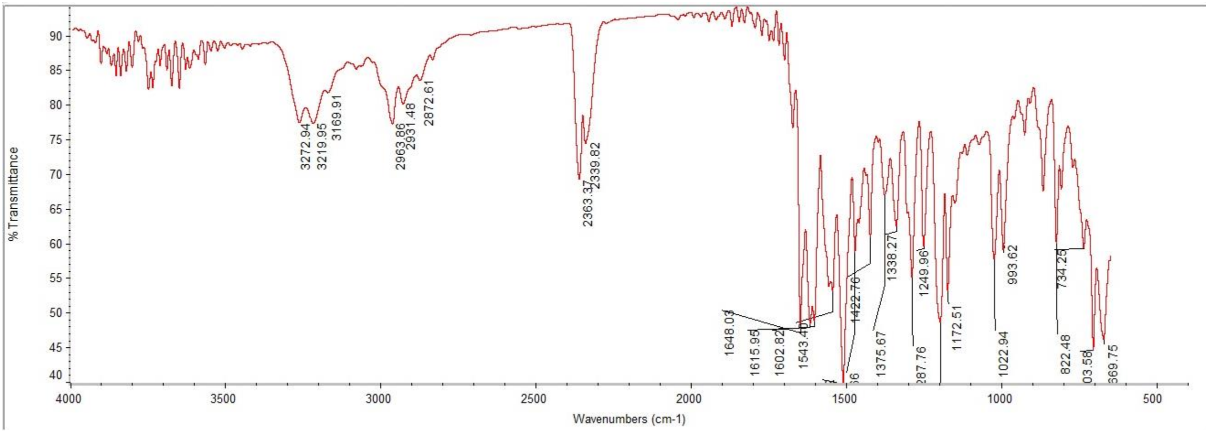

#### $^{1}$H NMR

This report was created by ACD/NMR Processor Academic Edition. For more information go to www.acdlabs.com/nmrproc/

**RAD-363**

09/11/2016 8:34:26 AM
Dr.Mustafa Sample : RAD-363 DMSO PROTON DMSO {D:\Magdy} nmr 13

| Formula $C_{21}H_{23}N_3O_3$ | FW 365.4256 |
|---|---|

| | | | | | | | |
|---|---|---|---|---|---|---|---|
| Acquisition Time (sec) | 2.6564 | Comment | Dr.Mustafa Sample : RAD-363 DMSO PROTON DMSO {D:\Magdy} nmr 13 | | | | |
| Date | 18 Jun 2015 14:41:36 | | | Date Stamp | 18 Jun 2015 14:41:36 | | |
| File Name | E:\Mostafa Alaraby Project\Oxazolone NMR\NMR final oxazolone\MUSTAFA RAD-363 18-06-2015\10\fid | | | | | Frequency (MHz) | 600.15 |
| Nucleus | 1H | Number of Transients | 32 | Origin | spect | Original Points Count | 32768 |
| Owner | nmr | Points Count | 32768 | Pulse Sequence | zg30 | Receiver Gain | 144.00 |
| SW(cyclical) (Hz) | 12335.53 | Solvent | DMSO-d6 | Spectrum Offset (Hz) | 3706.1750 | Spectrum Type | STANDARD |
| Sweep Width (Hz) | 12335.15 | Temperature (degree C) | 25.000 | | | | |

$^{1}$H NMR (600 MHz, DMSO-$d_6$) δ 9.72 (s, 1H), 8.69 (s, 1H), 8.47 (dd, *J* = 1.69, 4.71 Hz, 1H), 8.22 (t, *J* = 5.27 Hz, 1H), 7.91 (dd, *J* = 1.32, 8.09 Hz, 1H), 7.57 (d, *J* = 8.66 Hz, 2H), 7.46 (d, *J* = 15.43 Hz, 1H), 7.41 (dd, *J* = 4.71, 8.09 Hz, 1H), 7.02 (d, *J* = 8.66 Hz, 2H), 6.97 (s, 1H), 6.71 (d, *J* = 15.81 Hz, 1H), 3.81 (s, 3H), 3.10 - 3.16 (m, 2H), 1.49 (sxt, *J* = 7.23 Hz, 2H), 0.88 (t, *J* = 7.34 Hz, 3H)

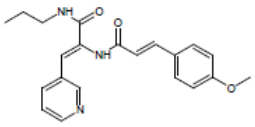

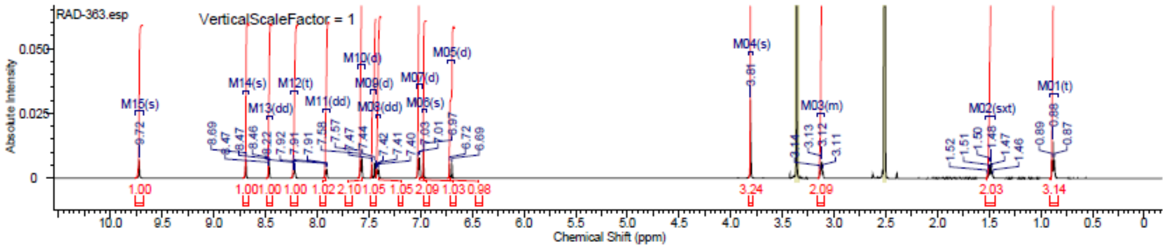


#### $^{13}$C NMR

This report was created by ACD/NMR Processor Academic Edition. For more information go to www.acdlabs.com/nmrproc/

**RAD-363**

20/09/2016 10:17:55 AM
Dr.Mustafa Sample ; RAD-363 DMSO

| Formula $C_{21}H_{23}N_3O_3$ | FW 365.4256 |
|---|---|

| | | | | | | | |
|---|---|---|---|---|---|---|---|
| Acquisition Time (sec) | 0.6423 | Comment | Dr.Mostafa Sample ; RAD-363 DMSO | | | Date | 19 Apr 2016 14:22:40 |
| Date Stamp | 19 Apr 2016 14:22:40 | | | | | | |
| File Name | E:\Projects\Mostafa Alaraby Project\Oxazolone\Oxazolone NMR\NMR final oxazolone\13 CNMR New 18-4-2016\MUSTAFA RAD-363 19-04-2016\10\fid | | | | | | |
| Frequency (MHz) | 213.77 | Nucleus | 13C | Number of Transients | 979 | Origin | spect |
| Original Points Count | 32768 | Owner | nmr | Points Count | 32768 | Pulse Sequence | zgpg30 |
| Receiver Gain | 186.93 | SW(cyclical) (Hz) | 51020.41 | Solvent | DMSO-d6 | Spectrum Offset (Hz) | 21292.2539 |
| Spectrum Type | STANDARD | Sweep Width (Hz) | 51018.85 | Temperature (degree C) | 25.001 | | |

$^{13}$C NMR (214 MHz, DMSO-$d_6$) δ 164.8, 164.4, 160.7, 150.1, 148.9, 140.1, 135.9, 132.3, 130.6, 129.5, 127.3, 123.7, 123.0, 118.8, 114.6, 55.4, 41.0, 39.8, 39.7, 39.6, 39.4, 39.3, 39.2, 22.4, 11.5

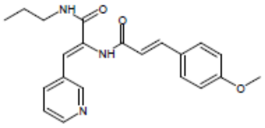

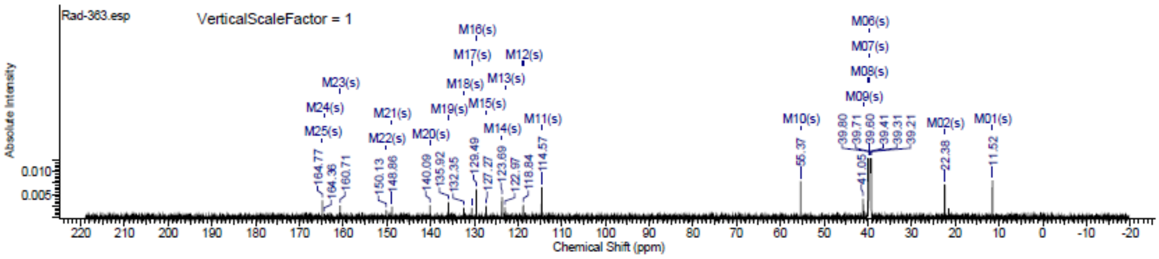

#### Spectra of (4512)

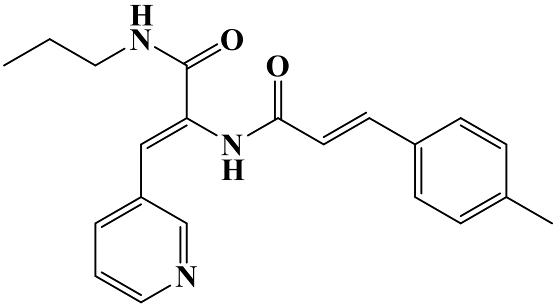


**LC/MS**

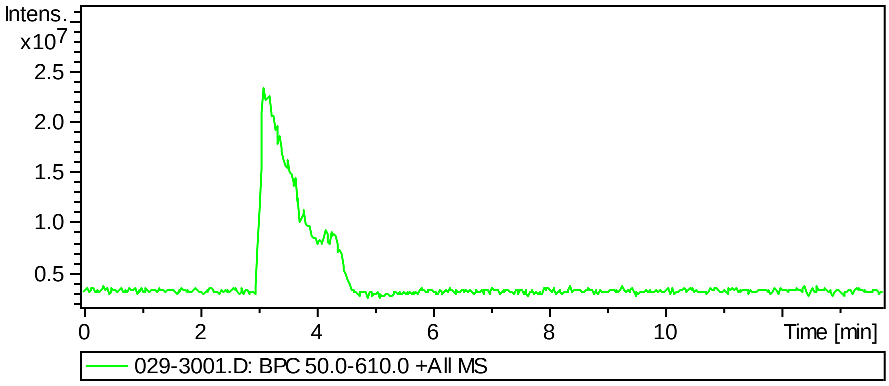


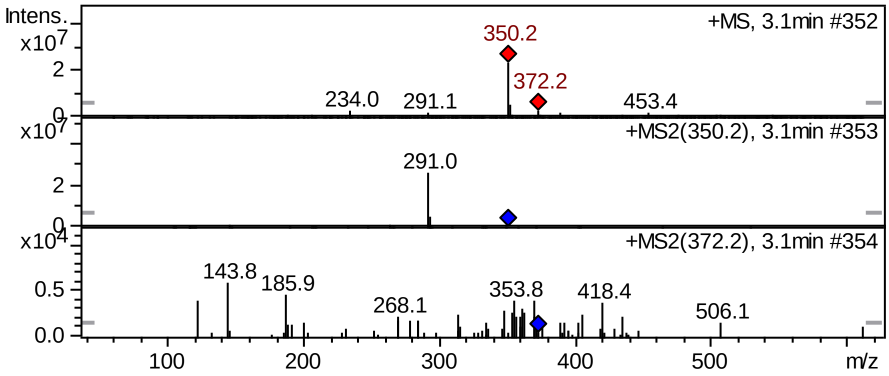

#### $^{1}$H NMR

This report was created by ACD/NMR Processor Academic Edition. For more information go to www.acdlabs.com/nmrproc/

**RAD-364**

09/11/2016 8:37:59 AM
Dr.Mustafa Sample : RAD-364 DMSO PROTON DMSO {D:\Magdy} nmr 21

| | |
|---|---|
| *Formula* $C_{21}H_{23}N_3O_2$ | *FW* 349.4262 |

| | | | | | | | |
|---|---|---|---|---|---|---|---|
| *Acquisition Time (sec)* | 2.6564 | *Comment* | Dr.Mustafa Sample : RAD-364 DMSO PROTON DMSO {D:\Magdy} nmr 21 | | | | |
| *Date* | 18 Jun 2015 15:22:08 | | | *Date Stamp* | 18 Jun 2015 15:22:08 | | |
| *File Name* | E:\Mostafa Alaraby Project\Oxazolone NMR\NMR final oxazolone\MUSTAFA RAD-364 18-06-2015\10\fid | | | | | *Frequency (MHz)* | 600.15 |
| *Nucleus* | 1H | *Number of Transients* | 32 | *Origin* | spect | *Original Points Count* | 32768 |
| *Owner* | nmr | *Points Count* | 32768 | *Pulse Sequence* | zg30 | *Receiver Gain* | 161.00 |
| *SW(cyclical) (Hz)* | 12335.53 | *Solvent* | DMSO-d6 | *Spectrum Offset (Hz)* | 3706.1750 | *Spectrum Type* | STANDARD |
| *Sweep Width (Hz)* | 12335.15 | *Temperature (degree C)* | 25.000 | | | | |

$^{1}$H NMR (600 MHz, DMSO-$d_6$) δ 9.77 (s, 1H), 8.69 (d, $J$ = 1.88 Hz, 1H), 8.47 (dd, $J$ = 1.69, 4.71 Hz, 1H), 8.23 (t, $J$ = 5.83 Hz, 1H), 7.92 (td, $J$ = 1.60, 8.09 Hz, 1H), 7.49 - 7.54 (m, $J$ = 7.91 Hz, 2H), 7.47 (d, $J$ = 15.81 Hz, 1H), 7.41 (dd, $J$ = 4.33, 8.09 Hz, 1H), 7.24 - 7.30 (m, $J$ = 7.91 Hz, 2H), 6.98 (s, 1H), 6.80 (d, $J$ = 15.81 Hz, 1H), 3.12 (q, $J$ = 6.65 Hz, 2H), 2.35 (s, 3H), 1.49 (sxt, $J$ = 7.30 Hz, 2H), 0.88 (t, $J$ = 7.34 Hz, 3H)

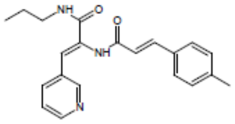


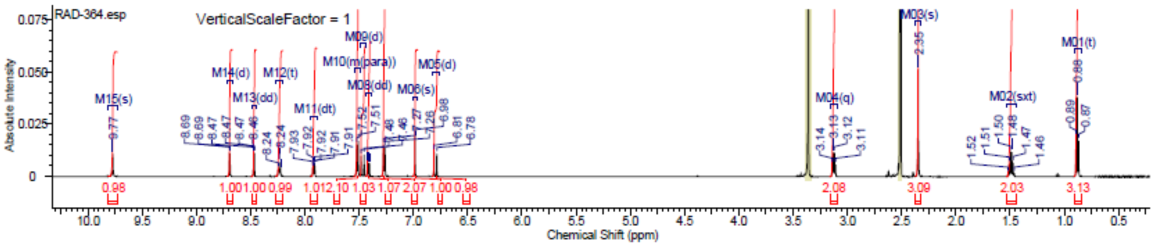


#### $^{13}$C NMR

This report was created by ACD/NMR Processor Academic Edition. For more information go to www.acdlabs.com/nmrproc/

**RAD-364**

20/09/2016 9:50:37 AM
Dr.Mostafa Sample ; RAD-364 DMSO

| | |
|---|---|
| *Formula* $C_{21}H_{23}N_3O_2$ | *FW* 349.4262 |

| | | | | | | | |
|---|---|---|---|---|---|---|---|
| *Acquisition Time (sec)* | 0.6423 | *Comment* | Dr.Mostafa Sample ; RAD-364 DMSO | | | *Date* | 20 Apr 2016 13:08:00 |
| *Date Stamp* | 20 Apr 2016 13:08:00 | | | | | | |
| *File Name* | E:\Projects\Mostafa Alaraby Project\Oxazolone\Oxazolone NMR\NMR final oxazolone\13 CNMR New 18-4-2016\MUSTAFA RAD-364 19-04-2016\110\fid | | | | | | |
| *Frequency (MHz)* | 213.77 | *Nucleus* | 13C | *Number of Transients* | 1147 | *Origin* | spect |
| *Original Points Count* | 32768 | *Owner* | nmr | *Points Count* | 32768 | *Pulse Sequence* | zgpg30 |
| *Receiver Gain* | 186.93 | *SW(cyclical) (Hz)* | 51020.41 | *Solvent* | DMSO-d6 | *Spectrum Offset (Hz)* | 21293.8125 |
| *Spectrum Type* | STANDARD | *Sweep Width (Hz)* | 51018.85 | *Temperature (degree C)* | 24.999 | | |

$^{13}$C NMR (214 MHz, DMSO-$d_6$) δ 164.7, 164.6, 150.1, 148.9, 140.3, 139.8, 135.9, 132.3, 132.0, 130.6, 129.7, 127.8, 123.7, 123.1, 120.4, 41.1, 39.8, 39.7, 39.6, 39.4, 39.3, 39.2, 22.4, 21.1, 11.5

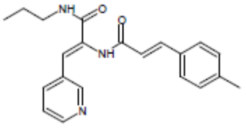


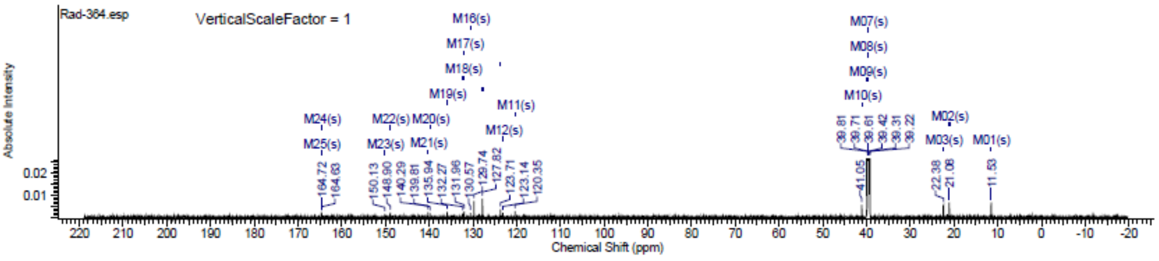

### Spectra of (4612)

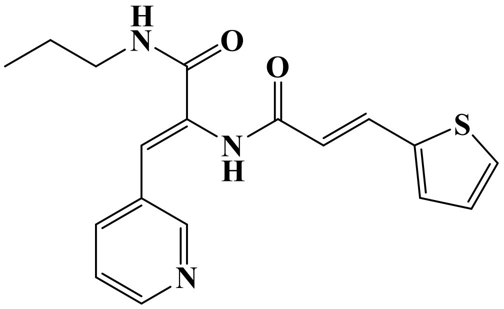

## LC/MS

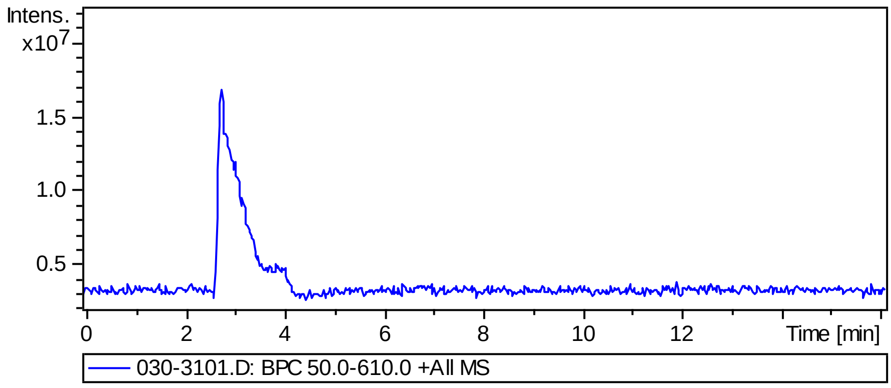


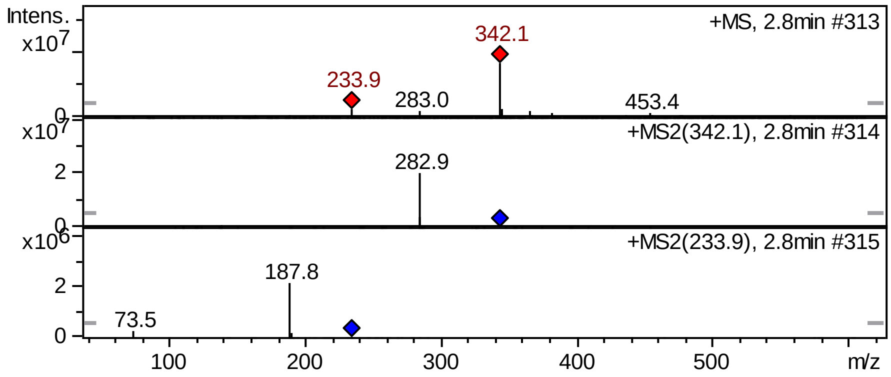


## FT-IR

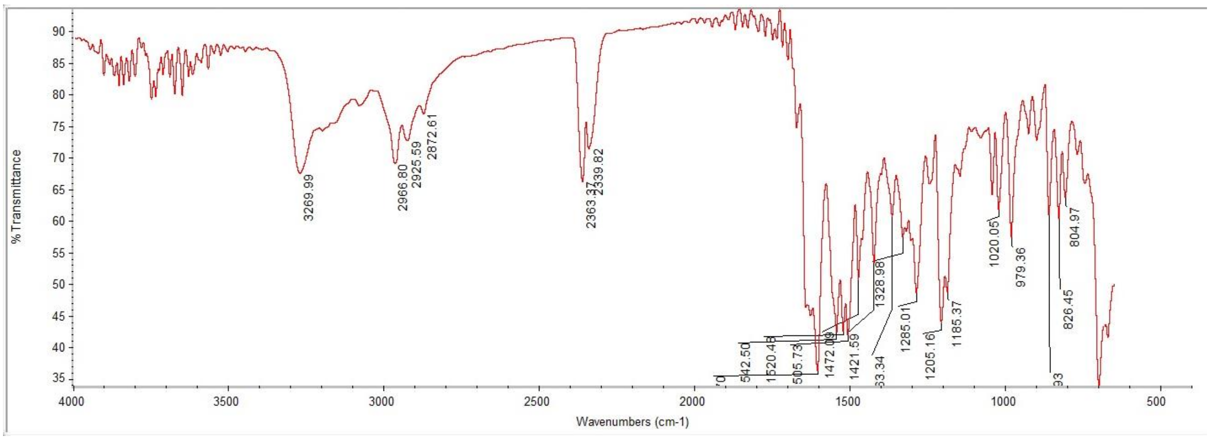

### [1]H NMR

This report was created by ACD/NMR Processor Academic Edition. For more information go to www.acdlabs.com/nmrproc/

#### RAD-365

09/11/2016 8:39:53 AM
Dr.Mustafa Sample : RAD-365 DMSO PROTON DMSO {D:\Magdy} nmr 22

| Formula $C_{18}H_{19}N_3O_2S$ | FW 341.4274 |
|---|---|

| | | | | | | | |
|---|---|---|---|---|---|---|---|
| Acquisition Time (sec) | 2.6564 | Comment | Dr.Mustafa Sample : RAD-365 DMSO PROTON DMSO {D:\Magdy} nmr 22 | | | | |
| Date | 18 Jun 2015 15:28:32 | | | Date Stamp | 18 Jun 2015 15:28:32 | | |
| File Name | E:\Mostafa Alaraby Project\Oxazolone NMR\NMR final oxazolone\MUSTAFA RAD-365 18-06-2015\10\fid | | | | | Frequency (MHz) | 600.15 |
| Nucleus | 1H | Number of Transients | 32 | Origin | spect | Original Points Count | 32768 |
| Owner | nmr | Points Count | 32768 | Pulse Sequence | zg30 | Receiver Gain | 144.00 |
| SW(cyclical) (Hz) | 12335.53 | Solvent | DMSO-d6 | Spectrum Offset (Hz) | 3706.1750 | Spectrum Type | STANDARD |
| Sweep Width (Hz) | 12335.15 | Temperature (degree C) | 25.000 | | | | |

[1]H NMR (600 MHz, DMSO-$d_6$) δ 9.78 (br. s., 1H), 8.68 (d, $J$ = 1.88 Hz, 1H), 8.47 (dd, $J$ = 1.51, 4.89 Hz, 1H), 8.23 (t, $J$ = 5.84 Hz, 1H), 7.91 (td, $J$ = 1.79, 8.09 Hz, 1H), 7.62 - 7.70 (m, 2H), 7.39 - 7.47 (m, 2H), 7.15 (dd, $J$ = 3.58, 5.08 Hz, 1H), 6.99 (s, 1H), 6.60 (d, $J$ = 15.81 Hz, 1H), 3.12 (q, $J$ = 6.65 Hz, 2H), 1.49 (sxt, $J$ = 7.23 Hz, 2H), 0.88 (t, $J$ = 7.53 Hz, 3H)

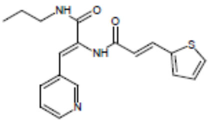


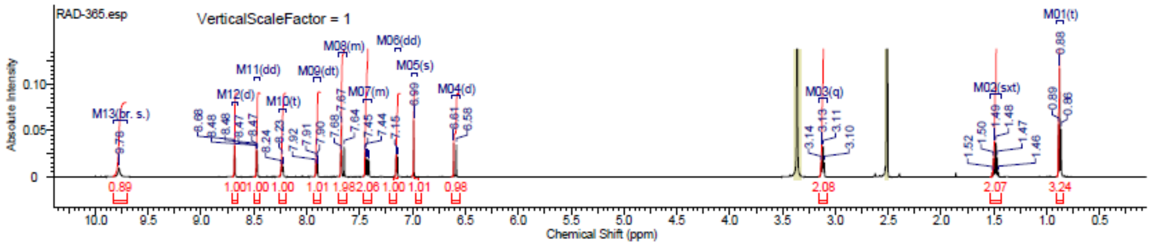


### [13]C NMR

This report was created by ACD/NMR Processor Academic Edition. For more information go to www.acdlabs.com/nmrproc/

#### RAD-365

20/09/2016 9:54:23 AM
Dr.Mostafa Sample ; RAD-365 DMSO

| Formula $C_{18}H_{19}N_3O_2S$ | FW 341.4274 |
|---|---|

| | | | | | | | |
|---|---|---|---|---|---|---|---|
| Acquisition Time (sec) | 0.6423 | Comment | Dr.Mostafa Sample ; RAD-365 DMSO | | | Date | 20 Apr 2016 17:24:00 |
| Date Stamp | 20 Apr 2016 17:24:00 | | | | | | |
| File Name | E:\Projects\Mostafa Alaraby Project\Oxazolone\Oxazolone NMR\NMR final oxazolone\13 CNMR New 18-4-2016\MUSTAFA RAD-365 19-04-2016\130\fid | | | | | | |
| Frequency (MHz) | 213.77 | Nucleus | 13C | Number of Transients | 3500 | Origin | spect |
| Original Points Count | 32768 | Owner | nmr | Points Count | 32768 | Pulse Sequence | zgpg30 |
| Receiver Gain | 186.93 | SW(cyclical) (Hz) | 51020.41 | Solvent | DMSO-d6 | Spectrum Offset (Hz) | 21284.4688 |
| Spectrum Type | STANDARD | Sweep Width (Hz) | 51018.85 | Temperature (degree C) | 25.000 | | |

[13]C NMR (214 MHz, DMSO-$d_6$) δ 164.6, 164.2, 150.1, 148.9, 139.7, 135.9, 133.3, 132.1, 131.5, 131.5, 130.5, 128.6, 123.7, 123.2, 120.1, 41.0, 39.8, 39.7, 39.6, 39.4, 39.3, 39.2, 22.3, 11.5

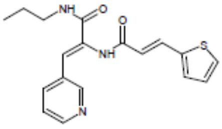


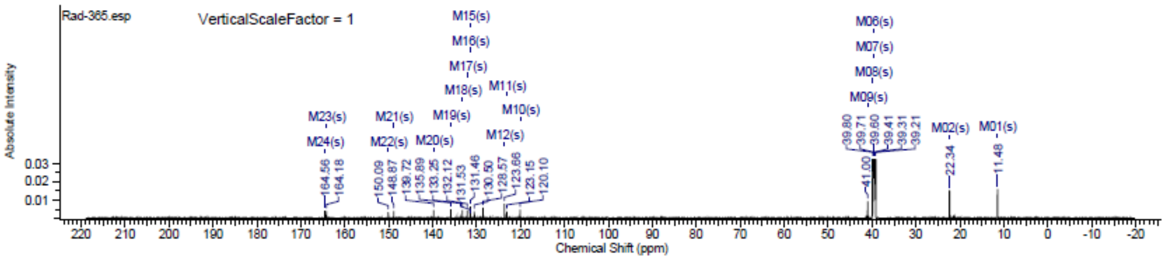

### Spectra of (4712)

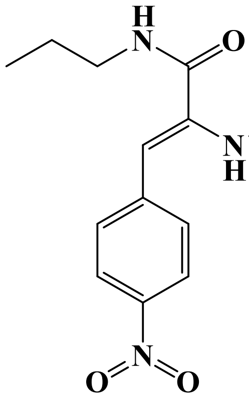

## LC/MS

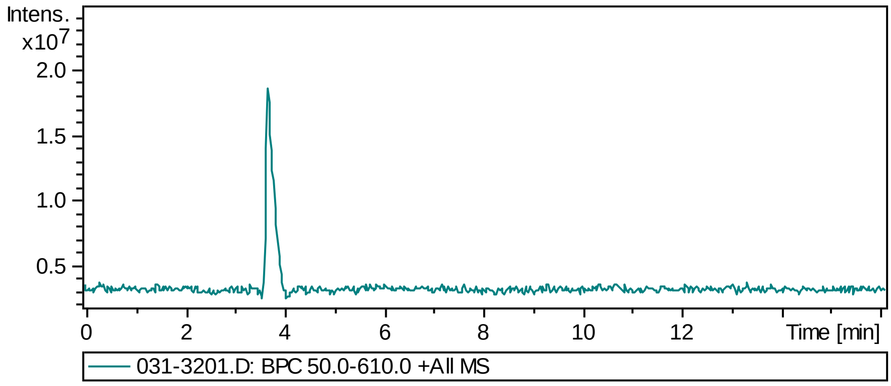


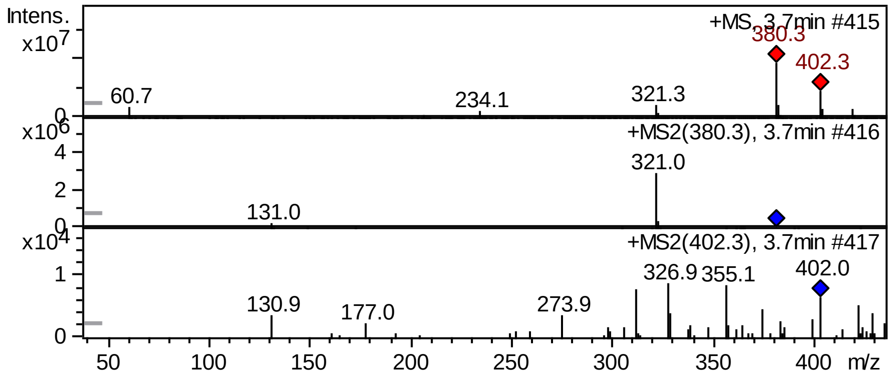


## FT-IR

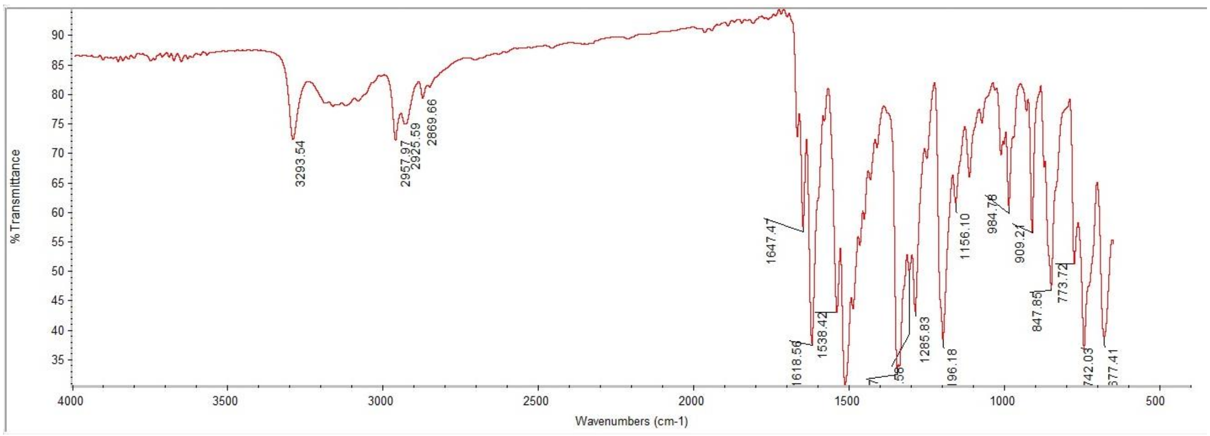

#### $^1$H NMR

This report was created by ACD/NMR Processor Academic Edition. For more information go to www.acdlabs.com/nmrproc/

##### RAD-371

11/10/2016 9:01:33 AM
Dr.Mustafa Sample : RAD-371 DMSO PROTON DMSO {D:\Magdy} nmr 17

| Formula $C_{21}H_{21}N_3O_4$ | FW 379.4091 |
|---|---|

| | | | | | | | |
|---|---|---|---|---|---|---|---|
| Acquisition Time (sec) | 2.6564 | Comment | Dr.Mustafa Sample : RAD-371 DMSO PROTON DMSO {D:\Magdy} nmr 17 | | | | |
| Date | 18 Jun 2015 15:00:48 | | | Date Stamp | 18 Jun 2015 15:00:48 | | |
| File Name | E:\Mostafa Alaraby Project\Oxazolone NMR\NMR final oxazolone\MUSTAFA RAD-371 18-06-2015\10\fid | | | | | Frequency (MHz) | 600.15 |
| Nucleus | 1H | Number of Transients | 32 | Origin | spect | Original Points Count | 32768 |
| Owner | nmr | Points Count | 32768 | Pulse Sequence | zg30 | Receiver Gain | 144.00 |
| SW(cyclical) (Hz) | 12335.53 | Solvent | DMSO-d6 | Spectrum Offset (Hz) | 3706.1750 | Spectrum Type | STANDARD |
| Sweep Width (Hz) | 12335.15 | Temperature (degree C) | 25.000 | | | | |

$^1$H NMR (600 MHz, DMSO-$d_6$) δ 9.94 (br. s., 1H), 8.34 (t, $J$ = 5.84 Hz, 1H), 8.21 - 8.26 (m, $J$ = 8.66 Hz, 2H), 7.74 - 7.79 (m, $J$ = 9.03 Hz, 2H), 7.62 (d, $J$ = 7.15 Hz, 2H), 7.51 (d, $J$ = 15.81 Hz, 1H), 7.43 - 7.48 (m, 3H), 6.96 (s, 1H), 6.86 (d, $J$ = 16.19 Hz, 1H), 3.10 - 3.17 (m, 2H), 1.50 (sxt, $J$ = 7.30 Hz, 2H), 0.89 (t, $J$ = 7.53 Hz, 3H)

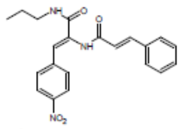


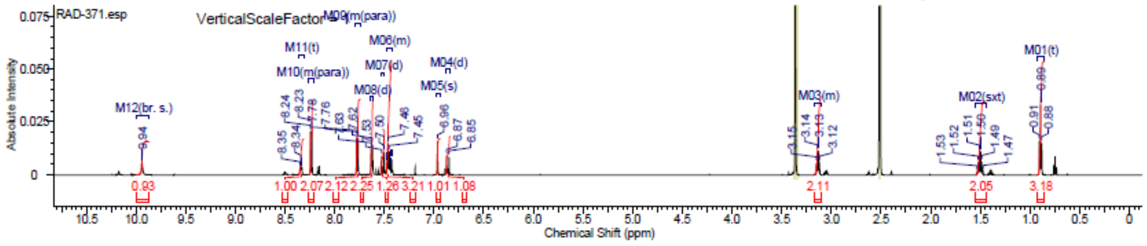


#### $^{13}$C NMR

This report was created by ACD/NMR Processor Academic Edition. For more information go to www.acdlabs.com/nmrproc/

##### RAD-371

20/09/2016 9:58:56 AM
Dr.Mostafa Sample ; RAD-371 DMSO

| Formula $C_{21}H_{21}N_3O_4$ | FW 379.4091 |
|---|---|

| | | | | | | | |
|---|---|---|---|---|---|---|---|
| Acquisition Time (sec) | 0.6423 | Comment | Dr.Mostafa Sample ; RAD-371 DMSO | | | Date | 21 Apr 2016 01:30:24 |
| Date Stamp | 21 Apr 2016 01:30:24 | | | | | | |
| File Name | E:\Projects\Mostafa Alaraby Project\Oxazolone\Oxazolone NMR\NMR final oxazolone\13 CNMR New 18-4-2016\MUSTAFA RAD-371 19-04-2016\160\fid | | | | | | |
| Frequency (MHz) | 213.77 | Nucleus | 13C | Number of Transients | 3500 | Origin | spect |
| Original Points Count | 32768 | Owner | nmr | Points Count | 32768 | Pulse Sequence | zgpg30 |
| Receiver Gain | 186.93 | SW(cyclical) (Hz) | 51020.41 | Solvent | DMSO-d6 | Spectrum Offset (Hz) | 21290.6973 |
| Spectrum Type | STANDARD | Sweep Width (Hz) | 51018.85 | Temperature (degree C) | 24.998 | | |

$^{13}$C NMR (214 MHz, DMSO-$d_6$) δ 164.7, 164.3, 146.4, 141.7, 140.5, 134.7, 133.8, 130.2, 130.0, 129.1, 127.8, 123.7, 123.0, 121.3, 41.1, 39.8, 39.7, 39.6, 39.4, 39.3, 39.2, 22.3, 11.5

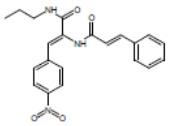


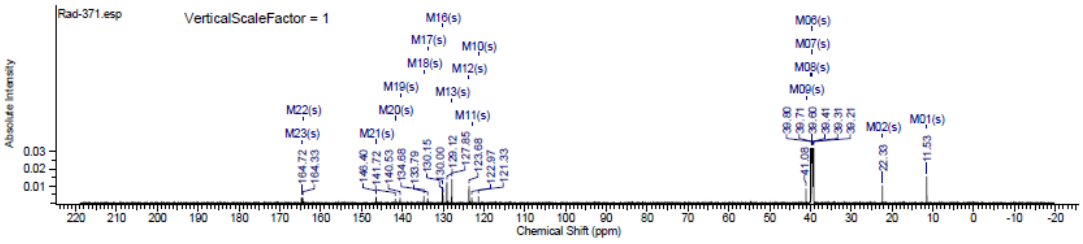

#### Spectra of (4812)

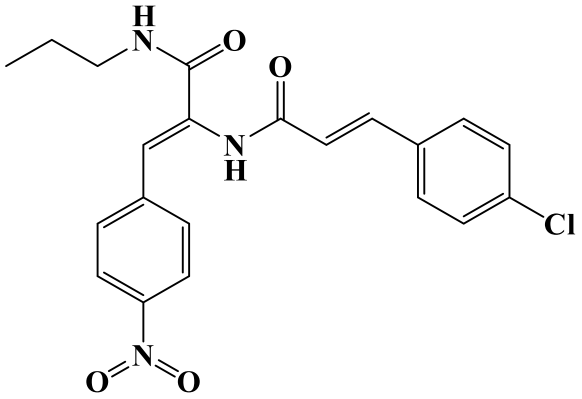


**LC/MS**

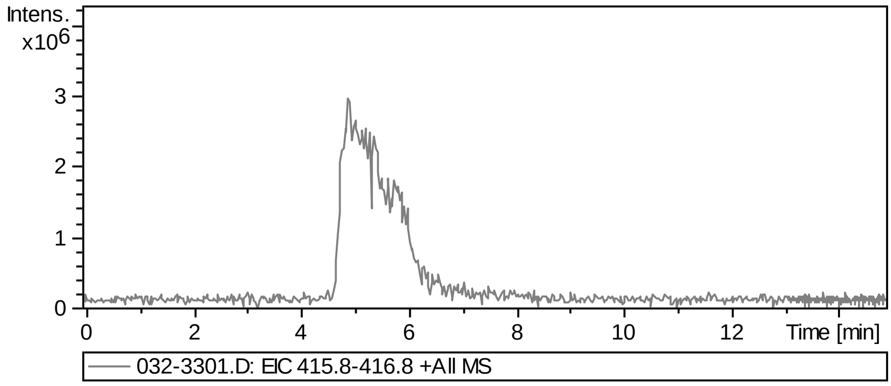


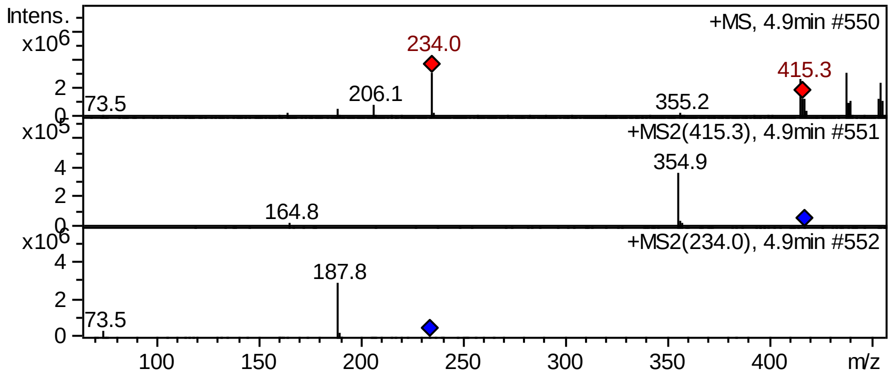


**FT-IR**

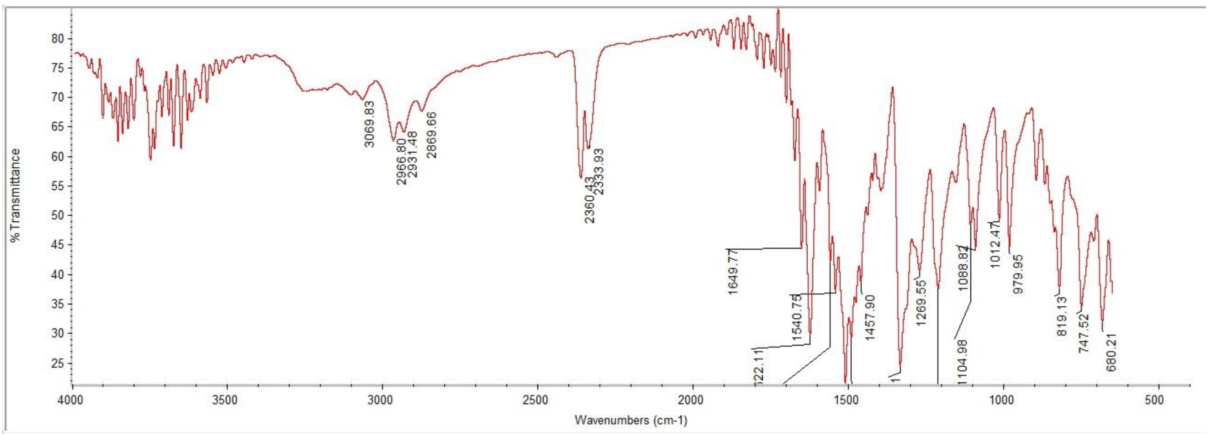

#### $^1$H NMR

This report was created by ACD/NMR Processor Academic Edition. For more information go to www.acdlabs.com/nmrproc/

**RAD-372**

11/10/2016 9:08:12 AM
Dr.Mustafa Sample : RAD-372 DMSO PROTON DMSO {D:\Magdy} nmr 20

| Formula | $C_{21}H_{20}ClN_3O_4$ | FW | 413.8542 |
|---|---|---|---|

| | | | | | | | |
|---|---|---|---|---|---|---|---|
| Acquisition Time (sec) | 2.6564 | Comment | Dr.Mustafa Sample : RAD-372 DMSO PROTON DMSO {D:\Magdy} nmr 20 | | | | |
| Date | 18 Jun 2015 15:17:52 | | | Date Stamp | 18 Jun 2015 15:17:52 | | |
| File Name | E:\Mostafa Alaraby Project\Oxazolone NMR\NMR final oxazolone\MUSTAFA RAD-372 18-06-2015\10\fid | | | | | Frequency (MHz) | 600.15 |
| Nucleus | 1H | Number of Transients | 32 | Origin | spect | Original Points Count | 32768 |
| Owner | nmr | Points Count | 32768 | Pulse Sequence | zg30 | Receiver Gain | 161.00 |
| SW(cyclical) (Hz) | 12335.53 | Solvent | DMSO-d6 | Spectrum Offset (Hz) | 3706.1750 | Spectrum Type | STANDARD |
| Sweep Width (Hz) | 12335.15 | Temperature (degree C) | 25.000 | | | | |

$^1$H NMR (600 MHz, DMSO-$d_6$) δ 9.95 (s, 1H), 8.34 (t, $J$ = 5.46 Hz, 1H), 8.21 - 8.25 (m, $J$ = 9.04 Hz, 2H), 7.75 - 7.79 (m, $J$ = 9.04 Hz, 2H), 7.65 (d, $J$ = 8.66 Hz, 2H), 7.50 - 7.55 (m, 3H), 6.97 (s, 1H), 6.86 (d, $J$ = 15.81 Hz, 1H), 3.13 (q, $J$ = 6.53 Hz, 2H), 1.50 (sxt, $J$ = 7.23 Hz, 2H), 0.89 (t, $J$ = 7.53 Hz, 3H)

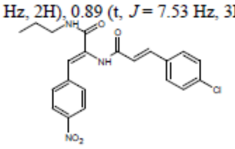


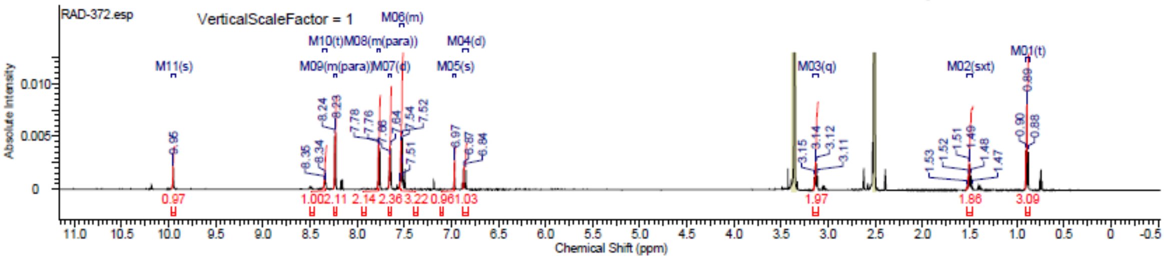


#### $^{13}$C NMR

This report was created by ACD/NMR Processor Academic Edition. For more information go to www.acdlabs.com/nmrproc/

**RAD-372**

20/09/2016 10:02:49 AM
Dr.Mostafa Sample : RAD-372 DMSO

| Formula | $C_{21}H_{20}ClN_3O_4$ | FW | 413.8542 |
|---|---|---|---|

| | | | | | | | |
|---|---|---|---|---|---|---|---|
| Acquisition Time (sec) | 0.6423 | Comment | Dr.Mostafa Sample : RAD-372 DMSO | | | Date | 19 Apr 2016 08:17:52 |
| Date Stamp | 19 Apr 2016 08:17:52 | | | | | | |
| File Name | E:\Projects\Mostafa Alaraby Project\Oxazolone\Oxazolone NMR\NMR final oxazolone\13 CNMR New 18-4-2016\MUSTAFA RAD-372 18-04-2016\90\fid | | | | | | |
| Frequency (MHz) | 213.77 | Nucleus | 13C | Number of Transients | 3072 | Origin | spect |
| Original Points Count | 32768 | Owner | nmr | Points Count | 32768 | Pulse Sequence | zgpg30 |
| Receiver Gain | 186.93 | SW(cyclical) (Hz) | 51020.41 | Solvent | DMSO-d6 | Spectrum Offset (Hz) | 21290.6973 |
| Spectrum Type | STANDARD | Sweep Width (Hz) | 51018.85 | Temperature (degree C) | 24.999 | | |

$^{13}$C NMR (214 MHz, DMSO-$d_6$) δ 164.7, 164.1, 146.4, 141.7, 139.1, 134.4, 133.7, 133.7, 130.2, 129.5, 129.2, 123.7, 123.1, 122.1, 41.1, 39.8, 39.7, 39.6, 39.4, 39.3, 39.2, 22.3, 11.5

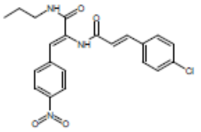


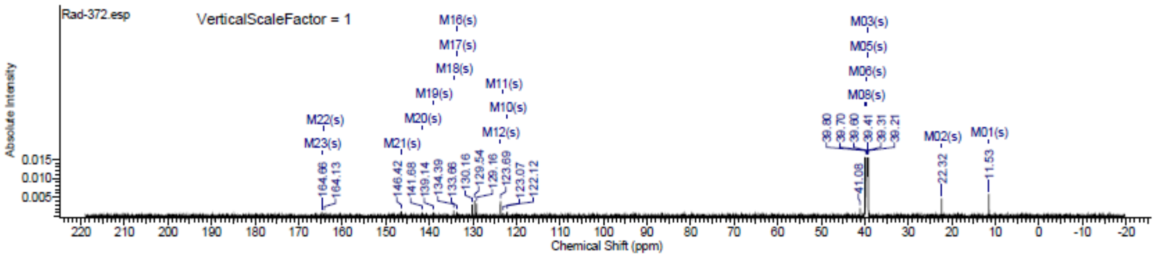

#### Spectra of (4912)

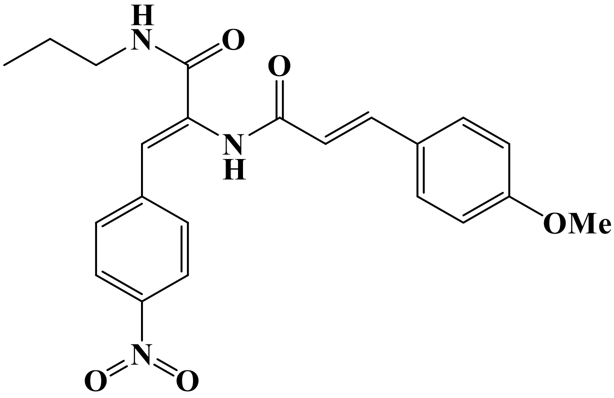


**FT-IR**

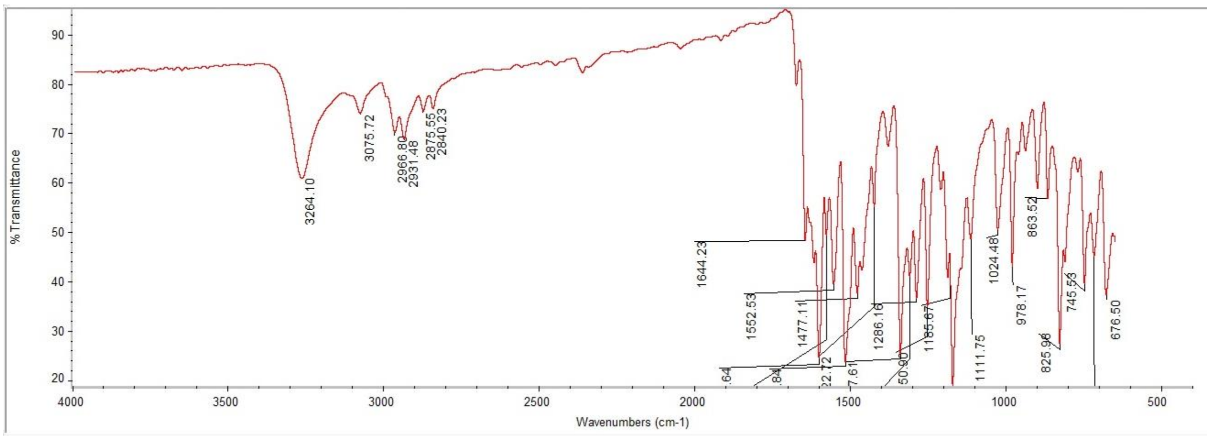

#### $^{1}$H NMR

This report was created by ACD/NMR Processor Academic Edition. For more information go to www.acdlabs.com/nmrproc/

##### RAD-373

09/11/2016 9:23:51 AM
Dr.Mustafa Sample : RAD-373 DMSO PROTON DMSO {D:\Magdy} nmr 25

| Formula $C_{22}H_{23}N_3O_5$ | FW 409.4351 |
|---|---|

| | | | | | | | |
|---|---|---|---|---|---|---|---|
| *Acquisition Time (sec)* | 2.6564 | *Comment* | Dr.Mustafa Sample : RAD-373 DMSO PROTON DMSO {D:\Magdy} nmr 25 | | | | |
| *Date* | 18 Jun 2015 15:43:28 | | | *Date Stamp* | 18 Jun 2015 15:43:28 | | |
| *File Name* | E:\Mostafa Alaraby Project\Oxazolone NMR\NMR final oxazolone\MUSTAFA RAD-373 18-06-2015\10\fid | | | | | *Frequency (MHz)* | 600.15 |
| *Nucleus* | 1H | *Number of Transients* | 32 | *Origin* | spect | *Original Points Count* | 32768 |
| *Owner* | nmr | *Points Count* | 32768 | *Pulse Sequence* | zg30 | *Receiver Gain* | 128.00 |
| *SW(cyclical) (Hz)* | 12335.53 | *Solvent* | DMSO-d6 | *Spectrum Offset (Hz)* | 3706.1750 | *Spectrum Type* | STANDARD |
| *Sweep Width (Hz)* | 12335.15 | *Temperature (degree C)* | 25.000 | | | | |

$^{1}$H NMR (600 MHz, DMSO-$d_6$) δ 9.84 (br. s., 1H), 8.31 (t, *J* = 5.65 Hz, 1H), 8.20 - 8.26 (m, *J* = 9.04 Hz, 2H), 7.73 - 7.79 (m, *J* = 8.66 Hz, 2H), 7.56 - 7.58 (m, *J* = 8.66 Hz, 2H), 7.46 (d, *J* = 15.81 Hz, 1H), 7.01 - 7.03 (m, *J* = 8.66 Hz, 2H), 6.93 (s, 1H), 6.71 (d, *J* = 15.81 Hz, 1H), 3.81 (s, 3H), 3.10 - 3.16 (m, 2H), 1.50 (sxt, *J* = 7.23 Hz, 2H), 0.89 (t, *J* = 7.53 Hz, 3H)

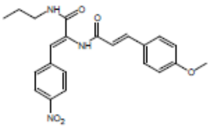

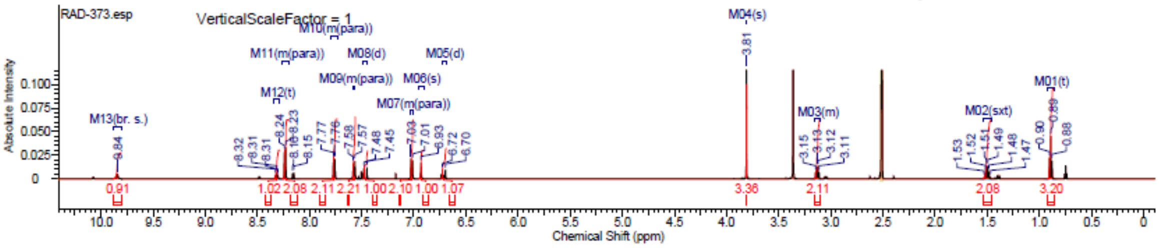


#### $^{13}$C NMR

This report was created by ACD/NMR Processor Academic Edition. For more information go to www.acdlabs.com/nmrproc/

##### RAD-373

20/09/2016 10:06:57 AM
Dr.Mostafa Sample : RAD-373 DMSO

| Formula $C_{22}H_{23}N_3O_5$ | FW 409.4351 |
|---|---|

| | | | | | | | |
|---|---|---|---|---|---|---|---|
| *Acquisition Time (sec)* | 0.6423 | *Comment* | Dr.Mostafa Sample : RAD-373 DMSO | | | *Date* | 19 Apr 2016 01:11:12 |
| *Date Stamp* | 19 Apr 2016 01:11:12 | | | | | | |
| *File Name* | E:\Projects\Mostafa Alaraby Project\Oxazolone\Oxazolone NMR\NMR final oxazolone\13 CNMR New 18-4-2016\MUSTAFA RAD-373 18-04-2016\60\fid | | | | | | |
| *Frequency (MHz)* | 213.77 | *Nucleus* | 13C | *Number of Transients* | 3072 | *Origin* | spect |
| *Original Points Count* | 32768 | *Owner* | nmr | *Points Count* | 32768 | *Pulse Sequence* | zgpg30 |
| *Receiver Gain* | 186.93 | *SW(cyclical) (Hz)* | 51020.41 | *Solvent* | DMSO-d6 | *Spectrum Offset (Hz)* | 21287.5840 |
| *Spectrum Type* | STANDARD | *Sweep Width (Hz)* | 51018.85 | *Temperature (degree C)* | 24.999 | | |

$^{13}$C NMR (214 MHz, DMSO-$d_6$) δ 164.8, 164.6, 160.7, 146.3, 141.8, 140.3, 134.0, 130.1, 129.5, 127.2, 123.7, 122.6, 118.7, 114.5, 55.4, 41.1, 39.8, 39.7, 39.6, 39.4, 39.3, 39.2, 22.3, 11.5

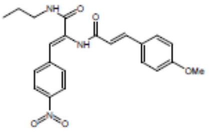

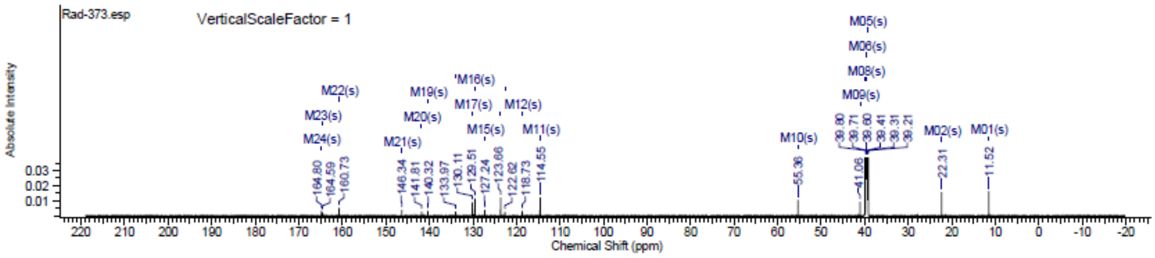

### Spectra of (5012)

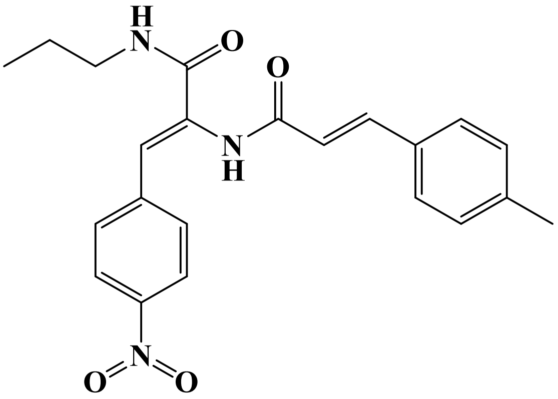


## LC/MS

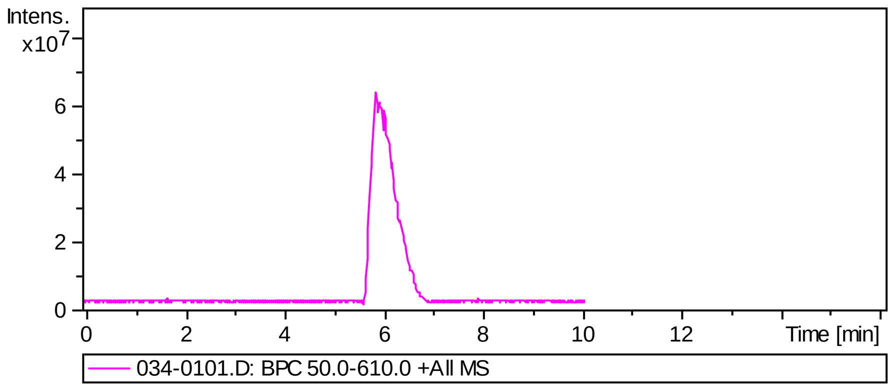


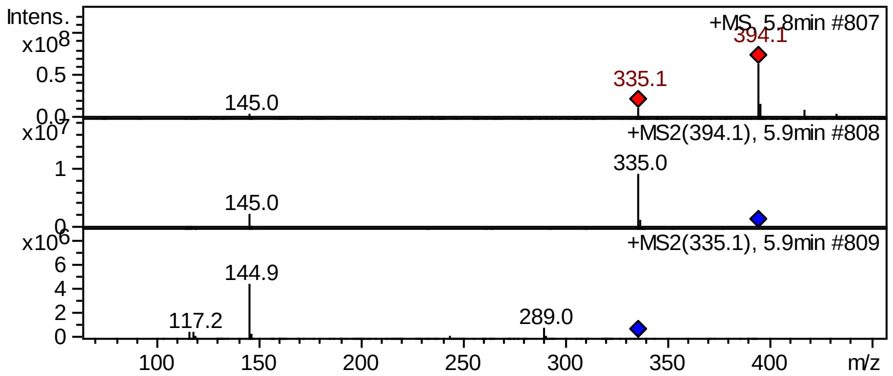


## FT-IR

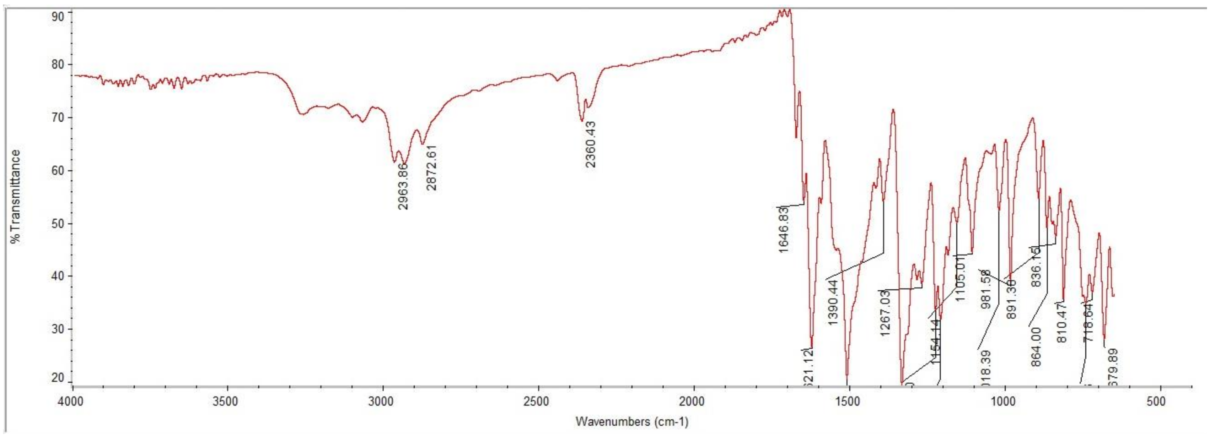

#### $^{1}$H NMR

This report was created by ACD/NMR Processor Academic Edition. For more information go to www.acdlabs.com/nmrproc/

##### RAD-374

09/11/2016 9:46:47 AM
Dr.Mustafa El-Araby Sample : RAD-314 DMSO

| Formula $C_{22}H_{24}N_2O_2$ | FW 348.4382 |
|---|---|

| | | | | | | | |
|---|---|---|---|---|---|---|---|
| Acquisition Time (sec) | 1.9268 | Comment | Dr.Mustafa El-Araby Sample : RAD-314 DMSO | | | Date | 08 Apr 2015 10:23:44 |
| Date Stamp | 08 Apr 2015 10:23:44 | | | | | | |
| File Name | E:\Mostafa Alaraby Project\Oxazolone NMR\NMR final oxazolone\MUSTAFA RAD-314 08-04-2015\40\fid | | | | | Frequency (MHz) | 850.15 |
| Nucleus | 1H | Number of Transients | 20 | Origin | spect | Original Points Count | 32768 |
| Owner | nmr | Points Count | 32768 | Pulse Sequence | zg30 | Receiver Gain | 9.04 |
| SW(cyclical) (Hz) | 17006.80 | Solvent | DMSO-d6 | Spectrum Offset (Hz) | 5250.0283 | Spectrum Type | STANDARD |
| Sweep Width (Hz) | 17006.28 | Temperature (degree C) | 25.000 | | | | |

$^{1}$H NMR (850 MHz, DMSO-$d_6$) δ 9.65 (br. s., 1H), 8.11 (t, *J* = 5.45 Hz, 1H), 7.48 - 7.57 (m, 4H), 7.46 (d, *J* = 15.57 Hz, 1H), 7.37 - 7.40 (m, 2H), 7.27 - 7.34 (m, 2H), 7.26 (d, *J* = 7.78 Hz, 1H), 6.98 - 7.01 (m, 1H), 6.82 (d, *J* = 15.57 Hz, 1H), 3.09 - 3.13 (m, 2H), 1.48 (td, *J* = 7.20, 14.14 Hz, 2H), 0.87 (t, *J* = 7.27 Hz, 3H)

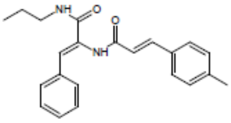


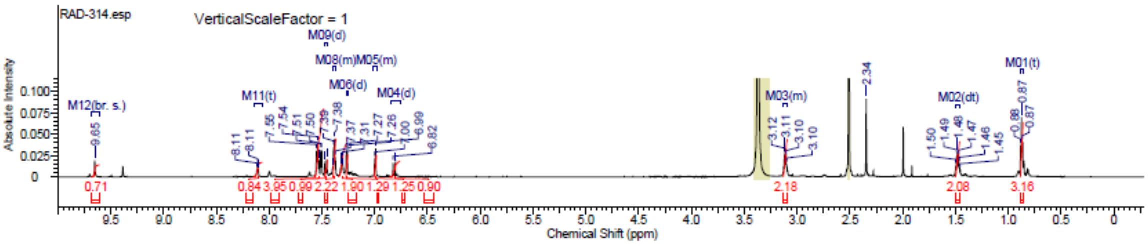


#### $^{13}$C NMR

This report was created by ACD/NMR Processor Academic Edition. For more information go to www.acdlabs.com/nmrproc/

##### RAD-374

20/09/2016 10:09:40 AM
Dr.Mostafa Sample ; RAD-374 DMSO

| Formula $C_{22}H_{23}N_3O_4$ | FW 393.4357 |
|---|---|

| | | | | | | | |
|---|---|---|---|---|---|---|---|
| Acquisition Time (sec) | 0.6423 | Comment | Dr.Mostafa Sample ; RAD-374 DMSO | | | Date | 20 Apr 2016 14:09:52 |
| Date Stamp | 20 Apr 2016 14:09:52 | | | | | | |
| File Name | E:\Projects\Mostafa Alaraby Project\Oxazolone\Oxazolone NMR\NMR final oxazolone\13 CNMR New 18-4-2016\MUSTAFA RAD-374 19-04-2016\120\fid | | | | | | |
| Frequency (MHz) | 213.77 | Nucleus | 13C | Number of Transients | 963 | Origin | spect |
| Original Points Count | 32768 | Owner | nmr | Points Count | 32768 | Pulse Sequence | zgpg30 |
| Receiver Gain | 186.93 | SW(cyclical) (Hz) | 51020.41 | Solvent | DMSO-d6 | Spectrum Offset (Hz) | 21292.2539 |
| Spectrum Type | STANDARD | Sweep Width (Hz) | 51018.85 | Temperature (degree C) | 25.001 | | |

$^{13}$C NMR (214 MHz, DMSO-$d_6$) δ 164.8, 164.5, 146.4, 141.8, 140.5, 139.9, 133.9, 131.9, 130.1, 129.7, 127.9, 123.7, 122.8, 120.3, 41.1, 39.8, 39.7, 39.6, 39.4, 39.3, 39.2, 22.3, 21.1, 11.5

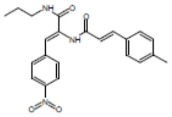


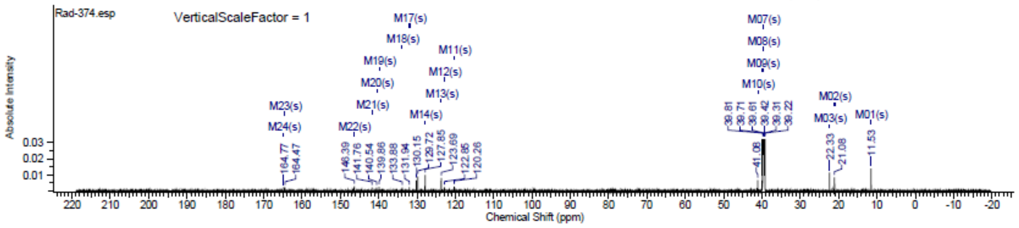

### Spectra of (5112)

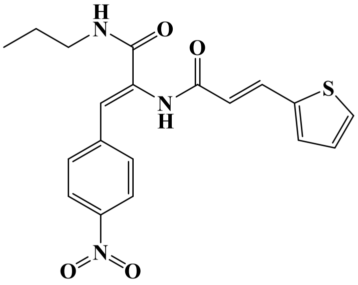


## LC/MS

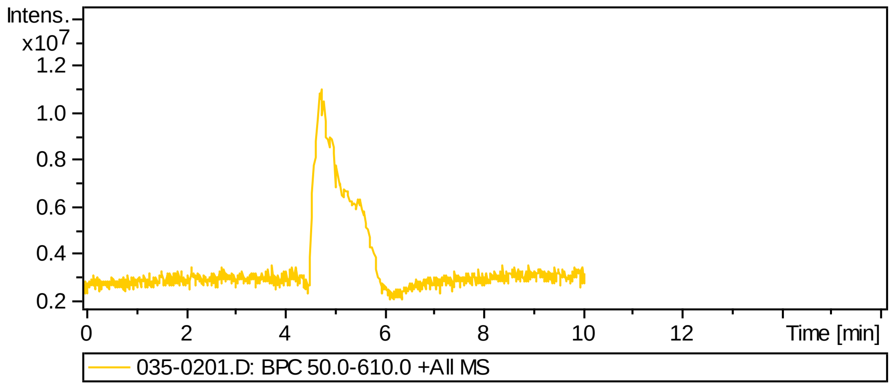


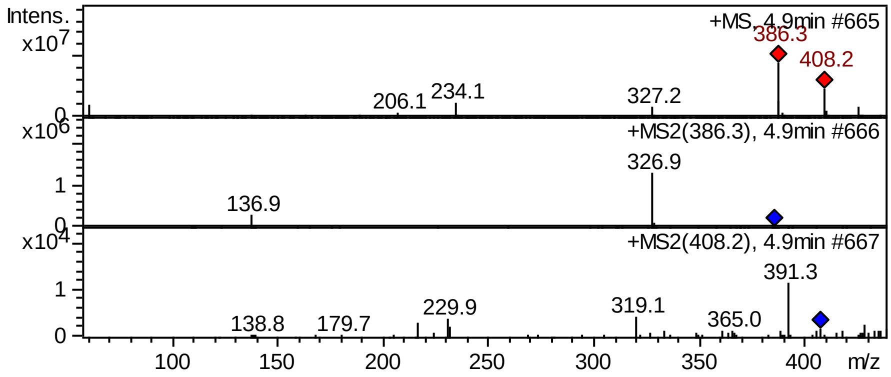


## FT-IR

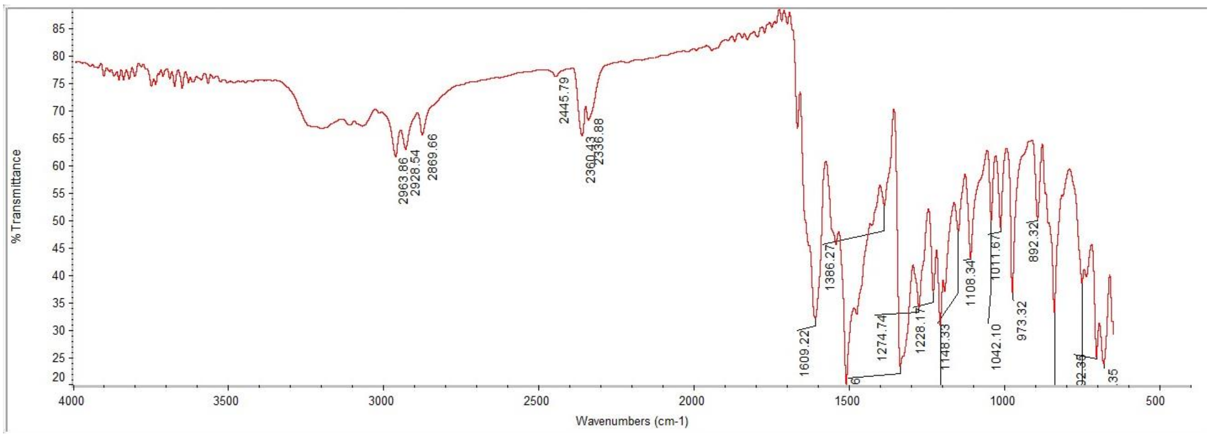

#### $^{1}$H NMR

This report was created by ACD/NMR Processor Academic Edition. For more information go to www.acdlabs.com/nmrproc/

**RAD-375**

09/11/2016 8:47:48 AM
Dr.Mustafa Sample : RAD-375 DMSO PROTON DMSO {D:\Magdy} nmr 24

| Formula $C_{19}H_{19}N_3O_4S$ | FW 385.4369 |
|---|---|

| | | | | | | | |
|---|---|---|---|---|---|---|---|
| Acquisition Time (sec) | 2.6564 | Comment | Dr.Mustafa Sample : RAD-375 DMSO PROTON DMSO {D:\Magdy} nmr 24 | | | | |
| Date | 18 Jun 2015 15:37:04 | | | Date Stamp | 18 Jun 2015 15:37:04 | | |
| File Name | E:\Mostafa Alaraby Project\Oxazolone NMR\NMR final oxazolone\MUSTAFA RAD-375 18-06-2015\10\fid | | | | | Frequency (MHz) | 600.15 |
| Nucleus | 1H | Number of Transients | 32 | Origin | spect | Original Points Count | 32768 |
| Owner | nmr | Points Count | 32768 | Pulse Sequence | zg30 | Receiver Gain | 144.00 |
| SW(cyclical) (Hz) | 12335.53 | Solvent | DMSO-d6 | Spectrum Offset (Hz) | 3706.1750 | Spectrum Type | STANDARD |
| Sweep Width (Hz) | 12335.15 | Temperature (degree C) | 25.000 | | | | |

$^{1}$H NMR (600 MHz, DMSO-$d_6$) δ 9.89 (s, 1H), 8.32 (t, $J$ = 5.65 Hz, 1H), 8.22 - 8.25 (m, $J$ = 9.04 Hz, 2H), 7.74 - 7.78 (m, $J$ = 9.03 Hz, 2H), 7.64 - 7.69 (m, 2H), 7.45 (d, $J$ = 3.39 Hz, 1H), 7.15 (dd, $J$ = 3.58, 5.08 Hz, 1H), 6.95 (s, 1H), 6.60 (d, $J$ = 15.43 Hz, 1H), 3.12 (q, $J$ = 6.40 Hz, 2H), 1.50 (sxt, $J$ = 7.30 Hz, 2H), 0.89 (t, $J$ = 7.53 Hz, 3H)

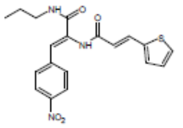


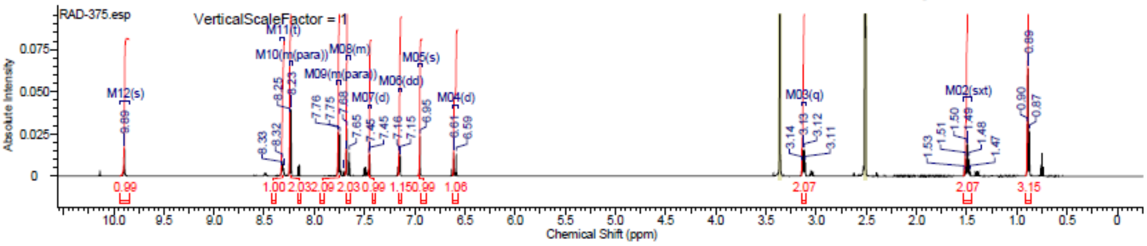


#### $^{13}$C NMR

This report was created by ACD/NMR Processor Academic Edition. For more information go to www.acdlabs.com/nmrproc/

**RAD-375**

20/09/2016 10:12:35 AM
Dr.Mostafa Sample : RAD-375 DMSO

| Formula $C_{19}H_{19}N_3O_4S$ | FW 385.4369 |
|---|---|

| | | | | | | | |
|---|---|---|---|---|---|---|---|
| Acquisition Time (sec) | 0.6423 | Comment | Dr.Mostafa Sample : RAD-375 DMSO | | | Date | 18 Apr 2016 22:48:16 |
| Date Stamp | 18 Apr 2016 22:48:16 | | | | | | |
| File Name | E:\Projects\Mostafa Alaraby Project\Oxazolone\Oxazolone NMR\NMR final oxazolone\13 CNMR New 18-4-2016\MUSTAFA RAD-375 18-04-2016\50\fid | | | | | | |
| Frequency (MHz) | 213.77 | Nucleus | 13C | Number of Transients | 3072 | Origin | spect |
| Original Points Count | 32768 | Owner | nmr | Points Count | 32768 | Pulse Sequence | zgpg30 |
| Receiver Gain | 186.93 | SW(cyclical) (Hz) | 51020.41 | Solvent | DMSO-d6 | Spectrum Offset (Hz) | 21293.8125 |
| Spectrum Type | STANDARD | Sweep Width (Hz) | 51018.85 | Temperature (degree C) | 24.998 | | |

$^{13}$C NMR (214 MHz, DMSO-$d_6$) δ 164.7, 164.1, 146.4, 141.7, 139.7, 133.7, 133.6, 131.6, 130.1, 128.7, 128.6, 123.7, 123.0, 120.0, 41.1, 39.8, 39.7, 39.6, 39.4, 39.3, 39.2, 22.3, 11.5

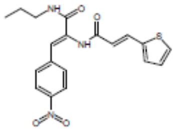


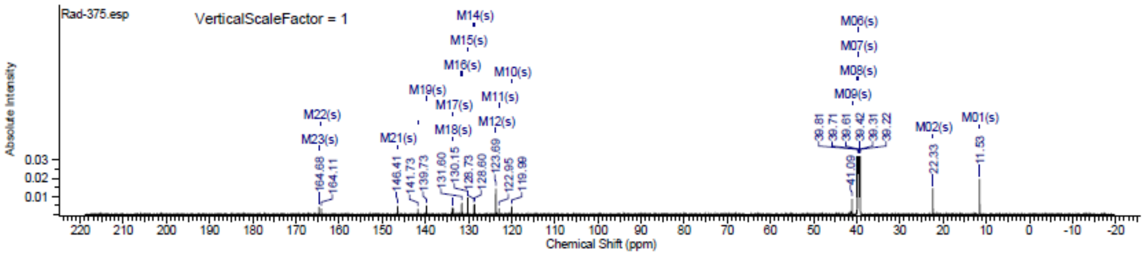

### II. Further Data on LD50 determination

**Chart of the OECD protocol, Guideline 423**
**OECD Guideline for testing of chemicals**
**Acute Oral Toxicity – Acute Toxic Class Method**
**https://ntp.niehs.nih.gov/iccvam/suppdocs/feddocs/oecd/oecd_gl423.pdf**

OECD/OCDE 423

ANNEX 2d: TEST PROCEDURE WITH A STARTING DOSE OF 2000 MG/KG BODY WEIGHT

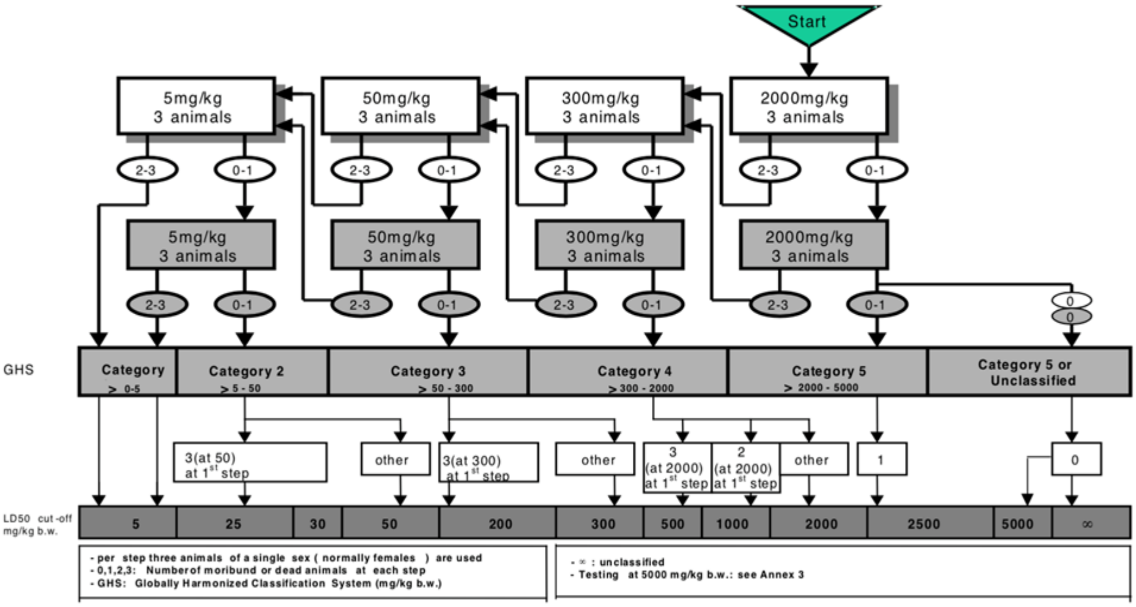
